# Conflict between heterozygote advantage and hybrid incompatibility in haplodiploids (and sex chromosomes)

**DOI:** 10.1101/196469

**Authors:** Ana-Hermina Ghenu, Alexandre Blanckaert, Roger K. Butlin, Jonna Kulmuni, Claudia Bank

## Abstract

In many diploid species the sex chromosomes play a special role in mediating reproductive isolation. In haplodiploids, where females are diploid and males haploid, the whole genome behaves similarly to the X/Z chromosomes of diploids. Therefore, haplodiploid systems can serve as a model for the role of sex chromosomes in speciation and hybridization. A previously described population of Finnish *Formica* wood ants displays genome-wide signs of ploidally and sexually antagonistic selection resulting from hybridization. Here, hybrid females have increased survivorship but hybrid males are inviable. To understand how the unusual hybrid population may be maintained, we developed a mathematical model with hybrid incompatibility, female heterozygote advantage, recombination, and assortative mating. The rugged fitness landscape resulting from the co-occurrence of heterozygote advantage and hybrid incompatibility results in a sexual conflict in haplodiploids, which is caused by the ploidy difference. Thus, whereas heterozygote advantage always promotes long-term polymorphism in diploids, we find various outcomes in haplodiploids in which the population stabilizes either in favor of males, females, or via maximizing the number of introgressed individuals. We discuss these outcomes with respect to the potential long-term fate of the Finnish wood ant population, and provide approximations for the extension of the model to multiple incompatibilities. Moreover, we highlight the general implications of our results for speciation and hybridization in haplodiploids versus diploids, and how the described fitness relationships could contribute to the outstanding role of sex chromosomes as hotspots of sexual antagonism and genes involved in speciation.

## Introduction

Haplodiploids are an emerging system for speciation genetics (Koevoets and Beukeboom, 2009; Kulmuni and Pamilo, 2014; Lohse and Ross, 2015; Knegt et al., 2017). Although ≈ 20% of animal species are haplodiploid (comprising most *Hymenopterans*, some arthropods, thrips and *Hemipterans*, and several clades of beetles and mites; Crozier and Pamilo, 1996; Evans et al., 2004; de la Filia et al., 2015), little evolutionary theory has been developed specifically for speciation in haplodiploids (Koevoets and Beukeboom, 2009). Under haplodiploidy with arrhenotoky (hereafter simply haplodiploidy; Suomalainen et al., 1987), males develop from the mother’s unfertilized eggs and are haploid, whereas eggs fertilized by fathers result in diploid females. Since this mode of inheritance is, from a theoretical viewpoint, similar to that of the X/Z chromosome, most work on speciation of haplodiploids draws on the rich literature of sex chromosome evolution (Jablonka and Lamb, 1991; Presgraves, 2008; Johnson and Lachance, 2012; Lohse and Ross, 2015). An important similarity between haplodiploids and X/Z chromosomes is that recessive mutations in the haploid sex are exposed to selection, but they are masked in diploids. This is expected to lead to faster evolution in the sex chromosomes (Charlesworth et al., 1987) that may partly underlie the large-X effect (Presgraves, 2008). The large-X effect refers to the observation that the sex chromosomes seem to play a special role in speciation by acting as the strongest barrier for gene flow between hybridizing lineages across different species (Höllinger and Hermisson, 2017). Similarly, haplodiploid species have been suggested to acquire reproductive isolation earlier and speciate faster than diploid species (Lohse and Ross, 2015; Lima, 2014). Although the factors influencing haplodiploid and X/Z chromosome evolution are not expected to be exactly the same (e.g. movement of sexually antagonistic genes to the sex chromosomes, dosage compensation between the sex chromosomes and autosomes, and turnover of sex chromosomes cannot occur in haplodiploids; Abbott et al., 2017), by studying haplodiploid models we can both improve our understanding of how speciation happens in the large subgroup of the animal kingdom that is haplodiploid, and gain new insights into the role of X/Z chromosomes in speciation for diploid species.

Recent studies have shown that hybridization and resulting gene flow between diverging populations may be important players in the speciation process since signs of hybridization and introgression are being observed ubiquitously in natural populations (Mallet, 2005; Dieckmann and Doebeli, 1999; Schluter, 2009; Schluter and Conte, 2009; Seehausen et al., 2014). When a hybrid population is formed, various selective forces may act simultaneously to either increase or decrease hybrid fitness, which dictate the fate of the population. One commonly documented finding is hybrid incompatibility (Presgraves, 2008; Fraïsse et al., 2014; Chen et al., 2016), where combinations of alleles at different loci interact to confer poor fitness when combined in a hybrid individual (Bateson, 1909; Dobzhansky, 1936; Muller, 1942; Orr, 1995). In a hybrid population, the existence of hybrid incompatibility reduces the mean population fitness. This deficit can be resolved either through reinforcement (evolution of increased premating isolation to avoid production of unfit hybrids; Servedio and Noor, 2003) or by purging (demographic swamping leading to extinction of one of the local populations/species or reinstatement of the ancestral allele combinations; Wolf et al., 2001). On the other hand, hybridization can transfer adaptive genetic variation from one lineage to another (Heliconius Genome Consortium, 2012; Song et al., 2011; Whitney et al., 2010) and may result in overall heterosis (also known as hybrid vigor): a higher fitness of hybrids as compared to their parents (Schwarz et al., 2005; Chen, 2013; Bernardes et al., 2017). Heterosis can stabilize polymorphisms by conferring a fitness advantage to hybrids and thereby favor the maintenance of hybridization either through the improved exploitation of novel ecological niches or the masking of recessive deleterious mutations. Therefore hybrid incompatibility acts to avert ongoing hybridization while heterosis favors the maintenance of hybrids.

One example of the simultaneous action of hybridization-averse and hybridization-favoring forces is found in a hybrid population of *Formica polyctena* and *F. aquilonia* wood ants in Finland (Kulmuni et al., 2010; Kulmuni and Pamilo, 2014; Beresford et al., 2017). Here, it has been reported that hybrid (haploid) males do not survive to adulthood, whereas (diploid) females have higher survivorship when they carry many introgressed alleles as heterozygotes (i.e., heterozygous for alleles originating from one of the parental species in a genomic background otherwise from the other parental species). Thus, a combination of hybrid incompatibility and heterosis seems to dictate the dynamics of the population in a ploidy-specific manner: hybrid haploid males suffer a fitness cost while diploid hybrid females can have a selective advantage over parental ones. Here, the differences in ploidy create an apparent sexual conflict (sensu Arnqvist and Rowe, 2005) between haploid males and diploid females, because their fitness landscapes (i.e., the complex relationship between genotypes and fitness created via hybrid incompatibility and heterozygote advantage) are different. This conflict is absent if the same rugged fitness landscape occurs in diploid autosomes.

When both hybridization-averse and hybridization-favoring forces are acting, the longterm resolution of a hybridizing population is difficult to foresee: will hybridization eventually result in either complete speciation or extinction of one of the populations involved? Alternatively, can it represent an equilibrium maintained stably on an evolutionary time scale? Furthermore, will the probability of these outcomes depend on ploidy? In other words, is one of these outcomes more probable when interacting genes are found on a “haplodiploid” X/Z chromosome than when they exist on a “diploid” autosome?

We here develop and analyze a population-genetic model of an isolated hybrid population in which both hybridization-averse and hybridization-favoring forces are acting, and we study the evolutionary outcomes in both haplodiploid and (fully) diploid genetic systems. The rich dynamics of the haplodiploid model can result in four possible evolutionary stable states depending on the strength of heterozygote advantage *versus* hybrid incompatibility, the strength of recombination, and the degree of assortative mating. This includes a case of symmetric coexistence (where all diversity is maintained) in which both alleles can be maintained despite the segregating hybrid incompatibility, and in which long-term hybridization is favored. We find that the dynamics differ between haplodiploid and diploid systems and that, unlike in previous models of sexual conflict in haplodiploid populations (Kraaijeveld, 2009; Albert and Otto, 2005), the conflict is not necessarily resolved in favor of the females. Indeed, a compromise may be reached at which the average fitness of females is decreased to rescue part of the fitness of males. Moreover, evaluation of the model using the data from the natural hybrid population suggests that, under the assumption of an equilibrium, the Finnish ant population may represent an example of compromise between male costs and female benefits through asymmetric coexistence. We discuss our findings with respect to the long-term effects of hybridization, the potential for speciation in haplodiploid versus diploid species, and with respect to their relevance for X- or Z-linked alleles in diploid individuals.

## Materials and Methods

### The model

We model an isolated haplodiploid or diploid hybrid population with individuals from two founder populations *P*_+_ and *P*_−_. Note that throughout the manuscript, we preferentially refer to (sub-)populations rather than species; in those instances in which we use the term ‘species’ it is in order to emphasize that the two populations have diverged sufficiently for (potentially strong) hybrid incompatibility to exist. We assume discrete generations and consider two loci, **A** and **B**. Each locus has two alleles, the ‘+’ allele (*A*_+_ or *B*_+_) inherited from population *P*_+_ and the ‘−’ allele (*A*_−_ or *B*_−_) inherited from population *P*_−_. We refer to ‘hybrids’ as individuals that carry two alleles from each of the two parental populations and cannot be assigned to either parental background. We refer to ‘introgressed’ individuals as those genotypes for which three of the four alleles are from the same parental population; these genotypes are identical to those produced by hybridization followed by backcrossing. We ignore new or recurrent mutation and genetic drift. Thus, male and female populations are of effectively infinite size; selection modifies the relative abundance of the different haplotypes/genotypes but not the number of individuals (soft selection). The life cycle is as follows (Fig. 1; see also Table 1 for a list of model parameters); consistent with the recursions defined below, we begin the life cycle at the adult stage:

1. mating, either randomly or via genotype matching with assortment strength *α* as detailed below;
2. recombination (in diploid individuals) at rate *ρ*;
3. viability (or survival) selection, where heterosis is modeled as a heterozygote advantage, *σ*, and hybrid incompatibility is modeled as a fully recessive negative epistasis, *γ*_1_ and *γ*_2_ (further details are provided below and in Figure 2).

**Figure 1:**
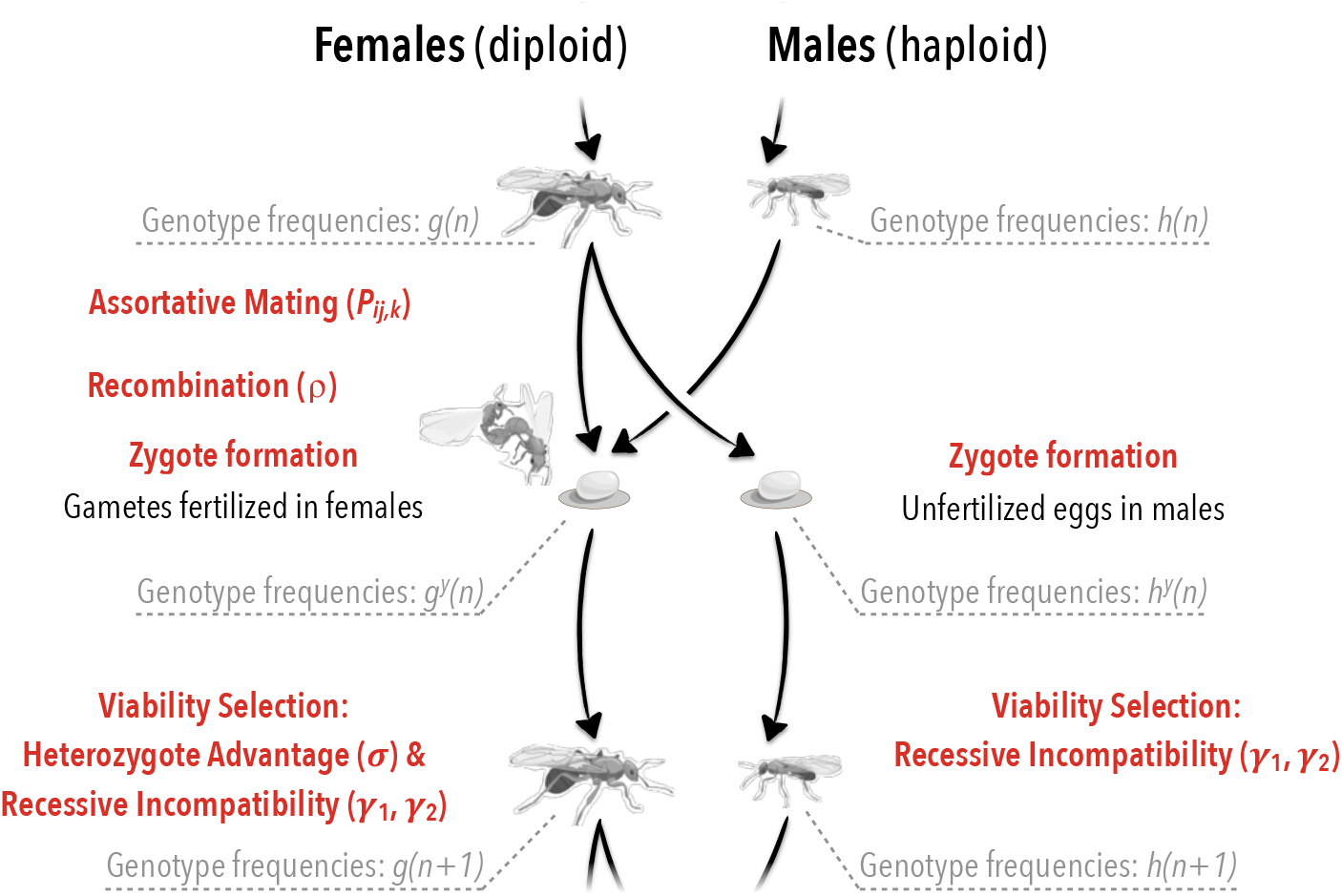
Illustration of the haplodiploid life cycle and its parametization

**Table 1:**
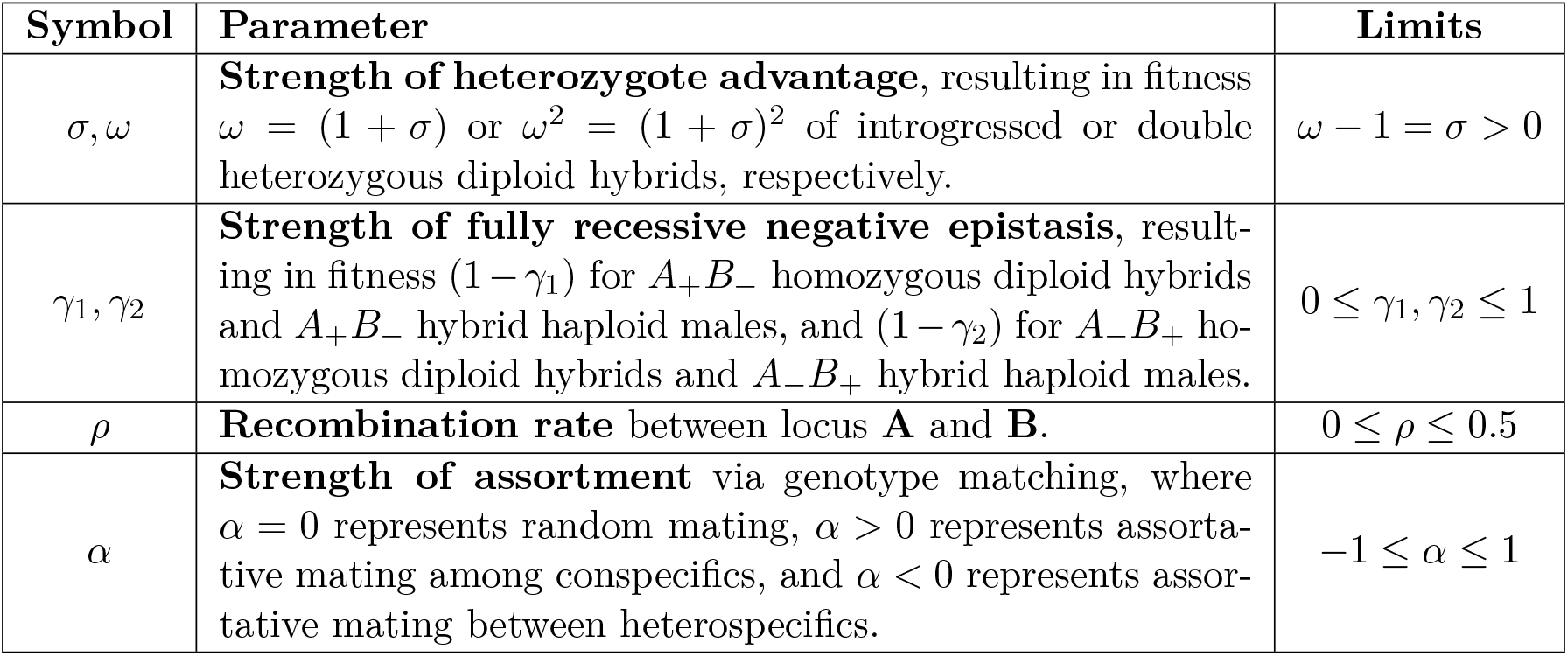
List of model parameters.

### Viability selection

The fitness landscape described here (Fig. 2) is inspired by the situation observed in Finnish *Formica* ants (Kulmuni et al., 2010; Kulmuni and Pamilo, 2014; Beresford et al., 2017). There, the authors discovered heterosis in the diploid females but recessive incompatibilities expressed in the haploid males. This creates a situation in which the same alleles that are favored in heterozygous females are selected against in hybrid haploid males and homozygous hybrid females. In the haplodiploid genetic system, males possess only one copy of each locus so they cannot be heterozygous and, therefore, cannot experience heterozygote advantage (Fig. 2(b)). Therefore, the fitness landscape with heterozygote advantage and recessive hybrid incompatibility expresses itself as an apparent sexual conflict when sexes differ in ploidy, as in haplodiploids or for X/Z chromosomes.

**Figure 2:**
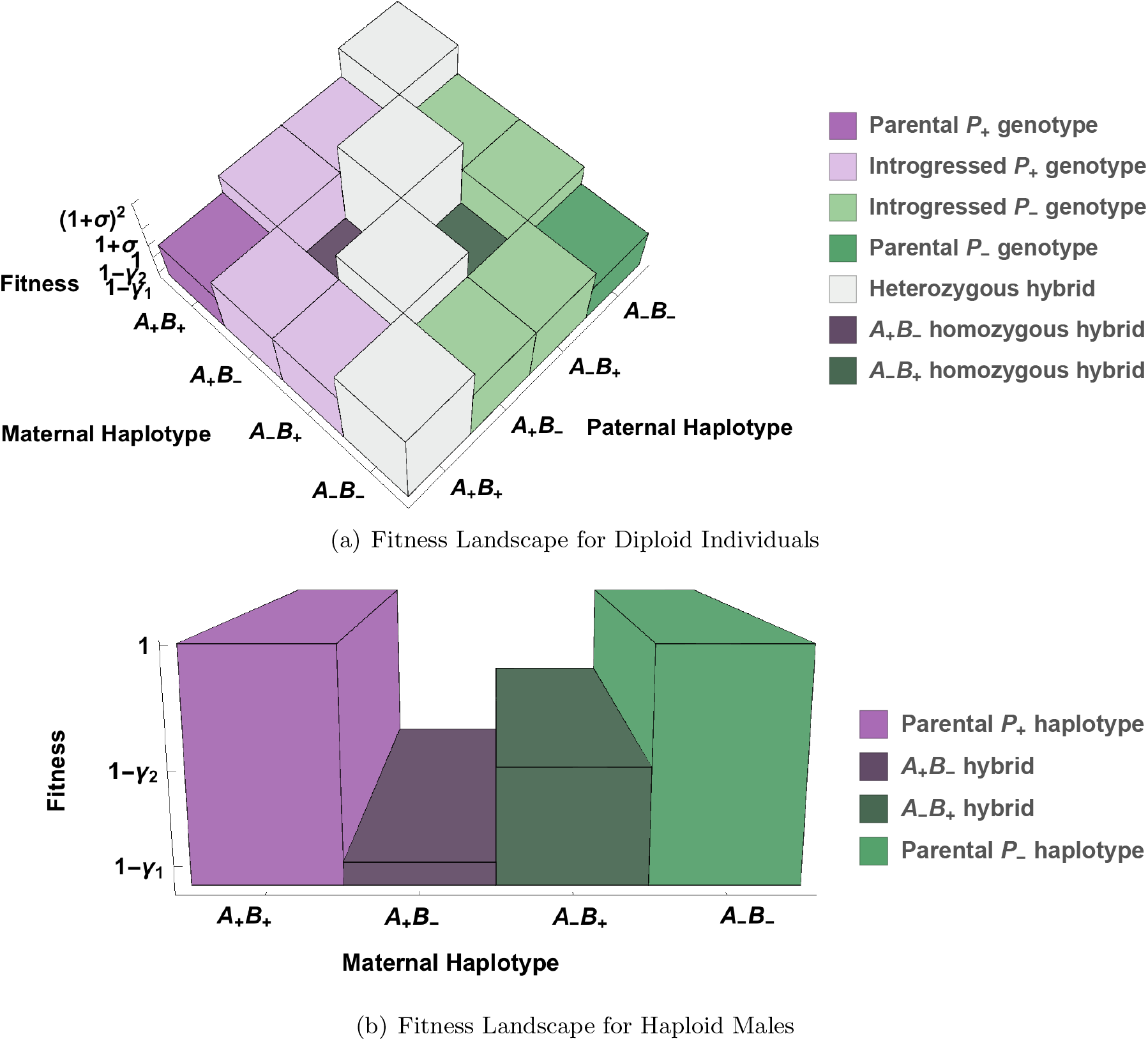
Three-dimensional fitness landscapes for the (a) diploid and (b) haploid genotypes. Panel a) corresponds to females in the haplodiploid model and all individuals in the diploid model. Individuals heterozygous at both loci (heterozygous hybrids) reside on a high fitness ridge (in white), whereas individuals homozygous at both loci (homozygous hybrids) suffer from reduced fitness due to negative epistasis. Panel b) shows the fitness landscape for haploid individuals (i.e. males) in the haplodiploid model. This landscape is identical to a transect from Panel a) for genotypes homozygous at both loci.

In our model, selection for heterozygous individuals is multiplicative with respect to the number of heterozygous loci: introgressed individuals with one heterozygous locus have fitness 1 + *σ*, whereas diploid hybrid individuals that are heterozygous at both loci have survivorship (1 + *σ*)^2^ (Fig. 2(a)). Finally, the recessive epistatic incompatibility parameter *γ*_1_ acts on individuals homozygous or haploid for the *A*_+_*B*_−_ haplotype, and *γ*_2_ acts on individuals homozygous or haploid for the *A*_−_*B*_+_ haplotype (without loss of generality, we assume *γ*_1_ ≥ *γ*_2_). Thus, epistasis in this model can be asymmetric, reflecting, for example, two Dobzhansky-Muller incompatibilities of different strength that have accumulated at a negligible recombination distance between the same chromosome pairs. Note that when *γ*_1_ = *γ*_2_ = 1, haploid hybrid males and homozygous hybrid zygotes are produced but do not survive to adulthood and that the classical case of a single Dobzhansky-Muller incompatibility is recovered when *γ*_2_ = 0.

### Assortative mating

Prezygotic isolation via assortative mating is an important mechanism that could mediate the detrimental effects to the population caused by the co-occurence of heterozygote advantage and epistasis modeled here. In the Finnish wood ant population that inspired our model (Kulmuni and Pamilo, 2014), almost all egg-laying queens collected had been inseminated by males of the same genetic group, indicating that prezygotic isolation barriers are likely operating to result in assortative mating. In this case, assortative mating could arise via choosiness of mating partners, via genotype-dependent development times, or via other post-mating prezygotic mechanisms. We implemented assortment via genotype matching (reviewed in Kopp et al. (2017)), where the proportion of matings depends on the genetic distance between two mating partners (and their respective frequencies in the population). We define the genetic distance between the genotypes of a mating pair as the average Hamming distance, i.e. the number of differences between 2 aligned sequences of characters, between all possible pairs of haplotypes with one parter from each sex. We use quadratic assortment (e.g., De Cara et al., 2008), which results in assortative mating without costs of choosiness but with sexual selection. The mating probability of a pair of male and female genotypes, *P_ij,k_* depends on the genetic distance between the two mates, the choosiness of the female, and the abundance of the different haplotype and genotypes as detailed below.

### Mathematical modeling and analysis

In a given generation *n*, the frequencies of the male and female adults are given by *h_k_*(*n*) and *g_ij_*(*n*), respectively, with *i* and *k* indicating the haplotype received maternally and *j* the one of paternal origin. Without loss of generality, we assign index *i* = 1 to haplotype *A*_+_*B*_+_, index *i* = 2 to haplotype *A*_+_*B*_−_, *i* = 3 to haplotype *A*_−_*B*_+_ and, *i* = 4 to *A*_−_*B*_−_. Below, we describe the modeled life cycle (illustrated in Fig. S1) which determines how frequencies change from one generation to the next.

1. As detailed in figure 1 the first step of the life cycle is the mating between two individuals. The mating probability between an *ij* female and a *k* male is given by:

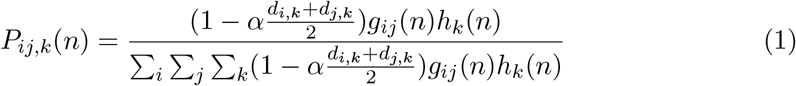

with *d_i,k_* the Hamming distance between two haplotypes. Note that for *α* = 0, this simplifies to random mating and thus becomes equivalent to the dynamics described in Supplementary material (S7).
2. The next step is the formation of the zygote. Recombination happens only in females. We denote the frequency of newly born females as 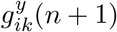.

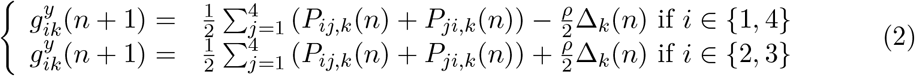

with Δ_*k*_(*n*) = *P*_14,*k*_(*n*) + *P*_41,*k*_(*n*) – *P*_23,*k*_(*n*) – *P*_32,*k*_(*n*). Males are composed from unfertilized female gametes, which have undergone recombination. The frequencies of newborn males are given by 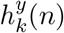:

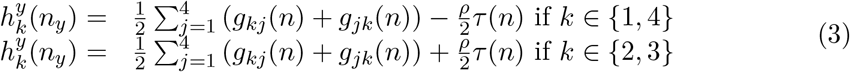

with *τ*(*n*) = *g*_14_(*n*) + *g*_41_(*n*) – *g*_23_(*n*) – *g*_32_(*n*).
3. Individuals of both sexes are under viability selection. The frequencies of male and female adults of the next generations are given by

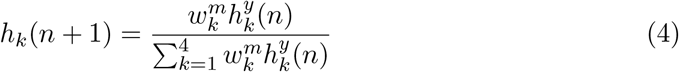

with 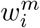 the fitness of haplotype *i* in males and:

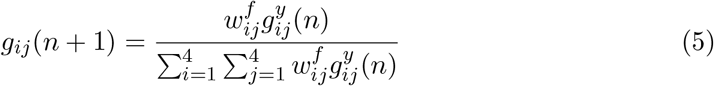

where 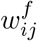 denotes the fitness of the *ij* genotype. Note that there are no parental effects: 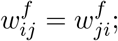 we maintain the distinction only for modeling convenience.

The complete recursion for females is obtained by substituting 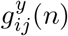 by its expression given in (2) in (5) and *P_ij,k_*(*n*) by (1). The complete recursion for males is given by substituting 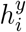 by its expression given in (3) in (4). For *α* = 0, the detailed recursion is given in Supplement (S7). Note that we use a different point of the life cycle (the gamete frequencies) as this is more easily tractable due to the reduced number of variables.

The diploid model can be obtained by applying equations (2) and (5) to males as well, with the corresponding relevant substitutions.

For the analysis, we focus on the equilibrium of the system defined by:

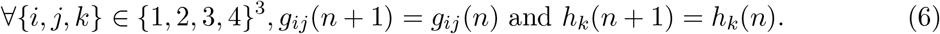

These equilibria can either be obtained by solving the system of equations presented above numerically, or by focusing on some of the known and potentially biological relevant equilibria, like fixation of a given haplotype. The stability of the equilibria is then obtained by computing the Eigenvalues of the Jacobian matrix at the focal equilibrium. If the absolute value of all Eigenvalues are below 1, the equilibrium is locally stable. For a more detailed explanation, see Otto and Day (2007, Chap. 7). We use this method to derive necessary and sufficient conditions for the existence and stability of the different evolutionary outcomes.

### Simulations

Derivations, simulations, and data fitting were performed in *Mathematica* (v 10.4.1.0; Wolfram Research, Inc., 2016). To enable complete reproducibility of the results, we provide an Online Supplement that documents all steps of the analysis as well as the code used for simulations and figures. Equilibrium genotype frequencies were obtained numerically when possible, or based on simulations until the differences between genotype frequencies of two consecutive generations were smaller than 10^−8^ (or stopped after 10^5^ generations without convergence).

### Fitting the model to a natural ant population

To compare our model with data from the natural, hybridizing Finnish ant population, we estimated the different genotype frequencies of parental *F. polyctena*-like and *F. aquilonia*-like individuals from the data. Assuming that the natural population is at equilibrium, we fit the data (Table S2) to the model by calculating the sum of squared differences between the observed data and predicted equilibrium frequencies. Complete details of data estimation and model fitting are given in the Supplementary Methods and Supplementary Results.

## Results

In this section, we describe the dynamics of a hybrid population under our model, with a particular focus on quantifying the differences between the haplodiploid and the diploid model. Two parameter domains are of particular interest:

1. The case of free recombination and strong epistasis (i.e., large *γ*_1_, *γ*_2_) most likely resembles that of the natural ant hybrid population that inspired the model. Here, the hybrid incompatibility loci are located on different chromosomes, and epistasis is strong enough to erase a large fraction of male zygotes during development.
2. The case of low recombination is most relevant for the effects of a fitness landscape with epistasis (i.e., a “rugged” landscape) in X or Z chromosomes. Here, epistasis could arise, for example, through interactions between regulatory regions and their respective genes.

### Evolutionary scenarios

Below, we describe four different types of evolutionary stable states (i.e., equilibrium scenarios) of the model, which represent long-term solutions to the opposing selective pressures of the hybridization-averse force of recessive negative epistasis and the hybridization-favoring heterozygote advantage. The population will attain these equilibria if no further pre- or post-zygotic barriers or other functional mutations appear. Next, we provide various necessary and sufficient analytical conditions for these scenarios. Figure 3 illustrates the potential equilibria by means of phase diagrams.

**Figure 3:**
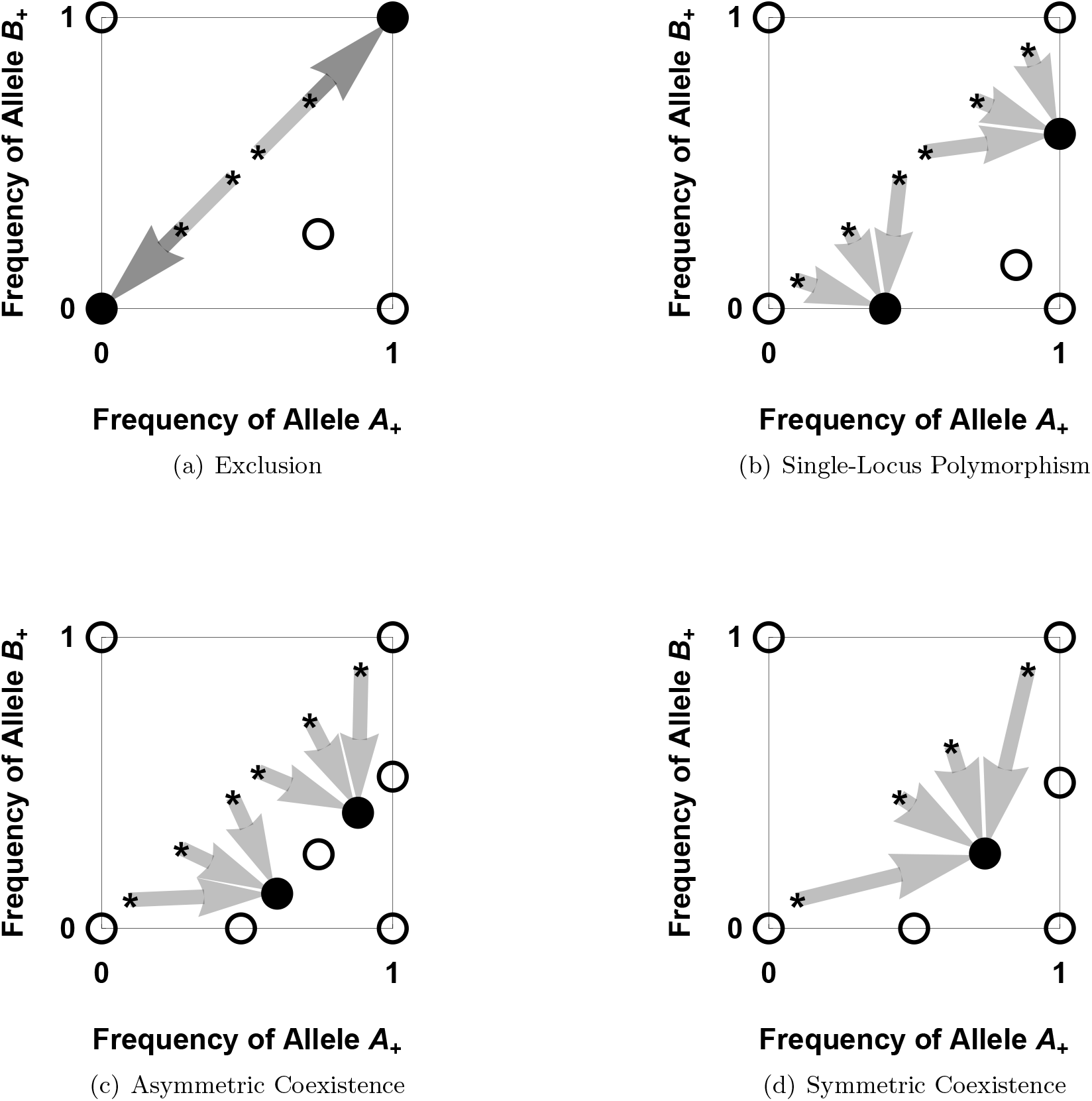
Phase-plane diagrams illustrating possible evolutionary scenarios in the haplodiploid model. The filled black dots show locally stable equilibria and the empty dots show unstable ones. The gray arrows show the basin of attraction starting from secondary contact scenarios (black asterisks on the line at *pB*_+_ = *PA*_+_). Panel (a) illustrates exclusion: There are 2 external locally stable equilibria, each corresponding to the fixation of a parental population haplotype. (Here, *σ* = 0.02, *γ*_1_ = 0.9, *γ*_2_ = 0.11, *ρ* = 0.5, and *α* = 0.) Panel (b) represents a single-locus polymorphism. Only one locus is polymorphic, leading to the maintenance of the weaker of the two incompatibilities (the *A*_−_*B*_+_ interaction). (Here, *σ* = 0.009, *γ*_1_ = 0.11, *γ*_2_ = 0.002, *ρ* = 0.5, and *α* = 0.) Panel (c) corresponds to asymmetric coexistence. Two internal equilibria are locally stable, with one allele close to fixation. This scenario minimize the expression of the strongest interaction *A*_+_*B*_−_. (Here, *σ* = 0.03, *γ*_1_ = 0.11, *γ*_2_ = 0.0013, *ρ* = 0.5, and *α* = 0.) Panel (d) shows symmetric coexistence. Frequencies of alleles *A*_−_ and *B*_−_ are symmetric around 0.5, with *pB*_+_ = 1 – *PA*_+_. This scenario maximizes the formation of female heterozygous hybrids. (Here, *σ* = 0.09, *γ*_1_ = 0.3, *γ*_2_ = 10^−4^, *ρ* = 0.5, and *α* = 0.)

### Exclusion

The *exclusion* scenario corresponds to the hybrid population becoming identical to one of the two parental populations, either *P*_+_ or *P*_−_, and the other parental population being therefore excluded. It occurs when both alleles from one of the founder subpopulations are purged, leading to a monomorphic stable state of the population (Fig. 3(a)). In this case, the initial frequency of *A*_+_*B*_+_ versus *A*_−_*B*_−_ individuals mainly determines the outcome (i.e., the population is swamped by the majority subpopulation). As a rule of thumb, this outcome is observed when recombination is frequent and when the hybridization-averse force of negative epistasis is strong as compared with the hybridization-favoring heterozygote advantage (*γ*_1_, *γ*_2_ ≫ *σ*).

With regard to the apparent sexual/ploidy conflict in the haplodiploid model, exclusion can be interpreted as a victory of the haploid males because all polymorphism is lost and no low-fitness hybrid males are produced. Conversely, since all polymorphism is lost, diploid females “lose” in this case and neither high-fitness introgressed (i.e., those individuals carrying only one ‘foreign’ allele) nor highest-fitness heterozygous hybrid females are produced. As discussed below, exclusion is never a possible outcome in the diploid model, in which there are no differences in ploidy.

### Single-locus polymorphism

A *single-locus polymorphism* occurs when one allele is purged from the population but the other locus remains polymorphic at equilibrium (Fig. 3(b)). Because this is possible for either of the two loci, two such equilibria exist simultaneously, which are reached depending on the initial haplotype frequencies. This outcome is observed when recombination is frequent, epistasis is asymmetric (*γ*_1_ ≠ *γ*_2_), and heterozygote advantage is small (*γ*_1_ ≫ *σ*). Like asymmetric coexistence below, this case represents a compromise between the hybridization-averse and hybridization-favoring forces of negative epistasis and heterozygote advantage, and is reached by maximizing the number of introgressed individuals of one founder subpopulation.

In the haplodiploid model, this scenario can be seen as a haploid-dominated compromise. Since one locus is fixed, one epistatic interaction has disappeared and few low-fitness hybrid males are produced. In females, high-fitness introgressed female frequencies are maximized but, since one locus is fixed, the highest-fitness heterozygous hybrid female genotypes are no longer available.

The single-locus polymorphism is never stable in the diploid model, i.e., when the ploidy difference is removed from the model. In a diploid population that resides transiently at single-locus polymorphism, a rare mutant at the second locus will always begin as heterozygote and therefore reap the advantage of being a heterozygote hybrid long before it suffers the epistatic cost of being a homozygote hybrid.

### Asymmetric coexistence

*“Asymmetric” coexistence* occurs when all four haplotypes remain in the population and the frequency of introgressed individuals of one founder subpopulation is maximized (Fig. 3(c)). Because this can be achieved in two ways, two possible equilibria reside off the diagonal line *p_B_* = 1 – *p_A_* (where *p_A_* and *p_B_* denote the allele frequencies of the ‘−’ allele at the respective locus), and the initial contribution of different haplotypes determines which equilibrium will be attained. Like the single-locus polymorphism, this equilibrium represents a compromise between hybridization-averse and hybridization-favoring forces that is reached by maximizing the number of introgressed individuals. Our simulations demonstrate that this scenario is rarely present in haplodiploids, and it generally involves asymmetric epistasis and intermediate-strength heterozygote advantage.

In the haplodiploid model, asymmetric coexistence can be seen as a compromise that is dominated by the diploids. Unlike in the single-locus polymorphism scenario, both loci are polymorphic and some double-heterozygous hybrid females are produced. But, unlike the symmetric coexistence scenario described below, females are not victorious over males because such high-fitness hybrid females are produced only at low frequencies.

### Symmetric coexistence

*Symmetric coexistence* occurs when a locally stable equilibrium exists on the diagonal *p_B_* = 1 – *p_A_*, such that the number of heterozygous hybrids is maximized (Fig. 3(d)). Our notion of “symmetric” refers to the total fraction of alleles from the *P*_+_ and *P*_−_ founder populations segregating at equilibrium, which is equal in this case. Here, prolonged hybridization is a mutual best-case scenario for both populations. This equilibrium is most likely when recombination is weak or when the hybridization-favoring force of heterozygote advantage is strong as compared with the hybridization-averse negative epistasis (*σ* ≥ *γ*_1_, *γ*_2_). In the haplodiploid model, symmetric coexistence represents a victory for the diploids, because they maximize their own fitness without regard to the production of unfit hybrid haploids.

The four evolutionary stable states described above usually result in either a single, globally stable equilibrium (in the case of symmetric coexistence) or a bistable system, in which two locally stable equilibria exist. In rare cases and close to bifurcation points, we observe cases of tristability, which are further described in Figure S2.

### Stability analysis of the model

Although the model dynamics are too complex to derive general analytical solutions, we were able to perform stability analyses for specific cases, which yield information about the general behavior of the model. In the following, our use of ‘>’ and ‘<’ does not necessarily imply strict inequalities; we merely did not explicitly study the limiting cases. For ease of notation, we refer to heterozygote advantage in terms of *ω* below; recall that *ω* = 1 + *σ*.

### Conditions for symmetric coexistence when epistasis is lethal

We begin by describing the equilibrium structure when epistasis is lethal, i.e. *γ*_1_ = *γ*_2_ = 1; this case may resemble that in the natural ant population, in which most hybrid males do not survive to reproduce. For the haplodiploid model, we obtain a full analytic solution of the identity, existence and stability of equilibria. Here, only two outcomes are possible: symmetric coexistence and exclusion (Fig. 4(a)). As necessary and sufficient criterion for exclusion, we obtain

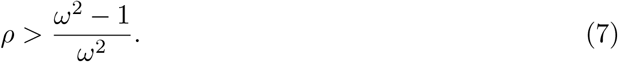

**Figure 4:**
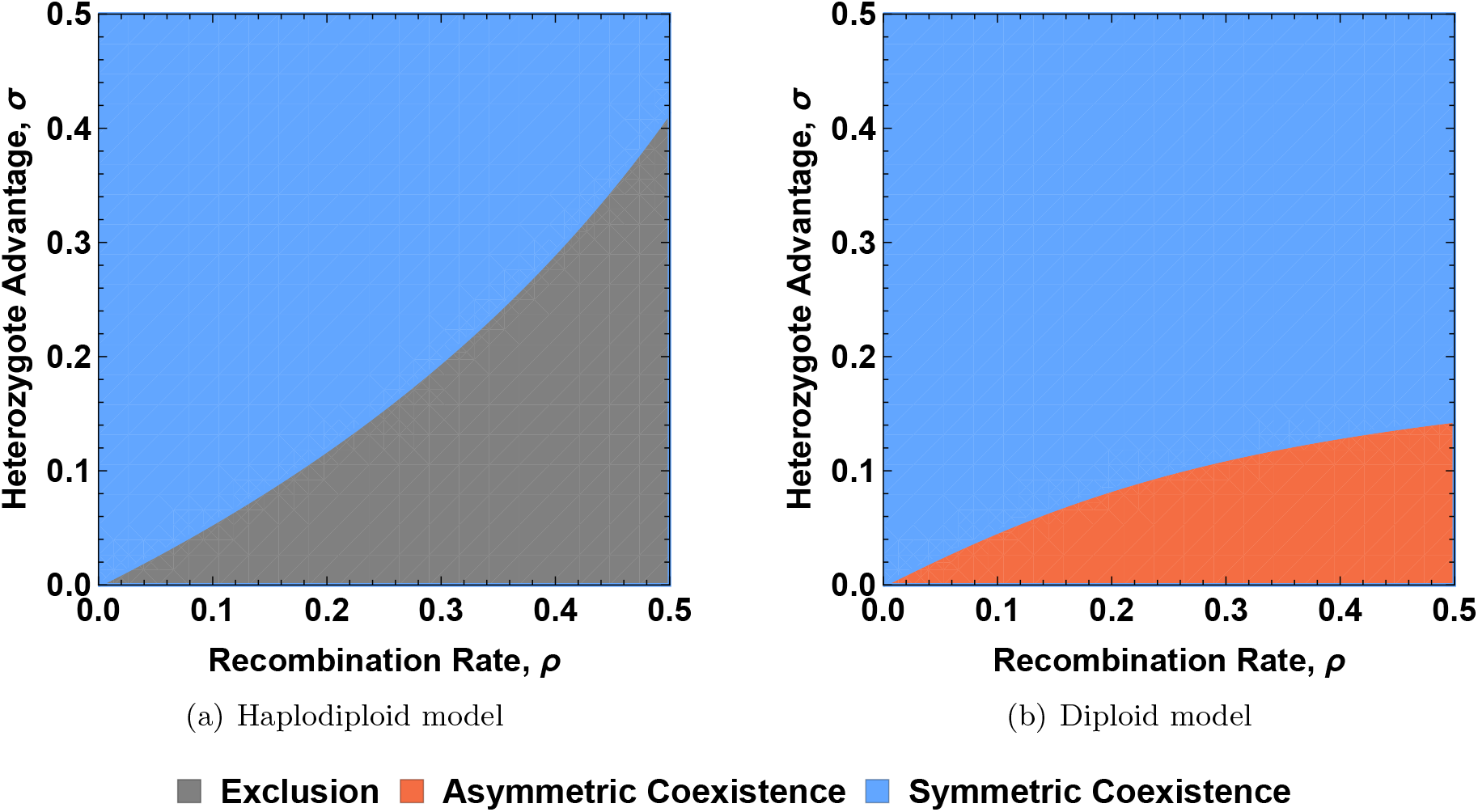
Symmetric coexistence can be locally stable if the heterozygote advantage, *σ*, is strong enough to compensate for recombination breaking up the parental haplotypes. Here we assume that epistasis is symmetric and lethal (*γ*_1_ = *γ*_2_ = 1). Panel (a) is an illustration of the condition for haplodiploids given in equation (7) and panel (b) of equation (8) for diploids.

Thus, exclusion is only possible if heterozygote advantage is not too strong, and if recombination is breaking up gametes sufficiently often to significantly harm the haploid males.

For the diploid model, we can show that no boundary equilibrium is ever stable; asymmetric and symmetric coexistence are the only two possible outcomes. Although it was not possible to perform a stability analysis on the internal equilibria, we were able to propose a condition for asymmetric coexistence, which has been evaluated numerically:

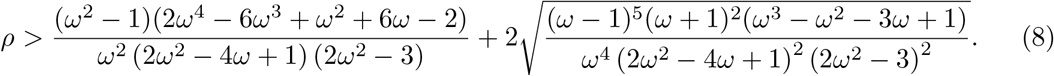

Although this expression is not very telling, its illustration in Figure 4(b) demonstrates how different this criterion is from that of the haplodiploid model. In the diploid model, males and females evolve on the same fitness landscape. Therefore, both males and females benefit from heterozygote advantage. This reduces the influence of the hybrid incompatibility on the optimal location of the population in genotype space, which thereby makes asymmetric coexistence less likely. Indeed, a heterozygote advantage of *ω* – 1 = *σ* >≈ 0.14 is sufficient to ensure symmetric coexistence for all recombination rates, whereas in the haplodiploid model, 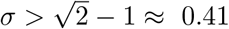 is necessary for symmetric coexistence independent of the recombination rate.

### General stability conditions in the haplodiploid model

Using the results derived for the case of lethal epistasis, and by means of critical examination of the existence and stability conditions that we were able to compute analytically, we arrived at several illustrative conjectures delimiting the evolutionary outcomes in the haplodiploid model when epistasis is not lethal (*γ*_1_, *γ*_2_ ≠ 1). These were all confirmed by extensive numerical simulations (see Mathematica Online Supplement). Note that assortative mating was not considered here.

Firstly, strong heterozygote advantage can always override the effect of epistasis. Specifically, if

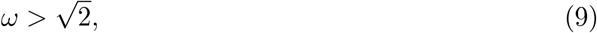

the evolutionary outcome is always symmetric coexistence, regardless of the values of *γ*_1_ and *γ*_2_. This is true not only for a single pair of interacting loci, but also for an arbitrary number of independent incompatibility pairs, because the detrimental effects caused by each incompatibility pair are eventually resolved independently (see also the section on multiple loci below). This result can be deduced from equation (7) for *ρ* = 0.5 and therefore corresponds to an upper bound: if heterozygote advantage is very strong, recombination no longer affects the outcome.

Secondly, recombination is a key player to determine whether compromise or exclusion can occur. In particular,

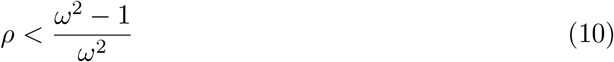

is a sufficient condition for the observation of symmetric coexistence, independent of the strength and symmetry of epistasis. This makes intuitive sense, because hybrid incompatibility is masked until gametes are broken up by recombination.

Thirdly, for symmetric epistasis (*γ*_1_ = *γ*_2_), there are three possible equilibrium patterns: symmetric coexistence, exclusion, and tristability of the two former types of equilibria. A necessary and sufficient condition for observation of anything but symmetric coexistence is

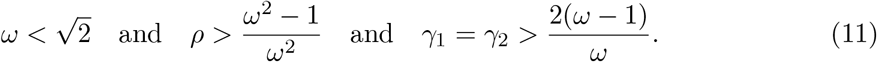

If the recombination rate *ρ* and the epistatic effects *γ*_1_, *γ*_2_ are very close to this limit, there is tristability; if they are far away, there is exclusion (cf. Fig. 5).

**Figure 5:**
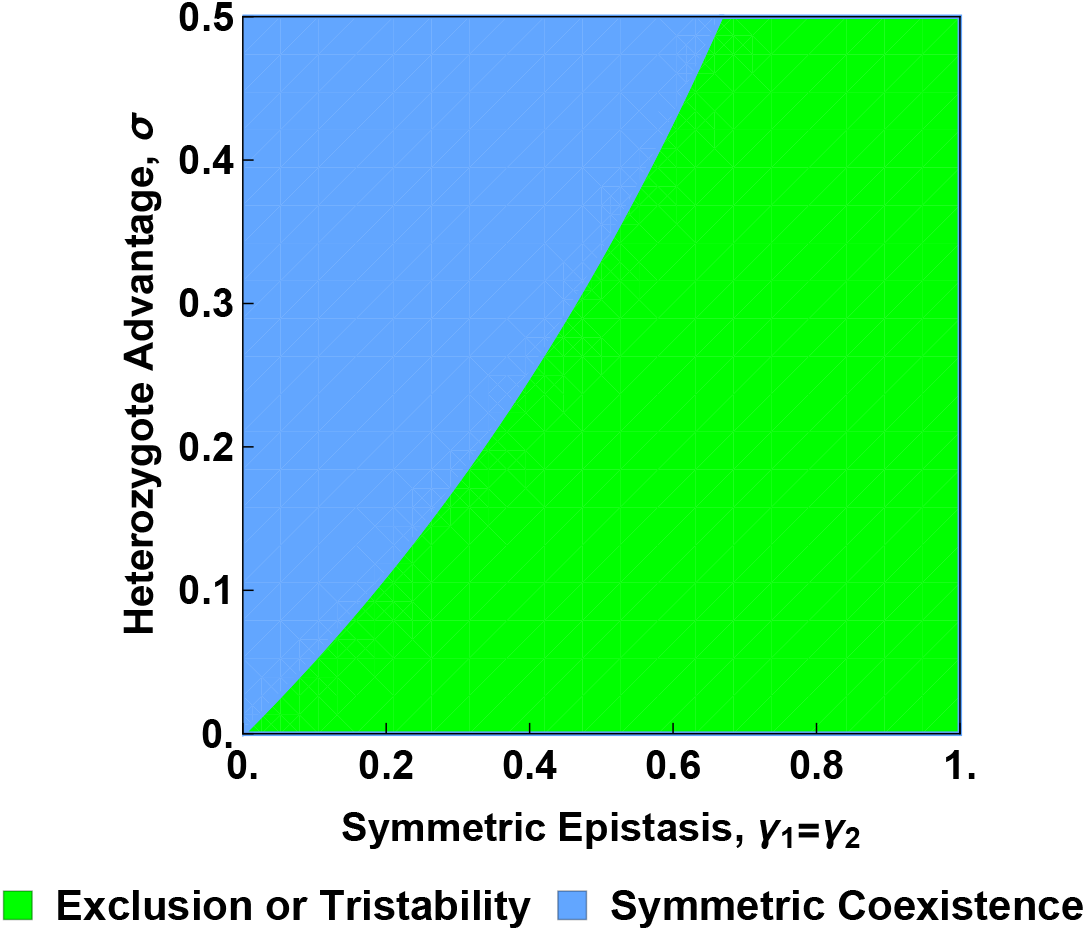
In haplodiploids, symmetric coexistence requires that heterozygote advantage, *σ*, is strong enough to both compensate for recombination such that the condition in equation 10 is fulfilled (see also Fig. 4(a)), and to overcome the deleterious effects of epistasis, as expressed by condition 11 for symmetric epistasis.

Finally, for asymmetric epistasis (*γ*_1_ ≠ *γ*_2_), the dynamics display the whole range of possible evolutionary outcomes: symmetric coexistence, asymmetric coexistence, single-locus polymorphism, exclusion, as well as tristability of exclusion *and* symmetric coexistence, and single-locus polymorphism *and* symmetric coexistence. The local stability criterion for the stability of the monomorphic equilibria (i.e., the criterion for exclusion, or tristability of exclusion and symmetric coexistence) is

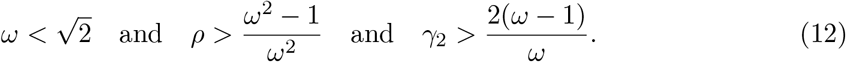

Thus, if epistasis is strong as compared with heterozygote advantage, no degree of asymmetry is sufficient to promote a compromise between males and females (i.e., single-locus polymorphism or asymmetric coexistence). In fact, we observe the following necessary (but not sufficient) condition for a single-locus polymorphism:

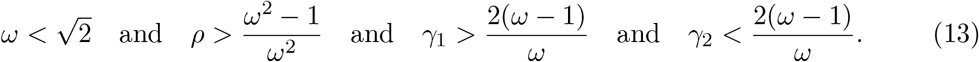

Hence, only a tight balance between the selective pressures of epistasis and heterozygote advantage in combination with asymmetry of the hybrid incompatibility promotes a longterm equilibrium with compromise.

### An extension to multiple loci

#### Incompatibilities involving four loci

Above, we have demonstrated that recombination is an essential player when determining whether exclusion or coexistence is the long-term outcome in the haplodiploid dynamics. In order to see how our results change in the (biologically relevant) case of multiple hybrid incompatibilities, we implemented the dynamics for four loci. Given the complexity of the system, we considered only lethal incompatibilities, i.e. *γ_i_* = 1 for all interactions *i*. With this extension, we consider two scenarios. Firstly, in the “*pairwise*” case we consider pairs of independent hybrid incompatibilities, where we assume that the incompatible loci are located next to each other (locus **A** interacts with locus **B** at recombination distance *ρ*_12_, and locus **C** with locus **D** at recombination distance *ρ*_34_), which leaves four viable male haplotypes (*A*_+_*B*_+_*C*_+_*D*_+_, *A*_+_ *B*_+_ *C*_−_*D*_−_, *A*_−_*B*_−_*C*_+_*D*_+_ and *A*_−_B_−_*C*_−_*D*_−_). Secondly, in the “*network*” case we assume that all loci interact such that only two viable male haplotypes exist *A*_+_*B*_+_*C*_+_*D*_+_ and *A*_−_*B*_−_*C*_−_*D*_−_. In both cases, heterozygote advantage is defined as before, now acting on all four loci multiplicatively.

Under this model, we derived the conditions under which exclusion (the purging of all foreign alleles resulting in a monomorphic equilibrium) is locally stable (cf. Mathematica Online Supplement). For the pairwise case, exclusion is stable only if heterozygote advantage is relatively weak:

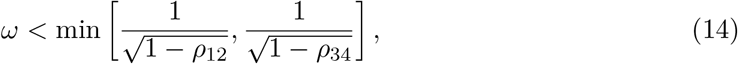

where *ρ_ij_* is the recombination rate between neighboring loci *i* and *j*. Note that this is independent of the recombination rate between non-interacting loci, here *ρ*_23_. If *ρ*_12_ = *ρ*_34_, this expression is equivalent to equation 7 (Fig. 4(a)). Overall, this condition indicates that exclusion, which we define as the fixation of one of the parental haplotypes, is less likely with four interacting loci than with two. This is because the fate of the two pairs of incompatibilities is decided independently, and exclusion requires that both pairs of incopatibilities fix for the same parental haplotype.

For the network case, the condition for stability of exclusion (see also Fig. S3) is

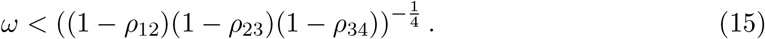

In this scenario, exclusion is a more likely outcome with two pairs of incompatibilities than with one. This is because there are more unfit intermediate types in this scenario as compared with the pairwise model. Specifically in males,14 out of the 16 possible haplotypes do not survive to adulthood. To compensate for this fitness cost, any alternative evolutionary outcome requires strong heterozygote advantage.

#### Incompatibilities involving an arbitrary number of loci

From the results for two and four loci, we derived a conjecture that generalizes to an arbitrary number of loci. For the pairwise case, equation 14 can be generalized to

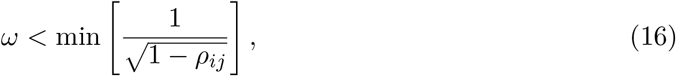

with *i* and *j* representing neighboring interacting loci. Note that this result holds only if interacting loci are next to each other on the same chromosome, or if all loci are unlinked (in which case it simplifies to 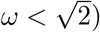.

For the network case, equation (15) generalizes to

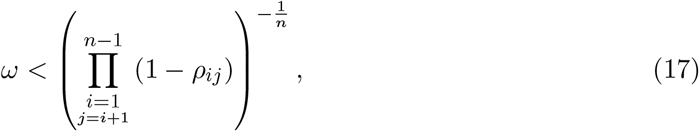

with *i* and *j* neighboring loci and *n* the total number of loci in the network. Unlike in the pairwise case, the results for the network case do not depend on the genetic architecture (here, the ordering of loci along the genome).

We can therefore deduce that, for the pairwise case, exclusion becomes increasingly unlikely as the number of pairs of independent hybrid incompatibilities involved in the genetic barrier increases. Conversely, the opposite result is observed for the network case: more loci make exclusion a more likely outcome, but each additional interaction contributes less (cf. Fig. S3).

### Increased assortative mating counteracts recombination and heterozygote advantage

Increasing the strength of assortative mating, *α* > 0, counteracts the hybridization-favoring effect of heterozygote advantage, because matings between individuals with the same genotype are more common under stronger, positive assortment. Under sufficiently large positive *α*, exclusion is unavoidable. In general, increasing *α* leads to less maintenance of polymorphism in the population (Fig. S4). Conversely, when *α* < 0, which means that individuals prefer to mate with those whose genotype is most different from their own, polymorphism is more likely to be maintained in the population.

Also with assortative mating, recombination remains a key player in determining the evolutionary outcome. When *α* < 0 and recombination is small, symmetric coexistence is possible even in the absence of heterozygote advantage (i.e., *σ* = 0; Fig. S4). Indeed, under these conditions and assuming epistasis is very strong, (almost) all hybrid males are dead and only parental males survive. This ‘disassortative’ mating (*α* < 0) creates a bias for the rare male haplotype. For example, if one female genotype increases in frequency, it will seek mainly the males of the other parental haplotype to reproduce with (which are currently rare, as their frequency is directly tied to the frequency of the females in the previous generation. This will increase their reproductive success, which leads to an increase of this haplotype frequency. Therefore, under this mate choice regime, we would observe a stable population composed almost exclusively of the *A*_+_*B*_+_ and *A*_−_*B*_−_ haplotypes.

### Differences between the haplodiploid and the diploid systems

As described above and illustrated in Figure 6, the resulting haplodiploid dynamics display a wider range of possible evolutionary outcomes than the diploid dynamics. Because both males and females profit from heterozygote advantage in the diploid model, polymorphism is always maintained; in other words, even the smallest amount of heterozygote advantage promotes the creation or maintenance of diversity in diploids (Table S3). Conversely, in the haplodiploid model, polymorphism can be lost either at one or both loci, resulting in a single-locus polymorphism or exclusion. Thus, alleles responsible for incompatibilities are more effectively purged in the haplodiploid model.

**Figure 6:**
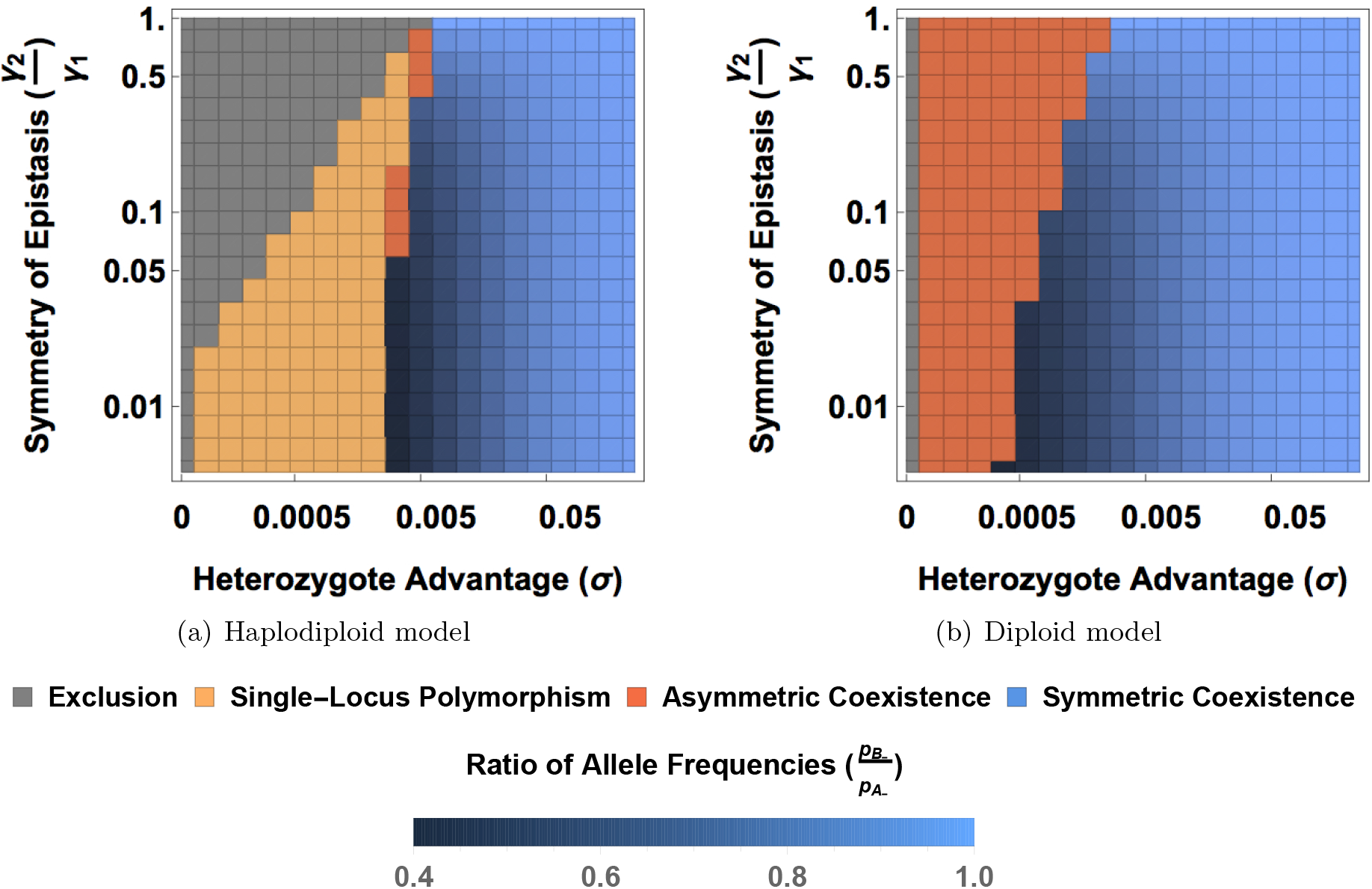
More evolutionary outcomes are possible in (a) the haplodiploid than (b) the diploid model. The y-axis shows the degree of asymmetry of epistasis, displayed as the ratio of the two epistasis parameters 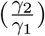 for a constant value of *γ*_1_ = 0.01. For symmetric coexistence, the locally stable equilibrium can be at any point on the diagonal *p*_*B*_−__ = 1 – *p*_*A*_−__, where *p*_*A*_−__ and *p*_*B*_−__ denote the allele frequencies of the – allele at the respective locus. Blue shading illustrates the location of the equilibrium at symmetric coexistence: darker shades correspond to a bigger disparity in allele frequencies. This is the case when the asymmetry of the two epistasis parameters is large (i.e. smaller values on the y-axis) because smaller values of *γ*_2_ favor the *A*_−_*B*_+_ haplotype over the *A*_+_*B*_−_ haplotype. (Here, *γ*_1_ = 0.01, *ρ* = 0.5, *α* = 0.)

In the diploid model, a single-locus polymorphism is never stable: Assume locus *α* is polymorphic and locus *B* is fixed for allele *B*_+_. Then, a new mutant carrying allele *B*_−_ will always have a selective advantage regardless of the genotype in which it first appears (Table S3). In contrast, in the haplodiploid model, this is no longer true as the mutant carrying allele *B*_−_ will have a much lower fitness in males when associated to allele A+. Therefore, if the cost of generating this unfit haplotype in males overrides the advantage in females, and allele *A*_+_ is at high frequency, then invasion of the *B*_+_ mutant may be prevented, leading to the stability of the single-locus polymorphism.

When polymorphism is maintained at both loci at equilibrium (i.e., asymmetric and symmetric coexistence), epistasis creates associations between the compatible alleles which results in elevated linkage disequilibrium (LD). Recombination breaks the association between alleles, thus high recombination decreases normalized LD (*D*′, where 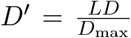 (Lewontin, 1964); Fig. S5). *D*′ increases with the strength of heterozygote advantage at low recombination rates because it maximizes the discrepancy between highly fit double-heterozygote females on the one hand that can, under low recombination rate, still produce many fit male offspring and introgressed females on the other, who are less fit and produce many unfit hybrid males

In Figure S6, we compare the normalized LD (i.e. *D*′) between the haplodiploid and diploid models. When polymorphism is maintained at both loci in both the haplodiploid and diploid model, normalized LD is always larger in haplodiploids than diploids. The difference in normalized LD between haplodiploids and diploids is maximized for intermediate recombination rates, where recombination is strong enough to create unfit hybrid genotypes, but not efficient enough to break the associations that are generated. Due to the increased selection against hybrid incompatibility in haploid males in the haplodiploid model, the normalized LD is usually 2–3 times higher in the haplodiploid as compared with the diploid model.

Thus, the hybrid incompatibility leaves a statistical signature in a population, even if the population finds itself at an equilibrium. The increased association across the genome, exhibited if the interacting loci are on the same chromosome, may also result in an underestimate of the recombination rate. Although both the diploid and the haplodiploid models display the elevated LD signal, it is much more pronounced in the haplodiploid scenario. This is because only an eighth of the possible diploid male genotypes suffer the cost of the incompatibility as compared to half of the possible haploid male genotypes.

## Discussion

Multiple recent studies have highlighted the pervasive nature of hybridization and its potential consequences for diversification and speciation (Abbott et al., 2013; Runemark et al., 2017; Montecinos et al., 2017). We here modeled the fate of a hybrid population in a scenario in which hybridization is simultaneously favored and selected against, inspired by a natural population of hybrid ants that simultaneously displays heterosis and hybrid incompatibility. In addition, both adaptive introgression and hybrid incompatibilities have been identified in natural systems (Heliconius Genome Consortium, 2012; Whitney et al., 2015; Corbett-Detig et al., 2013) and it is therefore likely that both processes may occur simultaneously during a single hybridization event. Furthermore, we were interested in comparing the long-term evolution of populations exposed to these opposing selective pressures under different ploidies (haplodiploid versus diploid), since it has been argued that haplodiploids might speciate more easily than diploids (Lohse and Ross, 2015). Finally, the comparison of ploidies can also be transferred to the case of diploid species with sex chromosomes, in which the described fitness landscape results in the diploid dynamics on the autosomes, and in the haplodiploid dynamics on the X/Z chromosome.

Our model considers a population in which heterozygote advantage and hybrid incompatibility act simultaneously on the same pair of loci, which creates a rugged fitness landscape with a ridge of high-fitness heterozygote genotypes, adjacent to which there are holes of incompatible double homozygotes (Fig. 2(a)). In haplodiploids, haploid males cannot profit from heterozygote advantage but suffer strongly from hybrid incompatibility (Fig. 2(b)). This results in a conflict of ploidies/sexes over the optimal location in the fitness landscape, because haploid males survive best if one parental haplotype is fixed whereas diploid females profit from maximum heterozygosity. Although females suffer from the same incompatibility as males, their presence is mainly masked in the diploid individuals because of the recessivity of the hybrid incompatibility. This is similar to Haldane’s rule (Charlesworth et al., 1987; Koevoets and Beukeboom, 2009).

### How ploidy matters

We found that, in the haplodiploid model, there exist four different stable outcomes of the conflict over hybrid status (Fig. 3): exclusion, where “males/haploids win”; symmetric coexistence, where “females/diploids win”; and two outcomes, single-locus polymorphism and asymmetric coexistence, where a compromise between male costs and female benefits is mediated by high frequencies of introgressed females. In fact, since low-frequency heterozygotes are favored both in males and in females in the diploid model, while only suffering the hybrid cost if introgressed alleles rise to high frequencies, exclusion and single-locus polymorphism never occur in the diploid model, which reduces the number of possible outcomes to asymmetric and symmetric coexistence. Therefore, consistent with Pamilo (1979); Pamilo and Crozier (1981); Patten et al. (2015), we found that introgression and maintenance of polymorphism, and thus long-term hybridization, are less likely in haplodiploids as compared to diploids.

Prior work has found that in haplodiploid species traditional sexual conflict tends to be resolved in favor of females because genes spend two thirds of their time in females (Albert and Otto, 2005). In our model, the co-occurrence of heterozygote advantage and hybrid incompatibility also creates an apparent sexual conflict that is caused by the difference in ploidy between the sexes. For several scenarios, we here derived the conditions for whether this conflict is resolved in favor of diploid females or haploid males. We find, that in addition to the strength of selection, recombination is a major player (cf. Fig. 4 and equation 12); only if recombination breaks up gametes, the hybrid incompatibility is expressed. With free recombination, i.e., if the interacting genes are found on separate chromosomes, heterozygote advantage has to be very strong to counteract the hybrid incompatibility. We find that it has to be on the same order of magnitude as the strength of the incompatibility, but can be slightly lower in its absolute value. For example, heterozygote advantage with strength 41% is sufficient to result in symmetric coexistence even if the incompatibility is lethal (Fig. 4B). Thus, under consideration of absolute magnitude across the full parameter range, our results are consistent with prior work. However, reported cases and potential mechanisms of hybrid incompatibility indicate that large effects are feasible, whereas observed cases of heterozygote advantage or heterosis of large effect are relatively rare (Hedrick, 2012). Therefore, it may well be that under natural circumstances, the conflict modeled here may indeed be likely to be resolved via purging of at least one incompatible allele and thus in favor of males/haploids.

As expected in the presence of epistasis, we observed that linkage disequilibrium (LD) is elevated at all polymorphic stable states (i.e., for symmetric and asymmetric coexistence) both in the diploid and haplodiploid models, especially at intermediate recombination rates. This is particularly true for haplodiploids, which display about 2-3 times the LD of the diploid model with the same parameters. Transferred to the context of X/Z chromosomes, this is consistent with observations of larger LD on the X chromosome as compared with autosomes (Wall et al., 2002; Sandor et al., 2006; Li and Merilä, 2010). It has been argued that this is because selection is more effective on X-linked loci: recessive deleterious mutations are more visible to selection in haploid individuals (Charlesworth et al., 1987). However, a hybrid incompatibility accompanied by heterosis/heterozygote advantage as in our model may not be purged but create a continuous high-LD signal in an equilibrium population. This can potentially result in less efficient recombination and in underestimates of recombination rates on X chromosomes (because recombined individuals are not observed).

### Generalization to multiple incompatibilities

Exclusion remains a stable solution when we extend the model to multiple loci and incompatibilities. We describe an interesting difference between multiple independent pairs of incompatibilities, and multiple loci that all interact with each other: in the latter case, exclusion becomes increasingly probable because the number of viable males decreases. This scenario of higher-order epistasis has recently received attention with regards to speciation (Paixão et al., 2014; Fraïsse et al., 2014; Kulmuni and Westram, 2017), and it will be interesting in the future to identify molecular scenarios (for example, involving biological pathways) that could result in such incompatibilities. In contrast, exclusion becomes less likely in the case of independent incompatibility pairs, where each incompatibility has to be purged independently, and in the same direction, for exclusion to occur. Here, mechanisms that reduce the recombination rate, such as inversions, could potentially invade and tilt the balance towards coexistence and thus maintenance of polymorphism in the hybrid population. It is important to note that the independent purging of incompatibilities, which leads to a decreasing probability of exclusion with increasingly many incompatibility pairs, is only true in effectively infinite-sized populations. In small populations, we expect that exclusion becomes a more likely scenario, especially if lethal incompatibility pairs are present.

### Model assumptions

We chose a classical population-genetic modeling approach (Bürger, 2000; Nagylaki et al., 1992) to study how the co-occurence of heterozygote advantage and hybrid incompatibility affect the long-term dynamics of a hybrid population. By treating the problem in a deterministic framework and considering only two loci throughout most of the manuscript, we greatly oversimplify the situation in the natural population that inspired our model. However, at the same time this allowed us to gain a general insight, (often by means of analytical expressions), into how opposing selective pressures in genomes may be resolved, and to contrast these outcomes between haplodiploid and diploid systems. In addition to some obvious mechanisms at play in natural populations, which we ignore in our model (e.g., random genetic drift), some extensions of the model could be interesting to elaborate on in the future. For example, the ant populations represent networks of interacting nests with many queens per nest, but potentially different mating flight timing that depends, for example, on sun exposure in the spring. Thus, for the purpose of population-genetic inference of the evolutionary history (and potential evolutionary fate) of the hybrid ant population in Finland, it would be desirable to incorporate population structure, uneven sex ratios at birth, and sex-biased dispersal into the model, and obtain population-genomic data to infer evolutionary parameters.

### Is the natural population at an equilibrium of asymmetric coexistence?

Model fitting results (see Supplementary Methods, Results, and Discussion) are inconclusive about the fate of the natural ant population that inspired our model. Our results suggest that the natural population might be approaching an evolutionary outcome that allows a compromise between male and female interests; either as single-locus polymorphism or via asymmetric coexistence. In particular, our model is able to explain the unusual skew in the population, where *F. aquilonia*-like parental genotypes far outnumber *F. polyctena*-like genotypes (see Supplement). Furthermore, the high recombination rates and strong prezygotic mechanisms operating in the natural population (Kulmuni et al., 2010; Kulmuni and Pamilo, 2014), are consistent with a parameter domain in our models at which asymmetric coexistence can be stably maintained over a wide range of values of female hybrid advantage. More complex models, for example including more than two incompatibility loci, may be better able to explain the high frequencies of introgressed as compared to parental females observed in the natural hybrid population. As argued in the Results, interactions at or between multiple loci should result in steeper differences of introgressed-allele frequencies across life stages than our model is able to produce.

### Implications for hybrid speciation

Our model illustrates how the co-occurrence of heterozygote advantage and hybrid incompatibility affects haplodiploid and diploid populations. We can hypothesize how these different outcomes may provide an engine to hybrid speciation, or which other long-term evolutionary scenarios we expect to arise. The case of exclusion, which is possible only in the haplodiploid model, will lead to loss of diversity in the hybrid population, and, in the two-locus case, should result in the reversion of the hybrid population into one of its parental species. However, if multiple pairs of interacting loci are resolved independently, they may be purged randomly towards either parent, which could result in a true hybrid species that is isolated from both its parental species (Buerkle et al., 2000; Butlin and Ritchie, 2013; Schumer et al., 2015). In fact, our finding that exclusion is less likely to occur in populations with multiple pairs of interacting loci may result from exactly this mechanism, but it is beyond the scope of this manuscript to explore this further.

The long-term fate of the population is less straightforward to anticipate in the case of polymorphic stable equilibria. For any of these, heterozygote advantage is strong enough to stabilize the polymorphism either at one or both loci. Without further occurrence of functional mutations, males (in the haplodiploid model) and double-homozygotes for the incompatible alleles will continue to suffer a potentially large fitness cost. Mechanisms that could reduce this cost would be increased assortative mating or decreased recombination. However, neither of these would necessarily cause isolation from the parental species, unless they involved additional hybrid incompatibilities which isolate the hybrid population from its parental species. Alternatively, mutations that lower the hybrid fitness cost could invade, which would result in a weakening of species barriers and promote further introgression from the parental species. This indicates that any scenario in which polymorphic equilibria are stable may indeed be an unlikely candidate for hybrid speciation. Considering that such stable polymorphism (either as symmetric or asymmetric coexistence) is the only possible outcome in the diploid model, this results in the prediction that hybrid speciation would be more likely in a haplodiploid scenario. This is an interesting observation that is in line with other predictions that haplodiploids speciate more easily, that X/Z chromosomes are engines of speciation (Lima, 2014), and that hybrid speciation is rare (Schumer et al., 2014).

### Relevance of the model for sex chromosomes

Haplodiploids and X/Z chromosomes have a similar mode of inheritance, where one sex carries a single copy of the chromosome, and the other carries two copies. Therefore, our results apply equally to cases of X-to-X or Z-to-Z hybrid incompatibilities (Lohse and Ross, 2015). Although haplodiploid systems do not include all of the unique evolutionary phenomena exhibited by sex chromosomes (Abbott et al., 2017), our results for haplodiploids are relevant for sex chromosomes. Our model predicts the long-term evolution of a population under the simultaneous influence of heterozygote advantage and hybrid incompatibility, and indicates the signatures that this type of fitness landscape could leave depending on whether it finds itself on an X chromosome or an autosome.

Firstly, the complex selection pressure imposed by the co-occurrence of heterozygote advantage and hybrid incompatibility manifests itself as an apparent sexual conflict on the X chromosome/in haplodiploids. This conflict is caused by the ploidy difference between the sexes. Here, the same fitness landscape that would be masked on an autosome and result in a stable polymorphism, creates a signal of sexually antagonistic selection on an X chromosome. Most importantly, this signal is created without the need for direct sexually antagonistic selection on single functional genes that have a sex-specific antagonistic effect. Thus, our model proposes an additional mechanism by which sex chromosomes can appear as hotspot of sexual conflict (e.g., Gibson et al., 2002; Pischedda and Chippindale, 2006).

Secondly, we find that purging of incompatibilities is more likely in the haplodiploid model, and thus on X/Z chromosomes. This is consistent with the faster-X theory (Charlesworth et al., 1987). However, only if recombination is strong enough, incompatibilities will become visible to selection and purged in the presence of heterozygote advantage. If they are not purged, they may persist as a long-term polymorphism, invisible to most empirical approaches, and confound population-genetic inference by creating signals of elevated linkage disequilibrium.

### Conclusion

Hybridization is observed frequently in natural populations, and can have both deleterious and advantageous effects. We here show how diverse outcomes can be produced even under a rather simple model of a single hybrid population, in which heterozygote advantage and hybrid incompatibility are occurring at the same time. Consistent with previous theory on haplodiploids and X/Z chromosomes, we found that incompatible alleles are more likely to be purged in a haplodiploid than in a diploid model. Nevertheless, our results suggest that longterm hybridization can occur even in the presence of hybrid incompatibility, and if there are many incompatible pairs or many loci involved in the incompatibility. The evolutionary fate of the Finnish hybrid ant population that inspired our model is difficult to predict; further population-genetic analysis will be necessary to gain a more complete picture of its structure and evolutionary history.

## Acknowledgements

We thank Laura Cêtre for her work on a previous version of the model. We thank Pekka Pamilo, Bret Payseur, two anonymous reviewers, and the members of the Bank and Kulmuni labs for discussion of the manuscript. This research was supported by the Fundação Calouste Gulbenkian and in part by the National Science Foundation under Grant No. NSF PHY-1125915. JK was supported by the Human Frontier Science Program, Finnish Cultural Foundation, Academy of Finland (252411 to CoE in Biological Interactions).

## Data Accessibility

The complete documentation of all steps of the analysis is available as a Mathematica Online Supplement. Ant colony data is provided as Supplementary Table S1; genotype frequency data were obtained from Kulmuni and Pamilo (2014).

## Author Contributions

CB, JK, and RB designed research, AB and CB developed the models, AHG performed simulations and data analysis, all authors interpreted the results and wrote the manuscript.

## Supplementary Methods

### Modeling of the life cycle in the absence of assortative mating

In the absence of assortative mating (i.e. *α* = 0), equations (1)–(5) still hold. Nevertheless, tracking genotypes is cumbersome and not necessary in this context (as the production of gametes is independent of their probability of forming a zygote). Therefore, we present here the system of equations used to derive the random mating case. The main difference is where we place the observation point in the life cycle. In the main manuscript, we observe the genotype frequencies of adults and track them over time. Here, we track the frequencies of the gametes (the point in the life cycle when both sexes are haploid), reducing the previous system of 18 variables to 6.

In a given generation n, the frequencies of the male and female gametes are given by *η_k_*(*n*) and *θ_i_*(*n*), respectively. Below, we present the modeled life cycle, in the absence of assortative mating, and how frequencies change from one generation to the next:

1. The first step is the formation of the zygote. In the absence of assortative mating, zygote formation depends only on the frequency of the different gametes in the population. In females, the frequency of a zygote carrying the *ij* genotype, with haplotype *i* inherited from the mother and *j* from the father is given by:

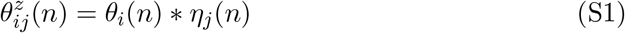 Males are formed from unfertilized female gametes. Therefore, the frequency of zygote males reflects perfectly the frequency of female gametes:

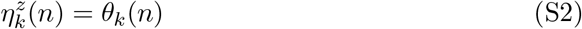
2. Individuals of both sex are under viability (or survival) selection. The frequencies of female and male adults are given by 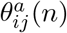 and 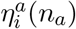, respectively:

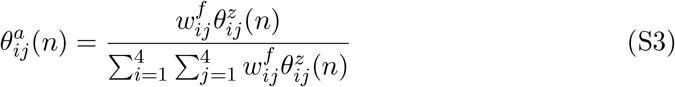

with 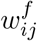 the fitness of the *ij* genotype. Note that there are no parental effects: 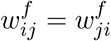; we maintain the distinction only for modeling convenience and:

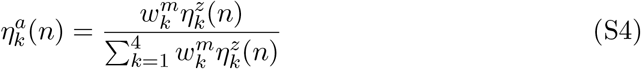

with 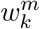 the fitness of haplotype *k* in males.
3. The last step corresponds to the production of gametes. As males are haploid, the gametes produced in the next generation reflect perfectly the distribution of adult males:

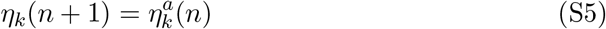 Adult females produce eggs. Recombination happens during this stage. The frequency of the female gametes of the next generation, 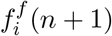, is given by:

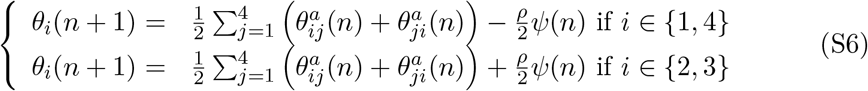

with 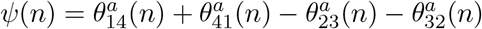. By combining the different equations detailed above (i.e., substituting 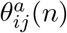 in equation (S6) and 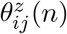 in (S3) for the female recursion and 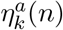 in (S5) as well as 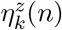 in (S4) for the male case), one obtains the following recursion when making the different fitness terms explicit:

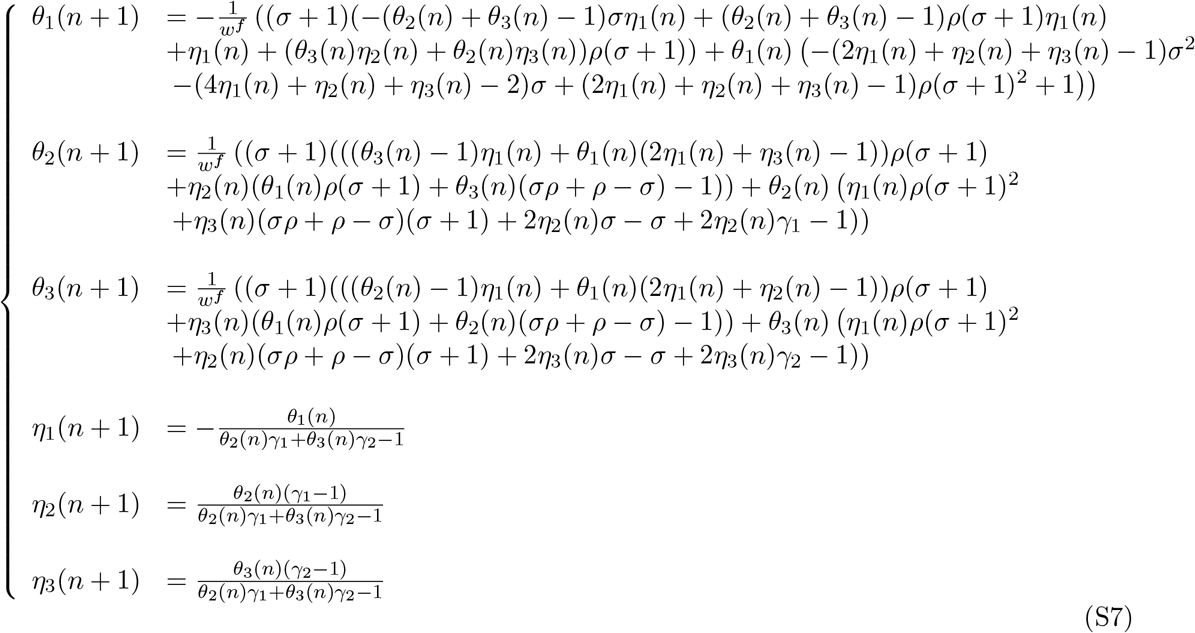

with *w^f^* the mean fitness of the female population:

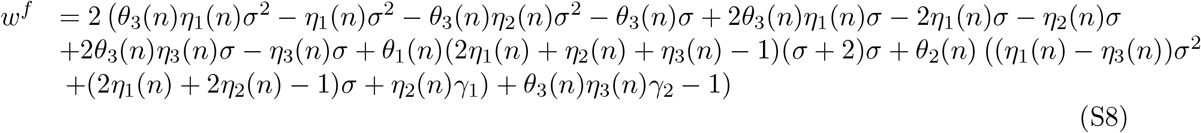

### Estimating genotype frequencies from a natural ant population

In order to compare our model with data from the natural, hybridizing Finnish ant population, we estimated the different genotype frequencies of parental *F. polyctena-like* and *F. aquilonia-like* individuals at pre-selection and post-selection life stages for males and females (Fig. S1(a)). We did not estimate the frequencies of introgressed or hybrid individuals. We used the genotype frequencies at different life-stages estimated in Kulmuni and Pamilo (2014) from nine microsatellite loci. For males, eggs were used to estimate pre-selection frequencies; the sum of adults and reproductive fathers was used to estimate post-selection frequencies. For females, eggs were used for pre-selection frequencies and the sum of young and old queens was used for post-selection frequencies. We used two different estimates for the number of parental females: individuals with exactly zero loci heterozygous for an introgressed allele, and individuals with one or more loci homozygous for the parental allele (i.e., the “diagnostic allele” in Kulmuni and Pamilo, 2014). In order to make these data comparable to our model, we rescaled the genotype frequencies such that 10.3% of the population is from the *F. polyctena-like* sub-population and 89.7% from the *F. aquilo-* nia-like sub-population, as estimated from the observed abundances of *F. polyctena-like* and *F. aquilonia-like* individuals from nests in the hybrid population collected between 1996–2012 (Table S1). Assuming that the natural population is at equilibrium, we fit the data (Table S2) to the model by calculating the sum of squared differences between the observed data and predicted equilibrium frequencies from 40600 parameter combinations.

## Supplementary Results

### Fitting the model to natural population frequencies

We compared the pre- and post-selection haplodiploid model (Fig. S1(a)) predictions with the estimated genotype frequencies of the natural, hybridizing *Formica* wood ant population for eggs and reproductive life-stages of males and females (Table S2). The model predictions from the best-fit models are shown in Figures S7 and S8. The best-fit models had parameter values corresponding to single-locus polymorphism or asymmetric coexistence, regardless of how the female frequencies were estimated (Fig. S9). Since these outcomes can occur at a variety of parameter combinations, we were not able to infer any specific parameter estimates other than that large values appear to be preferred for *γ*_1_ and recombination (Fig. S10–S13), consistent with the genomic architecture of the natural population, where multiple incompatibilities are likely to be spread across chromosomes (Kulmuni and Pamilo, 2014). Our model predicts less change in the genotype frequencies before vs. after selection as compared to the differential observed in the data for eggs vs. reproductive adults (Fig. S7(c) and S8(c)).

## Supplementary Discussion

We fitted our model to the data from the natural ant population described in Kulmuni and Pamilo (2014) and Table S1 in a rather crude approach. In the fitting procedure, we ignored that the data contain information from marker loci rather than the selected alleles, and we summarized the data in categories to resemble our case of a two-locus interaction. Our model fitting results indicate that the unequal ratio of *F. polyctena*-like and *F. aquilona-like* types that is observed in the natural population could represent a stable equilibrium of asymmetric coexistence. In fact, the high recombination rates among diagnostic alleles and strong prezygotic mechanisms producing within-group zygotes exhibited in the natural population (Kulmuni et al., 2010; Kulmuni and Pamilo, 2014) correspond with an area in the parameter space where asymmetric coexistence can be stably maintained over a wide range of values for female hybrid advantage.

Our model fit does not perform well at predicting the number of introgressed and hybrid females in the population. We were not able to estimate the population frequencies for introgressed and hybrid females with data from Kulmuni and Pamilo (2014), but we know from Kulmuni et al. (2010) that the vast majority of both *F. polyctena-like* and *F. aquilonia-like* females exhibit some introgression. Contrary to this observation in the natural population, our model fit predicts that introgressed *F. polyctena-like* females should be rare (< 15%) and that pure *F. aquilona-like* females should be only slightly less common than the introgressed *F. polyctena-like* females (Fig. S8).

## Supplementary Tables

**Table S1:**
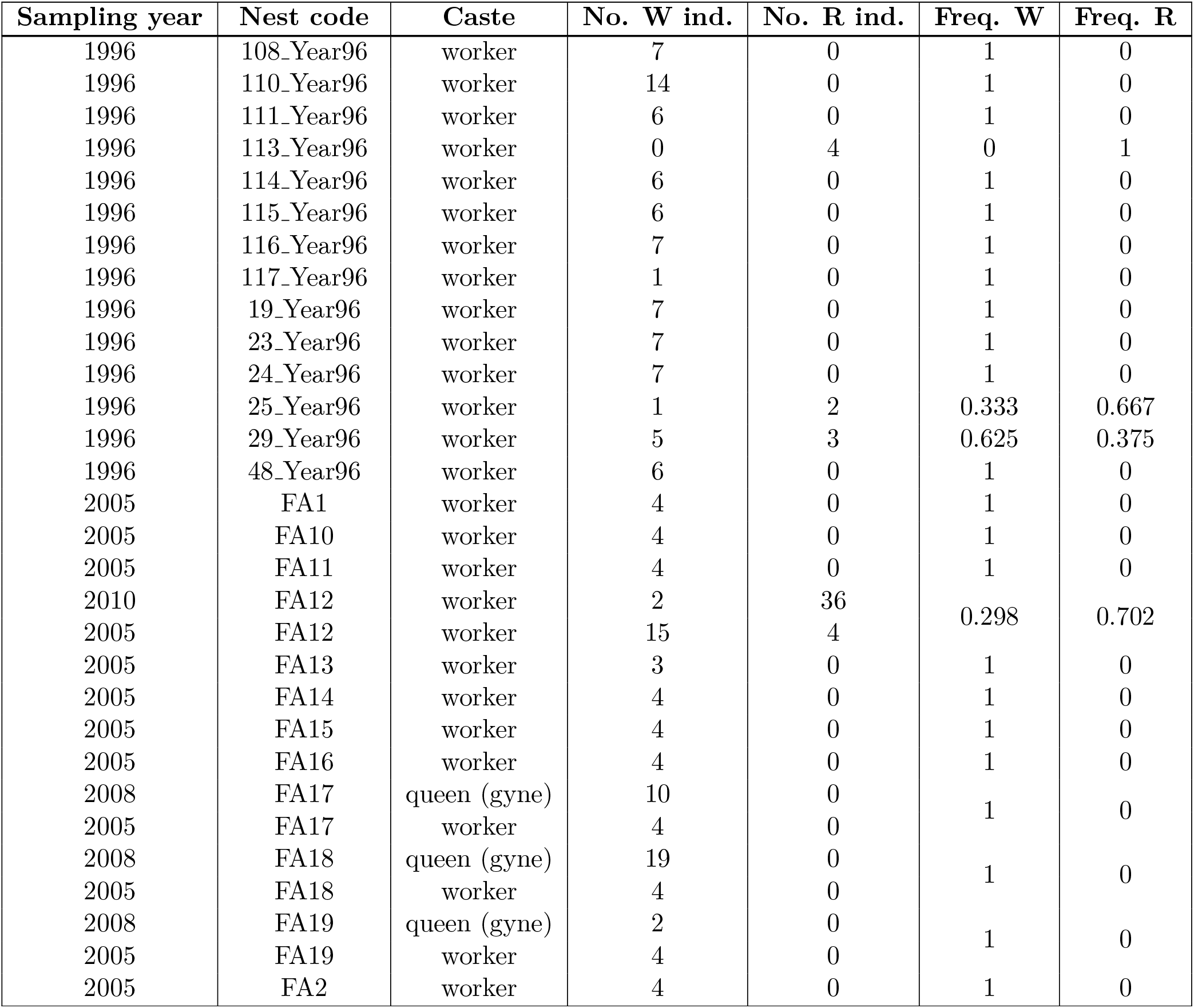
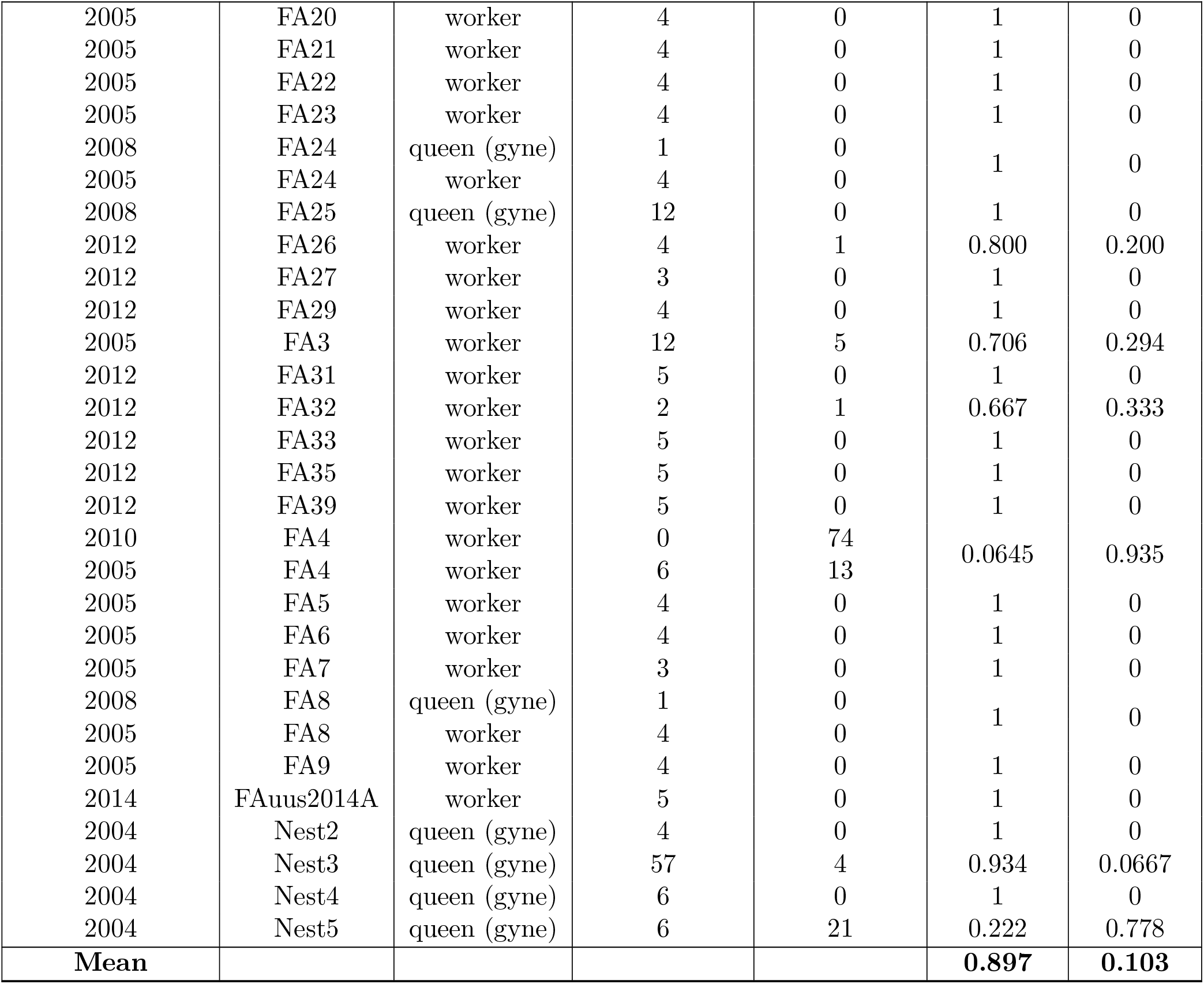
Table of adult frequencies for ant colonies in Långholmen in southern Finland. Individuals were genotyped and assigned to W- or R-type, where W indicates *F. aquilonia-* like parental individuals and R indicates *F. polyctena-like* parental individuals. Mean frequencies of *F. aquilonia-like* and *F. polyctena-like* types were calculated across castes and sampling years.

**Table S2:** Table of estimated genotype frequencies used for data fitting. This data was calculated using the colony data from Table S1 above and the within-group genotype frequencies from Tables 1 & S7 of Kulmuni and Pamilo (2014). Following this same reference, “Introgressed” females refers to indviduals inferred to be purebred based on having no loci heterozygous for introgressed alleles. “Diagnostic” females refers to individuals inferred to be purebred based on having at least one locus homozygous for diagnostic alleles.

| Males |  |  |
| --- | --- | --- |
| “Types” | Before selection (Eggs) | After selection (Reproductive males) |
| Parental *F. aquilonia*-like | 0.637 | 0.828 |
| Parental *F. polynecta*-like | 0.0412 | 0.102 |
| Summed hybrids | 0.322 | 0.0701 |

| “Introgressed” Females |  |  |
| --- | --- | --- |
| “Types” | Before selection (Eggs) | After selection (Young and old queens) |
| Parental *F. aquilonia*-like | 0.511 | 0.424 |
| Parental *F. polynecta*-like | 0.0216 | 0 |
| Summed all others | 0.467 | 0.576 |

| “Diagnostic” Females |  |  |
| --- | --- | --- |
| “Types” | Before selection (Eggs) | After selection (Young and old queens) |
| Parental *F. aquilonia*-like | 0.314 | 0.189 |
| Parental *F. polynecta*-like | 0.0834 | 0.00214 |
| Summed all others | 0.603 | 0.809 |

**Table S3:** Table that demonstrates why single-locus polymorphism can be invaded in diploids. This serves to explain why single-locus polymorphism is never stable in diploids.

| Genotype | Mutant | Fitness of the mutant / fitness of the mutated genotype |
| --- | --- | --- |
| $A_+ A_+ B_+ B_+$ | $A_- A_+ B_+ B_+$ | $(1 + \sigma) > 1$ |
| $A_+ A_+ B_+ B_-$ | $A_- A_+ B_+ B_-$ | $(1 + \sigma) > 1$ |
| $A_+ A_+ B_- B_-$ | $A_- A_+ B_- B_-$ | $(1 + \sigma)/(1 - \gamma_1) > 1$ |
| $A_+ B_+$ | $A_- B_+$ | $(1 - \gamma_2) < 1$ |
| $A_+ B_-$ | $A_- B_-$ | $1/(1 - \gamma_1) > 1$ |

## Supplementary Figures

**Figure S1:**
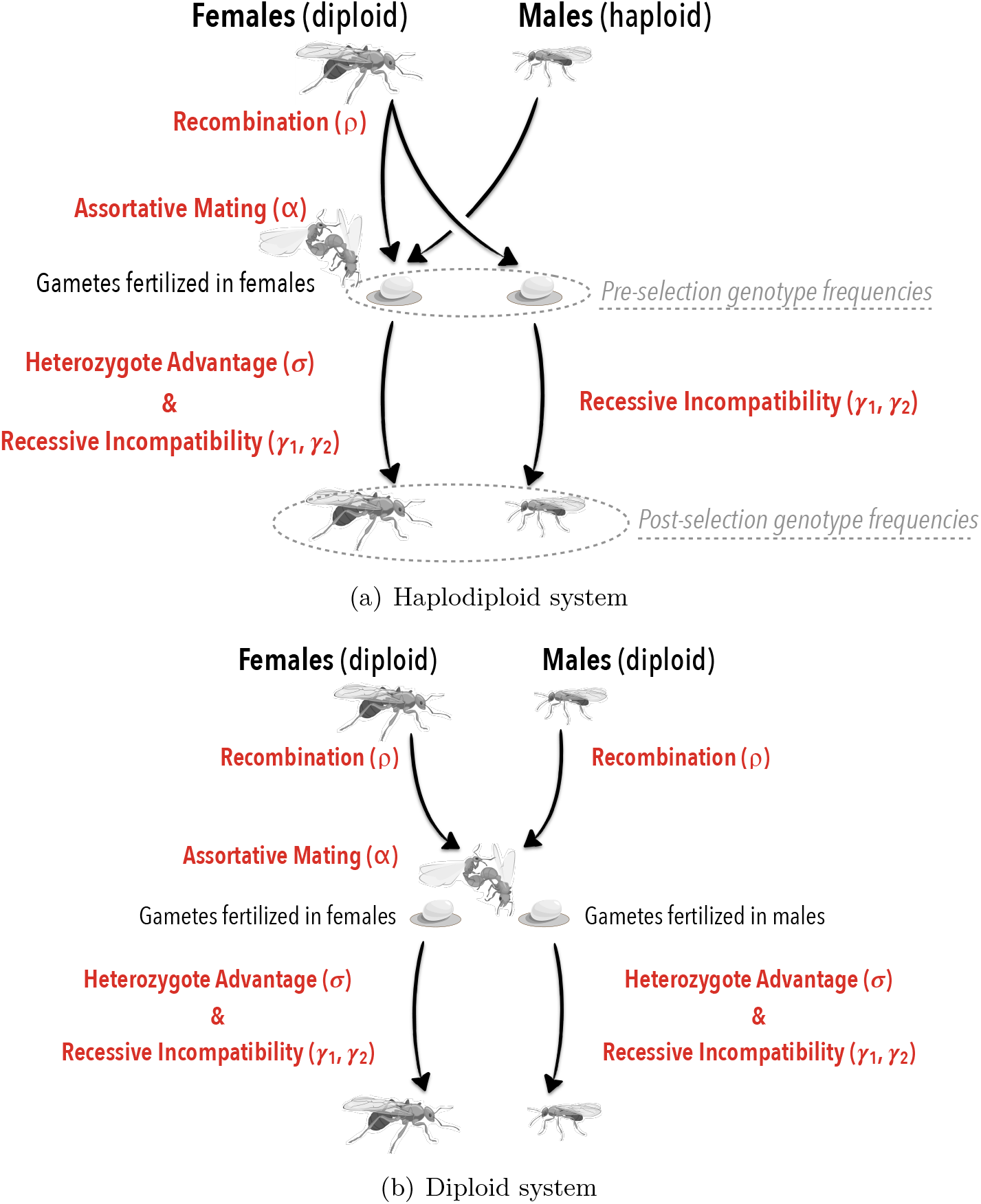
Panel (a) shows the lifecycle of haplodiploids, panel (b) shows the lifecycle of diploids. Events that occur in either females and/or males are illustrated separately on the left and right sides, respectively, in order to highlight the differences between the lifecycles. The pre- and post-selection genotype frequencies for haplodiploids are indicated because they were used for model fitting.

**Figure S2:**
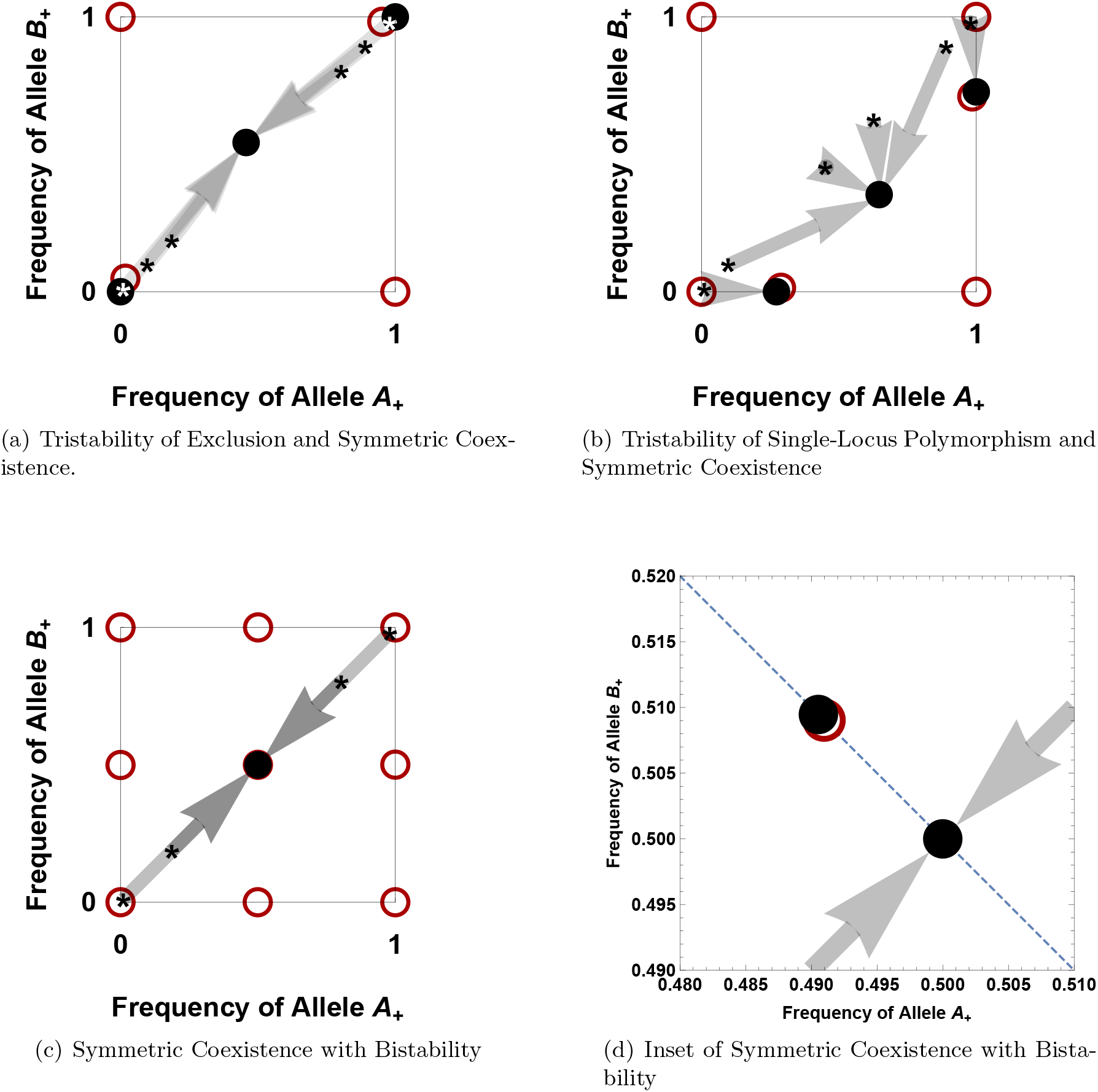
Phase-plane diagrams of evolutionary scenarios in the haplodiploid model that are not discussed in the main text. Figures are as described in figure 3 except that unstable equilibia are red. Panel (a): *σ* = 0.05, *γ*_1_ = 0.2, *γ*_2_ = 0.11, *ρ* = 0.099, *α* = 0. Panel (b): *σ* = 0.143, γ_1_ = 0.46, *γ_2_* = 0.083, *ρ* = 0.455, *α* = 0. Panel (c): *σ* = 0.2, γ_1_ = 0.019, *γ*_2_ = 0.011, *ρ* = 0.000001, *α* = 0. Panel (d): inset of panel (c) with a dashed line at *y* = 1 – *x*.

**Figure S3:**
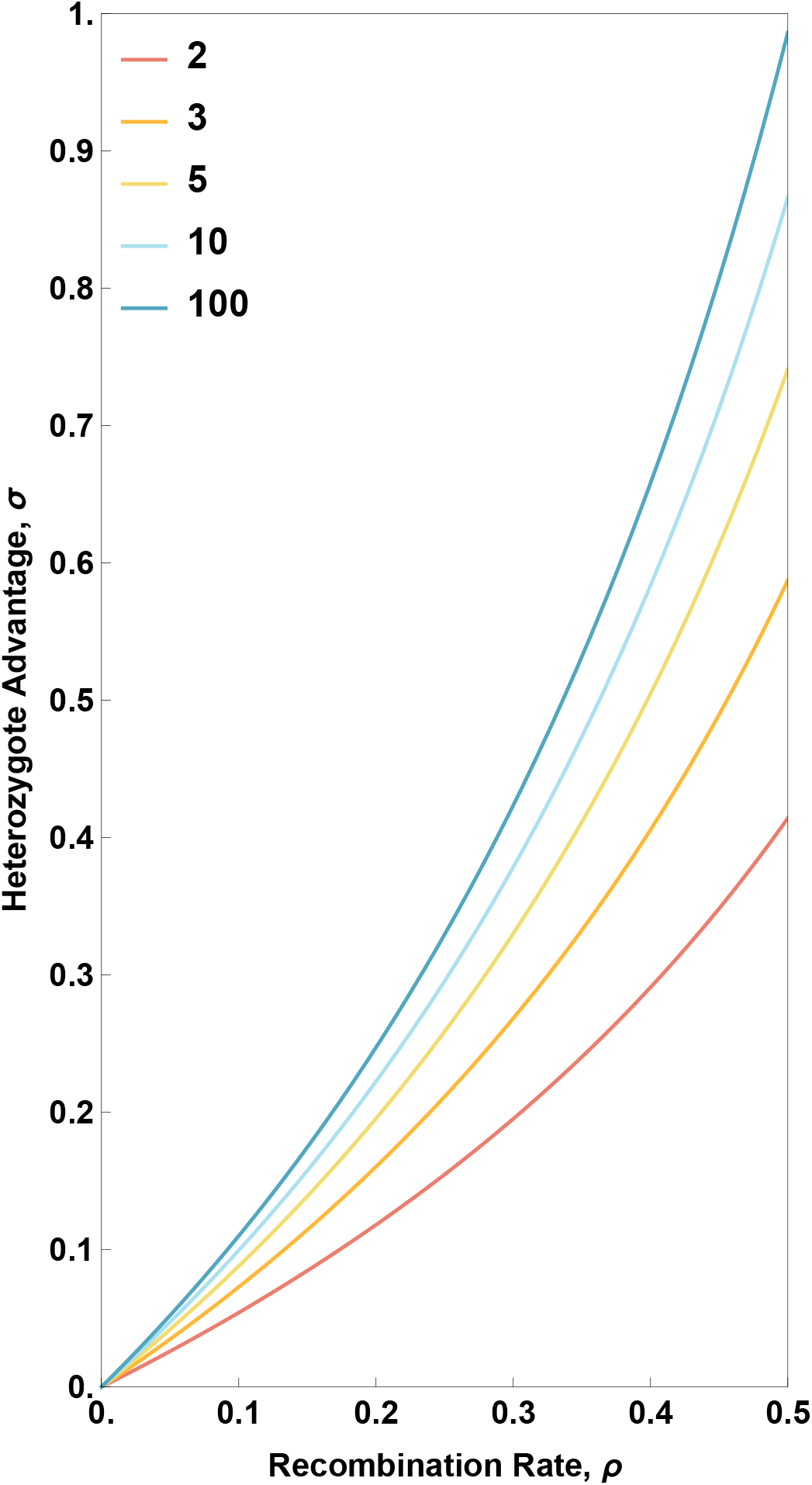
Conditions for stability of exclusion in the network case for haplodiploids with an increasing number of loci, as indicated in the legend. Exclusion represents a locally stable equilibrium for any parameter combination below the curve.

**Figure S4:**
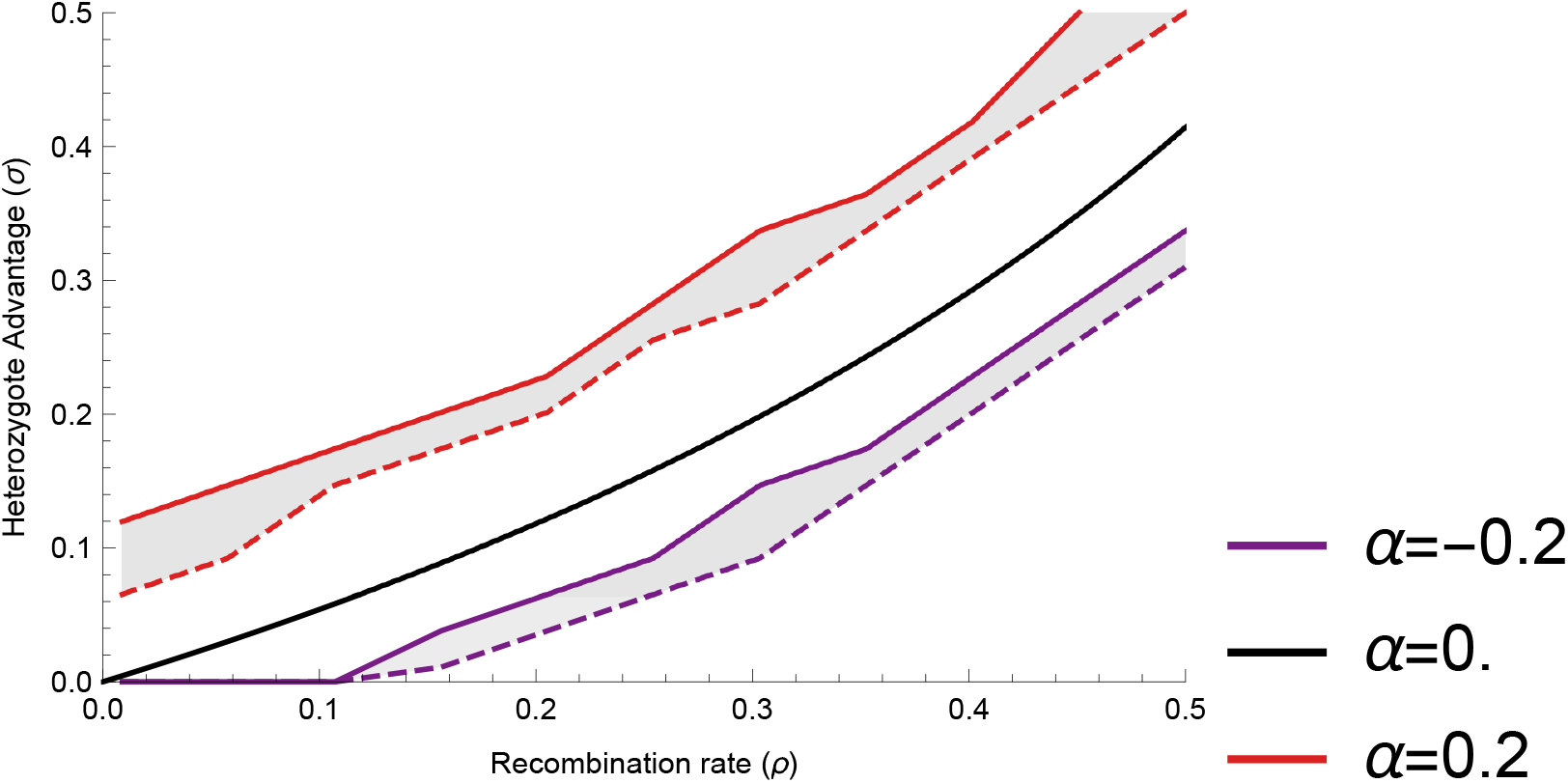
Plot of recombination (*ρ*) vs heterozygote advantage (*σ*) for lethal epistasis (*γ*_1_ = *γ*_2_ = 1) and varying strength of assortment, *α*. Solid lines show values at and above which symmetric coexistence occurs; dashed lines show values at and below which exclusion occurs; the shaded region between the solid and dashed lines shows values where asymmetric coexistence occurs. The result random mating (*α* = 0) is given in equation 7 and so the black line is exactly the same as figure 4(a). This figure was generated with a precision of only ≈ 0.05 units of *σ*.

**Figure S5:**
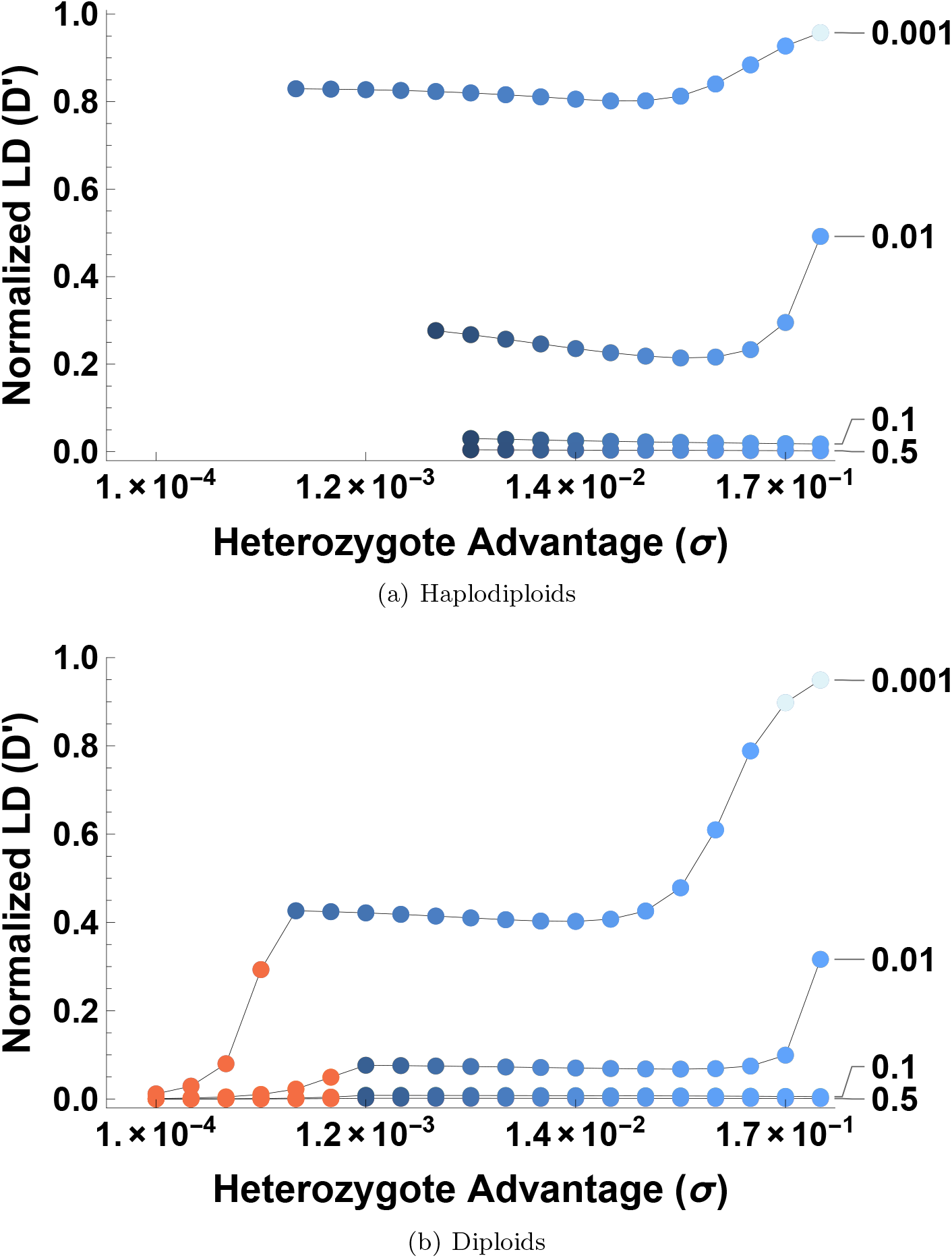
The effect of heterozygote advantage on the normalized linkage disequilibrium 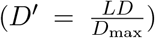 for (a) haplodiploids and (b) diploids under varying recombination rates (values for *ρ* indicated at right of plot). Each point is shaded according to its evolutionary outcome following the color-scheme in figure 6. Normalization of LD is required since allele frequencies at the two loci can vary considerably (see shading of blue points in the plots above and in figure 6A). No values are shown for the single-locus polymorphism or exclusion outcomes because LD cannot exist when polymorphism is lost at one or both loci. (*γ*_1_ = 0.01, *γ*_2_ = 0.002, *α* = 0).

**Figure S6:**
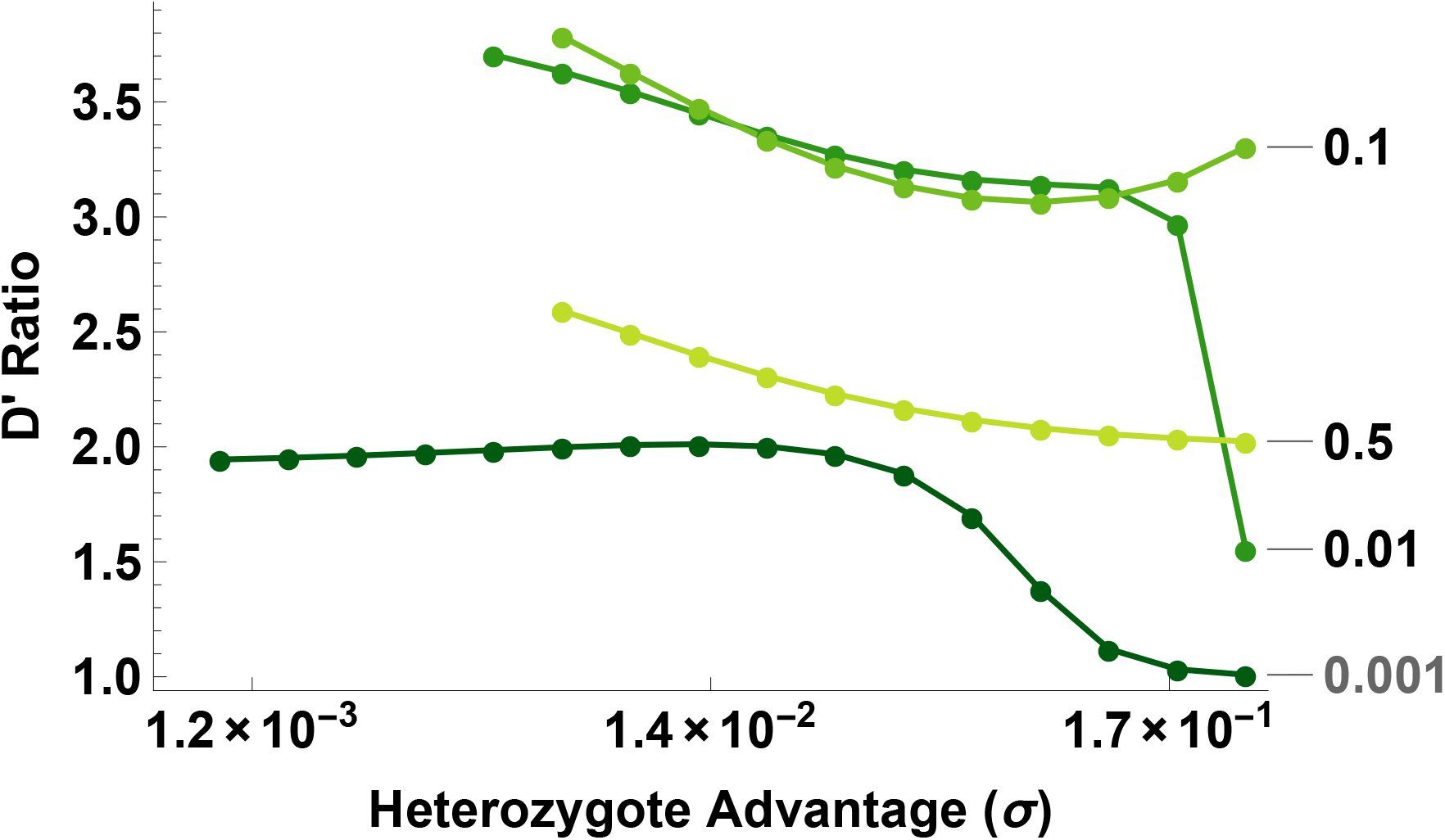
The effect of heterozygote advantage on the ratio of normalized linkage disequilibrium for haplodiploids over diploids 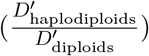 under varying recombination rates. A diploids ratio greater than 1 indicates more LD in haplodiploids; when polymorphism is maintained at both alleles, haplodiploids always have more or equal LD to diploids. Intermediate recombination values maximize the ratio. (*γ*_1_ = 0.01, *γ*_2_ = 0.002, *α* = 0)

**Figure S7:**
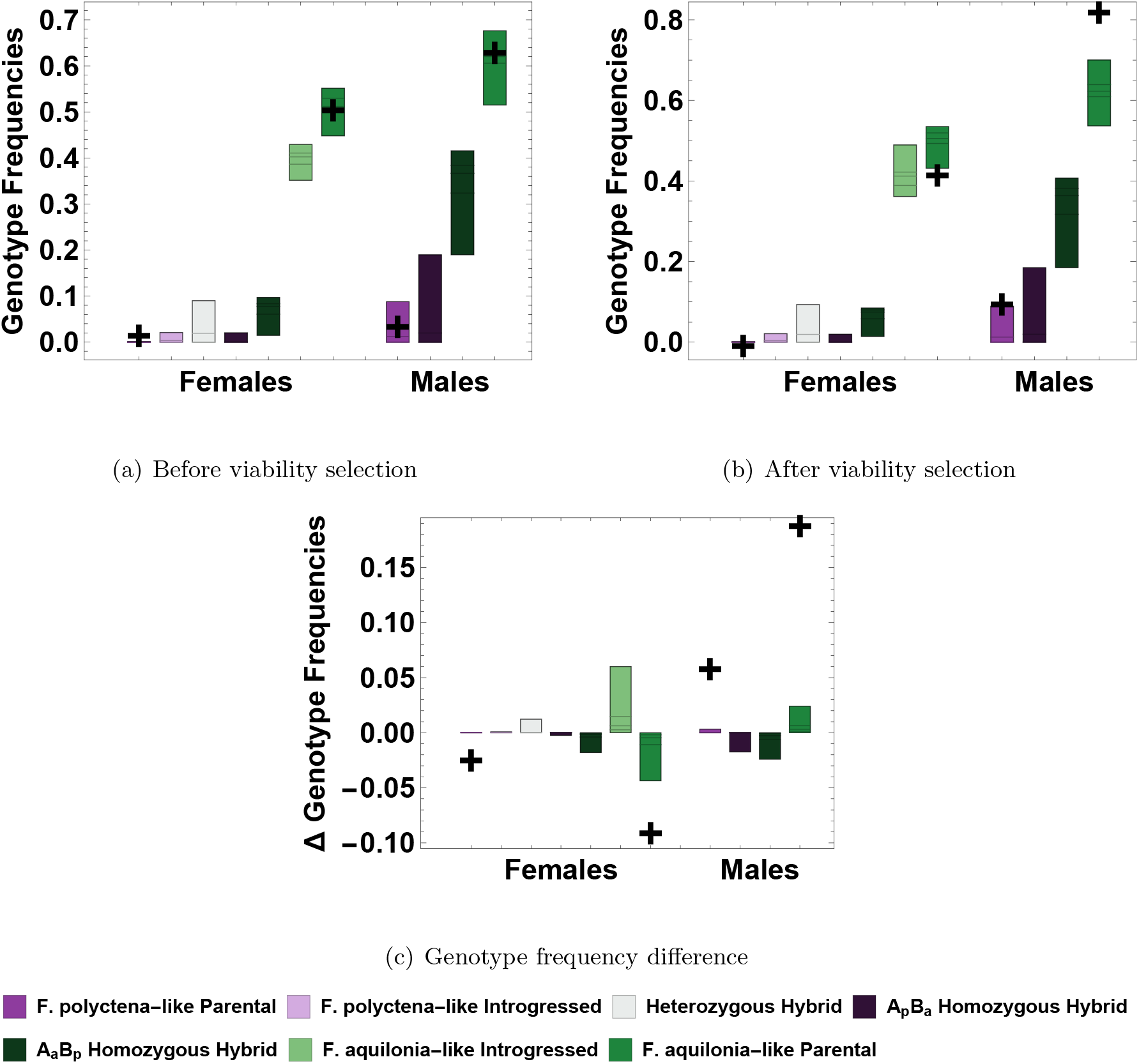
Comparison of model predictions (boxplots) to the data used for fitting the model (+) shows that the model is able to capture the high frequency of *F. aquilona-* like alleles (green shades) in the population. Boxplots show the genotype frequencies for females and males that are predicted from the distribution of the best fitting models. Model predictions before selection, after selection, and delta selection (after selection − before selection). For “introgressed markers” data with random mating (*α* = 0).

**Figure S8:**
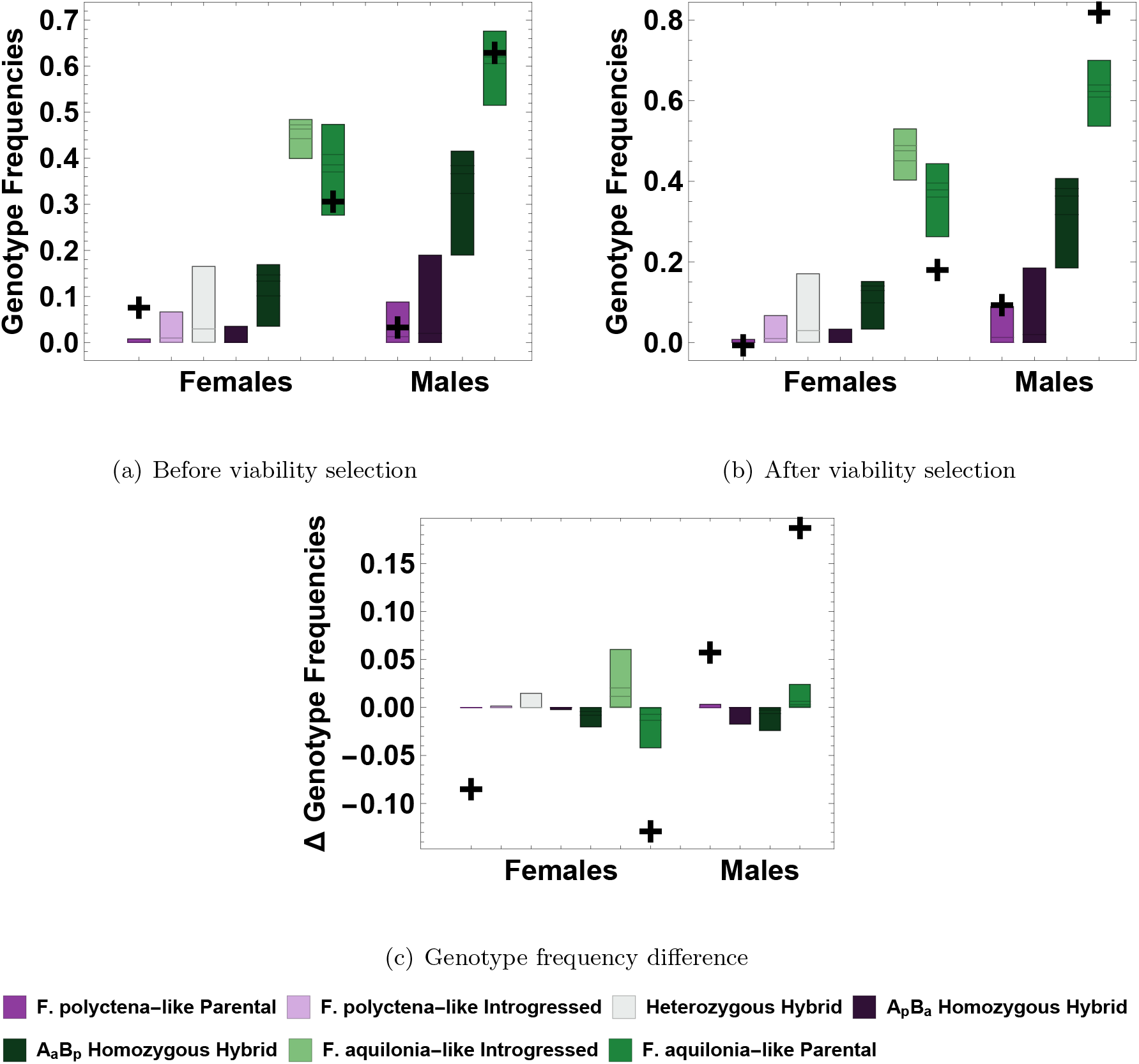
Comparison of model predictions (boxplots) to the data used for fitting the model (+) shows that the model is able to capture the high frequency of *F. aquilona*-like alleles (green shades) in the population. Boxplots show the genotype frequencies for females and males that are predicted from the distribution of the best fitting models. In this case parental genotype frequencies (shown on plot as +) are estimated using individuals with one or more loci homozygous for the parental allele (“diagnostic markers” data). Model predictions before selection, after selection, and delta selection (after selection – before selection). For “diagnostic markers” data with random mating (*α* = 0).

**Figure S9:**
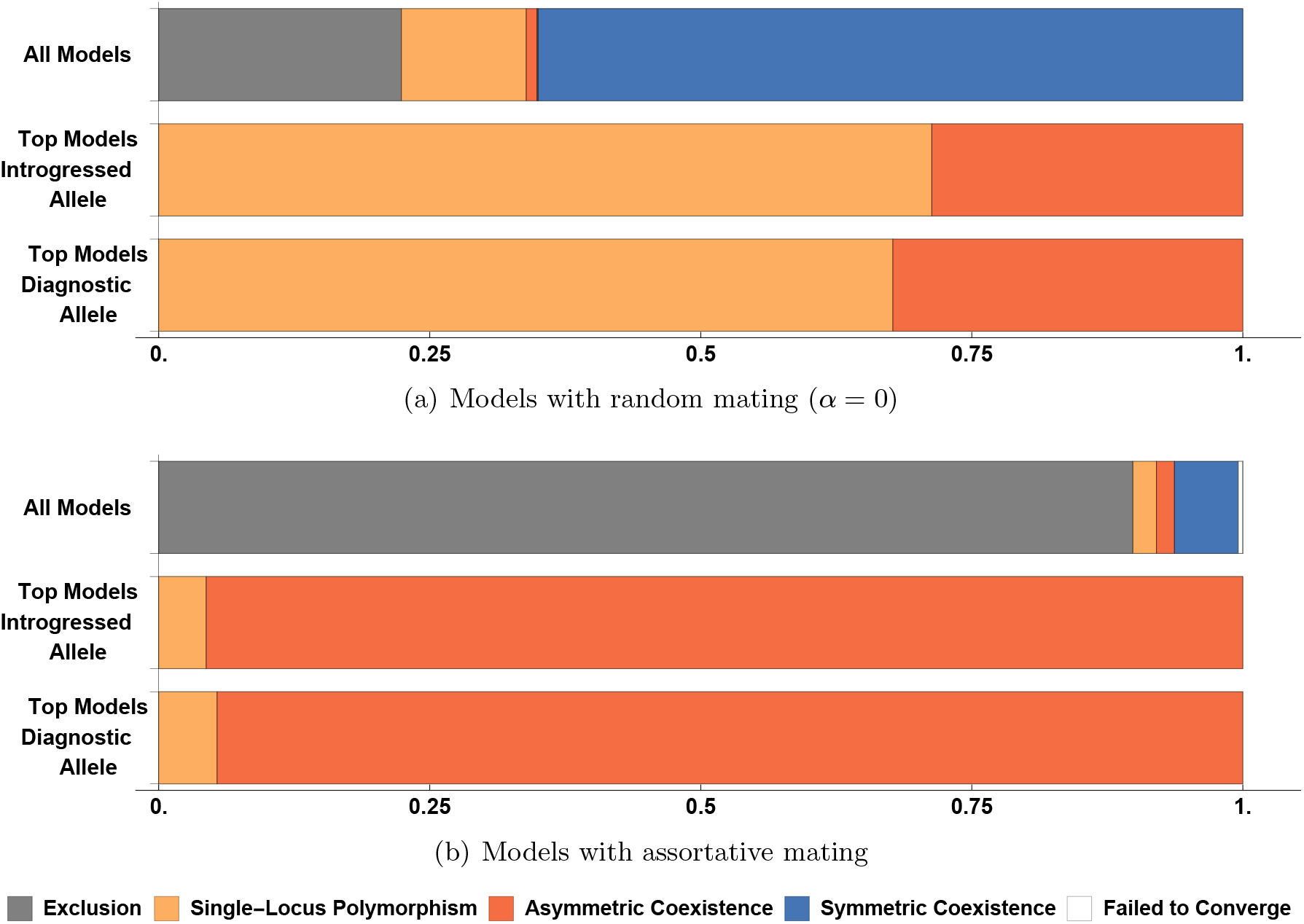
Bar plots show the proportion of parameter combinations that correspond to different evolutionary outcomes. “All models” is for all parameter combinations investigated, “Top models introgressed allele” is for model fitting to the “introgressed markers” data, and “Top models diagnostic allele” is for model fitting to the “diagnostic markers” data.

**Figure S10:**
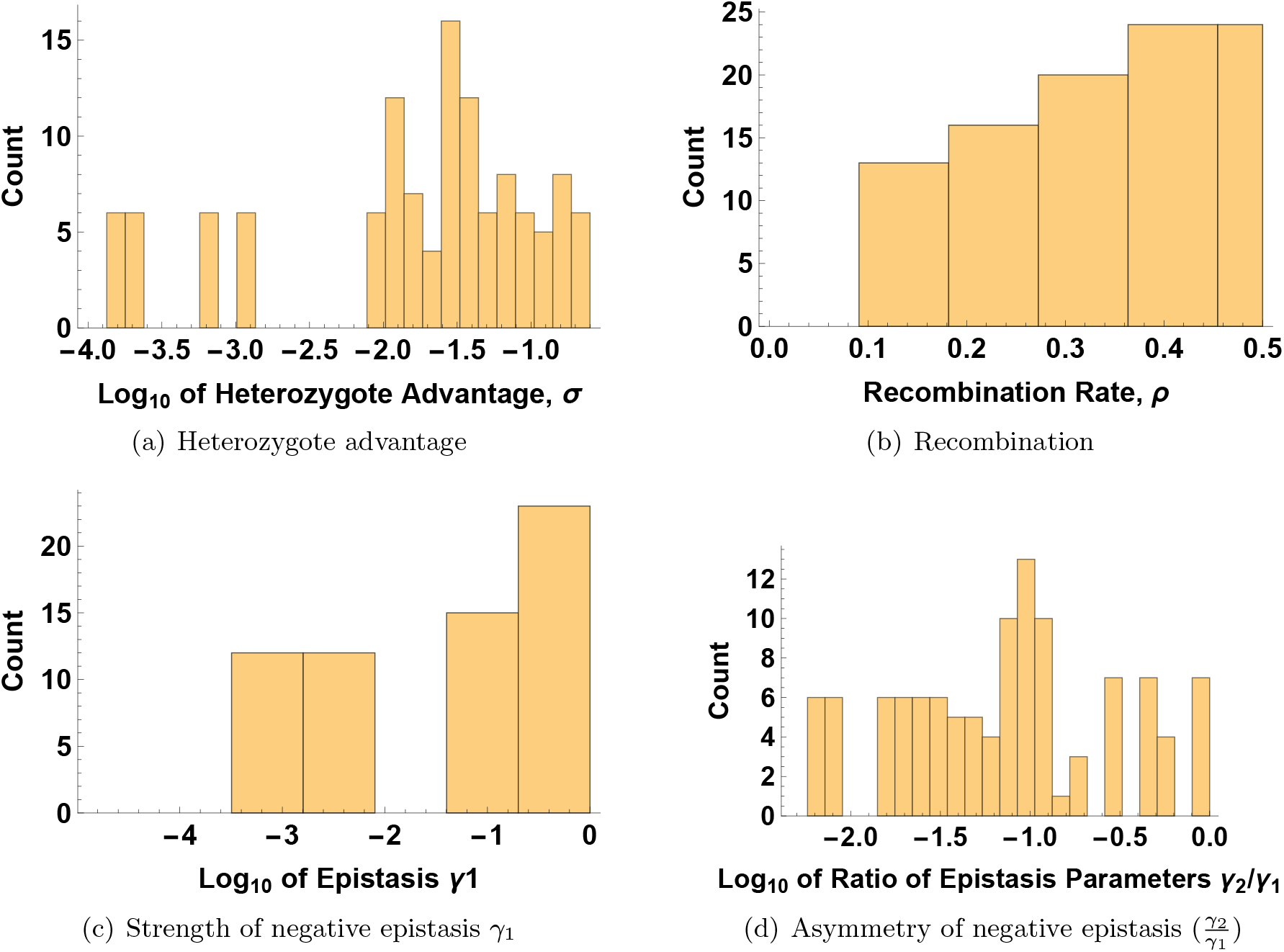
Distribution of the estimated parameter values for the model fitting “introgressed markers” data with random mating (*α* = 0).

**Figure S11:**
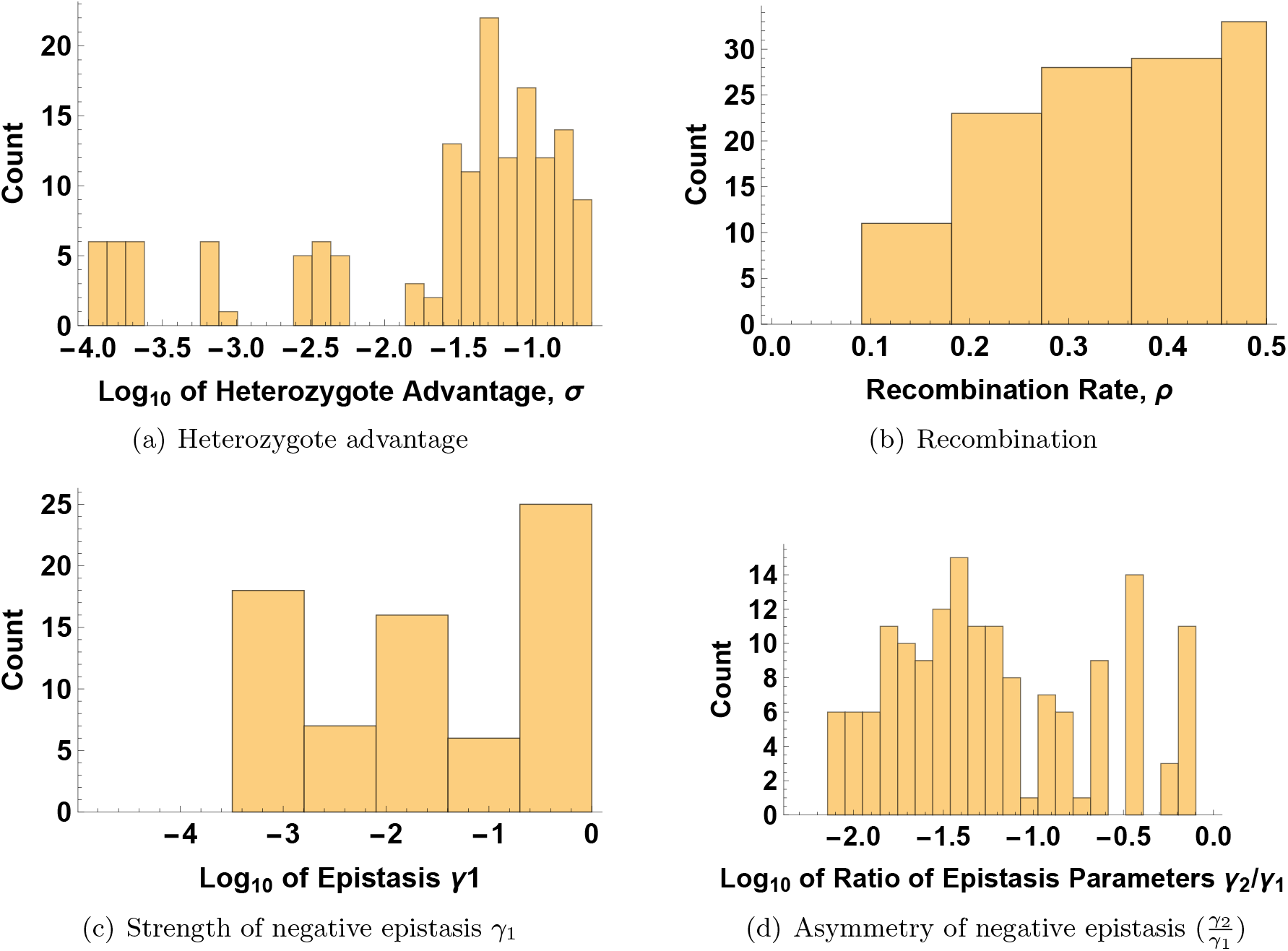
Distribution of the estimated parameter values for the model fitting “diagnostic markers” data with random mating (*α* = 0).

**Figure S12:**
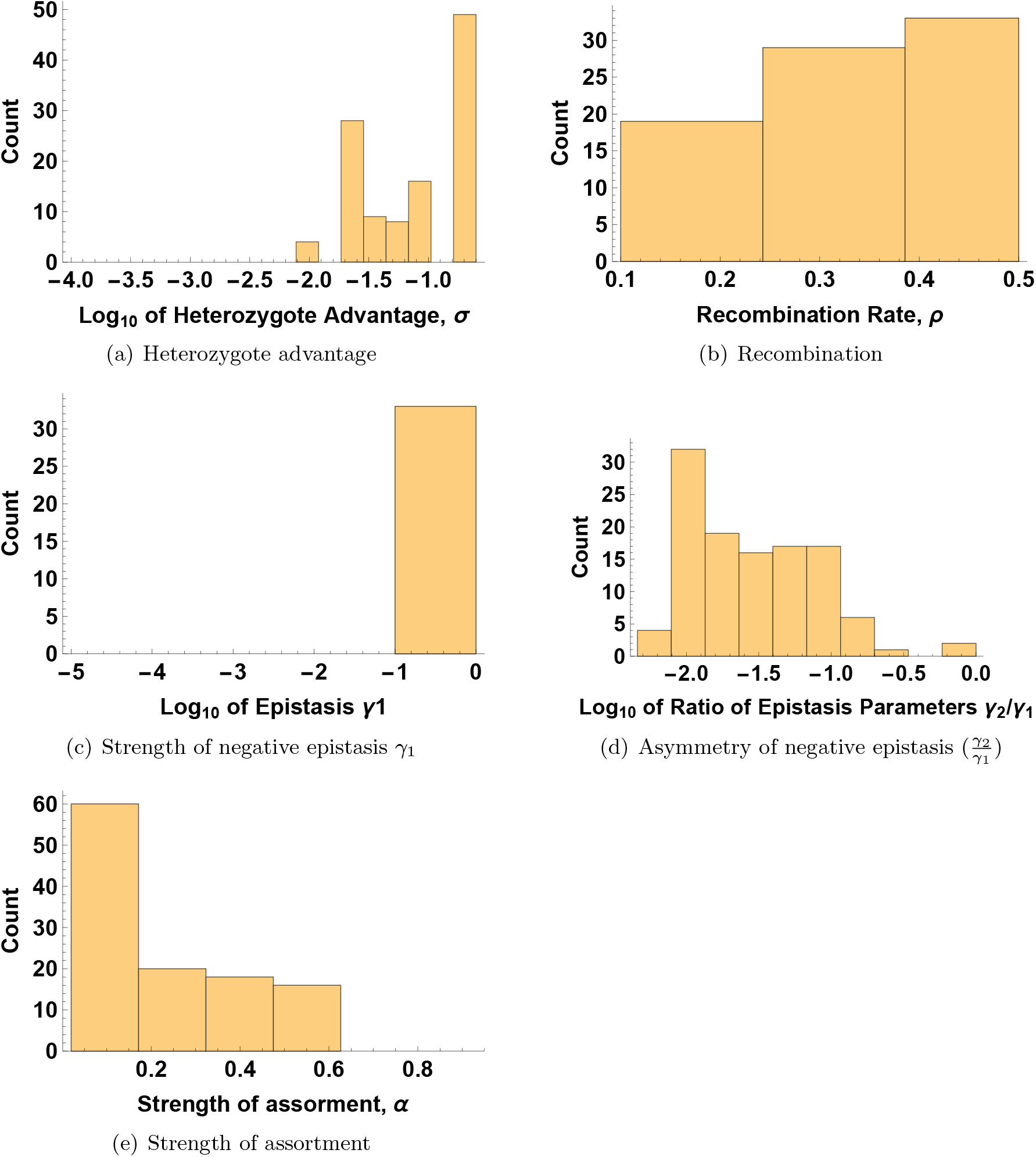
Distribution of the estimated parameter values for the model fitting “introgressed markers” data with assortative mating.

**Figure S13:**
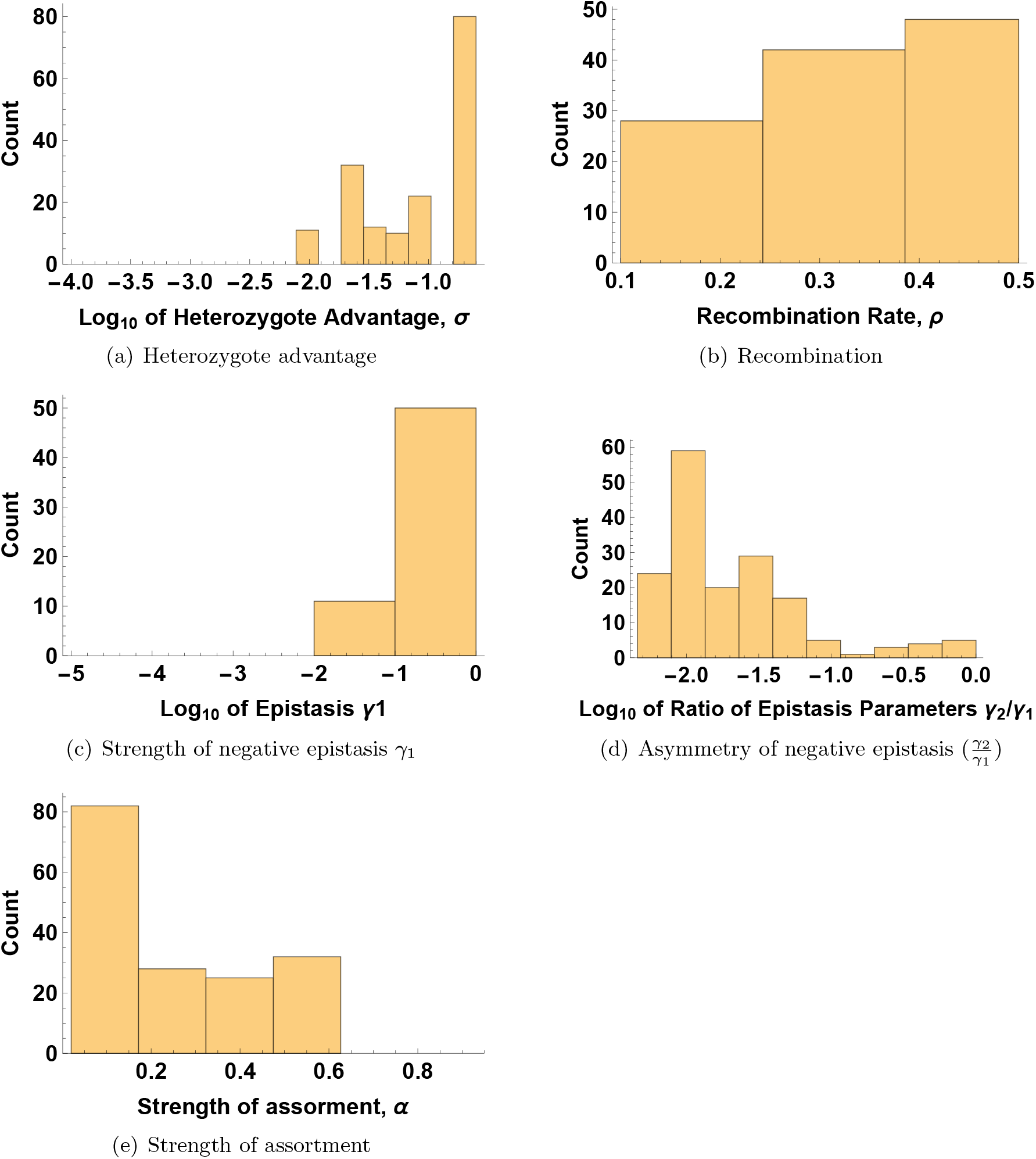
Distribution of the estimated parameter values for the model fitting “diagnostic markers” data with assortative mating.

## Mathematica Online Supplement

The executable version of this file is available here: https://www.dropbox.com/s/w6jcx30im5t5597/Ant-hybrids_Mathematica_file_for_re-submission.nb?dl=0

**Author contributions for coding and analysis**

Chapter 1: CB

Chapter 2: AHG with help from AB

Chapter 3: CB

Chapter 4: AB

Chapter 5: AHG with help from CB, AB

Chapter 6: AHG

### Chapter 1: Specifying the model

#### ▀ Dynamics without assortment

##### Life cycle for discrete dynamics & multiplicative selection

Before we begin, here is a handy function to improve display of lists:

~~~
grid[tab_] := Grid[tab, ItemSize → Full] (* this way it pretends that
 the document is of infinite width and doesn’t break lines in tables *)
~~~

Assume that we have female gamete frequencies

~~~
gamF = {g1, g2, g3, g4};
~~~

and adult male frequencies

~~~
hapM = {h1, h2, h3, h4};
~~~

These create the next generation of female genotypes

~~~
genF = Table [gamF [[i]] hapM [[j]], {i, 4}, {j, 4};] // Flatten
{g1 hi, g1h2, g1h3, g1h4, g2h1, g2h2, g2h3,
 g2h4, g3h1, g3h2, g3h3, g3 h4, g4h1, g4h2, g4h3, g4 h4}
~~~

and male genotypes/haplotypes (which are simply the female gamete frequencies).

~~~
genM = gamF
 {g1, g2, g3, g4}
~~~

At first we will write down the model for general epistasis, but then study asymmetric recessive epistasis throughout the manuscript.

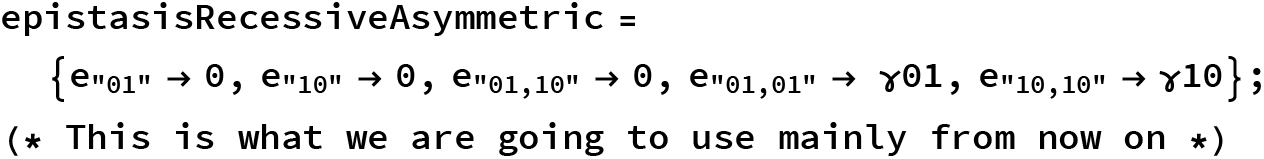

###### Viability selection in males

Male frequencies are entirely determined by female gamete frequencies in the previous generation.

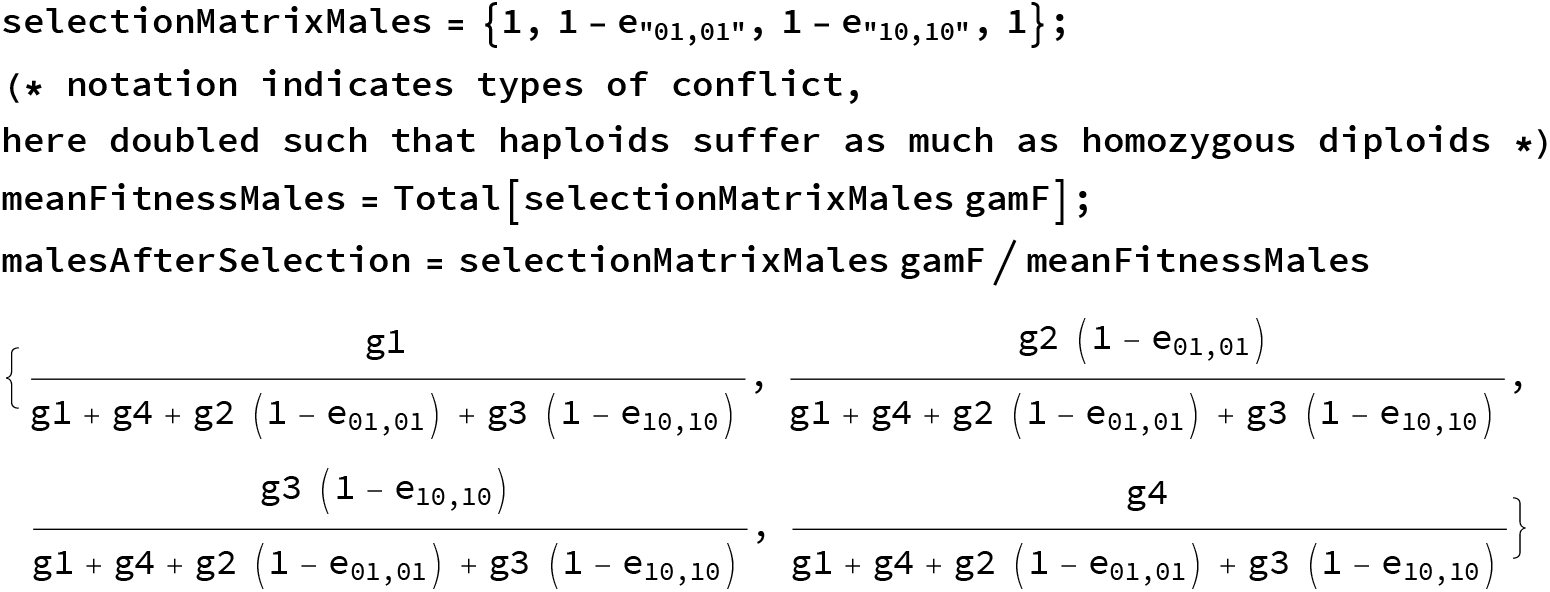

###### Viability selection in females, based on genotypes

To allow for strong selection, this scheme should be multiplicative. Also, we can integrate different types of epistasis. This is the general form:

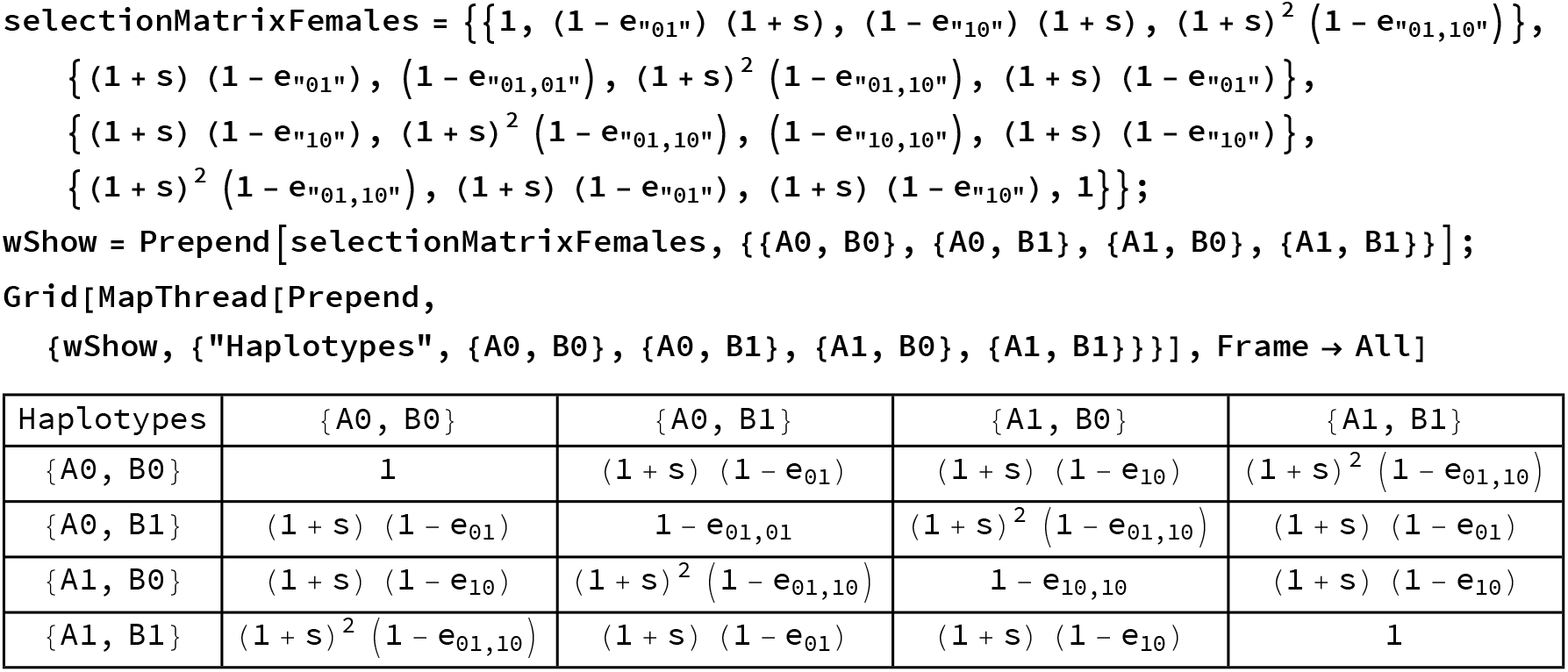

Selection of genotypes happens as follows

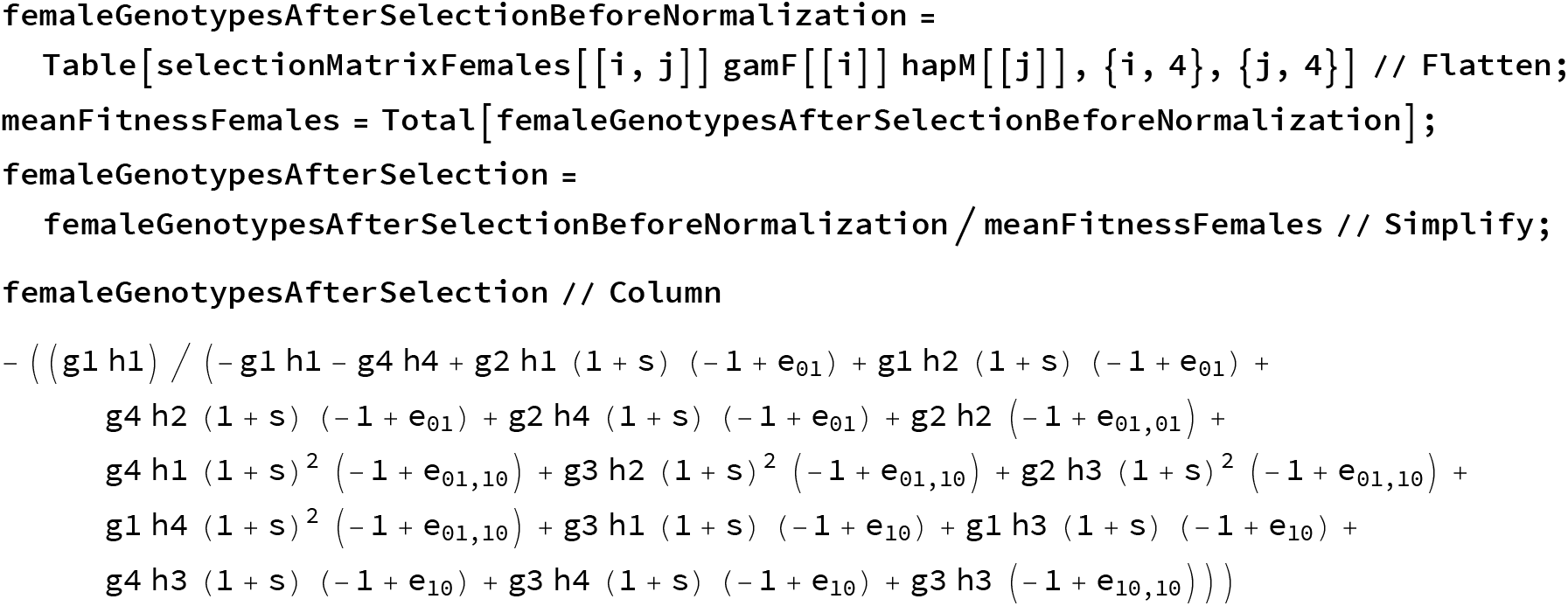
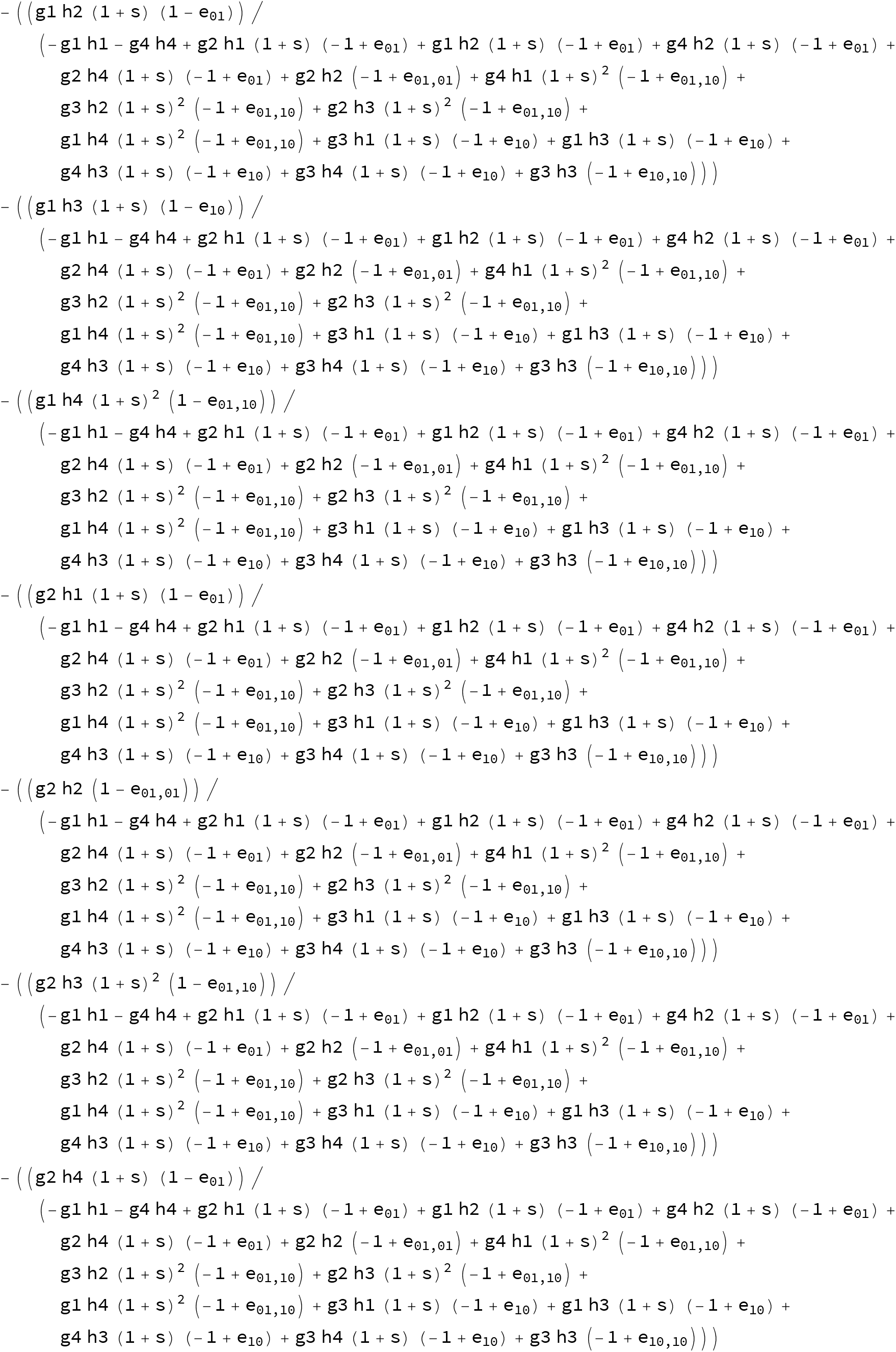
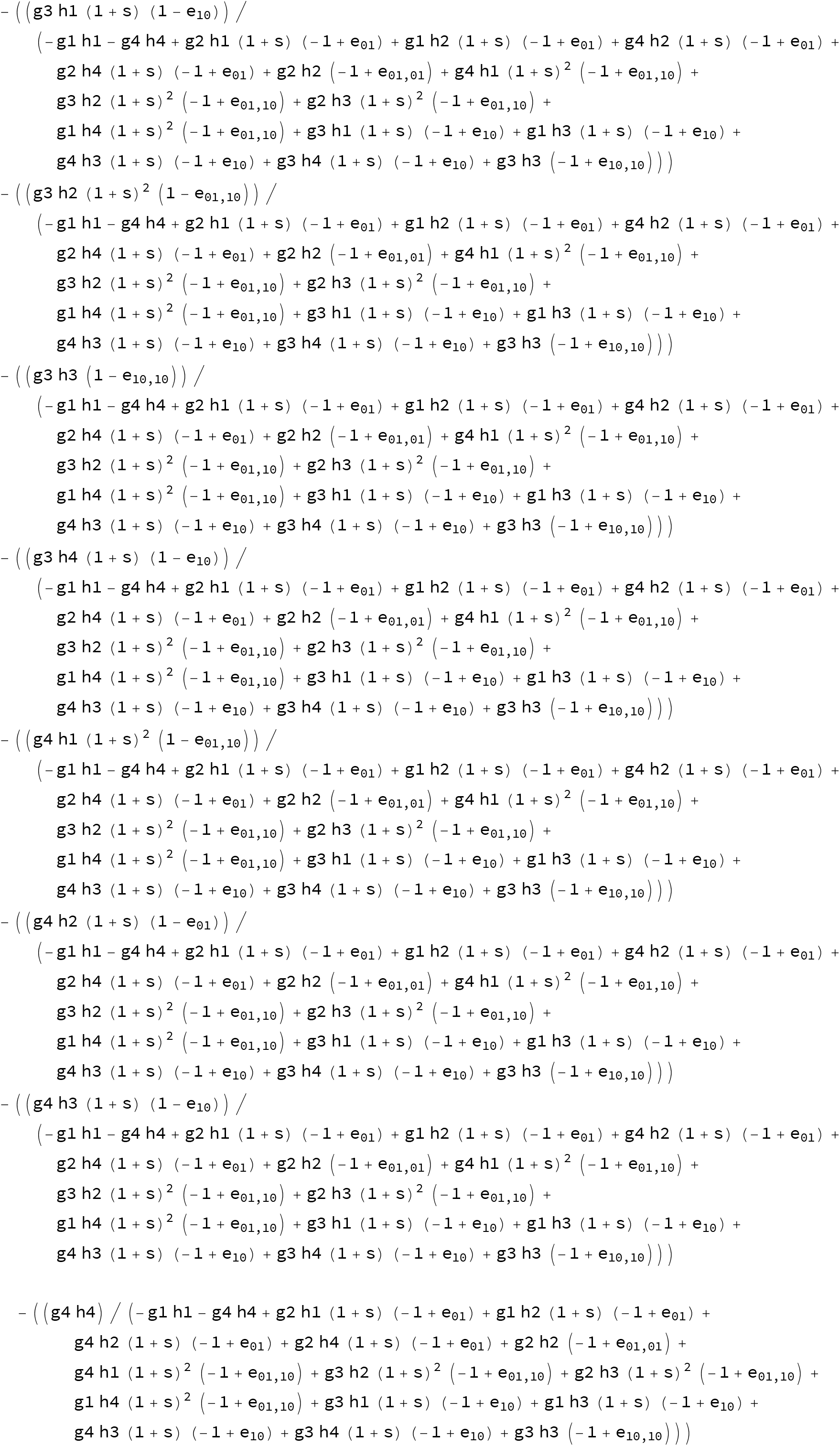

###### Recombination

Define different haplotypes and genotypes

~~~
haplotypes = Tuples[{0, 1}, 2]
genotypes = Tuples [haplotypes, 2]
{{0, 0}, {0, 1}, {1, 0}, {1, 1}}
{{{0, 0}, {0, 0}}, {{0, 0}, {0, 1}}, {{0, 0}, {1, 0}}, {{0, 0}, {1, 1}},
   {{0, 1}, {0, 0}}, {{0, 1}, {0, 1}}, {{0, 1}, {1, 0}}, {{0, 1}, {1, 1}},
   {{1, 0}, {0, 0}}, {{1, 0}, {0, 1}}, {{1, 0}, {1, 0}}, {{1, 0}, {1, 1}},
   {{1, 1}, {0, 0}}, {{1, 1}, {0, 1}}, {{1, 1}, {1, 0}}, {{1, 1}, {1, 1}}}
~~~

Define general frequency vectors

~~~
genFemale = Table[genotype_k_, {k, genotypes}]
gametes = Table[gamete_k_, {k, haplotypes}]
{genotype_{{0,0},{0,0}}_, genotype_{{0,0},{0,1}}_, genotype_{{0,0},{1,0}}_, genotype_{{0,0},{1,1}}_,
  genotype_{{0,1},{0,0}}_, genotype_{{0,1},{0,1}}_, genotype_{{0,1},{1,0}}_, genotype_{{0,1},{1,1}}_,
 genotype_{{1,0},{0,0}}_, genotype_{{1,0},{0,1}}_, genotype_{{1,0},{1,0}}_, genotype_{{1,0},{1,1}}_,
 genotype_{{1,1},{0,0}}_, genotype_{{1,1},{0,1}}_, genotype_{{1,1},{1,0}}_, genotype_{{1,1},{1,1}}_}
{gamete_{0,0}_, gamete_{0,1}_, gamete_{1,0}_, gamete_{1,1}_}
~~~

Now make gametes out of genotypes after selection

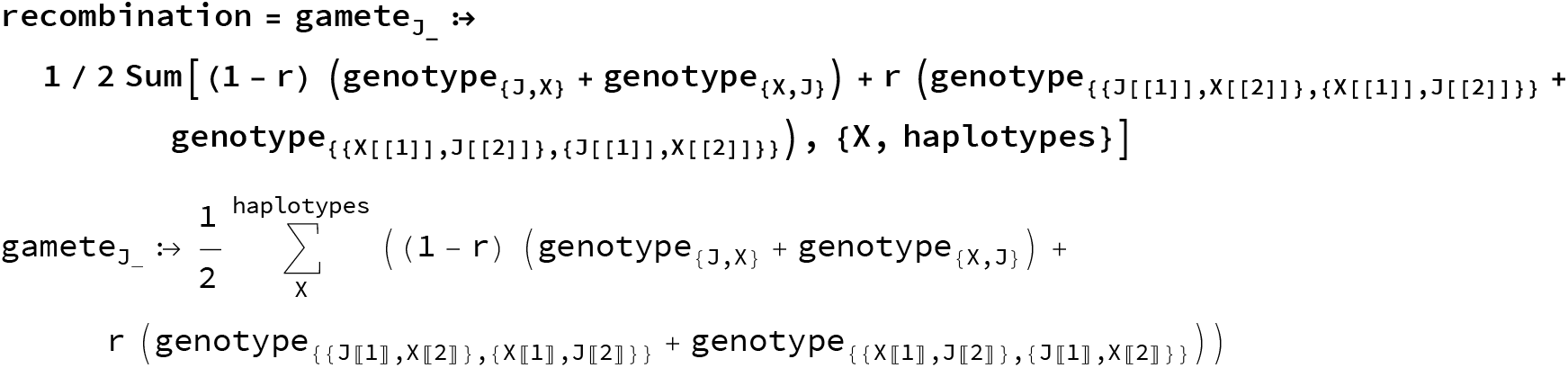

Here is the printout of what happens:

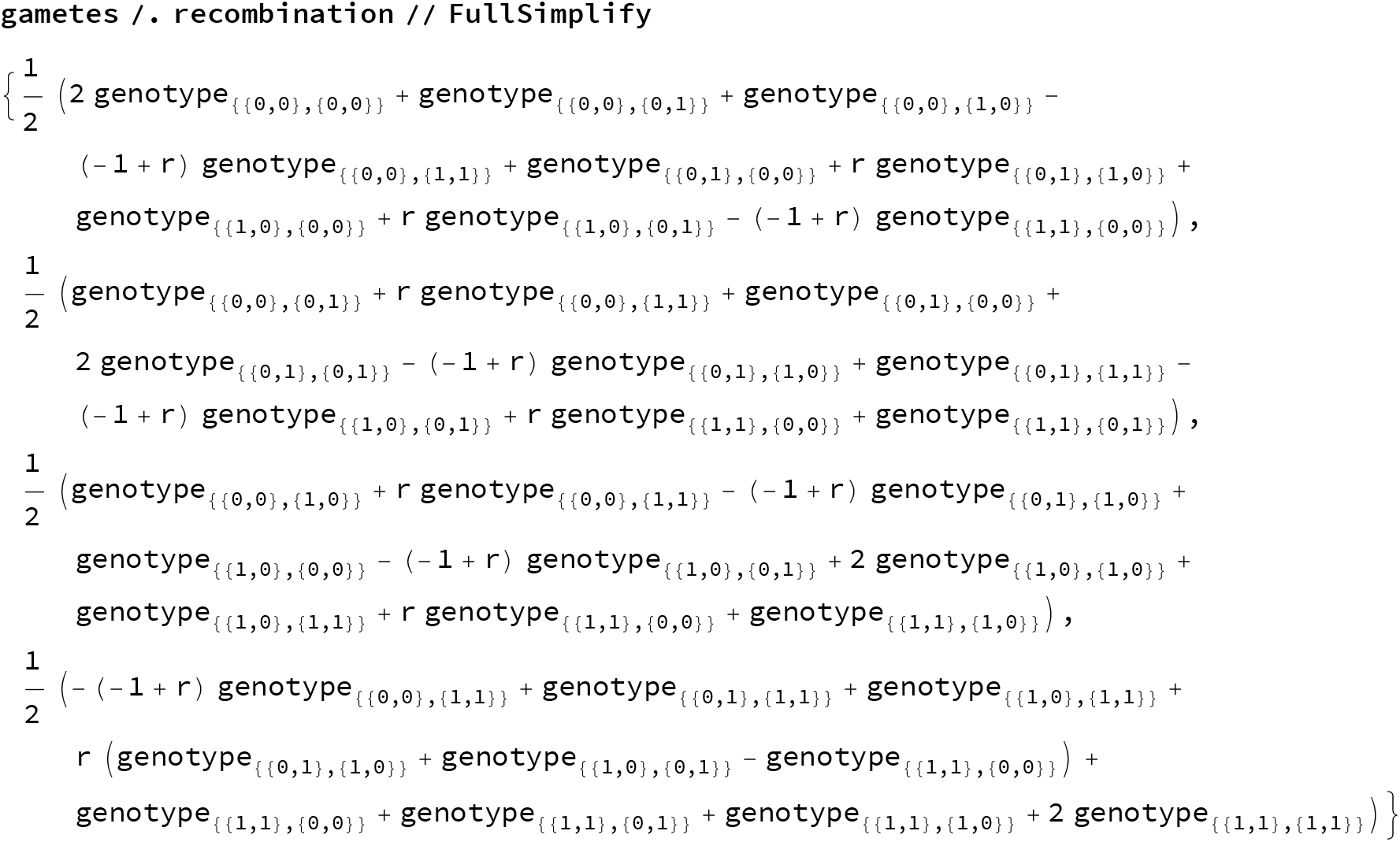

As a test, combine selection and recombination in females, and try some special cases.

~~~
femaleLifeCycle = gametes /. recombination /.
 Thread[genFemale → femaleGenotypesAfterSelection]//Simplify;
~~~

No selection or recombination

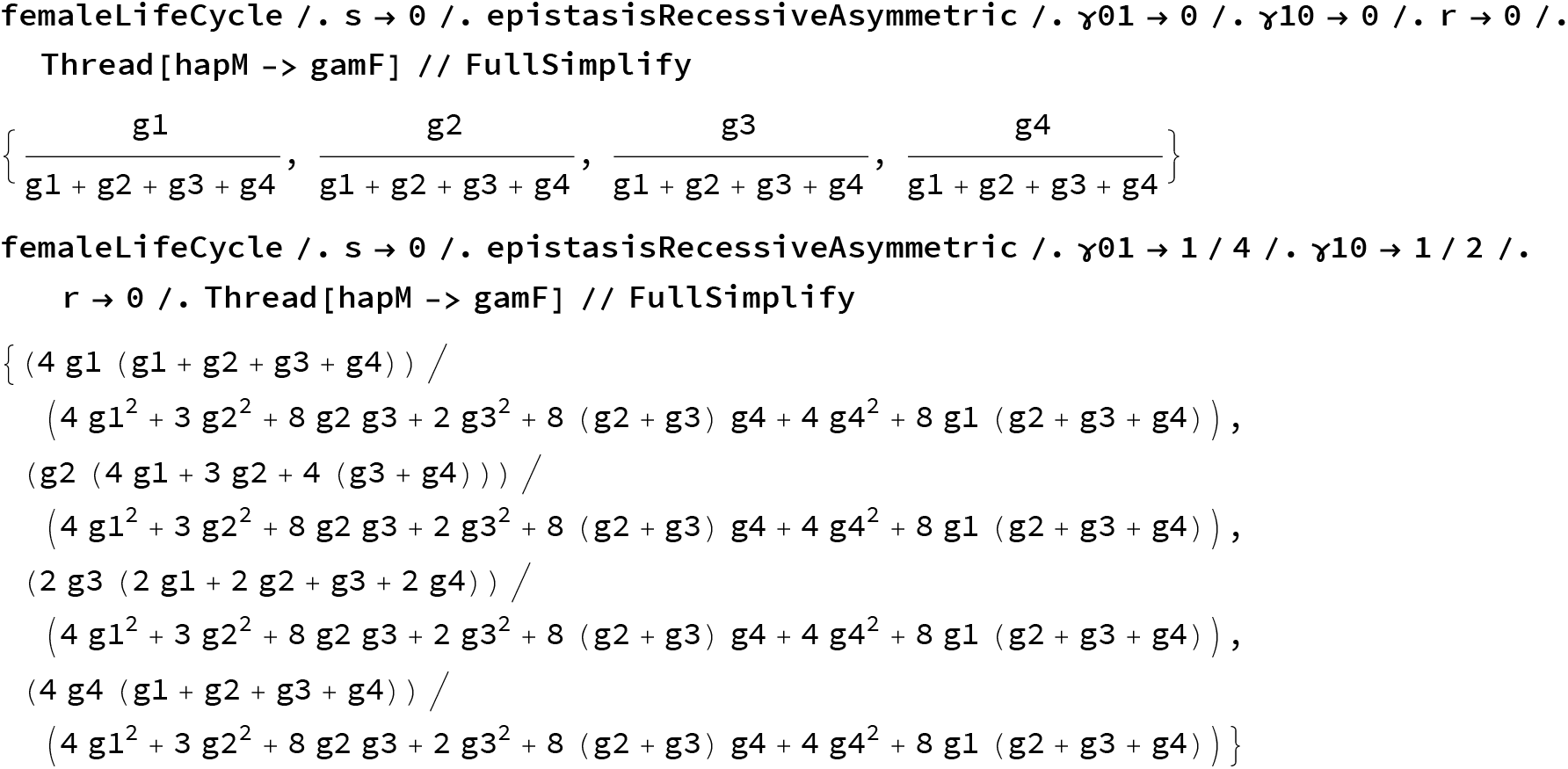

Lethal epistasis and no recombination

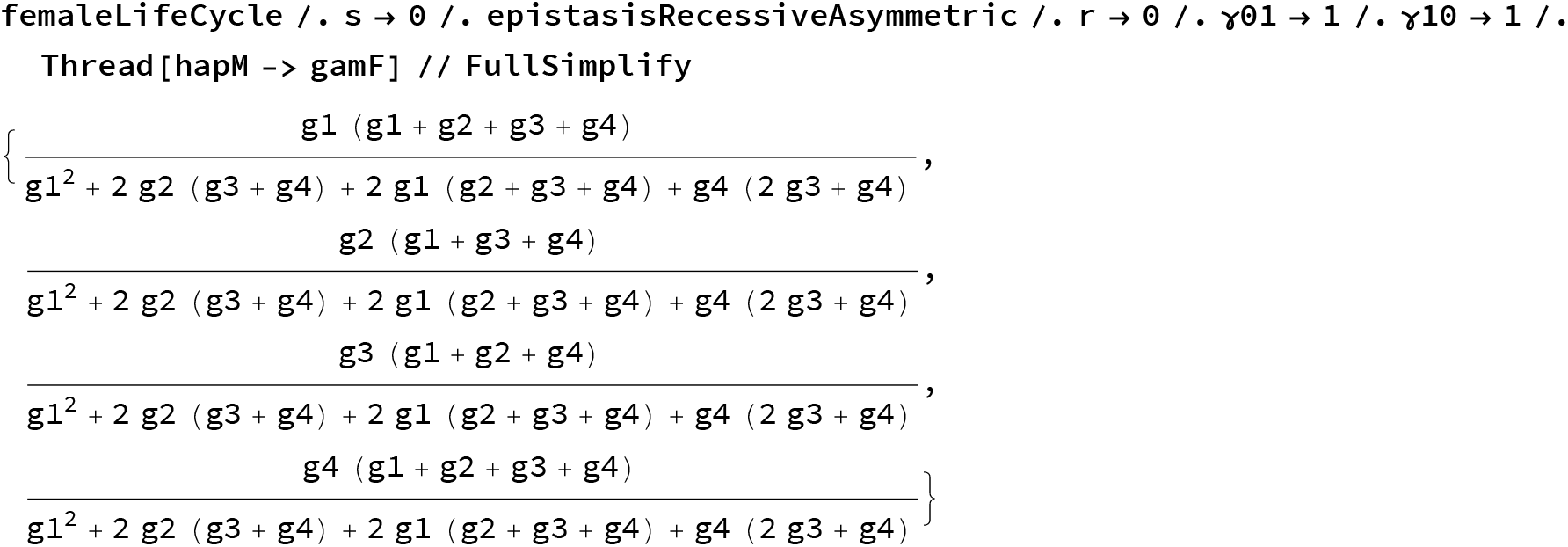

Death of all heterozygotes

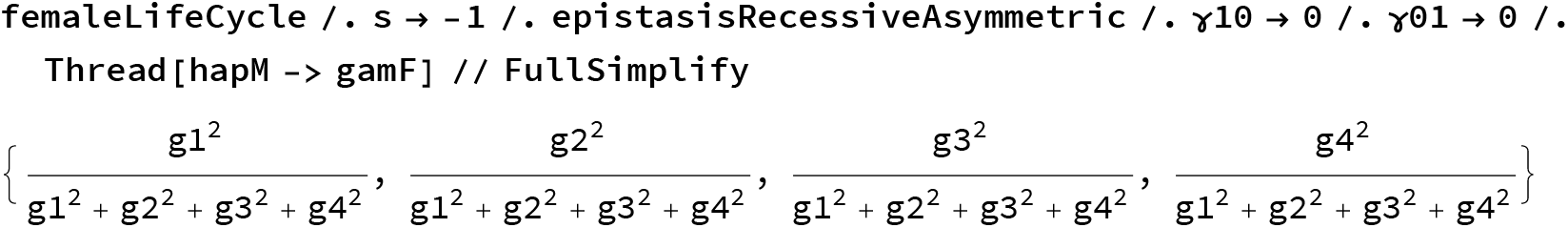

Death of heterozygotes & incompatibility

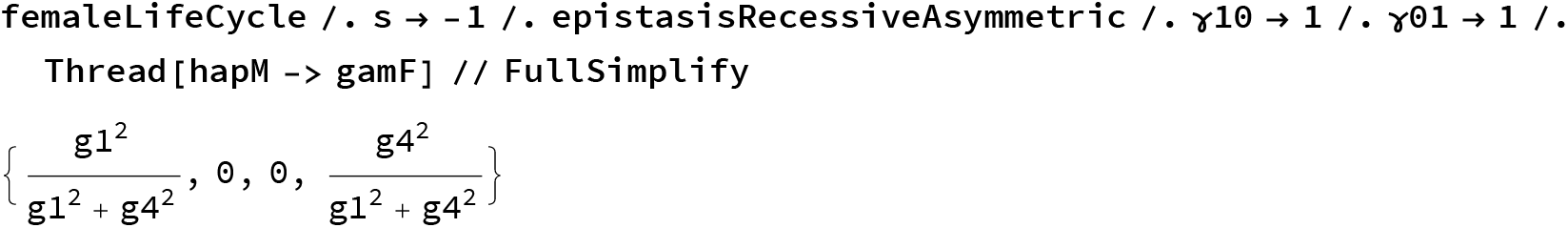

Makes all sense.

#### Dynamics as functions of haplotype frequencies

Here we define functions that describe the dynamics int he haplodiploid model.

##### Haplodiploid dynamics as function of female gametes and male frequencies after selection

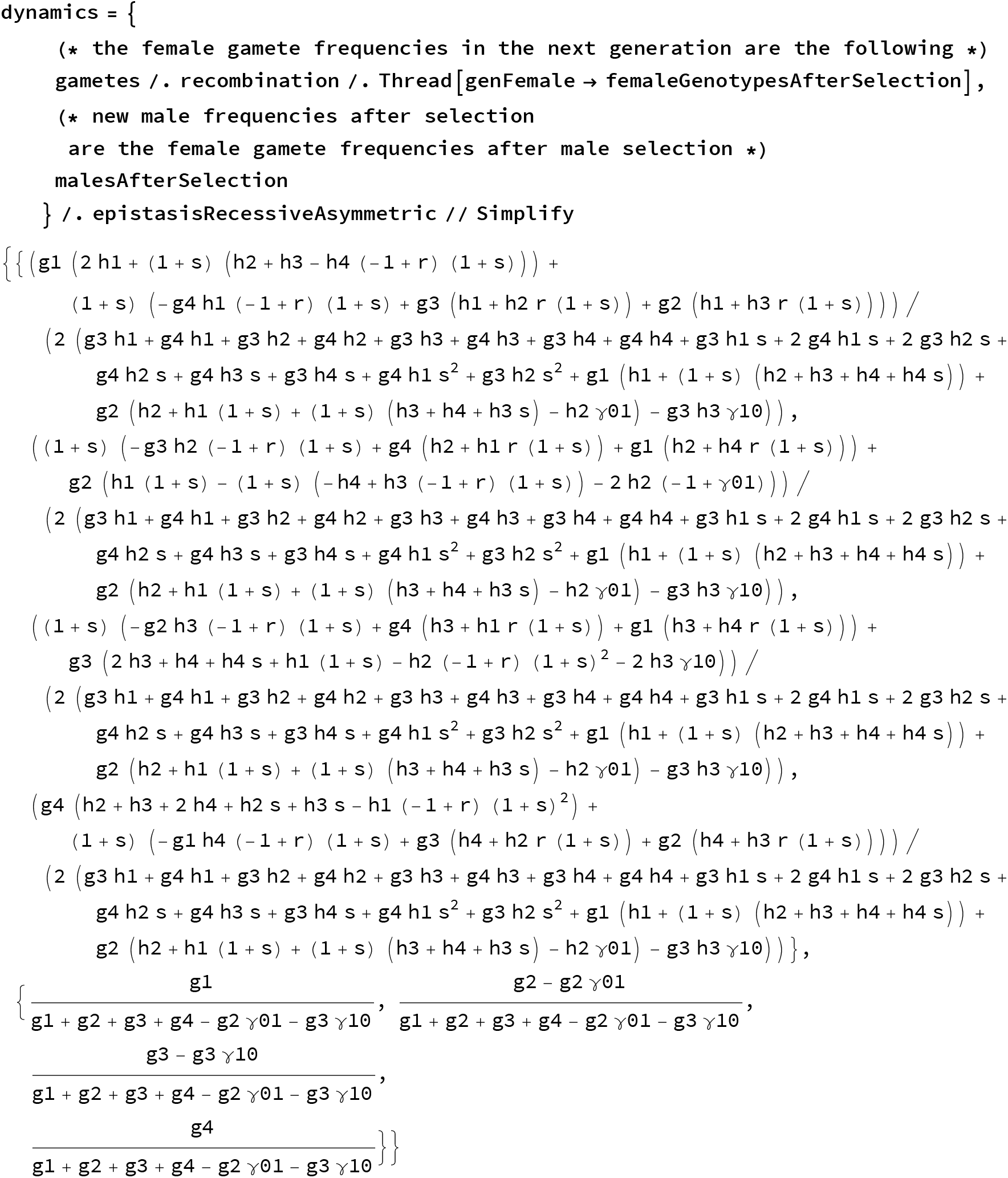

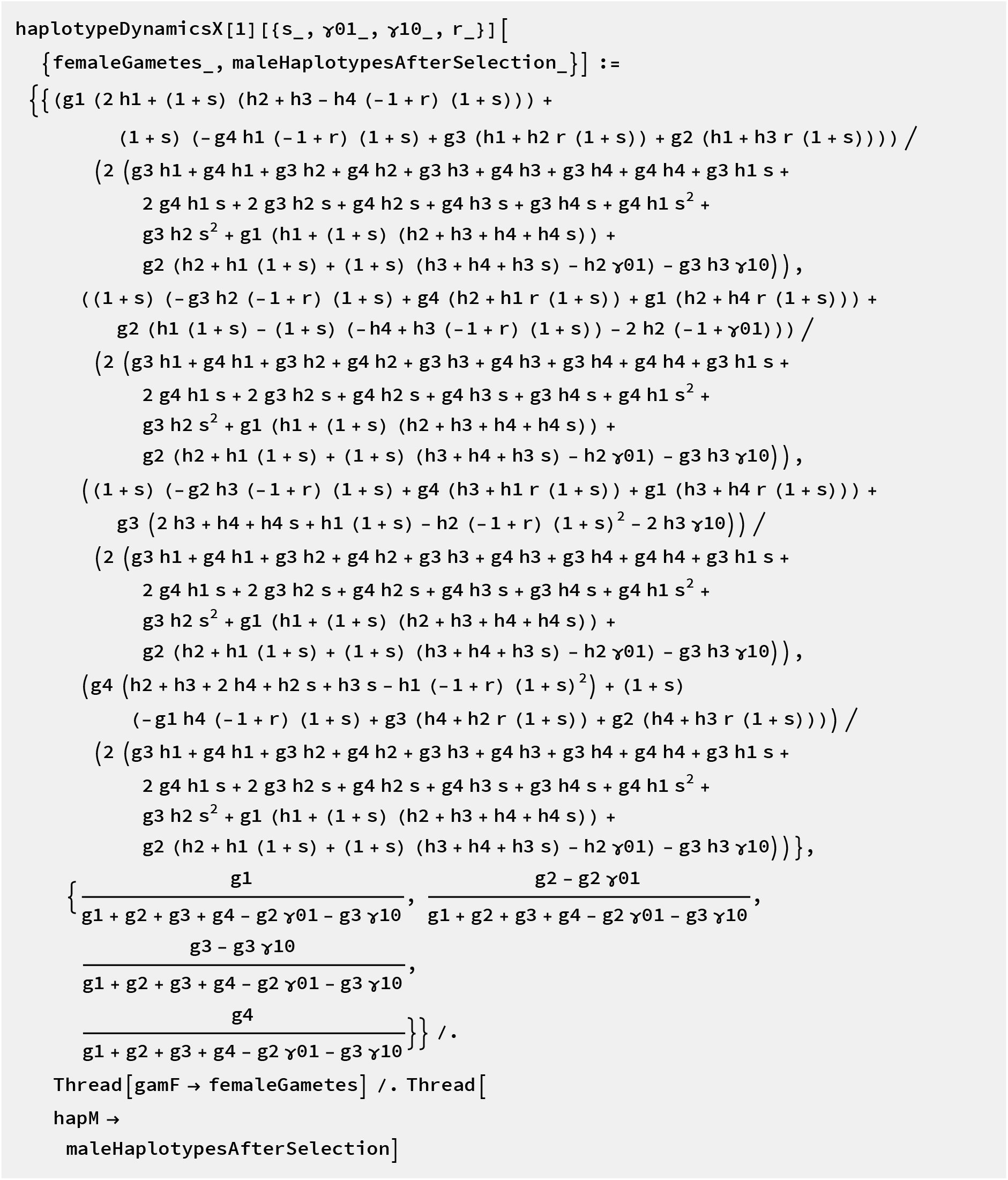

But we can get rid of 2 variables here because frequencies sum up to 1.

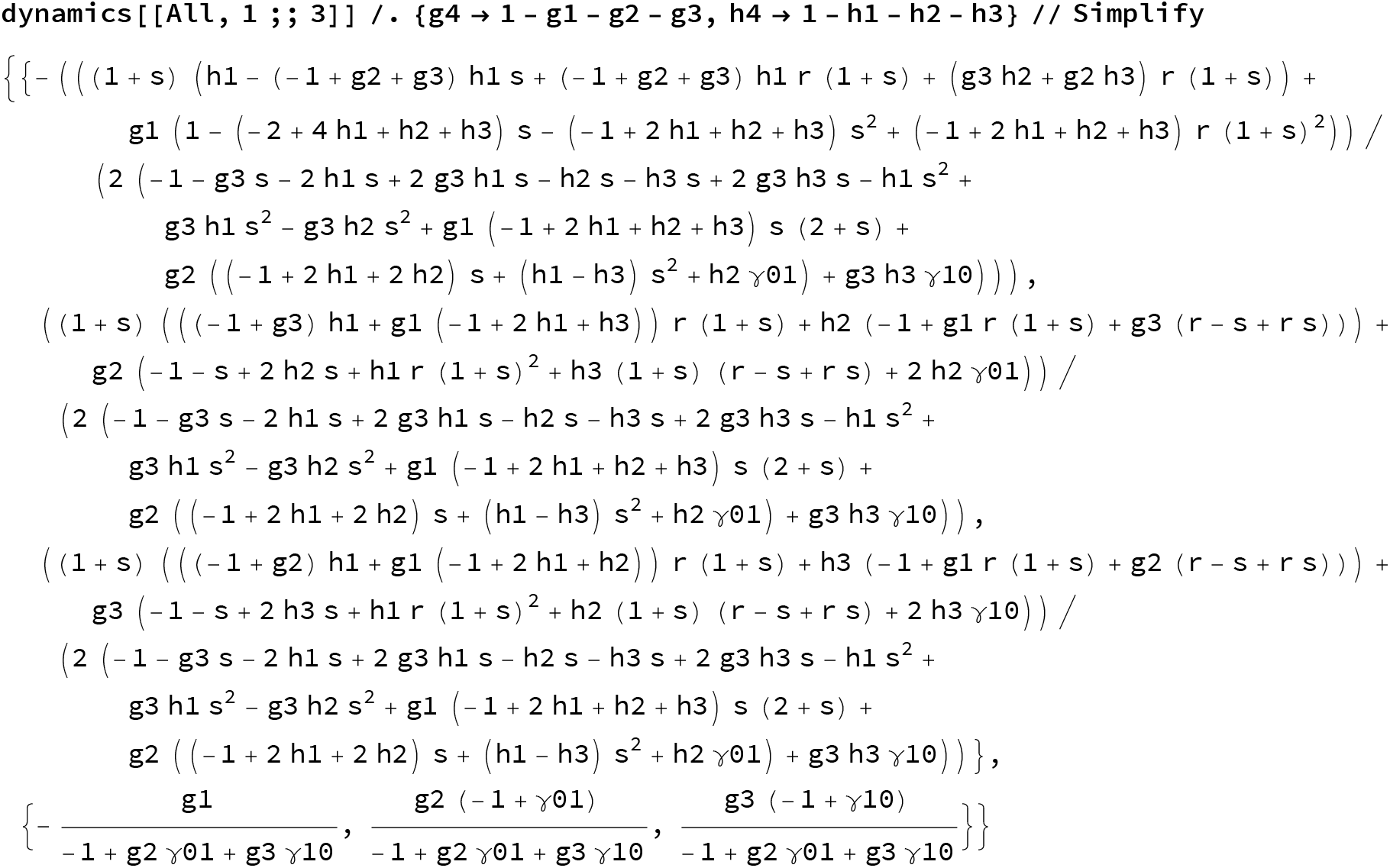

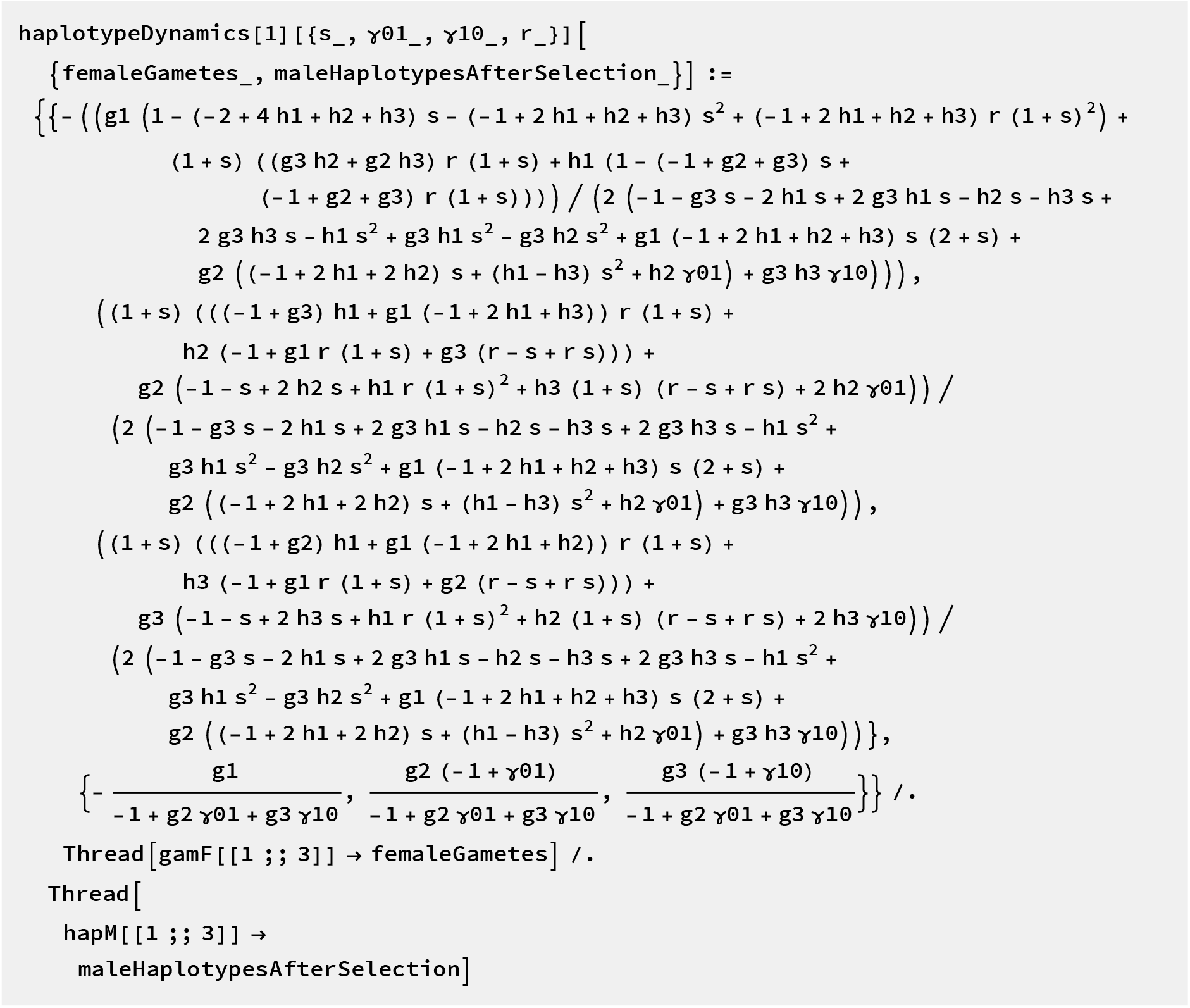

##### Diploid dynamics as function of female gametes

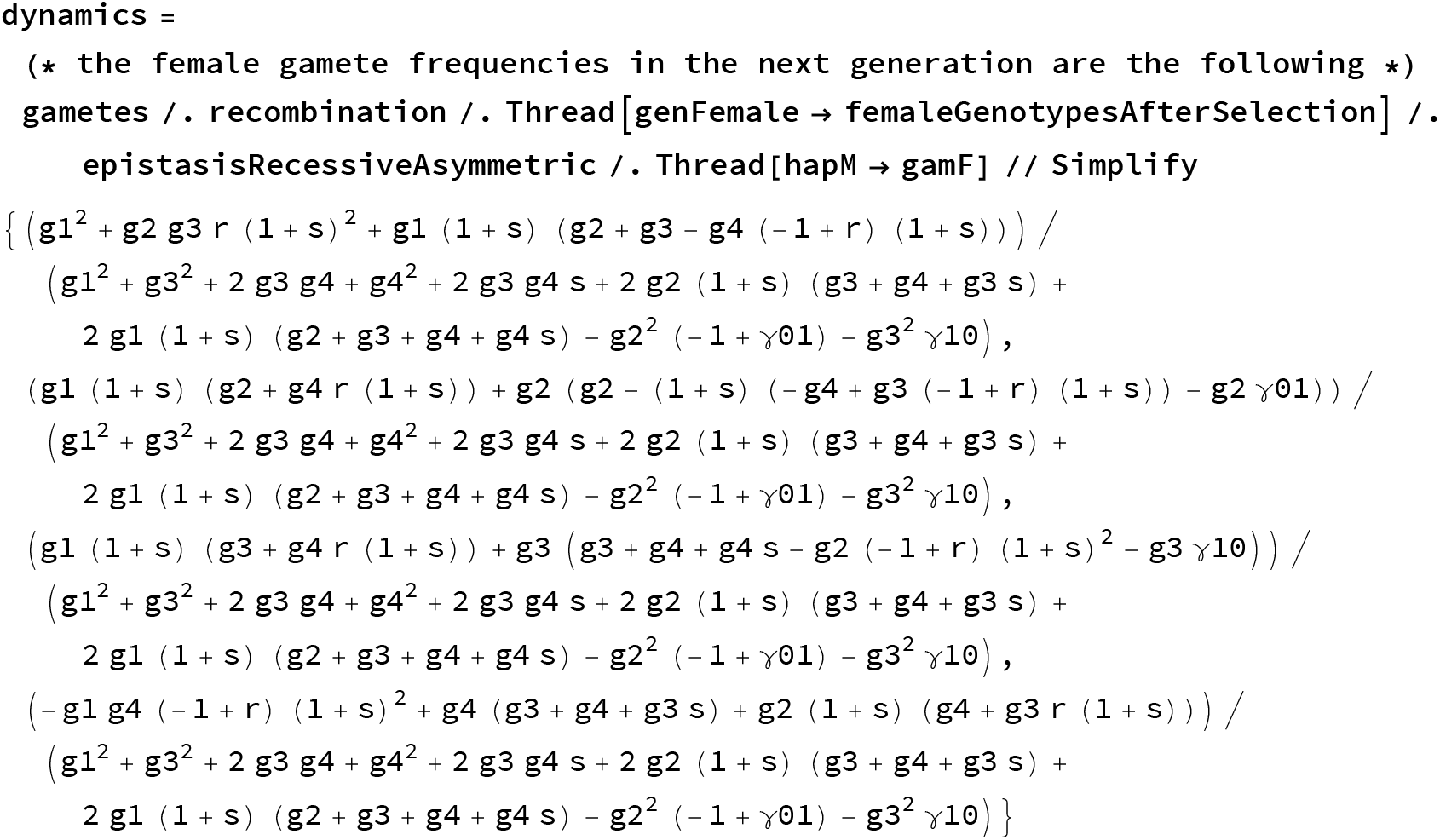

Index 1 stands for the haplodiploid model, index 2 stands for the diploid model. We write down the complete (but redundant) version too (with an X), but only for checking.

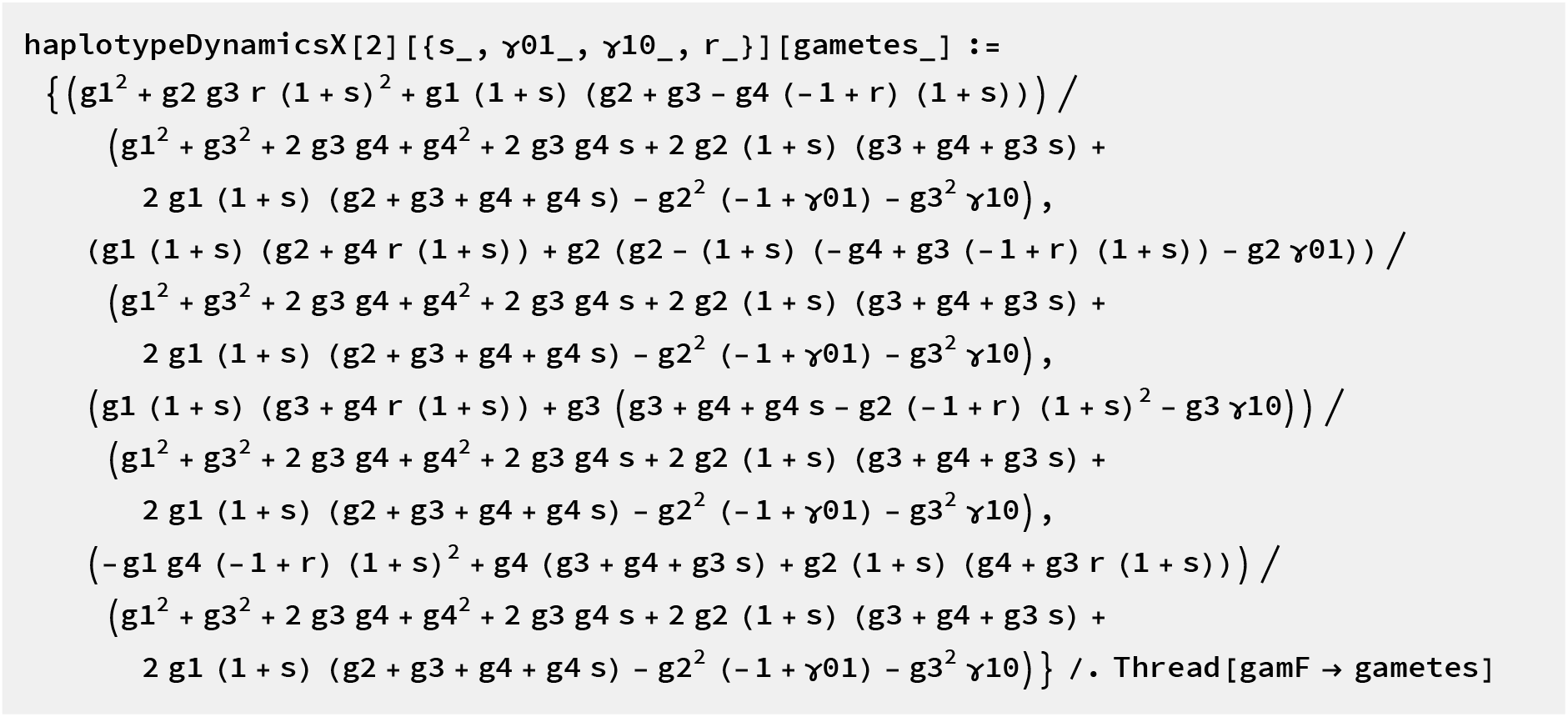

Get rid of redunancy

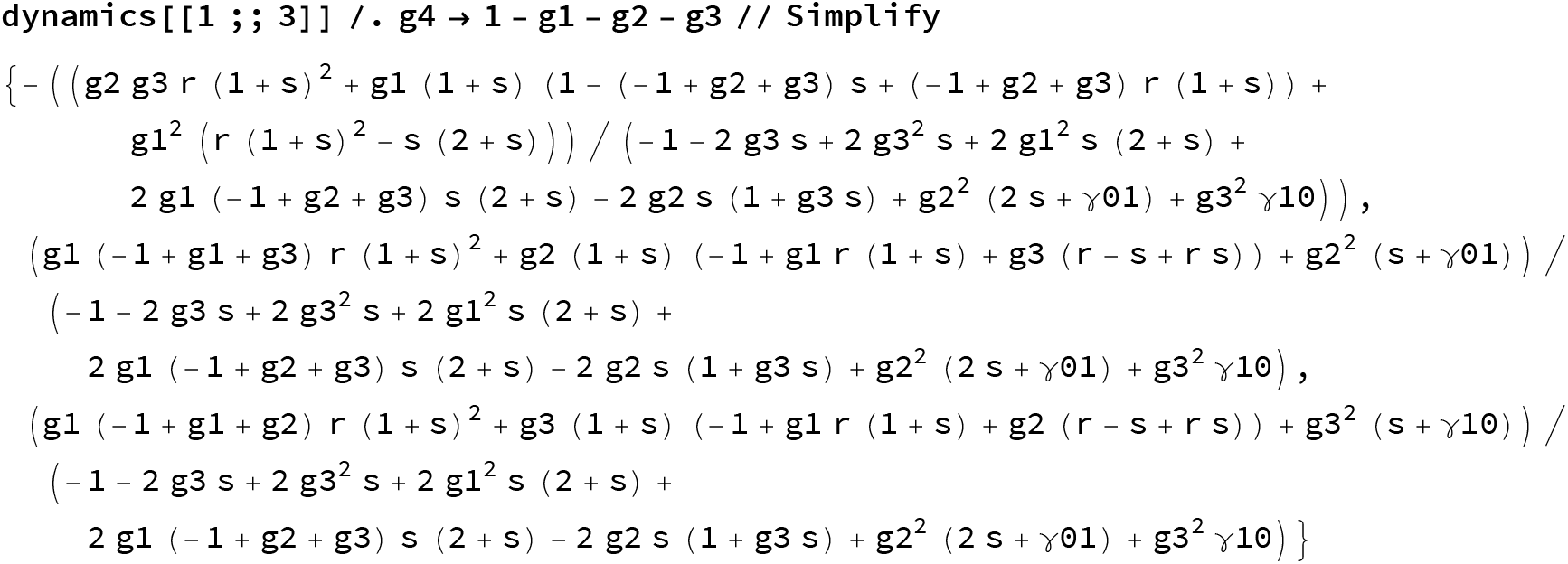

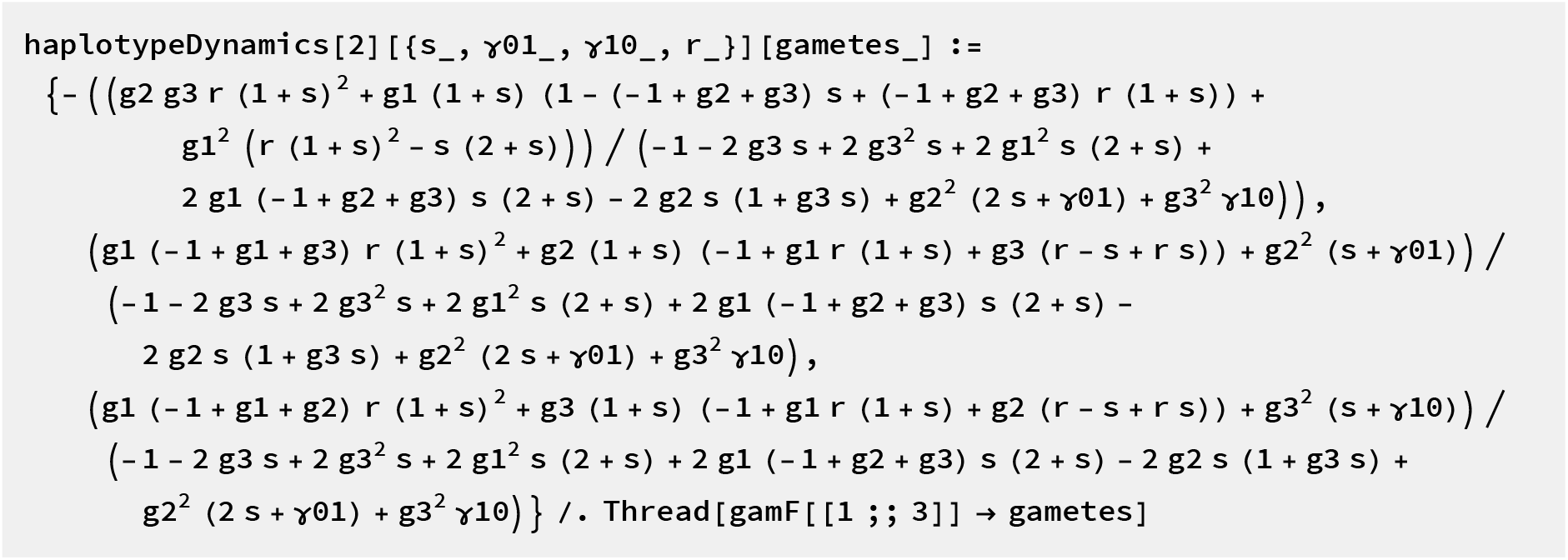

##### Haploid dynamics

For completeness we write down the haploid dynamics too.

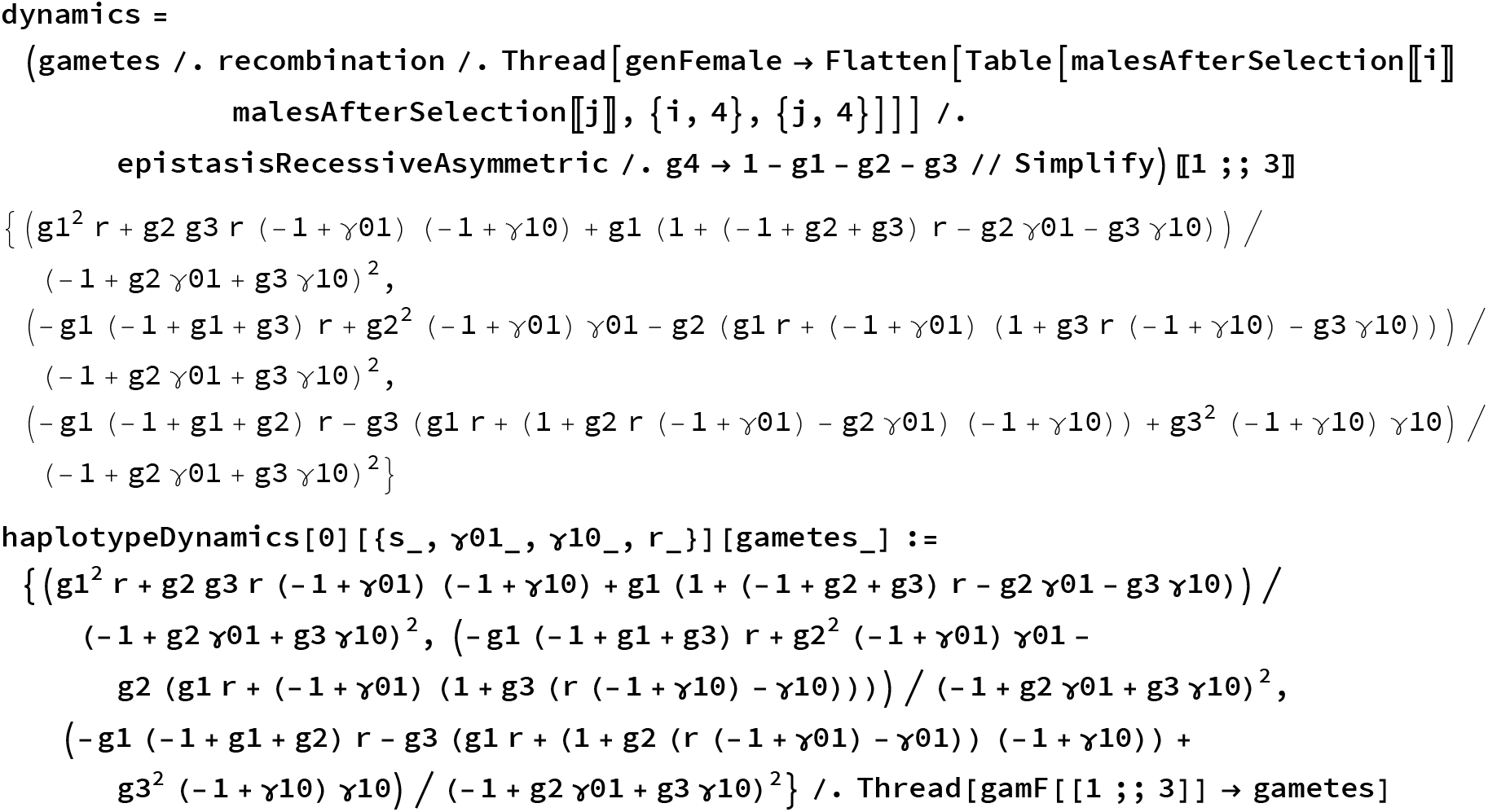

### Dynamics as functions of allele frequencies

Here, we translate the haplotype dynamics into allele frequency dynamics. That can be useful for analytical work later.

#### Transfer rules

p is the frequency of 1 at locus A/0, q is the frequency of allele 1 at locus B/1.

~~~
transferpRulegFemales =
 Solve [{pf == g3 + g4, qf == g2 + g4, LDf == g1 g4 - g2 g3, g1 + g2 + g3 + g4 == 1},
  {g1, g2, g3, g4}] // Flatten
transferpRulegMales = Solve [ {pm == h3 + h4, qm == h2 + h4, LDm == h1 h4 - h2 h3,
   h1 + h2 + h3 + h4 == 1}, {h1, h2, h3, h4}] // Flatten
{g1 → 1 + LDf - pf - qf + pf qf, g2 → - LDf + qf - pf qf, g3 → - LDf + pf - pf qf, g4 → LDf + pfqf}
{h1 → 1 + LDm - pm - qm + pm qm, h2 → - LDm + qm - pm qm, h3 → - LDm + pm - pm qm, h4 → LDm + pm qm}
~~~

#### Conversion function

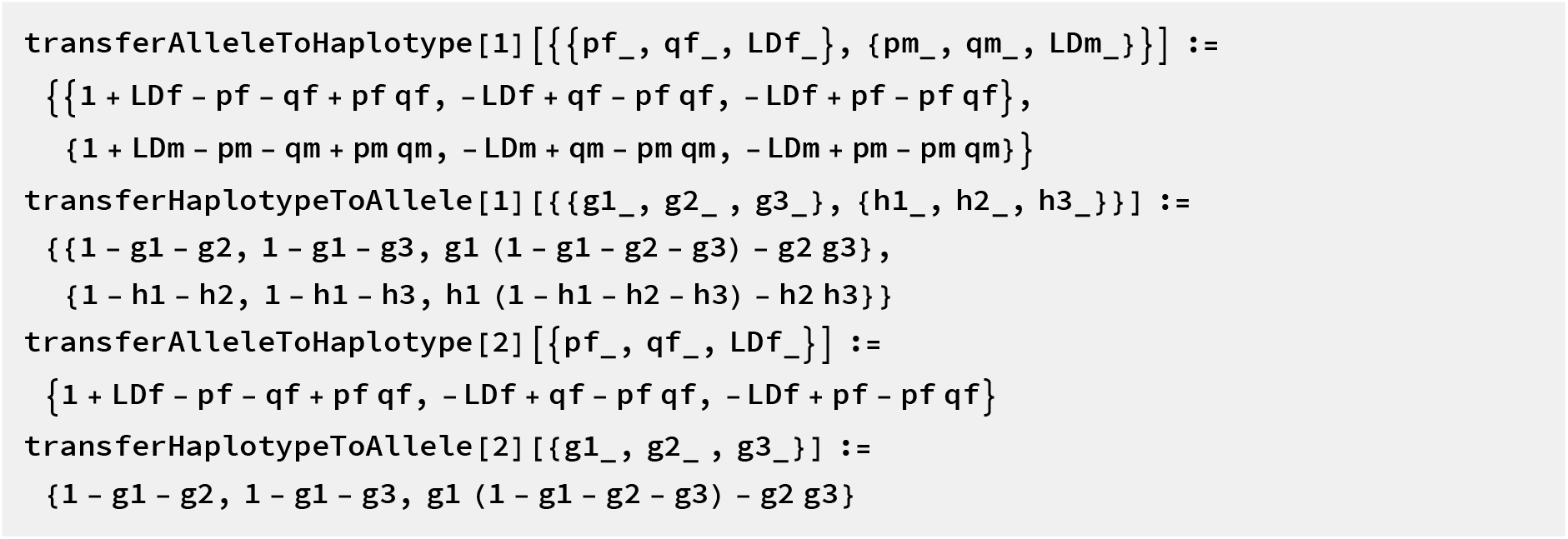

#### Haplodiploid dynamics of allele frequencies and LD

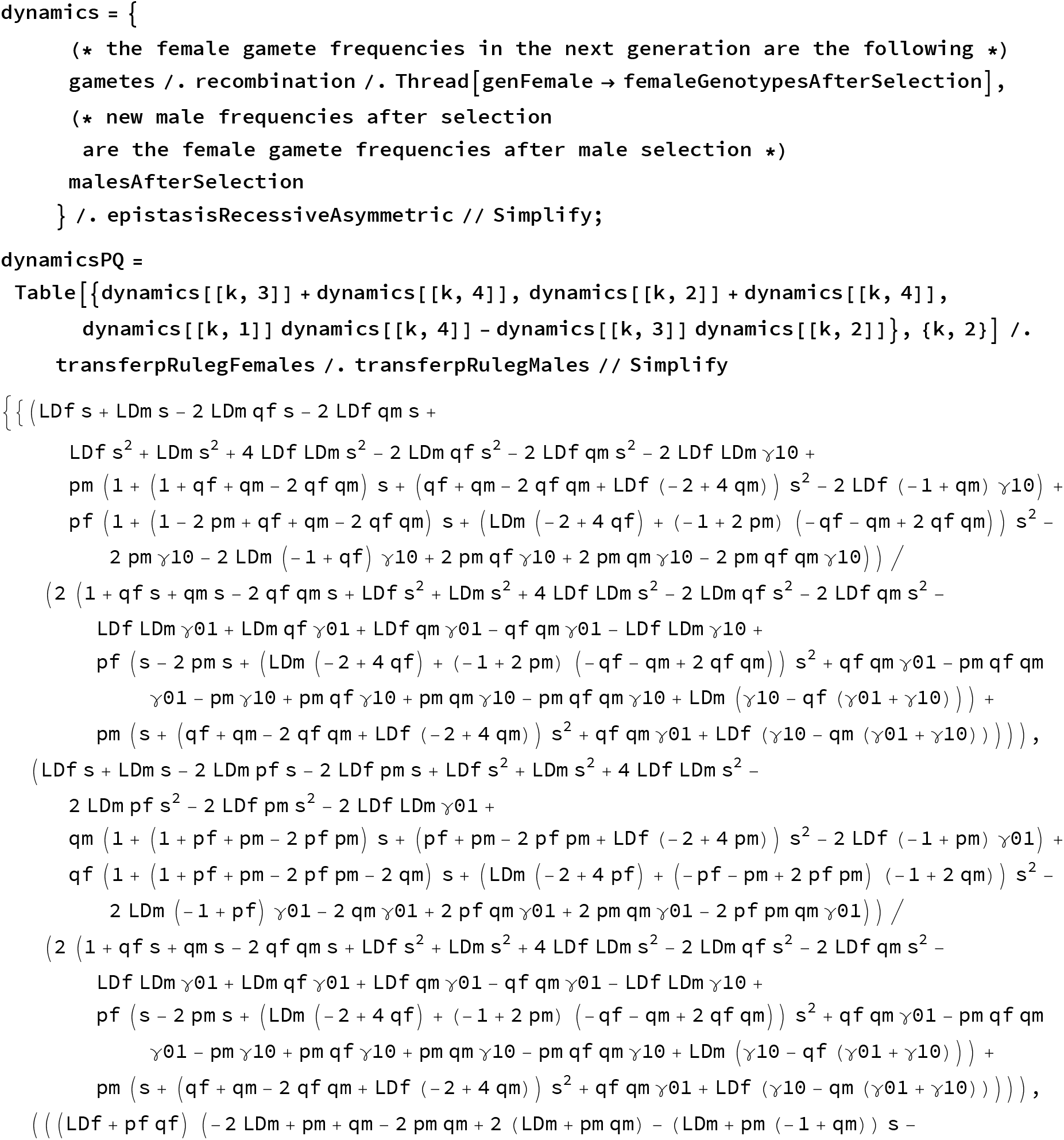
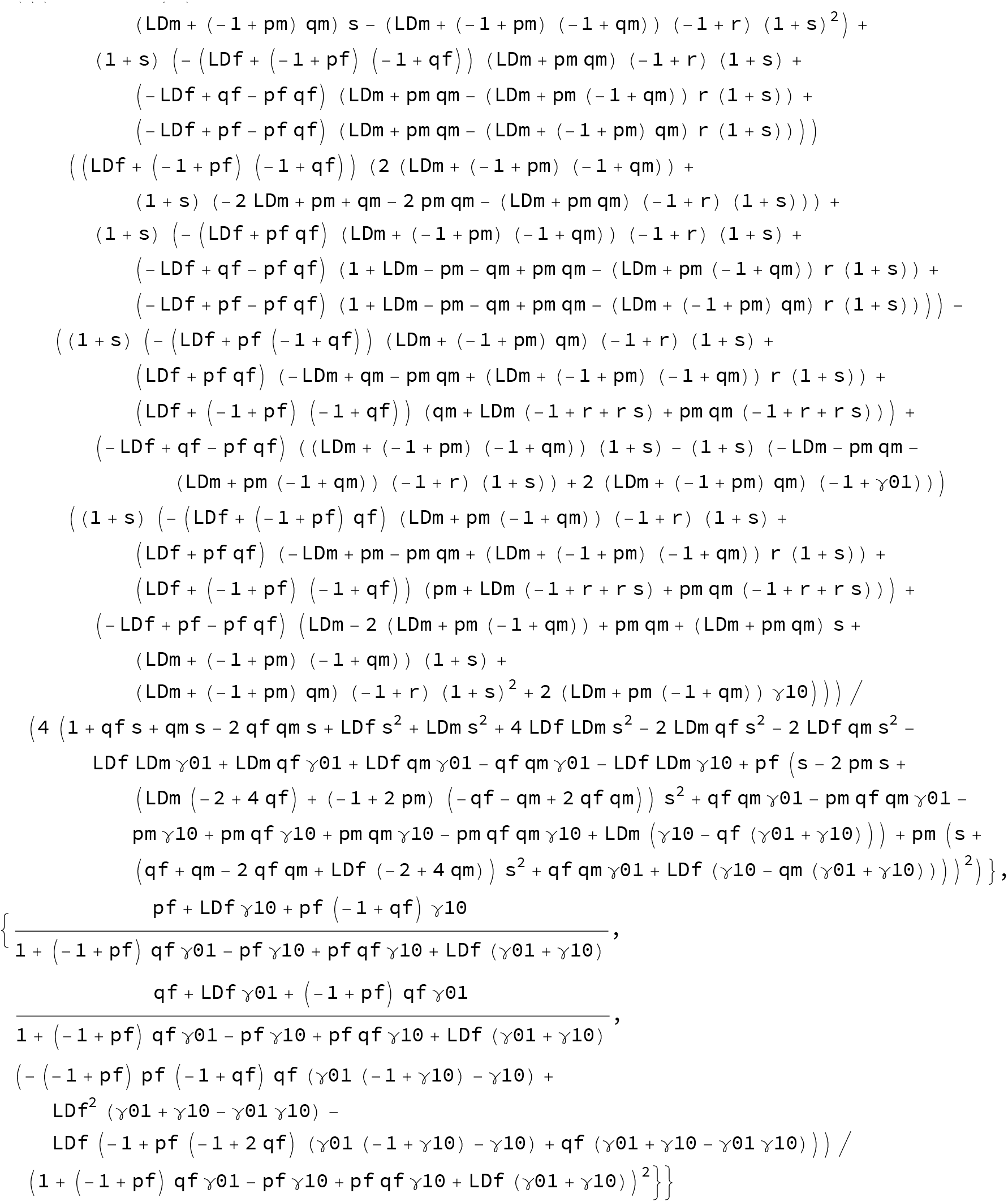

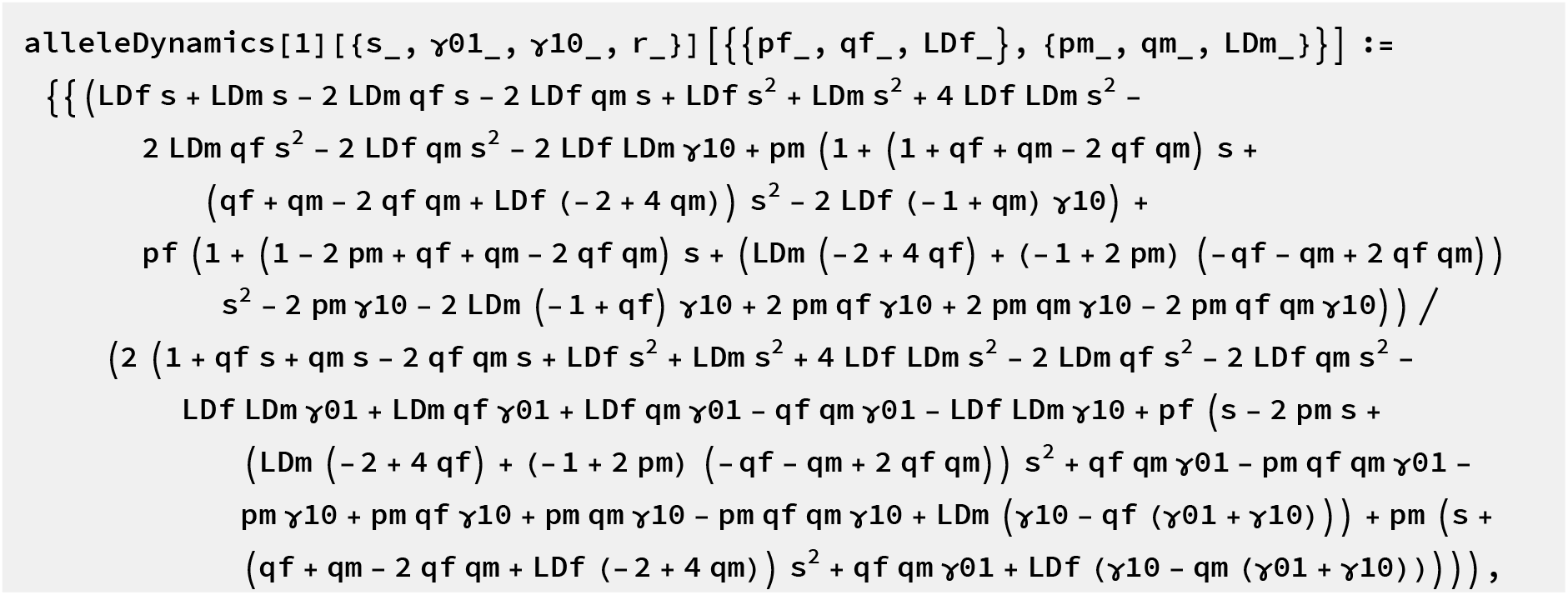
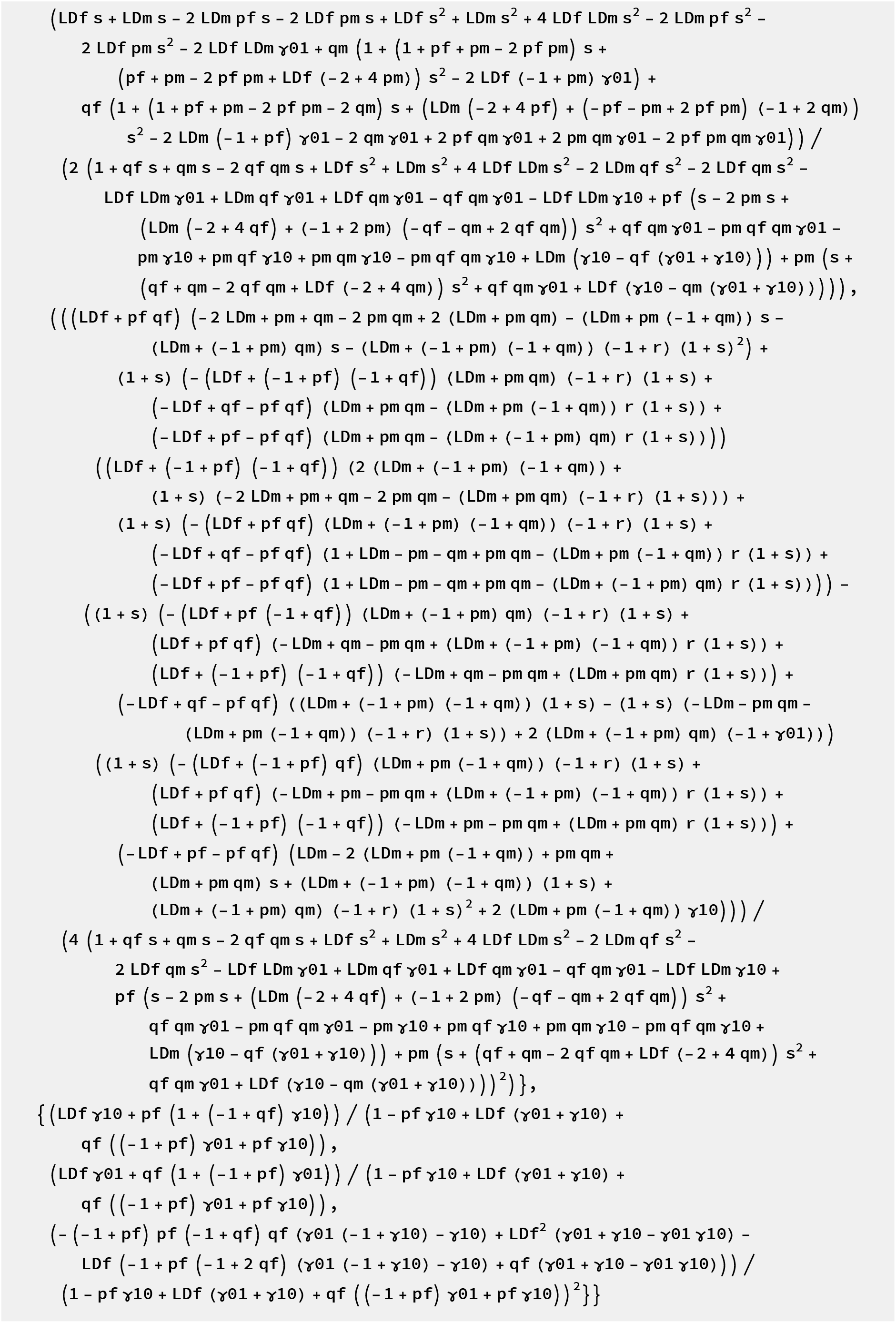

#### Diploid dynamics of allele frequencies and LD

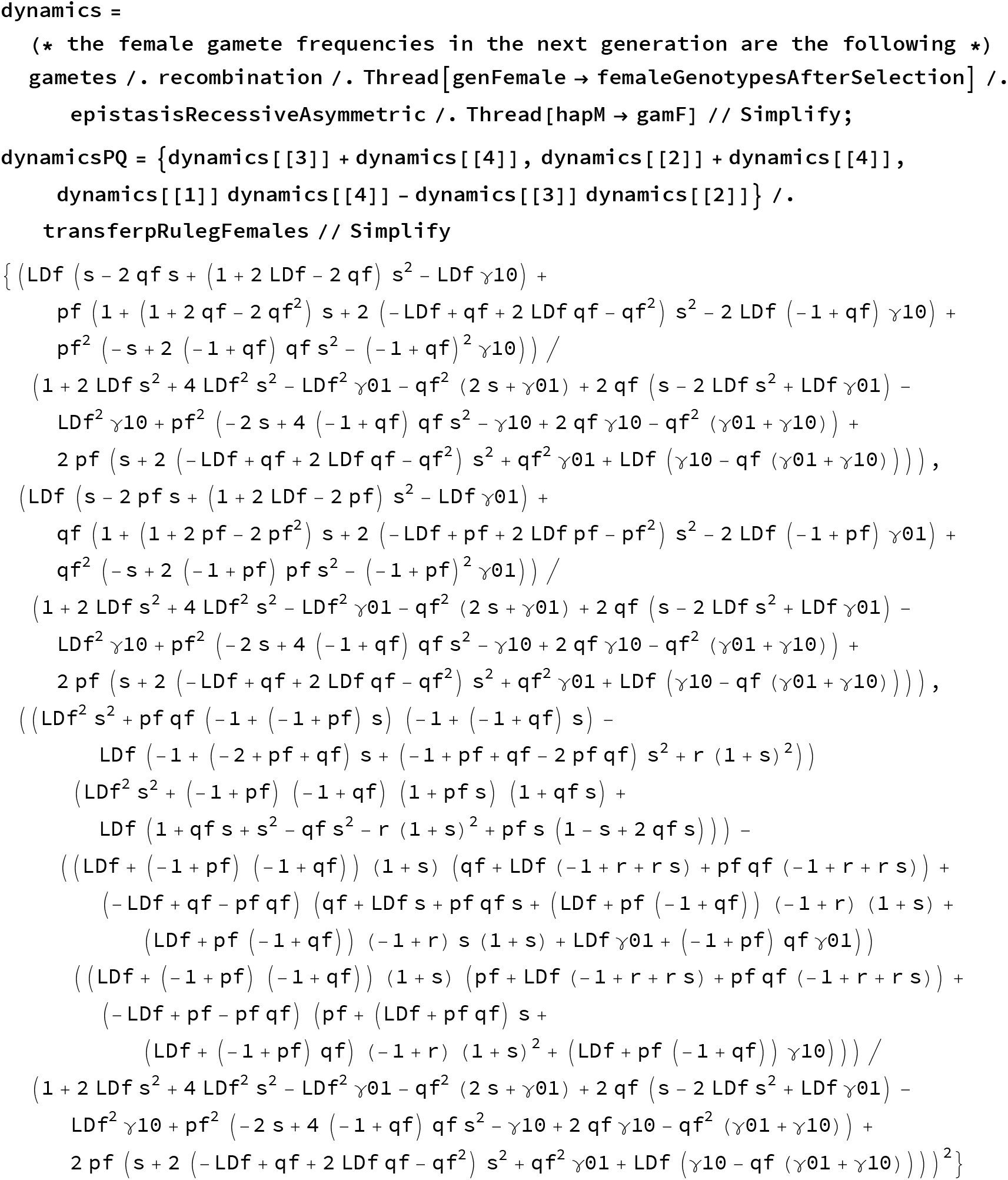

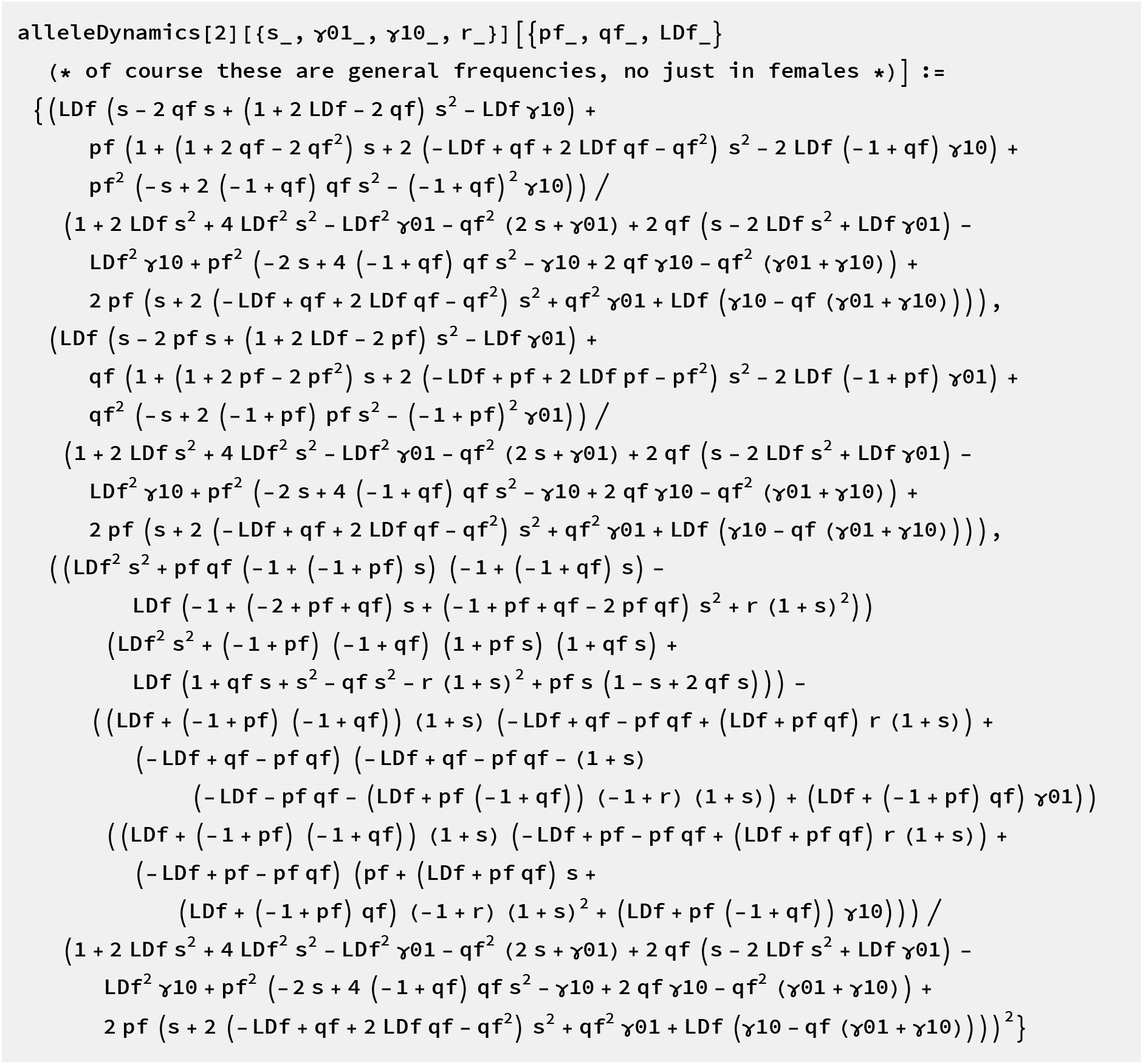

#### Haploid dynamics of allele frequencies and LD

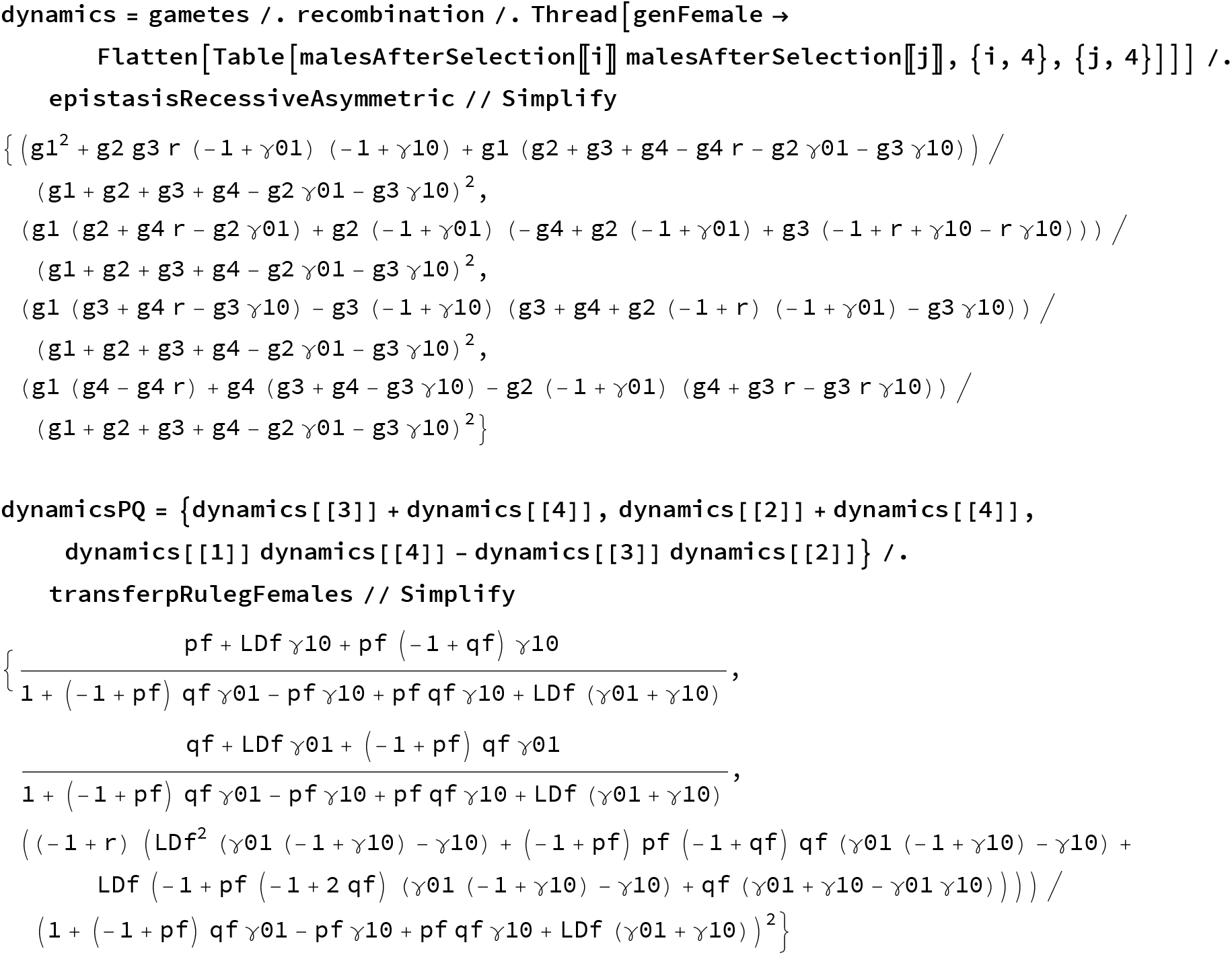

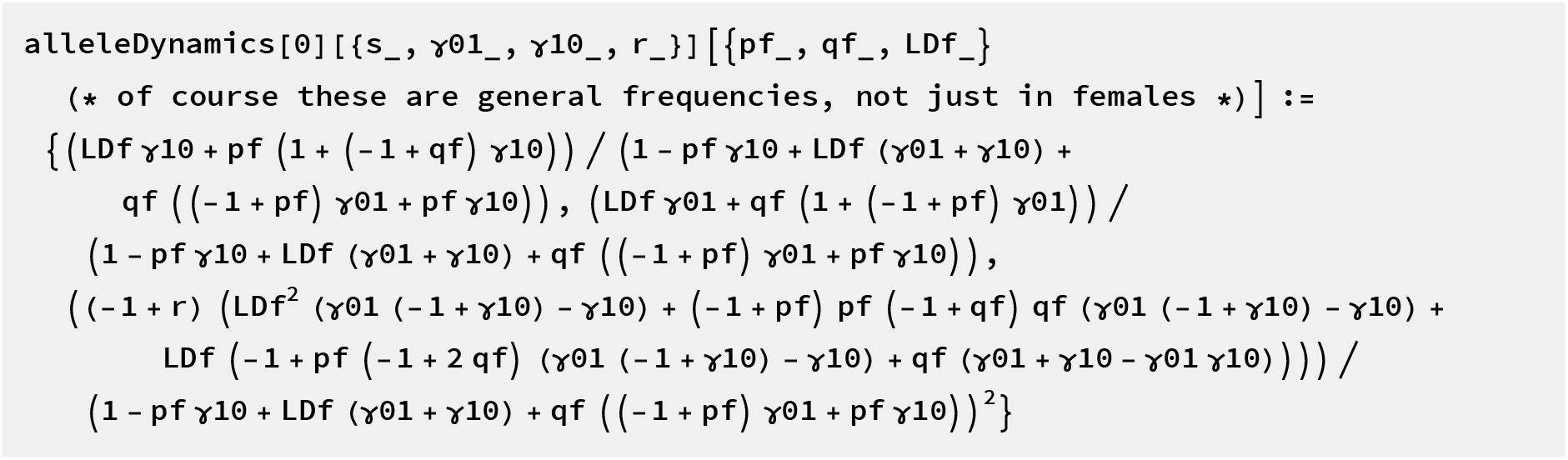

### Some sanity checks of dynamical equations

Make sure to use the X when you want to enter full haplotypes.

~~~
haplotypeDynamicsX [1] [{0, 0, 0, 0}] [{{0.5, 0.4, 0.1, 0}, {0.25, 0.25, 0.25, 0.25}}]
haplotypeDynamics [1] [{0, 0, 0, 0}][{{0.5, 0.4, 0.1}, {0.25, 0.25, 0.25}}]
{{0.375, 0.325, 0.175, 0.125}, (0.S, 0.4, θ.1, 0.}}
{{θ.375, 0.325, 0.175}, {0.5, 0.4, 0.1}}
~~~

#### Iterate dynamics

Should not change if all pars are 0

~~~
startfreq = {0.5, 0.4, 0.1, 0};
pars = {0, 0, 0, 0};
freq = startfreq;
Do[freq = haplotypeDynamicsX[2][pars][freq];
 Print[freq],
 {k, 5}]
 {0.5, 0.4, 0.1, 0. }
 {0.5, 0.4, 0.1, 0. }
 {0.5, 0.4, 0.1, 0. }
 {0.5, 0.4, 0.1, 0. }
 {0.5, 0.4, 0.1, 0. }
startfreq = {{0.5, 0.4, 0.1, 0}, {0.5, 0.4, 0.1, 0}};
pars = {0, 0, 0, 0} ;
freq = startfreq;
Do[freq = haplotypeDynamicsX[1][pars][freq];
 Print[freq],
 {k, 5}]
 {{0.5, 0.4, 0.1, 0.}, {0.5, 0.4, 0.1, 0.}}
 {{0.5, 0.4, 0.1, 0.}, {0.5, 0.4, 0.1, 0.}}
 {{0.5, 0.4, 0.1, 0.}, {0.5, 0.4, 0.1, 0.}}
 {{0.5, 0.4, 0.1, 0.}, {0.5, 0.4, 0.1, 0.}}
 {{0.5, 0.4, 0.1, 0.}, {0.5, 0.4, 0.1, 0.}}
startfreq = {0.5, 0.4, 0, 0.1};
pars = {0.4, 1, 1, 1 /3};
freq = startfreq;
Do[freq = haplotypeDynamicsX[2][pars][freq] // Chop;
 If[IntegerQ[k / 1000], Print[freq]; Print[Total[freq]]],
 {k, 5000}]
transferHaplotypeToAllele[2][freqĮ1 ;; 3ļ]
{0.2913, 0.2087, 0.2087, 0.2913}
1.
{0.2913, 0.2087, 0.2087, 0.2913}
1.
{0.2913, 0.2087, 0.2087, 0.2913}
1.
{0.2913, 0.2087, 0.2087, 0.2913}
1.
{0.2913, 0.2087, 0.2087, 0.2913}
1.
{0.5, 0.5, 0.0412997}
~~~

Should go to exclusion with only *γ*

~~~
startfreq = {0.5, 0.4, 0, 0.1};
pars = {0, 0.2, 0.2, 1 / 2};
freq = startfreq;
Do[freq = haplotypeDynamicsX[2][pars][freq] // Chop;
 If[IntegerQ[k / 1000], Print[freq]; Print[Total[freq]]],
 {k, 5000}]
{0.989231, 0.00538911, 0.00535054, 0.0000292745}
1.
{0.994777, 0.0026125, 0.00260356, 6.85181×10^−6^}
1.
{0.99656, 0.00172064, 0.00171678, 2.96825×10^−6^}
1.
{0.997437, 0.00128196, 0.00127982, 1.64659×10^−6^}
1.
{0.997958, 0.00102125, 0.0010199, 1.04456×10^−6^}
1.
~~~

Because of the recessivity it is really slow at the end. It’s much faster for haplodiploids:

~~~
startfreq = {{0.5, 0.4, 0, 0.1}, {0.5, 0.4, 0, 0.1}};
pars = {0, 0.2, 0.2, 1 / 2};
freq = startfreq;
Do[freq = haplotypeDynamicsX[1][pars][freq] // Chop;
 If[IntegerQ[k / 1000], Print[freq]],
 {k, 5000}]
transferHaplotypeToAllele[1][freq[[All, 1 ;; 3]]]
{{1., 0, 0, 0}, {1., 0, 0, 0}}
{{1., 0, 0, 0}, {1., 0, 0, 0}}
{{1., 0, 0, 0}, {1., 0, 0, 0}}
{{1., 0, 0, 0}, {1., 0, 0, 0}}
{{1., 0, 0, 0}, {1., 0, 0, 0}}
{{0., 0., 0.}, {0., 0., 0.}}
startfreq = {{0.1, 0, 0, 0.9}, {0.1, 0, 0, 0.9}};
pars = {0, 0.3, 1, 1 / 7}; (* “asymm. DMI!” *)
freq = startfreq;
Do[freq = haplotypeDynamicsX[1][pars][freq] // Chop;
 If[IntegerQ[k / 1000], Print[freq]],
 {k, 5000}]
transferHaplotypeToAllele[1][freq[[All, 1 ;; 3]]]
{{0, 0, 0, 1.}, {0, 0, 0, 1.}}
{{0, 0, 0, 1.}, {0, 0, 0, 1.}}
{{0, 0, 0, 1.}, {0, 0, 0, 1.}}
{{0, 0, 0, 1.}, {0, 0, 0, 1.}}
{{0, 0, 0, 1.}, {0, 0, 0, 1.}}
{{1, 1, 0}, {1, 1, 0}}
startfreq = {{0.1, 0, 0, 0.9}, {0.1, 0, 0, 0.9}};
pars = {.61, 0, 1, 1 / 2}; (* “asymm. DMI!” *)
freq = startfreq;
Do[freq = haplotypeDynamicsX[1][pars][freq] // Chop;
 If[IntegerQ[k / 1000], Print[freq]],
 {k, 5000}]
transferHaplotypeToAllele[1][freq[[All, 1 ;; 3]]]
{{0.23736, 0.42985, 0.0954301, 0.23736}, {0.262401, θ.475198, θ, 0.262401}}
{{θ.23736, θ.42985, θ.θ9543θ1, θ.23736}, {θ.2624θ1, θ.475198, θ, θ.2624θ1}}
{{θ.23736, θ.42985, θ.θ9543θ1, θ.23736}, {θ.2624θ1, θ.475198, θ, θ.2624θ1}}
{{θ.23736, θ.42985, θ.θ9543θ1, θ.23736}, {θ.2624θ1, θ.475198, θ, θ.2624θ1}}
{{θ.23736, θ.42985, θ.θ9543θ1, θ.23736}, {θ.2624θ1, θ.475198, θ, θ.2624θ1}}
{{θ.33279, θ.66721, θ.θ153192}, {θ.2624θ1, θ.737599, θ.θ688543}}
~~~

#### Define Jacobian

~~~
JacobianAllele[1][par_][x_] :=
D[Flatten[alleleDynamics[1][par][{{pf, qf, LDf}, {pm, qm, LDm}}], 2],
  {{pf, qf, LDf, pm, qm, LDm}}] /. x
JacobianAllele[2][par_][x_] :=
 D[alleleDynamics[2][par][{pf, qf, LDf}], {{pf, qf, LDf}}] /. x
JacobianHaplotype[1][par_][x_] :=
 D[Flatten[haplotypeDynamics[1][par][{{g1, g2, g3}, {h1, h2, h3}}], 2],
  {{g1, g2, g3, h1, h2, h3}}] /. x
JacobianHaplotype[2][par_][x_] :=
 D[haplotypeDynamics[2][par][{g1, g2, g3}], {{g1, g2, g3}}] /. x
JacobianAllele[0][par_][x_] :=
 D[alleleDynamics[0][par][{pf, qf, LDf}], {{pf, qf, LDf}}] /. x
JacobianHaplotype[0][par_][x_] :=
 D[haplotypeDynamics[0][par][{g1, g2, g3}], {{g1, g2, g3}}] /. x
~~~

As a test, we can compute the eigenvalues of a monomorphic equilibrium, both from a haplotype perspective and an allele perspective.

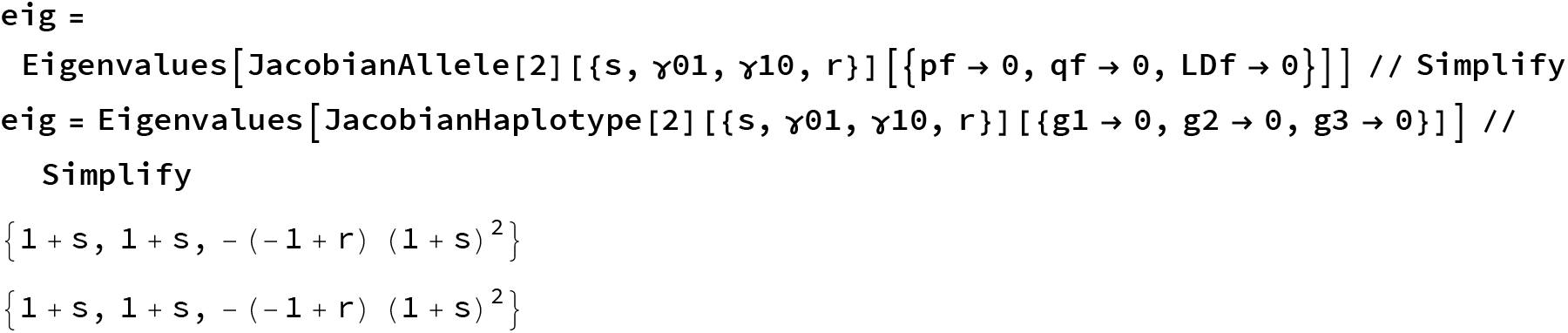

#### A function that determines equilibria and stability numerically

For haplodiploid model:

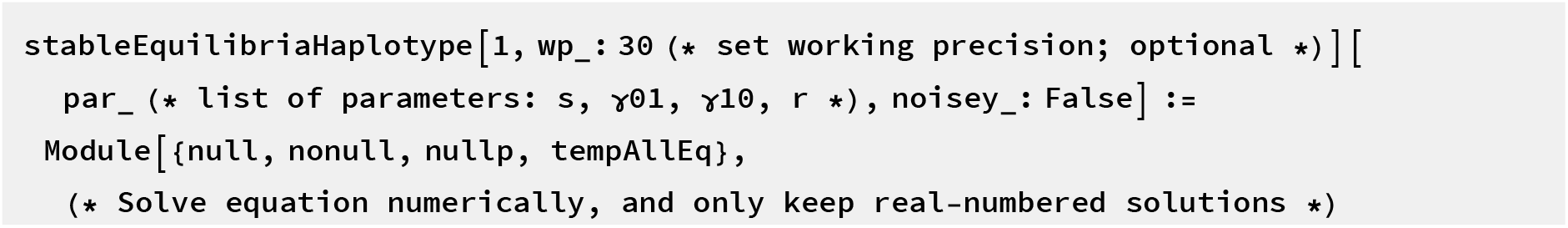
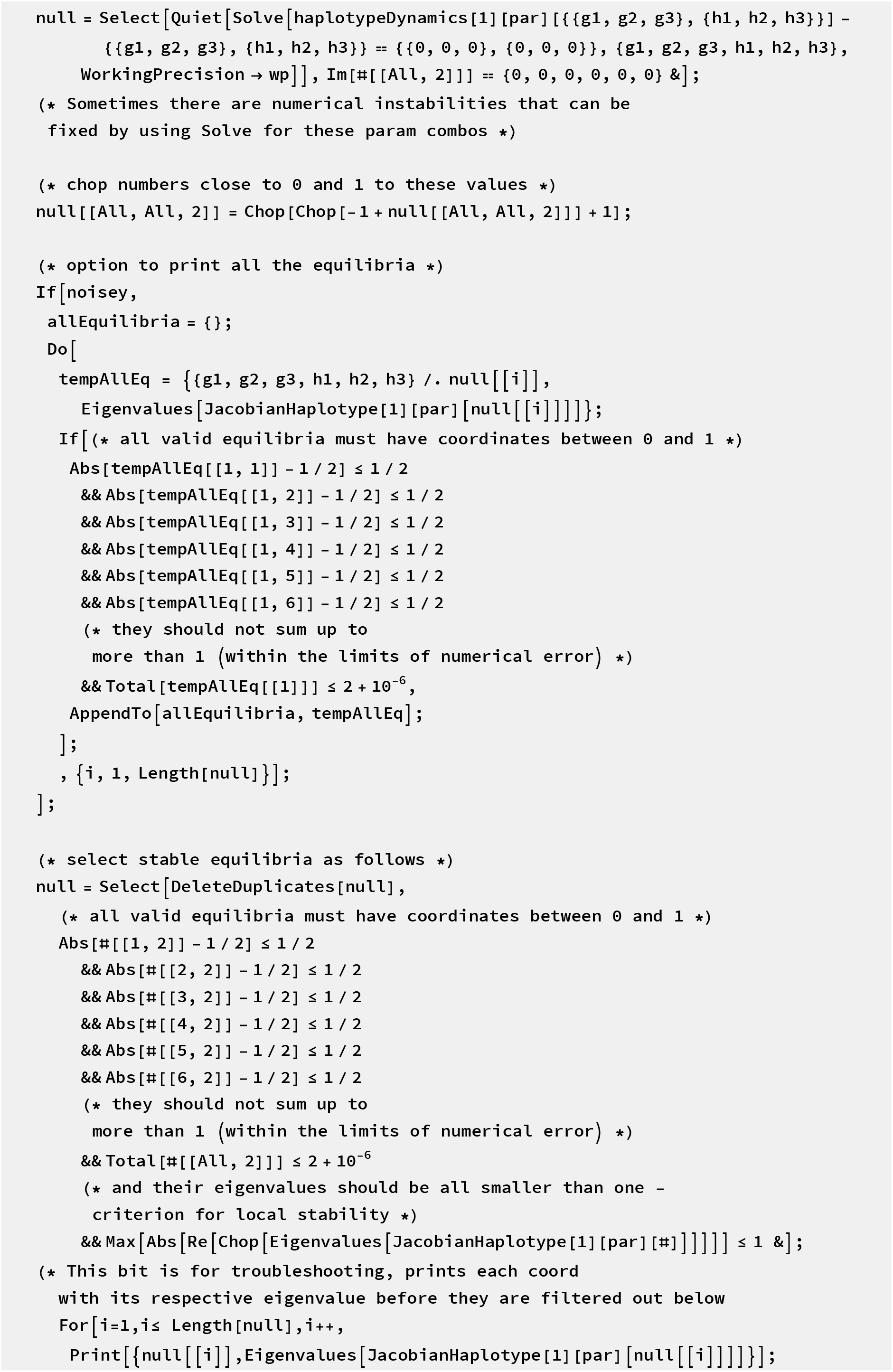
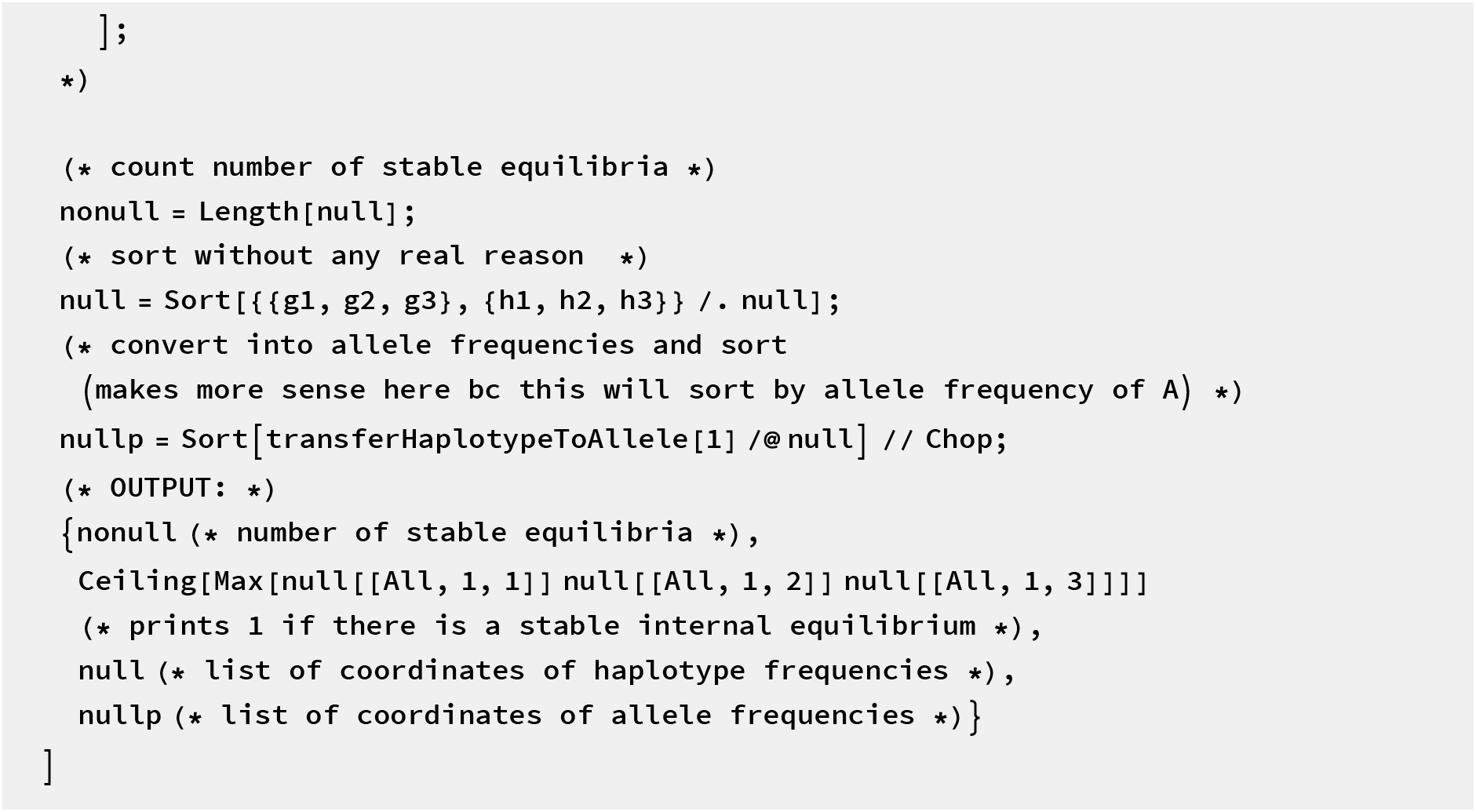

For diploid model:

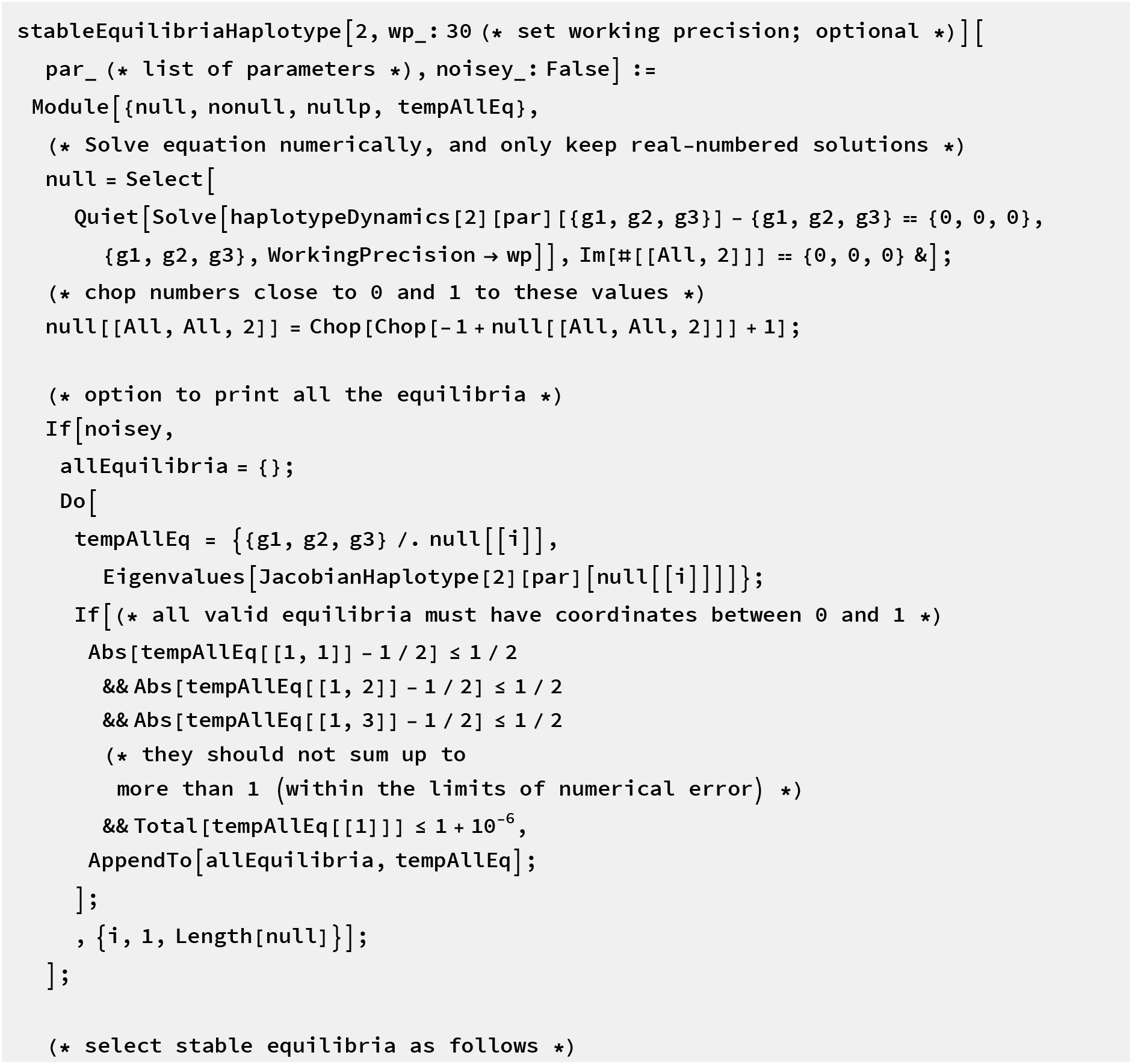
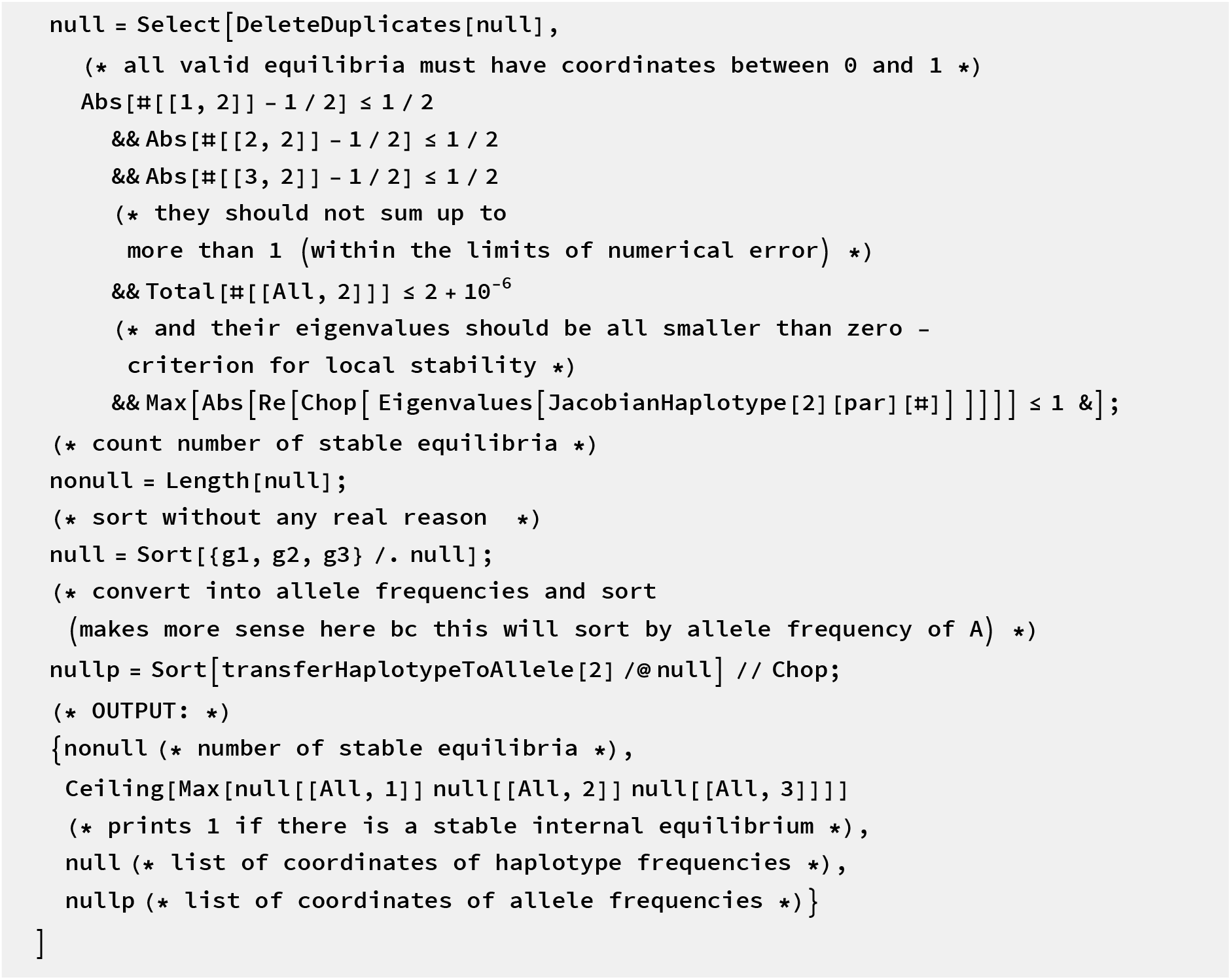

##### Some checks

~~~
Round[stableEquilibriaHaplotype[1][{0.3, 0, 1, 0.5}], 10^−6^] // N
{1., 1., {{{0.2096, 0.52539, 0.05541}, {0.221895, 0.55621, 0.}}},
 {{{0.26501, 0.73499, 0.01482}, {0.221895, 0.778105, 0.049237}}}}
Round[stableEquilibriaHaplotype[2][{0.1, 0.01, 0.05, 0.5}], 10^−6^] // N
{1., 1., {{0.251079, 0.268493, 0.229349}}, {{0.480428, 0.519572, 0.001462}}}
~~~

Looks like we can also find (tri-/)bistability here! (??)

~~~
Round[stableEquilibriaHaplotype [1][{0.05, 0.9, 0.01, 0.1}], 10^−6^] // N
{2., 0., {{{0., 0., 0.409049}, {0., 0., 0.406622}},
 {{0.590951, 0., 0.409049}, {0.593378, 0., 0.406622}}},
{{{0.409049, 0., 0.}, {0.406622, 0., 0.}}, {{1., 0.590951, 0.}, {1., 0.593378, 0.}}}}
Round[stableEquilibriaHaplotype[1][{0.05, 0.8, 0.01, 0.1}], 10^−6^] // N
{3., 1., {{{0., 0., 0.409049}, {0., 0., 0.406622}},
 {{0.238495, 0.015059, 0.50795}, {0.242651, 0.003064, 0.511633}},
  {{0.590951, 0., 0.409049}, {0.593378, 0., 0.406622}}},
{{{0.409049, 0., 0.}, {0.406622, 0., 0.}}, {{0.746445, 0.253555, 0.049231},
 {0.754284, 0.245716, 0.057312}}, {{1., 0.590951, 0.}, {1., 0.593378, 0.}}}}
~~~

#### ▀ Dynamics with assortment

##### Life cycle with assortment via matching function

As compared with the no-preference version, we operate now on genotype and mating pair level rather than genotype and gamete. However, there are some redundancies with the above subsection where we define the dynamics without the preference.

~~~
haplotypes = Tuples[{0, 1}, 2]
genotypes = Tuples[haplotypes, 2]
~~~

~~~
{{0, 0}, {0, 1}, {1, 0}, {1, 1}}
{{{0, 0}, {0, 0}}, {{0, 0}, {0, 1}}, {{0, 0}, {1, 0}}, {{0, 0}, {1, 1}},
 {{0, 1}, {0, 0}}, {{0, 1}, {0, 1}}, {{0, 1}, {1, 0}}, {{0, 1}, {1, 1}},
 {{1, 0}, {0, 0}}, {{1, 0}, {0, 1}}, {{1, 0}, {1, 0}}, {{1, 0}, {1, 1}},
 {{1, 1}, {0, 0}}, {{1, 1}, {0, 1}}, {{1, 1}, {1, 0}}, {{1, 1}, {1, 1}}}
~~~

General genotypes of males and females:

~~~
genotypeFemale = Table [genotypeF_genotypes[[i]]_, {i,16}]
genotypeMale = Table [genotypeM_haplotypes[[i]]_, {i,4}]
~~~

~~~
{genotypeF_{{0,0},{0,0}}_, genotypeF_{{0,0},{0,1}}_, genotypeF_{{0,0}, {1,0}}_, genotypeF_{{0,0}, {1,1}}_,
 genotypeF_{{0,1}, {0,0}}_, genotypeF_{{0,1}, {0,1}}_, genotypeF_{{0,1},{1,0}}_, genotypeF_{{0,1},{1,1}}_,
 genotypeF_{{1,0},{0,0}}_, genotypeF_{{1,0}, {0,1}}, genotypeF_{{1,0},{1,0}}_, genotyPeF_{{1,0},{1,1}}_,
 genotypeF_{{1,1},{0,0}}_, genotypeF_{{1,1>,{0,1}}_, genotypeF_{{1,1>,{1,0}}_, genotypeF_{{1,1},{1,1}}}_
{genotypeM_{0,0}_, genotypeM_{0,1}_, genotypeM_{1,0}_, genotypeM{1,1}_}
~~~

Later we will deal with general diploid genotypes:

~~~
genotypeDiploid = Table[genotypeD_genotypes〚i〛_, {i, 16}]
~~~

~~~
{genotypeD_{{0,0},{0,0}}_, genotypeD_{{0,0},{0,1}}_, genotypeD_{{0,0},{1,0}}_, genotypeD_{{0,0},{1,1}}_,
genotypeD_{{0,1},{0,0}}_, genotypeD_{{0,1}, {0,1}}_, genotypeD_{{0,1},{1,0}}_, genotypeD_{{0,1},{1,1}}_,
genotypeD_{{1,0},{0,0}}_, genotypeD_{{1,0}, {0,1}}_, genotypeD_{{1,0},{1,0}}_, genotypeD_{{1,0},{1,1}}_,
genotypeD_{{1,1},{0,0}}_, genotypeD_{{1,1}, {0,1}}_, genotypeD_{{1,1},{1,0}}_, genotypeD_{{1,1},{1,1}}}_
~~~

We now also need the level of mating pairs:

~~~
matingPairs = Table [matingPair_genotypes〚i〛,haplotypes〚j〛_, {i, 16}, {j, 4}] // Flatten
~~~

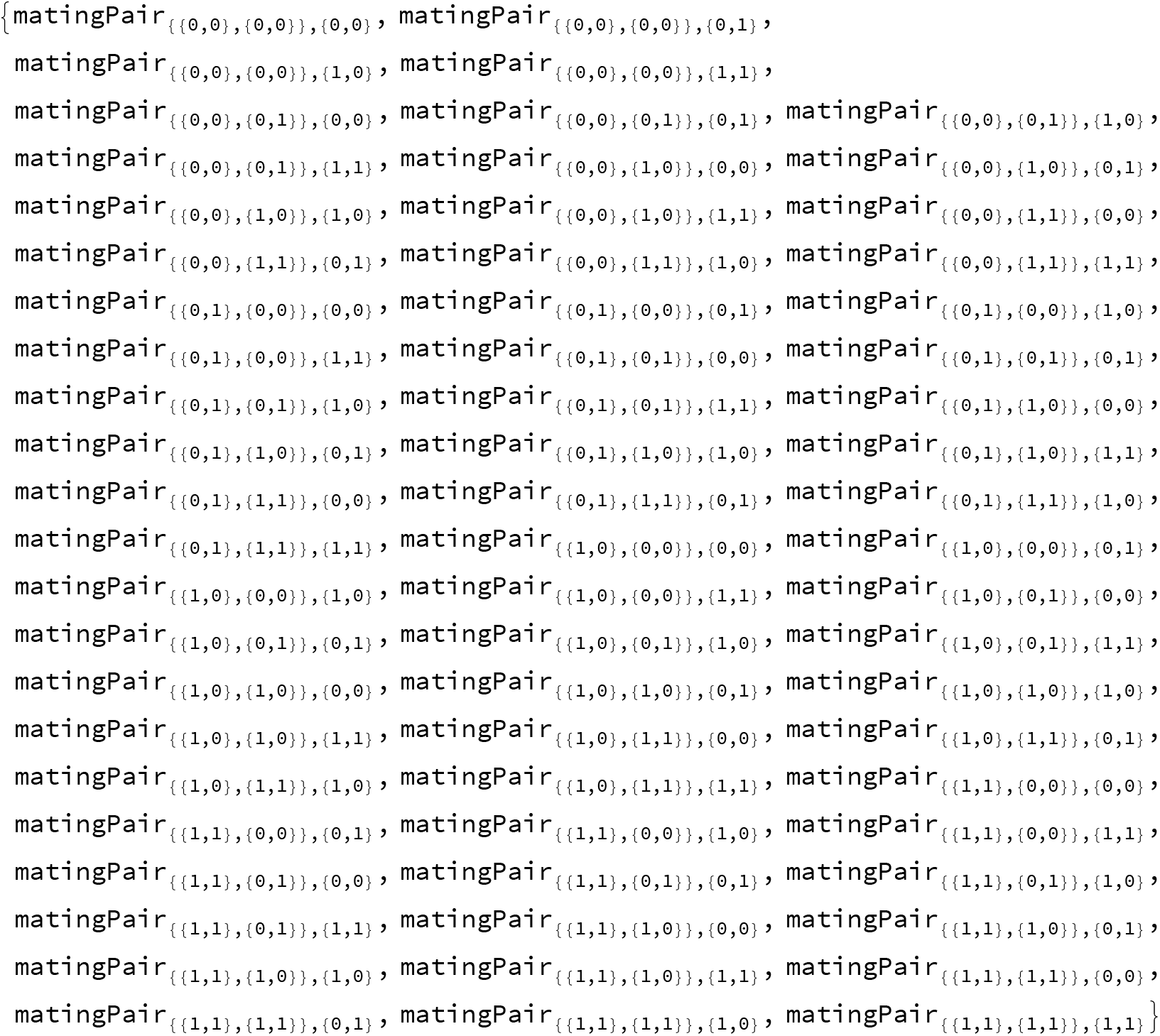

and for the diploid case

~~~
matingPairs = Table [matingPair_genotypes〚i〛,haplotypes〚j〛_, {i, 16}, {j, 4}] // Flatten;
~~~

##### Epistasis

Later, we will usually deal with the case of completely recessive epistasis (using parameter *γ* because ϵ is always handy for Taylor approximations)

~~~
epistasisRecessiveAsymmetric =
 {e_“01”_ → 0, e_“10”_ → 0, e_“01,10”_ → 0, e_“01,10”_ → γ01, e_“10,10”_ → γ10};
~~~

##### Viability selection in males

Note that the frequencies are purely determined by female gamete frequencies in the previous generation.

~~~
selectionMatrixMales = -1, 1 - e”_01_,_01_”, 1 - e”_10_,_10_”, 1} ;
(* notation indicates types of conflict,
here doubled such that haploids suffer as much as homozygous diploids *) meanFitnessMales = Total [selectionMatrixMales genotypeMale] malesAfterSelection = selectionMatrixMales genotypeMale? meanFitnessMales
~~~

~~~
genotypeM_{0;0}_ + (1 - e_01,01_) genotypeM_{0,1}_ + 1 - e_10,10_) genotypeM_{1,0}_ + genotypeM{1,1}
{genotypeM_{0,0}_ /
  (genotypeM_{0,0}_ + (1 - e_01,01_) genotypeM{0,1}, + (1 - e_10,10_) genotypeM{_1,0}_ + genotypeM{1,1}),
 ((1 - e_01,01_) genotypeM_{0,1}_) /
(genotypeM_{0,0}_ + (1 - e_01,01_) genotypeM_{0,1}_ + (1 - e_10,10_) genotypeM_{1,0}_ + genotypeM_{1,1}_), ((1 - e_10,10_) genotypeM_{1,0}_) (genotypeM_{0,0}_ + (1 - e_01,01_) genotypeM_{0,1}_ + (1 - e_10,10_) genotypeM_{1,0}_ + genotypeM_{1,1}_), genotypeM_{1,1}_/
(genotypeM_{0,0}_ + (1 - e_01,01_) genotypeM_{0,1}_ + (1 - e_10,10_) genotypeM_{1,0}_ + genotypeM_{1,1}_)}
~~~

##### Viability selection in females, based on genotypes

For strong selection, this scheme should be multiplicative. Also, we can integrate different types of epistasis. This is the general form:

~~~
selectionMatrixFemales = {{1, (1 - e_“01”_) (1 + s), (1 - e_“10”_) (1 + s), (1 + s)^2^ (1 - e_“01,10”_)},
 {(1 + s) (1 - e_“01”_), (1 - e_“01,01”_), (1 + s)^2^ (1 - e_“01,10”_), (1 + s) (1 - e_“01”_)},
 {(1 + s) (1 - e_“10”_), (1 - e_“01,10”_), (1 + s)^2^ (1 - e_“10,10”_), (1 + s) (1 - e_“01”_)},
 {(1 + s)^2^ (1 - e_“01,10”_), (1 + s) (1 - e_“10”_),}},
selectionTableFemales = Flatten[selectionMatrixFemales];
wShow = Prepend[selectionMatrixFemales, {{A0, B0}, {A0, B1}, {A1, B0}, {A1, B1}}];
Grid[MapThread[Prepend,
  {wShow, {“Haplotypes”, {A0, B0}, {A0, B1}, {A1, B0}, {A1, B1}}}], Frame → All]
~~~

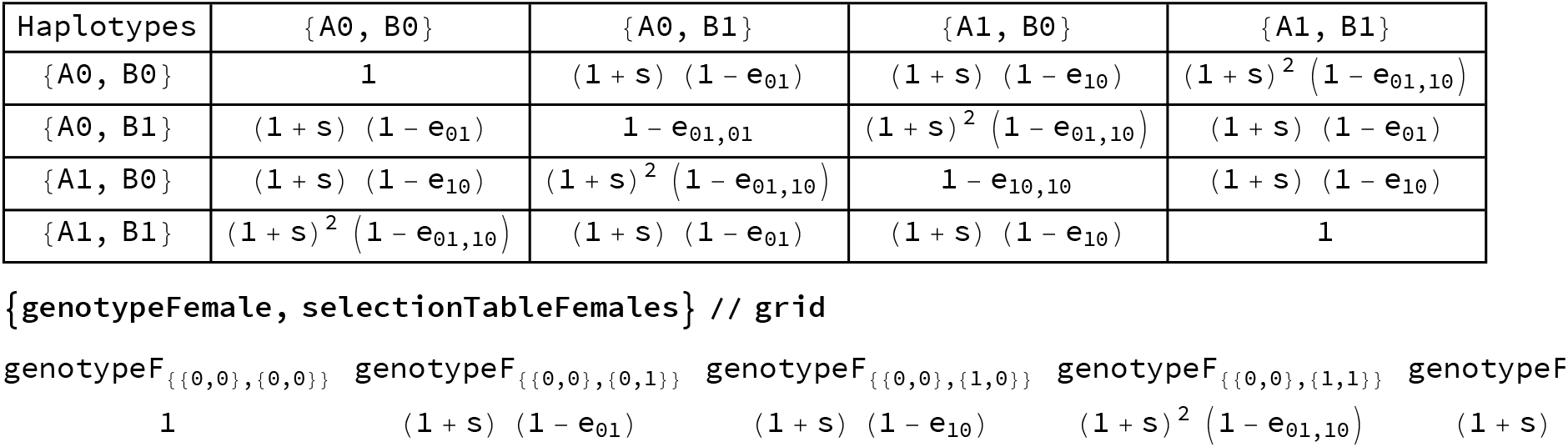

Selection of genotypes

~~~
femaleGenotypesAfterSelectionBeforeNormalization =
 selectionTableFemales genotypeFemale;
meanFitnessFemales = Total[femaleGenotypesAfterSelectionBeforeNormalization];
femaleGenotypesAfterSelection =
 femaleGenotypesAfterSelectionBeforeNormalization? meanFitnessFemales // Simplify;
~~~

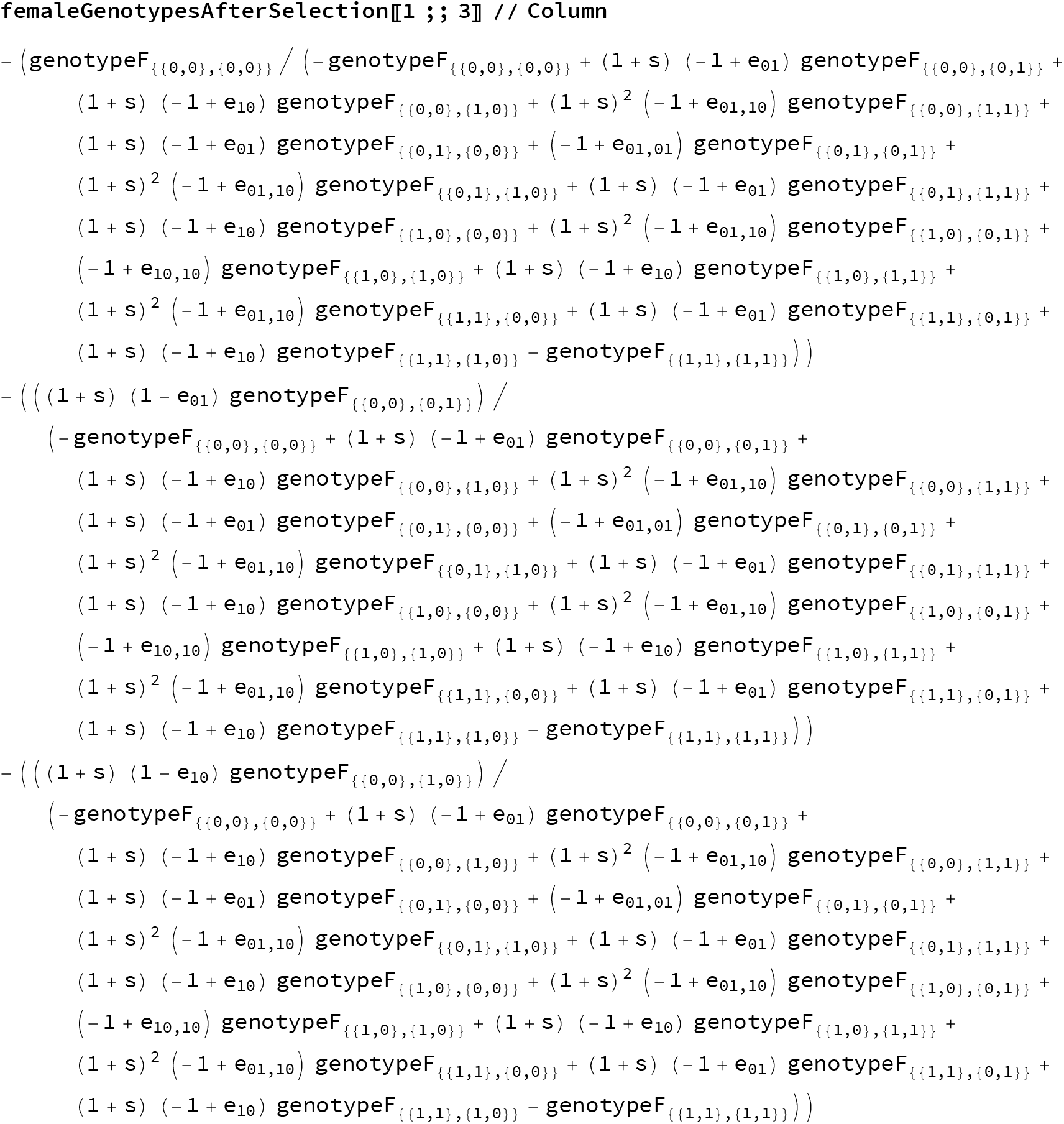

That looks correct! The same is of course happening in the diploid model, we just need to replace the variable:

~~~
diploidGenotypesAfterSelection =
  femaleGenotypesAfterSelection /. {genotypeF_x__ → genotypeD_x_};
~~~

##### Assortment and recombination

~~~
geneticDistance[F_, M_] := Total[Abs[Total [F] /2 - M]]
(* Total[F] will sum the values inside each of the two “locus” tuples,
then divide this value by 2 because females have
 a genome that’s twice the size of that of the males’.
For example, Total[{{0,0},{1,1}}]/2 will return {0.5,0.5}.
Then get the sum of the distances for female vs male at locus1 and female vs male at locus2. *)
~~~

For diploids

~~~
geneticDistanceDiploid[F_, M_] := Total[Abs[Total [F] - Total[M]] / 2]
~~~

~~~
geneticDistance[{{1, 1}, {0, 0}}, {0, 0}]
1
geneticDistanceDiploid[{{1, 1}, {0, 0}}, {{1, 1}, {1, 1}}]
1
~~~

##### Assortative mating

This is what they call “Gaussian assortative mating” in De Cara et al. 2008.

~~~
assortmentCoefficientsGaussian [F_, M_] := (1 - A) ^geneticDistance[F,M]^
~~~

And this is what they call “quadratic assortative mating”:

~~~
assortmentCoefficientsQuadratic[F_, M_] := 1 - AgeneticDistance[F, M] /2
assortmentCoefficientsQuadraticDiploid[F_, M_] := 1 - AgeneticDistanceDiploid[F, M] /2
~~~

We will be using quadratic assortment for now, because Gaussian assortative mating is problematic for introgressed females (for A=1, they would not be able to mate at all). The maximum assortment coefficient is 1 for both of these function, i.e. A ranges from 0 (random mating) to 1 (strong assortment, no mating with opposite type).

~~~
assortmentCoefficientsTable =
Table[assortmentCoefficientsQuadratic[matingPairs〚i, 2〛, matingPairs〚i, 3〛],
 {i, Length[matingPairs]}]
~~~

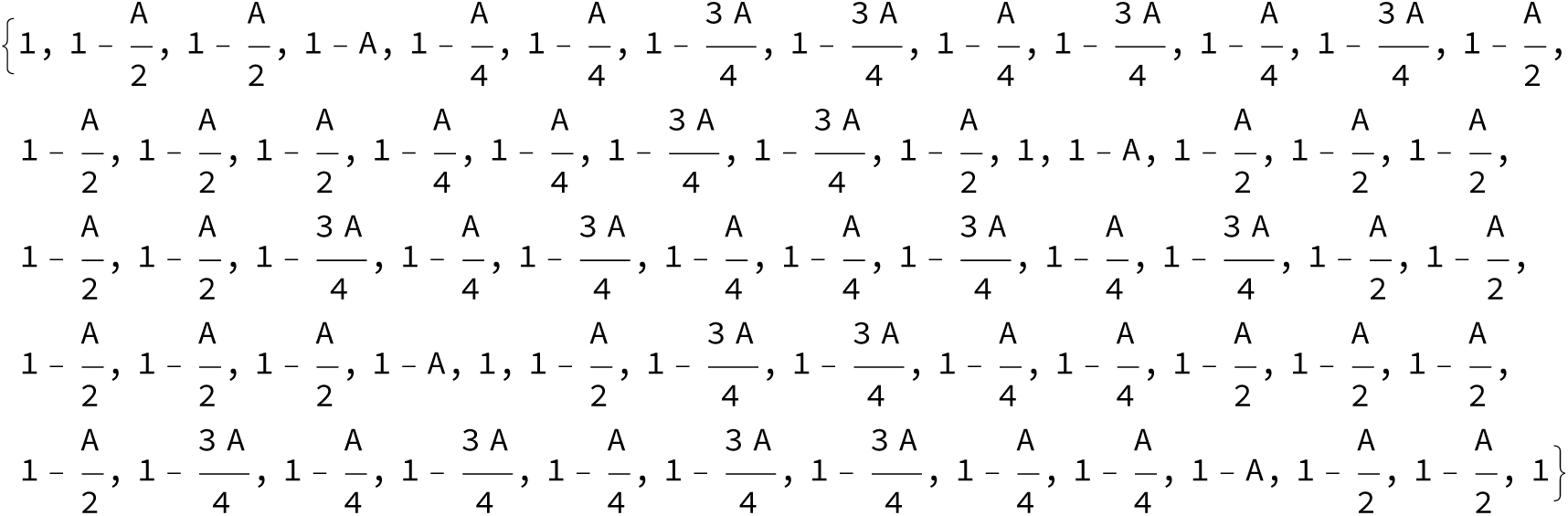

For diplo-diploids

~~~
assortmentCoefficientsTableDiploid =
 Table[assortmentCoefficientsQuadraticDiploid[matingPairsDiploid〚i, 2〛,
  matingPairsDiploid〚i, 3〛], {i, Length[matingPairsDiploid]}];
~~~

Mating pairs are created from males and females after selection and then weighed by the assortment coefficients. Be aware that this is very slow, but the simplification is needed to subsequently speed up the simulation code.

~~~
matingPairFrequenciesBeforeNormalization =
 assortmentCoefficientsTable Flatten[Table[femaleGenotypesAfterSelection〚i〛
    malesAfterSelection〚j〛, {i, 16}, {j, 4j]] // Simplify;
totalForNormalization = Total[matingPairFrequenciesBeforeNormalization] // Simplify;
matingPairFrequencies =
  matingPairFrequenciesBeforeNormalization/ totalForNormalization/ Simplify; matingPairFrequencies〚3〛
~~~

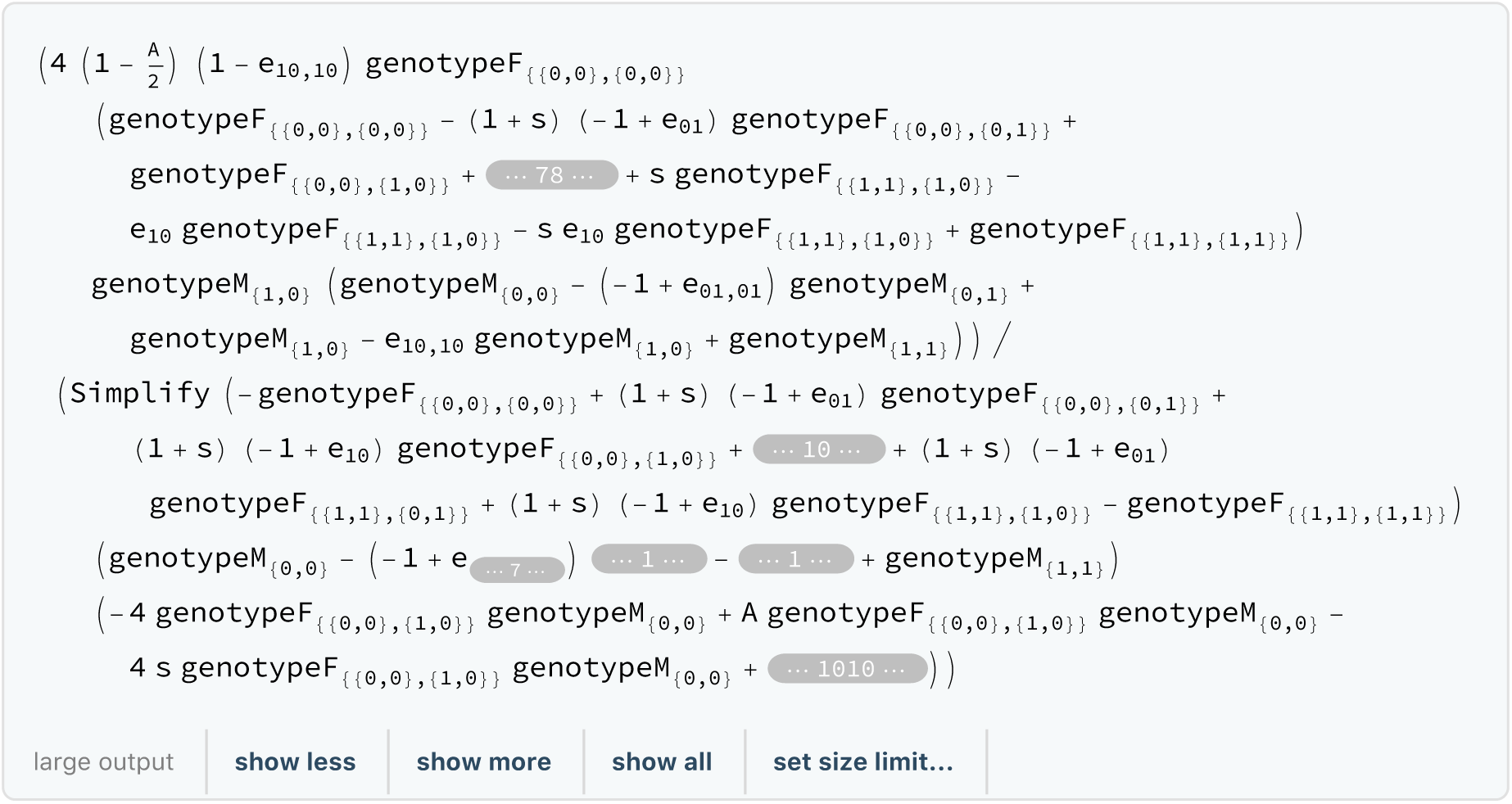

~~~
Sum[matingPairFrequencies〚i〛, {i, Length[matingPairs]}] // Simplify
1
~~~

Same for diploids:

~~~
matingPairFrequenciesDiploid = assortmentCoefficientsTableDiploid
 Flatten[Table[diploidGenotypesAfterSelection〚i〛
  diploidGenotypesAfterSelection〚j〛, {i, 16j, {j, 16j]];
matingPairFrequenciesDiploid〚3〛 // Simplify
~~~

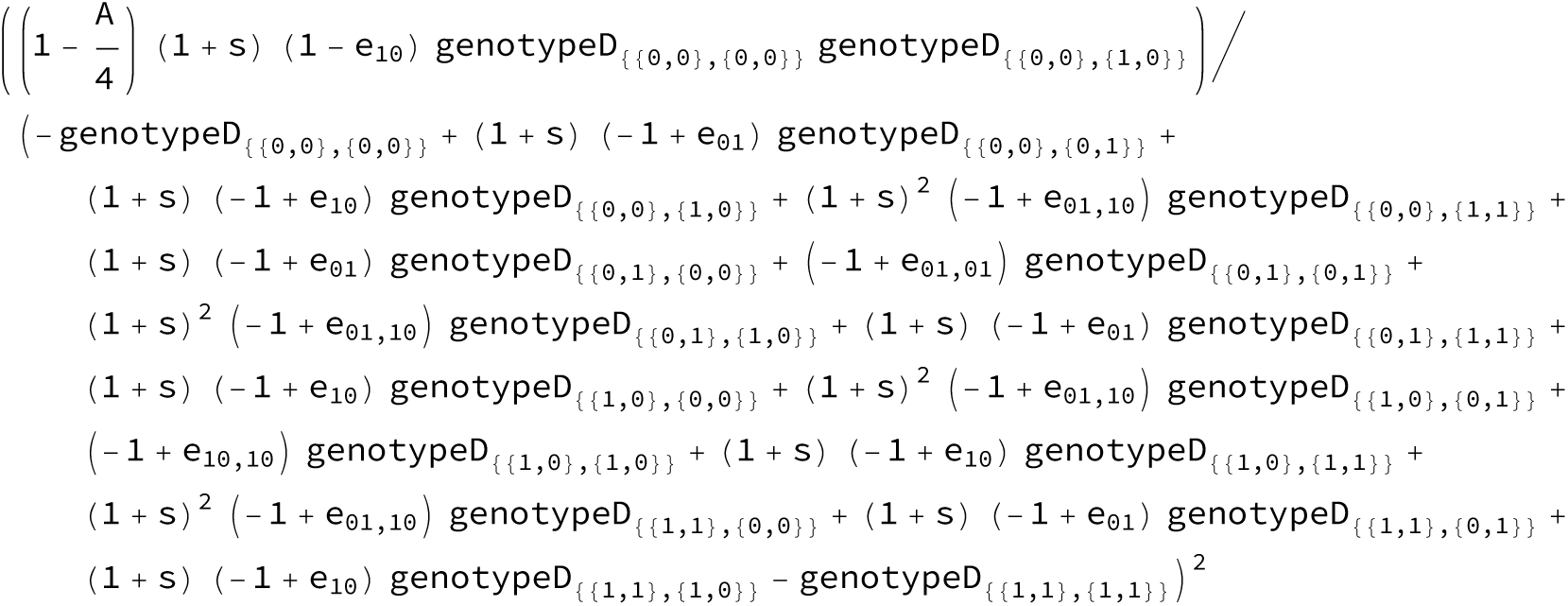

~~~
matingPairFrequenciesDiploidBeforeNormalization =
  assortmentCoefficientsTableDiploid Flatten[Table[diploidGenotypesAfterSelection〚i〛
      diploidGenotypesAfterSelection〚j〛, {i, 16}, {j, 16}]] // Simplify;
totalForNormalizationDiploid = Total[
    matingPairFrequenciesDiploidBeforeNormalization] // Simplify;
matingPairFrequenciesDiploid = matingPairFrequenciesDiploidBeforeNormalization /
    totalForNormalizationDiploid // Simplify;
matingPairFrequenciesDiploid〚
3〛
~~~

$Aborted

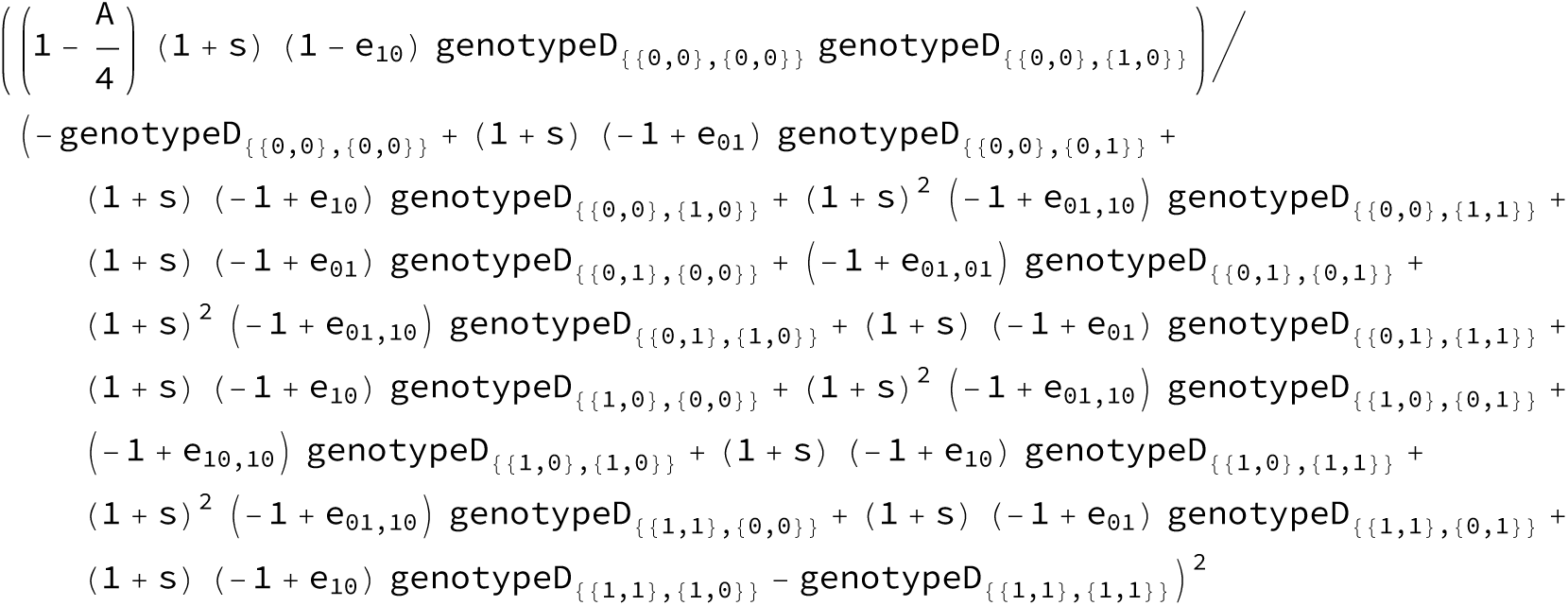

~~~
Sum[matingPairFrequenciesDiploid〚i〛, {i, Length[matingPairsDiploid]}] // Simplify
1
~~~

##### Recombination

Now, new females are created from recombined female gametes and male haplotypes of mating pairs.

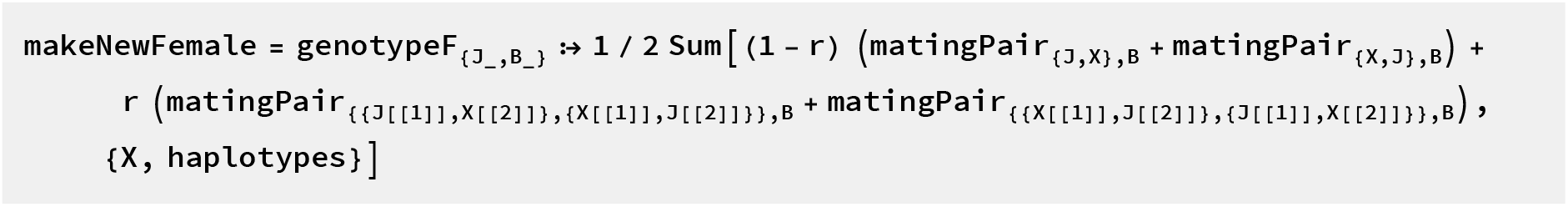

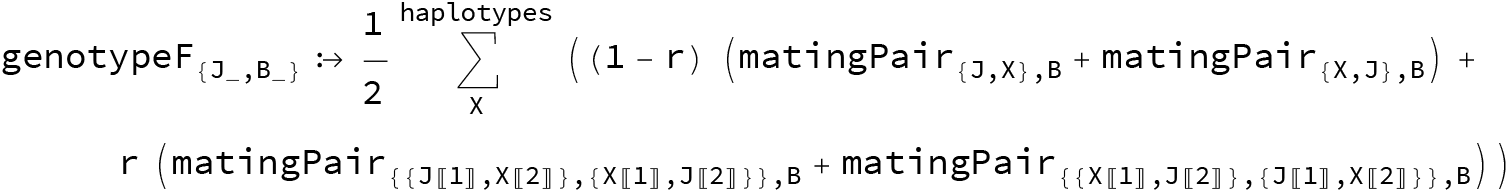

Males are created only from recombined female gametes.

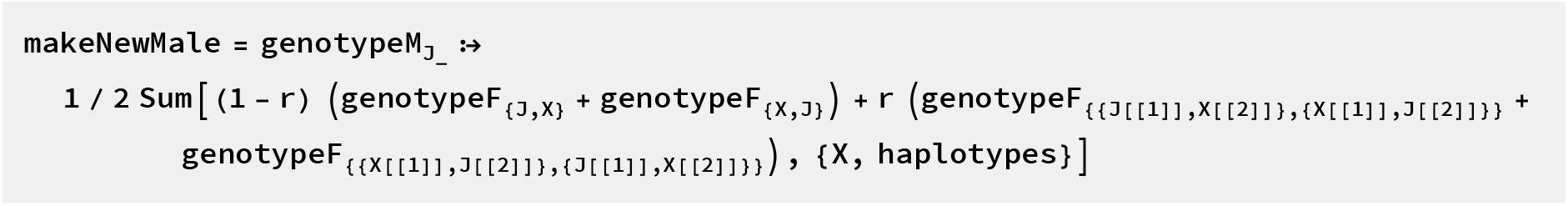

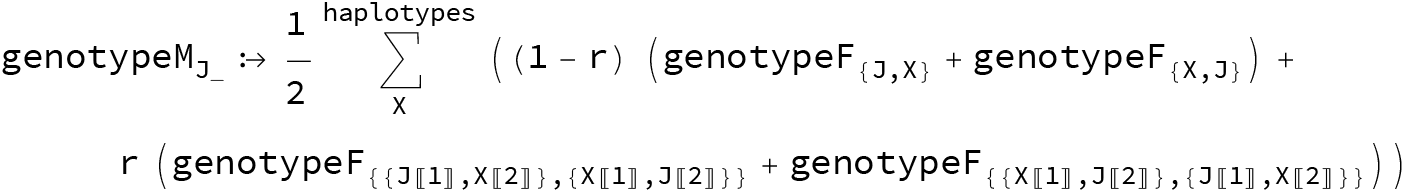

Try out:

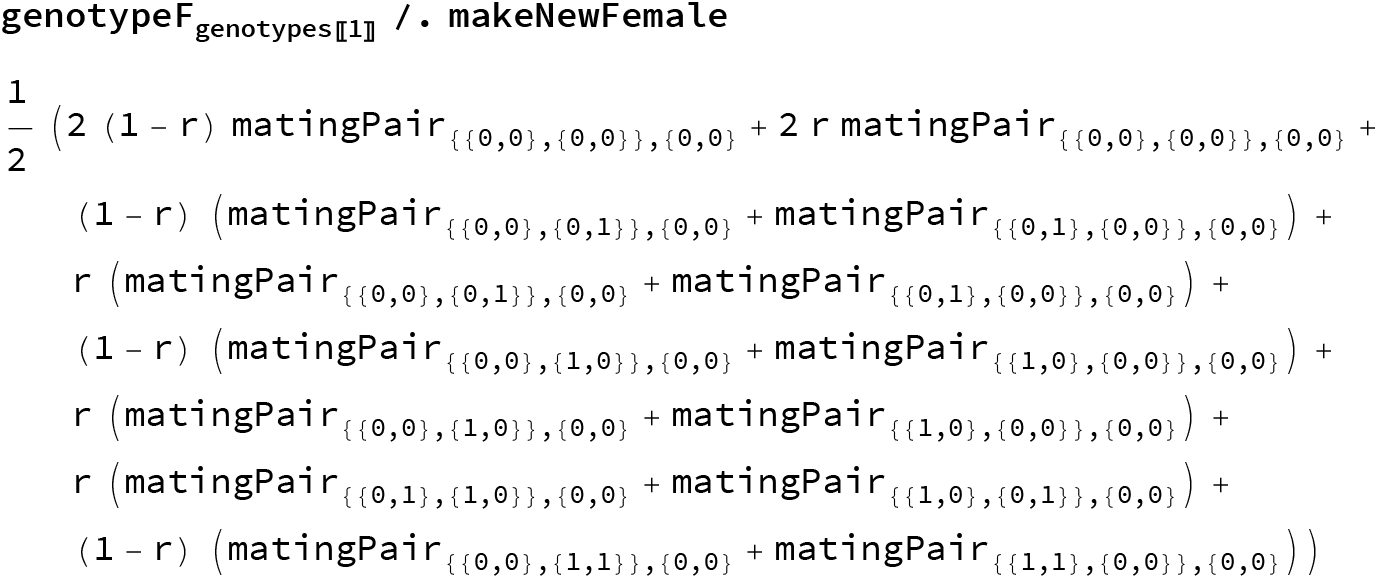

It is a bit more difficult to make new diploids, because both male and female parents can recombine:

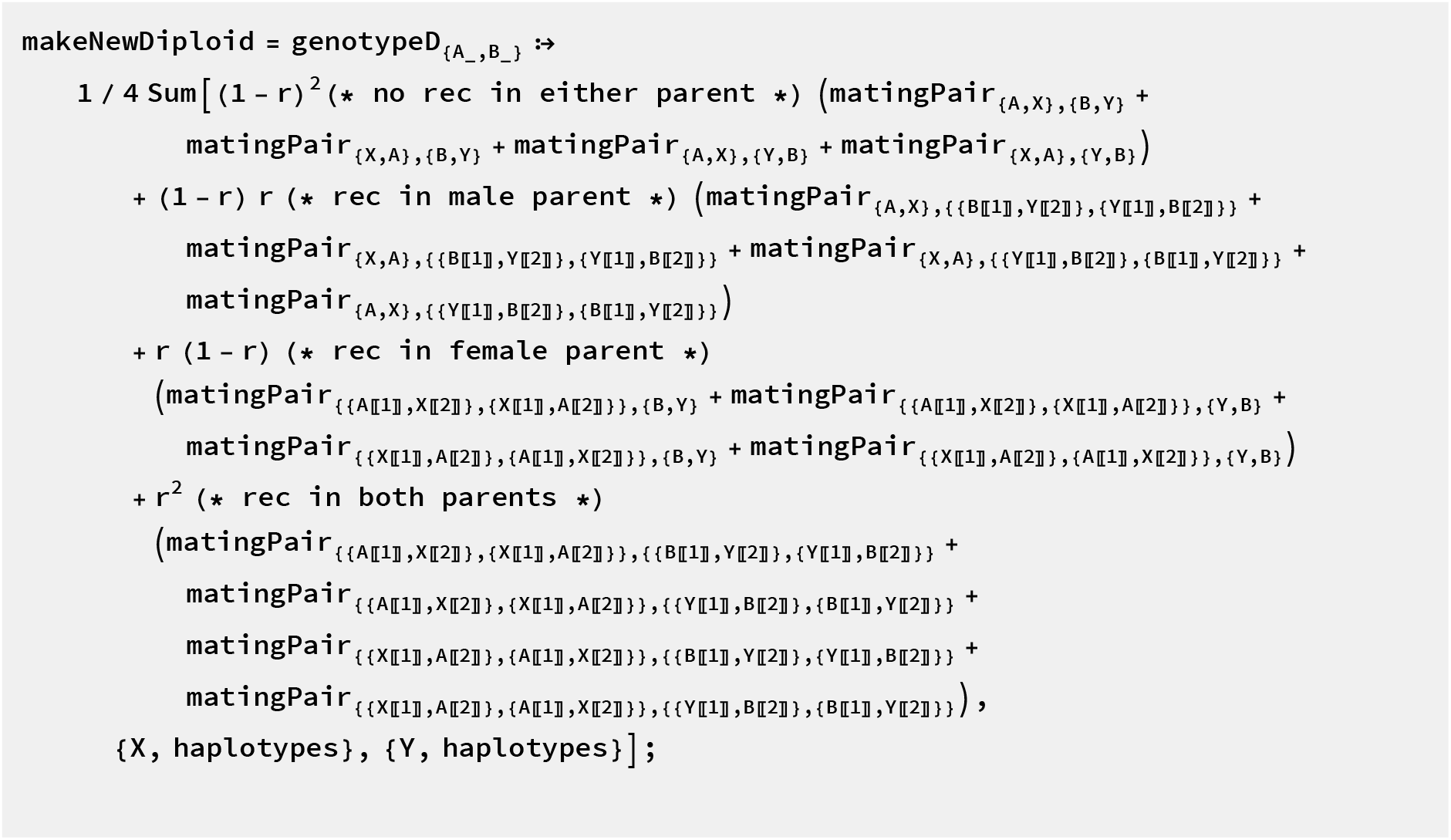

We can now combine the pieces of the life cycle to the actual dynamics:

~~~
dynamicsUnnormalizedWithMatching = {(* new female genotype arises from *)
  genotypeFemale /. makeNewFemale(* recombination and sampling from
    mating pairs *) /. Thread[matingPairs → matingPairFrequencies]
  (* input of actual mating pair frequencies after selection *),
  (* new male genotype arises from *)
   genotypeMale /. makeNewMale(* recombination of female gametes *) /.
   Thread[genotypeFemale → femaleGenotypesAfterSelection]
   (* input of female frequencies after selection *)j /.
epistasisRecessiveAsymmetric(* reduce epistasis coefficients *) // Simplify;
(* keep in mind that this has to be normalized
to 1 *)
dynamicsTotal = Total /@ dynamicsUnnormalizedWithMatching // Simplify
 (* compute total for normalization *) ;
~~~

An example:

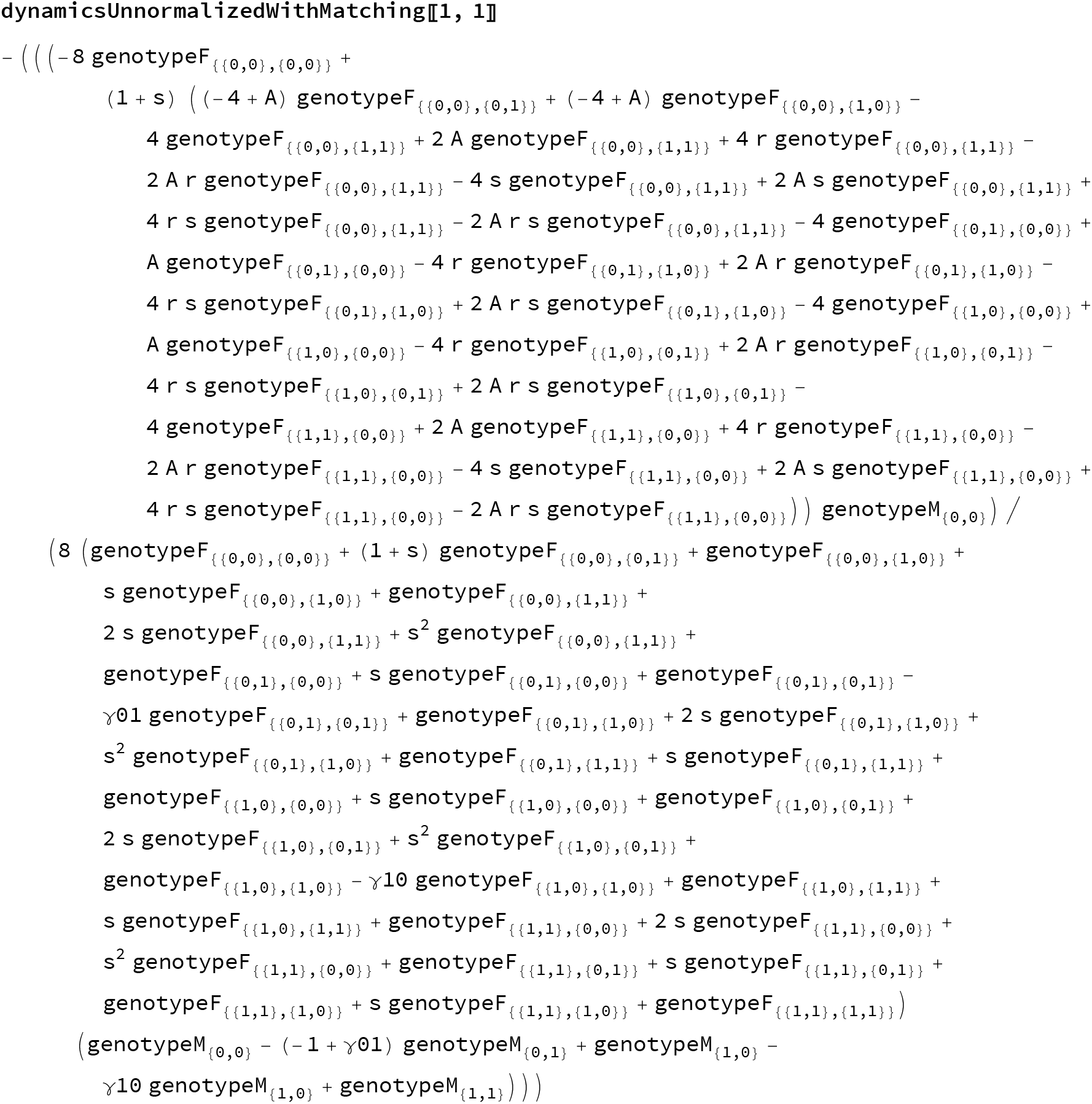

This takes a while to run. You can save the dynamical equations for later, or cut a couple of simplification steps.

~~~
Import[ToString[NotebookDirectory[]] <>“dynamicsWithPref_saved”]
(* this imports the file called “dynamicsWithPref_saved”
so long as it is in the same directory as this notebook *)
~~~

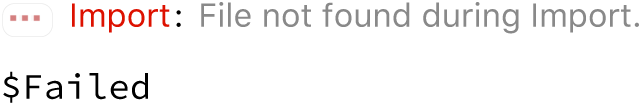

The diploid dynamics are slightly simpler:

~~~
dynamicsUnnormalizedWithMatchingDiploid = (* new female genotype arises from *)
 genotypeDiploid /. makeNewDiploid (* recombination and sampling from mating
    pairs *) /. Thread[matingPairsDiploid → matingPairFrequenciesDiploid]
 (* input of actual mating pair frequencies after selection *) /.
 epistasisRecessiveAsymmetric(* reduce epistasis coefficients *) // Simplify;
dynamicsTotalDiploid = Total /@dynamicsUnnormalizedWithMatchingDiploid
 (* compute total for normalization *) ;
$Aborted
~~~

#### Dynamics as functions of haplotype frequencies

##### Haplodiploid dynamics as function of female gametes and male frequencies after selection

Create some genotype frequencies for testing purposes:

~~~
testF = RandomVariate[UniformDistribution[], 16];
testF = testF / Total[testF];
testM = RandomVariate[UniformDistribution[], 4] ;
testM = testM / Total[testM];
~~~

Dynamical equations:

~~~
dynamicsHaplodiploidWithMatching[{α_, σ_, γ1_, γ2_, R_}] [
  {femaleGenotypes_, maleGenotypes_}] :=
dynamicsUnnormalizedWithMatching/ dynamicsTotal /.
  Thread[genotypeFemale → femaleGenotypes] /. Thread[
  genotypeMale → maleGenotypes] /. Thread[{A, s, γ01, γ10, r} -> {α, σ, γ1, γ2, R}]
~~~

Test:

~~~
dynamicsHaplodiploidWithMatching[{1, 0.1, 0.01, 0.02, 0.5}][{testF, testM}]
{{0.0840051, 0.0790078, 0.0348801, 0.0214843, 0.0918598, 0.175376, 0.0243312, 0.0530175,
0.072139, 0.0358112, 0.0673231, 0.038687, 0.0319385, 0.090978, 0.0380982, 0.0610633},
{0.209395, 0.315888, 0.239392, 0.235324}}
~~~

#### Diploid dynamics as function of gamete frequencies after selection

~~~
dynamicsDiploidWithMatching[{α_, σ_, γ1_, γ2_, R_}][genotypes_] :=
 dynamicsUnnormMatchingDiploidSimplified / dynamicsTotalDiploid /. Thread[
   genotypeDiploid → genotypes] /. Thread[{A, s, γ01, γ10, r} -> {α, σ, γ1, γ2, R}]
~~~

Test:

~~~
dynamicsDiploidWithMatching[{1, 0.1, 0.01, 0.02, 0.5}][testF] // N
{0.116377, 0.13204, 0.111305, 0.101934, 0.13204, 0.165403, 0.114043, 0.136624,
0.111305, 0.114043, 0.132534, 0.112855, 0.101934, 0.136624, 0.112855, 0.130467}
~~~

You can also save everything in the notebook that you have calculated so far, in order to “resurrect” it later.

~~~
DumpSave[ToString[NotebookDirectory[]] <>
   “ant-dynamics_haplodiploid&diploid_with-preference.mx”, “Global”] ;
~~~

### Chapter 2: Evolutionary scenarios

For the model without preference, use the function **stableEquilibriaHaplotype** to solve for all of the equilibria. Then investigate the stable equilibria in order to assign the given parameter combination to a specific evolutionary scenario.

Here are two functions (for haplodiploids and diploids seperately) that will classify the different evolutionary scenarios from a given parameter combination. Recall that parameter combinations must be inputted as a list in the following order: {*σ*, *γ*1, *γ*2, *ρ*}

~~~
IdentifyCase[pars_] := Module[{allMaleFreqs},
 equilOutput = stableEquilibriaHaplotype[1][pars];
 allMaleFreqs = Sort[Flatten@equilOutput[[4, All, 2, 1 ;; 2]]];
~~~

~~~
(* Now catch the 6 possible cases as well as some putative error cases: *)
Which[equilOutput[[1]] == 1
 && Abs[equilOutput[[4, 1, 1, 1]] − 0.5] < 0.01 &&
 Abs [equilOutput[[4, 1, 1, 2]] − 0.5] < 0.01,
"F.5" (* females win with 0.5 allele freq for both loci *),
equilOutput[[1]] == 1, "F", (* females win but NOT
 near 0.5 allele freq for both loci *)
 equilOutput[[1]] == 5, "MCF"(* depending on intial it's possible that males win,
compromise, or females win *),
equilOutput [[1]] == 2
 && allMaleFreqs == {0, 0, 1, 1}, "M" (* males win *),
equilOutput [[1]] == 2
 && allMaleFreqs[[{1, 4}]] == {0, 1} && allMaleFreqs[[2 ;; 3]] ≈ {0, 1},
"C" (* compromise *),
equilOutput [[1]] == 2
 && Total @ Abs [equilOutput [[4, 1, 1, 1 ;; 2]] − equilOutput[[4, 2, 1, 1 ;; 2]]] < 0.05
 (* stable equilibria are almost on top of each other *),
 "F2" (* near the bistability so 2 equilibria are
 almost about to converge to F case *),
equilOutput [[1]] == 2
 && equilOutput[[2]] == 1, "A" (* asymmetric coexistence *),
equilOutput [[1]] == 2,
"weird bistable" (* if it's bistable but NEITHER C, M, nor A *),
equilOutput [[1]] == 3
 && allMaleFreqs [[{1, 2, 5, 6}]] == {0, 0, 1, 1}
 && 0 < allMaleFreqs[[3]] < 1 && 0 < allMaleFreqs[[4]] < 1,
"MC" (* depending on intial males win or compromise *),
equilOutput [[1]] == 3
 && allMaleFreqs[[{1, 6}]] == {0, 1}
 && 0 < allMaleFreqs[[3]] < 1 && 0 < allMaleFreqs[[4]] < 1,
"CF" (* depending on intial compromise or females win *),
equilOutput[[1]] == 3, "weird tristable" (* if it'
 s tristable but NEITHER CF nor MC *),
True, “weird other” (* none of the above cases are met *)]
]
~~~

~~~
IdentifyCaseDIPLOID[pars_] := Module[{},
 equilOutput = stableEquilibriaHaplotype [2][pars];
~~~

~~~
Which[equilOutput[[1]] == 1
  && Abs[equilOutput[[4, 1, 1]] − 0.5] < 0.01 &&
  Abs [equilOutput[[4, 1, 2]] − 0.5] < 0.01 (* both loci near 0.5 *)
  && Chop[1 − equilOutput[[4, 1, 1]] − equilOutput[[4, 1, 2]], 10^− 6] == 0
 (* point sits on line y=1-x *),
 "F.5" (* coexistence near 0.5 allele freq for both loci *), equilOutput [[1]] == 1
  && Chop[1 − equilOutput[[4, 1, 1]] − equilOutput[[4, 1, 2]], 10^− 6] == 0
 (* point sits on line y=1-x *),
 "F" (* coexistence but not near 0.5 allele freq for both loci *),
 equilOutput[[1]] == 1, "weird unistability",
 equilOutput[[1]] == 2
  && equilOutput[[4, 1, 1]] < 0.01 && equilOutput[[4, 1, 2]] < 0.01
 (* both loci within 1% of 0 or 1 allele freq *)
  && equilOutput[[4, 2, 1]]> 0.99 && equilOutput[[4, 2,2]]> 0.99,
 "M'(* exclusion-like state *),
 equilOutput[[1]] == 2
  && Total@Abs [equilOutput [[4, 1, 1 ;; 2]] − equilOutput[[4, 2, 1 ;; 2]]] < 0.05
 (* stable equilibria are almost on top of each other *)
, "F2" (* near the bistability so 2 equilibria are
  almost about to converge to F.5 case *),
 equilOutput [[1]] == 2
  && Chop[1 − equilOutput [[4, 1,2]] − equilOutput [[4, 2, 1]], 10^ −6] == 0 &&
  Chop[1 − equilOutput [[4, 1,1]] − equilOutput [[4, 2, 2]], 10^ −6] == 0
 (* equilibria are symmetrical within numerical precision*)
, "A" (* asymmetric coexistence *),
 equilOutput [[1]] == 2, "weird bistability",
 equilOutput [[1]] ==3, “weird tristability”
 ]
]
~~~

Here are two functions (for haplodiploids and diploids seperately) that will plot the phase plane for a given parameter combination.

Note that the phase plane diagrams given in the manuscript also show the basin of attraction. The basin of attraction was done by simulation; see Chapter 5 section “Basin of attraction” below.

~~~
PlotPhasePlane[pars_, PlotFemales_: False, allBlack_: True] : =
 Module[{equilFreqFemalesMales, unstableEquil, unstableHaps},
  (* get the equilibria *)
  equilFreqFemalesMales =
   stableEquilibriaHaplotype[1][pars, True][[4, All, All, 1 ;; 2]];
  (* calculate the allele freq for stable *)
  If [PlotFemales == True,
   (* just use the female allel freq *)
   stableFreq = equilFreqFemalesMales[[All, 1, All]];,
   (* use the allele freq for males and females combined *)
   stableFreq = 2/3 *equilFreqFemalesMales[[All, 1, All]] +
    1/3 *equilFreqFemalesMales[[All, 2, All]];
];
~~~

~~~
unstableEquil = Select[DeleteDuplicates[allEquilibria],
   (* one of the eigenvalues should be bigger than one -
   criterion for local instability *)
  Max[Re[Chop[#[[2]] ]]] > 1 &][[All, 1]];
(* keep just the equilibrium frequencies of the haplotypes *)
(* count allele frequencies then combine by males and females *) unstableHaps = Partition[#, 3] & /@ unstableEquil;
If [PlotFemales == True,
  (* just use the female allele freq *)
  unstableFreq =
   Thread[transferHaplotypeToAllele[1][unstableHaps]][[All, 1, 1 ;; 2]];,
  (* use the allele freq for males and females combined *)
  unstableFreq = Total[#*{2 / 3,1 / 3}] & /@
   Thread[transferHaplotypeToAllele[1][unstableHaps]][[All, All, 1 ;; 2]];
];
~~~

~~~
Which [allBlack == True,
 plot = Show[ListPlot[stableFreq, Frame → True, AspectRatio → 1,
     PlotRange → {{0, 1}, {0, 1}}, PlotRangeClipping → False, ImagePadding → 65,
     FrameTicks → {{{{0,”0 “}, {1, “1 “}}, None}, {{{0, “\n0”}, {1, “\n1”}}, None}},
     FrameLabel → (Style[#, FontSize → 14.5] &/@
        {“Frequency of Allele A_w_”, “Frequency of Allele B_w_”}), PlotLabel → pars,
     LabelStyle → {FontFamily -> “Times”, FontSize → 16}, ImageSize → 250],
   Graphics[{PointSize[0.15], Point[stableFreq]}],
   Graphics[{PointSize[0.15], Point[unstableFreq]}],
   Graphics[{White, PointSize[0.12], Point[unstableFreq]}]]
,
 allBlack == False,
 plot = Show[ListPlot[stableFreq, Frame → True, AspectRatio → 1,
    PlotRange →{{0, 1}, {0, 1}}, PlotRangeClipping → False, ImagePadding → 65,
    FrameTicks →{{{{0,”0 “}, {1, “1 “}}, None}, {{{0, “\n0”}, {1, “\n1”}}, None}},
    FrameLabel → (Style[#, FontSize → 14.5] &/@
        {“Frequency of Allele A_w_”, “Frequency of Allele B_w_”}), PlotLabel → pars,
    LabelStyle → {FontFamily -> “Times”, FontSize → 16}, ImageSize → 250],
   Graphics[{PointSize[0.15], Point[stableFreq]}],
   Graphics[{Darker[Red], PointSize[0.15], Point[unstableFreq]}],
   Graphics[{White, PointSize[0.12], Point[unstableFreq]}]]
 ];
 Return[plot];
]
~~~

~~~
PlotPhasePlaneDIPLOID[pars_] := Module[{unstableEquil, unstableFreq},
 (* get the equilibria *)
 stableFreq = stableEquilibriaHaplotype[2][pars, True][[4, All, 1 ;; 2]];

 (* get the unstable equilibria *)
 unstableEquil = Select[DeleteDuplicates[allEquilibria],
    (* one of the eigenvalues should be bigger than one - criterion for local instability *)
    Max[Re[Chop[#[[2]] ]]] > 1 &][[All, 1]];
 (* keep just the equilibrium frequencies of the haplotypes *)
 (* convert to allele freq *)
 unstableFreq = Thread[transferHaplotypeToAllele[2][unstableEquil]][[All, 1 ;; 2]];

 Show[ListPlot[stableFreq, Frame → True, AspectRatio → 1,
   PlotRange →{{0, 1}, {0, 1}}, PlotRangePadding → 0.02, FrameLabel →
    {“Frequency of Allele 1 at Locus 1”, “Frequency of Allele 1 at Locus 2”},
   PlotLabel → “Diploid “ <> ToString@pars],
  Graphics[{PointSize[0.04], Point[stableFreq]}],
  Graphics[{Darker[Red], PointSize[0.025], Point[unstableFreq]}],
  Graphics[{White, PointSize[0.01], Point[unstableFreq]}]]
]
~~~

If *α*=0 (no preference) and we don’t care about the time to convergence or any specific starting conditions, we can use solve to get the haplotype frequencies before and after selection, as well as the LD at the equilibria.

~~~
noPrefEquilGenotypes[pars_, identifyCase_: False, summarizeGen_: True] : =
 Module[{equilOutput, allMaleFreqs, temp, out, gentempF, gentemp, gentempAfterSel,
   femalesPostSel, malesPostSel, LD}, (* pars are: σ, γ1, γ2, r *)
  equilOutput = stableEquilibriaHaplotype[1][pars // N];

  (* this is the part that gets the genotypes before and after selection from gamete frequencies *)
  temp = equilOutput [[3]] // N;
  out = {};
  (* functions to apply selection on newborns *)
  femalesPostSel = femaleGenotypesAfterSelection /. epistasisRecessiveAsymmetric /.
     Thread [{s, γ01, γ10} → pars [[1 ;; 3]]];
  malesPostSel = malesAfterSelection /. epistasisRecessiveAsymmetric /.
     Thread [ {γ01, γ10} → pars [[2 ;; 3]]];
~~~

~~~
Do [
  (* go from haplotype frequencies to genotype frequencies *)
  AppendTo[temp[[i, 1]], 1 - Total@temp[[i, 1]]];
  AppendTo[temp[[i, 2]], 1 - Total@temp[[i, 2]]];
  gentempF = Flatten[Table[temp[[i, 1, j]] * temp[[i, 2, k]], {j, 4}, {k, 4}]];
  gentemp = {gentempF, temp[[i, 1]]};
  (* apply selection on newborns *)
  gentempAfterSel = {femalesPostSel /. Thread[genotypeFemale → gentemp[[1]] ],
     malesPostSel /. Thread[genotypeMale → gentemp[[2]] ]};
~~~

~~~
(* save to output *)
If [summarizeGen == True,
  AppendTo[out, {Join[fem7Catfreq@gentemp, temp[[i, 1]]],
    Join[fem7Catfreq@gentempAfterSel, gentempAfterSel[[2]]]}],
  AppendTo[out, {gentemp, gentempAfterSel}]
],
{i, Length@temp}];
~~~

~~~
(* this is the part that categorizes what’s going on at these param values *)
If [identifyCase == True,
  allMaleFreqs = Sort[Flatten@equilOutput[[4, All, 2, 1 ;; 2]]];
  (* Now catch the 6 possible cases as well as some putative error cases: *)
  outputCase = Which[equilOutput[[1]] == 1
     && Abs[equilOutput[[4, 1, 1, 1]] - 0.5] < 0.01 && Abs[equilOutput[[4, 1, 1, 2]] - 0.5] 0.01, “F.5” (* females win with 0.5 allele freq for both loci *),
    equilOutput[[1]] == 1, “F”, (* females win but NOT near
      0.5 allele freq for both loci *)
    equilOutput[[1]] == 5, “MCF”(* depending on intial it’s
     possible that males win, compromise, or females win *),
    equilOutput [[1]] == 2
     && allMaleFreqs == {0, 0, 1, 1}, “M” (* males win *), equilOutput [[1]] == 2 && allMaleFreqs[[{1, 4}]] == {0, 1} && allMaleFreqs[[2 ;; 3]] Φ {0, 1},
   “C” (* compromise; single locus polymorphism *),
   equilOutput [[1]] == 2
     && Total @ Abs [equilOutput [[4, 1, 1, 1 ;; 2]] - equilOutput [[4, 2, 1, 1 ;; 2]]] < 0.05
   (* stable equilibria are almost on top of each other *),
   “F2” (* near the bistability so 2 equilibria are almost about to converge to F case *),
   equilOutput [[1]] == 2
     && equilOutput[[2]] == 1, “A” (* asymmetric coexistence *),
   equilOutput [[1]] == 2,
   “weird bistable” (* if it’s bistable but NEITHER C, M, nor A *),
   equilOutput [[1]] == 3
     && allMaleFreqs[[{1, 2, 5, 6}]] == {0, 0, 1, 1}
     && 0 < allMaleFreqs[[3]] < 1 && 0 < allMaleFreqs[[4]] < 1,
   “MC” (* depending on intial males win or compromise *),
   equilOutput [[1]] == 3
     && allMaleFreqs[[{1, 6}]] == {0, 1}
     && 0 < allMaleFreqs [ [3]] < 1 && 0 < allMaleFreqs[[4]] < 1,
   “CF” (* depending on intial compromise or females win *),
   equilOutput[[1]] == 3, “weird tristable” (* if it’s
     tristable but NEITHER CF nor MC *),
    True, “weird other” (* none of the above cases are met *)];
     LD =
   Partition[Flatten[equilOutput[[4]]][[3 * Range[2 * Length@equilOutput[[4]]]]], 2];
     LD = (2 / 3) #[[1]] + (1 / 3) #[ [2] ] & /@ LD;
  (* weighted average among males and females *)
  Return[{out, outputCase, LD}],
  Return[out] (* this outputs a table of all equilibrium
  freq. First giving BEFORE selection freq then AFTER selection freq. *)
 ]
]
~~~

### Chapter 3: Stability analysis of the model

#### Equilibria and stability analysis for γ=1 in haplodiploids

Write out dynamics for lethal epistasis:

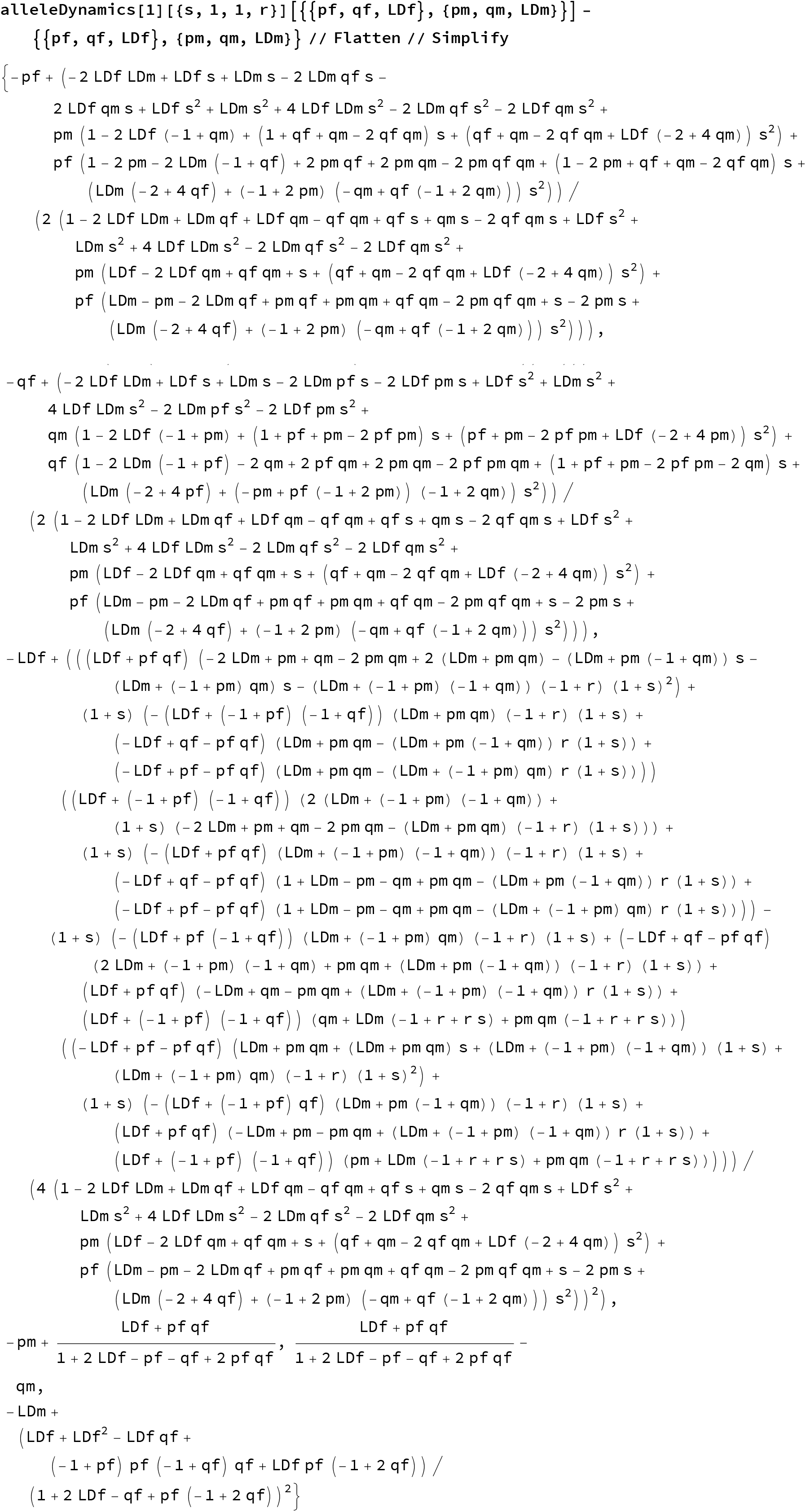

Find equilibria:

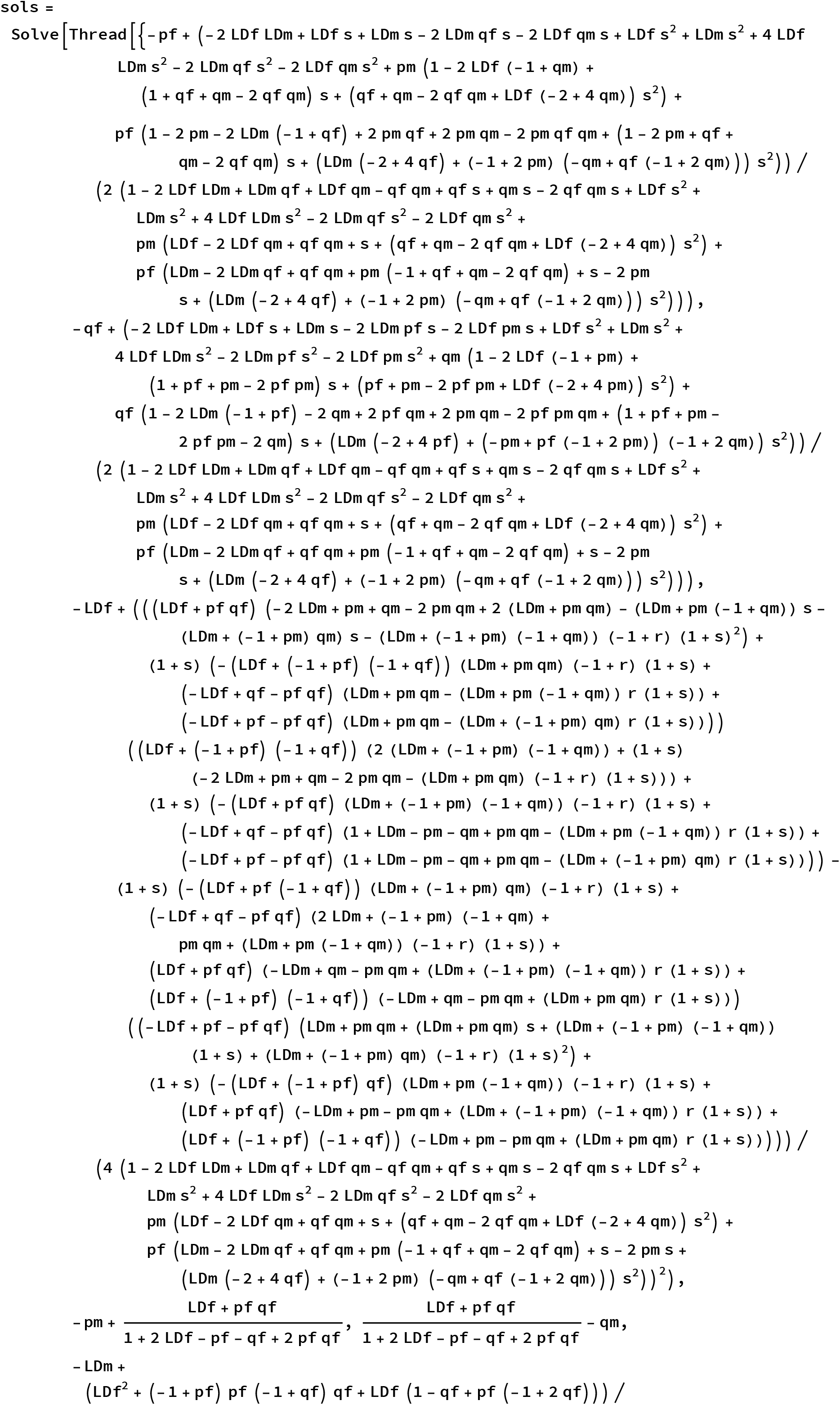
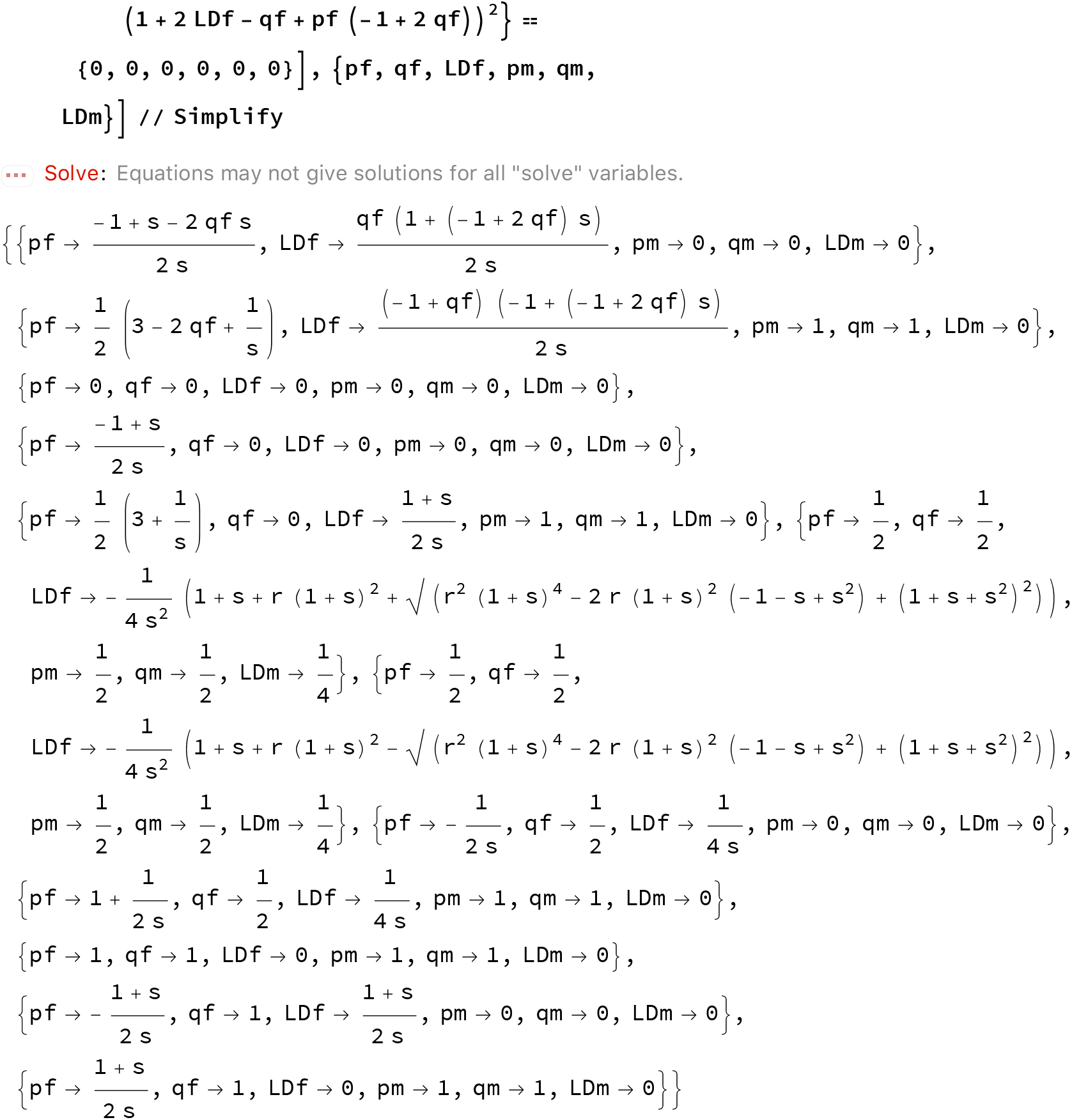

For systematic exploration add solution for qf

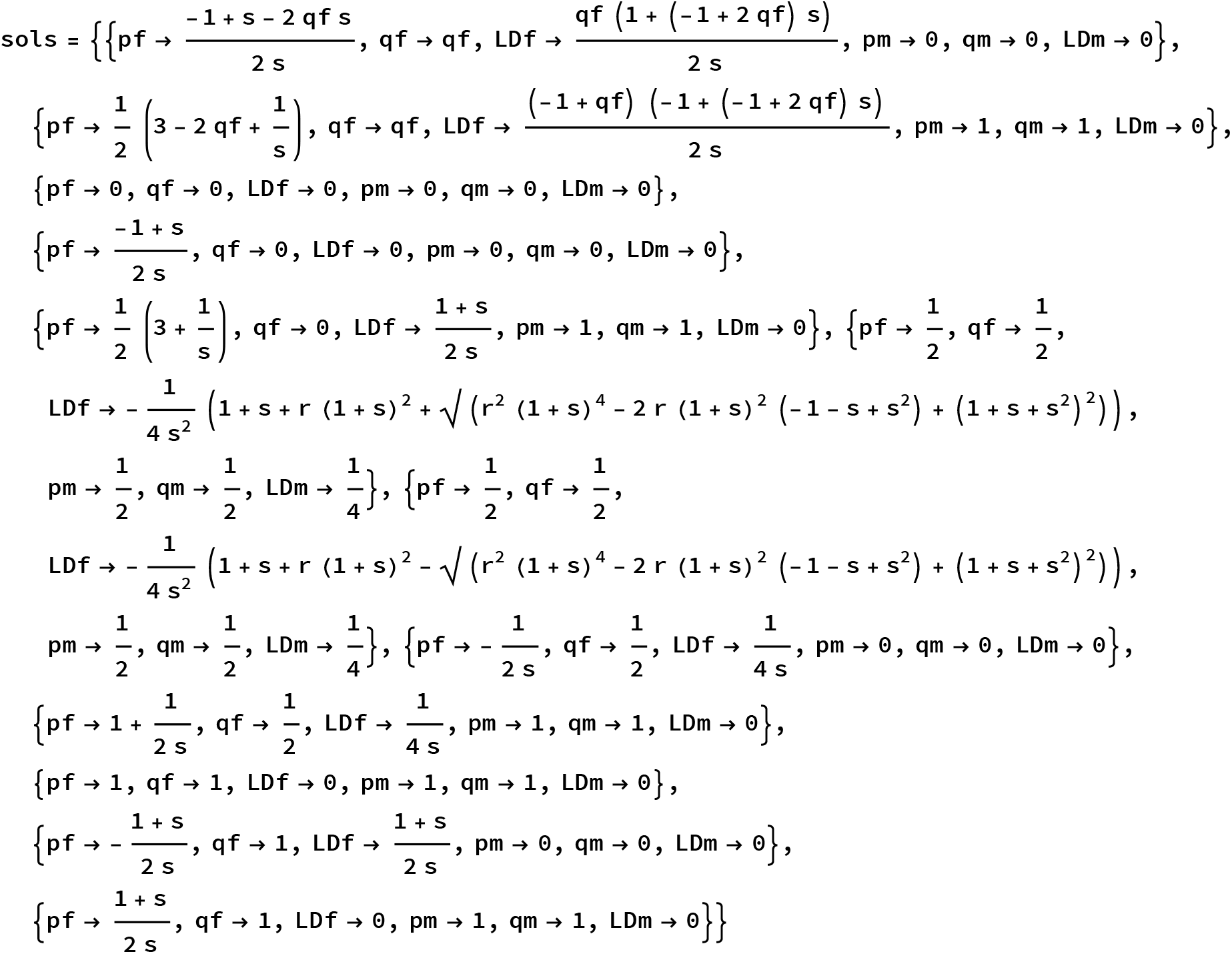
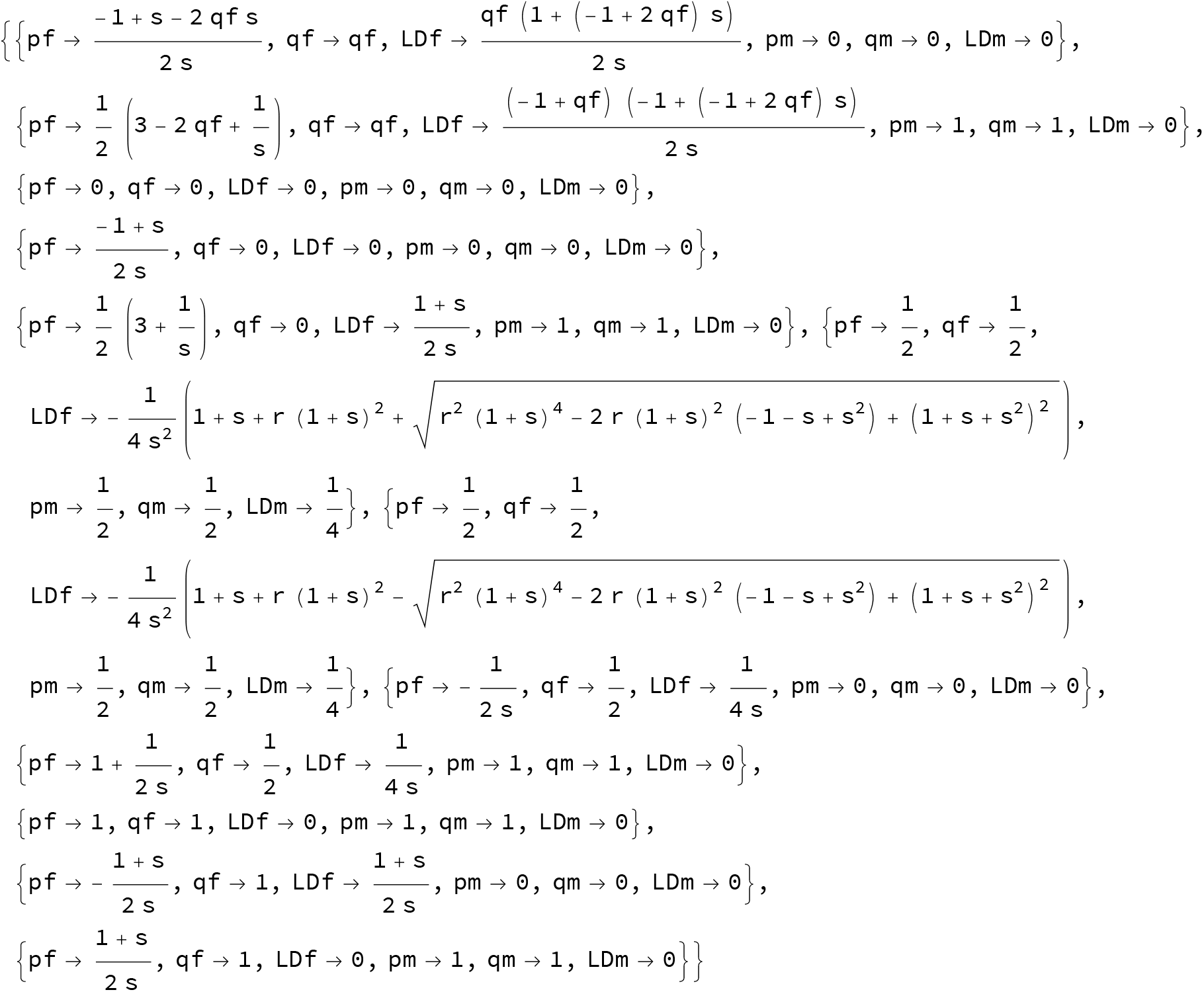

Test existence and stability (based on eigenvalues) for all solutions: updated

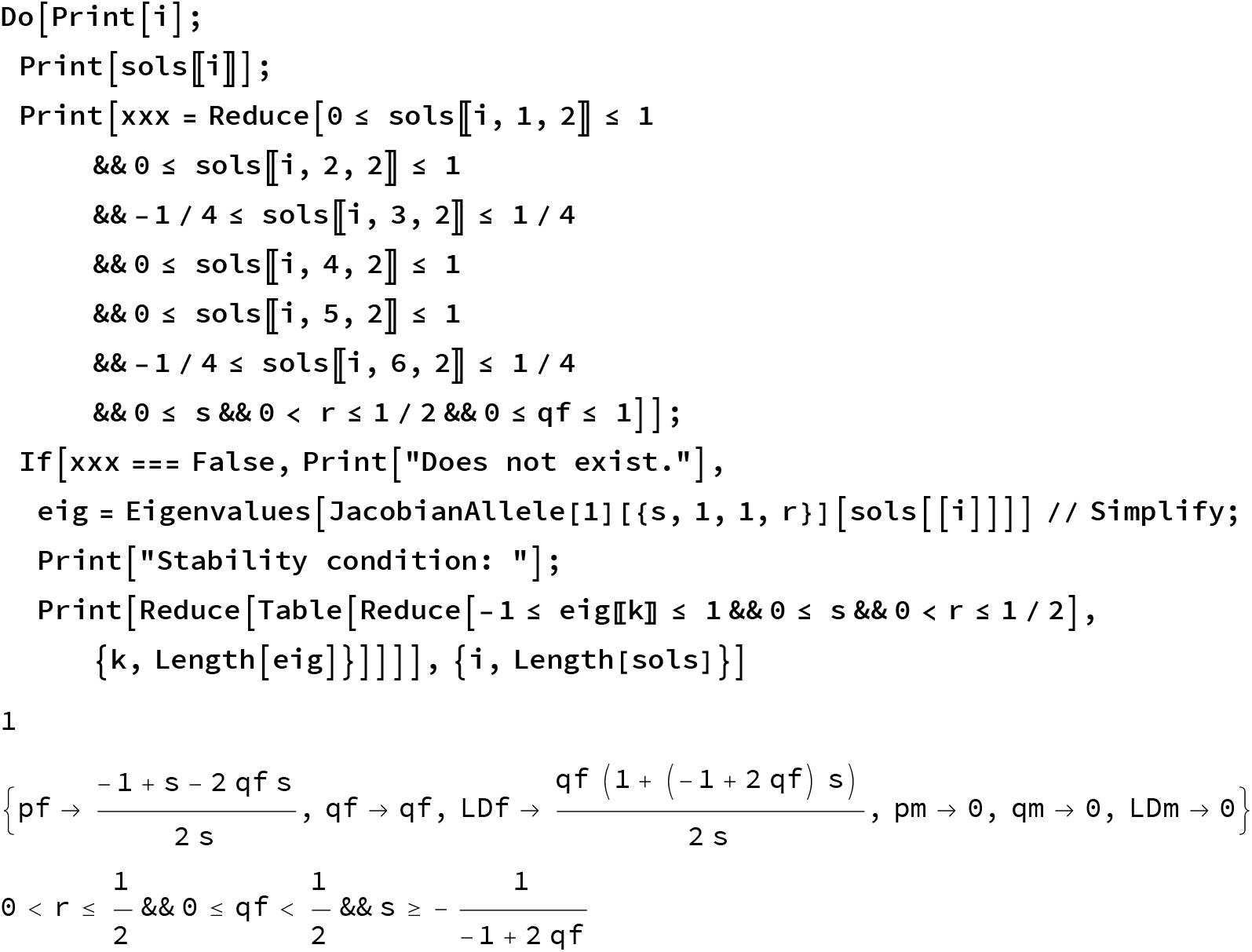
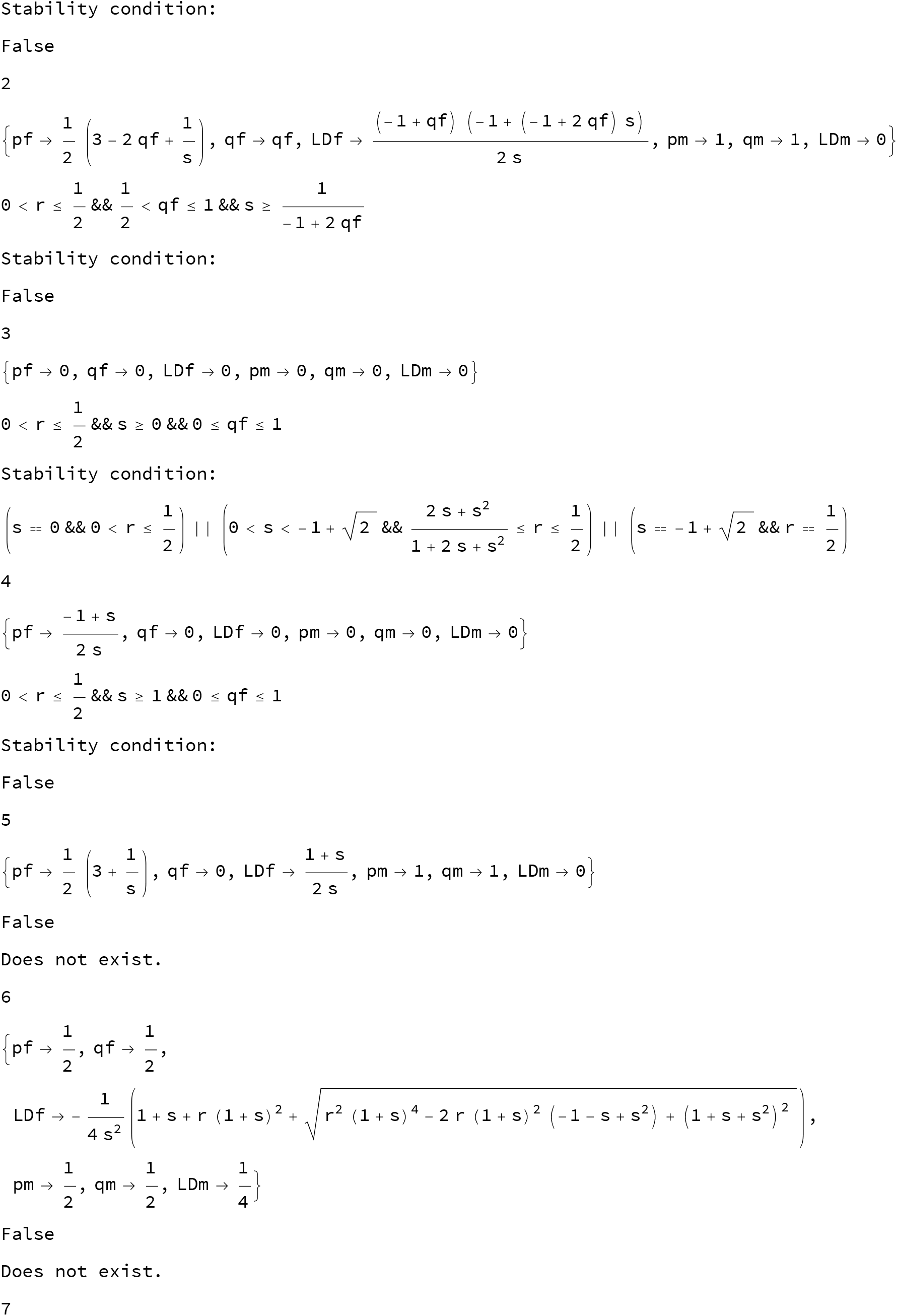
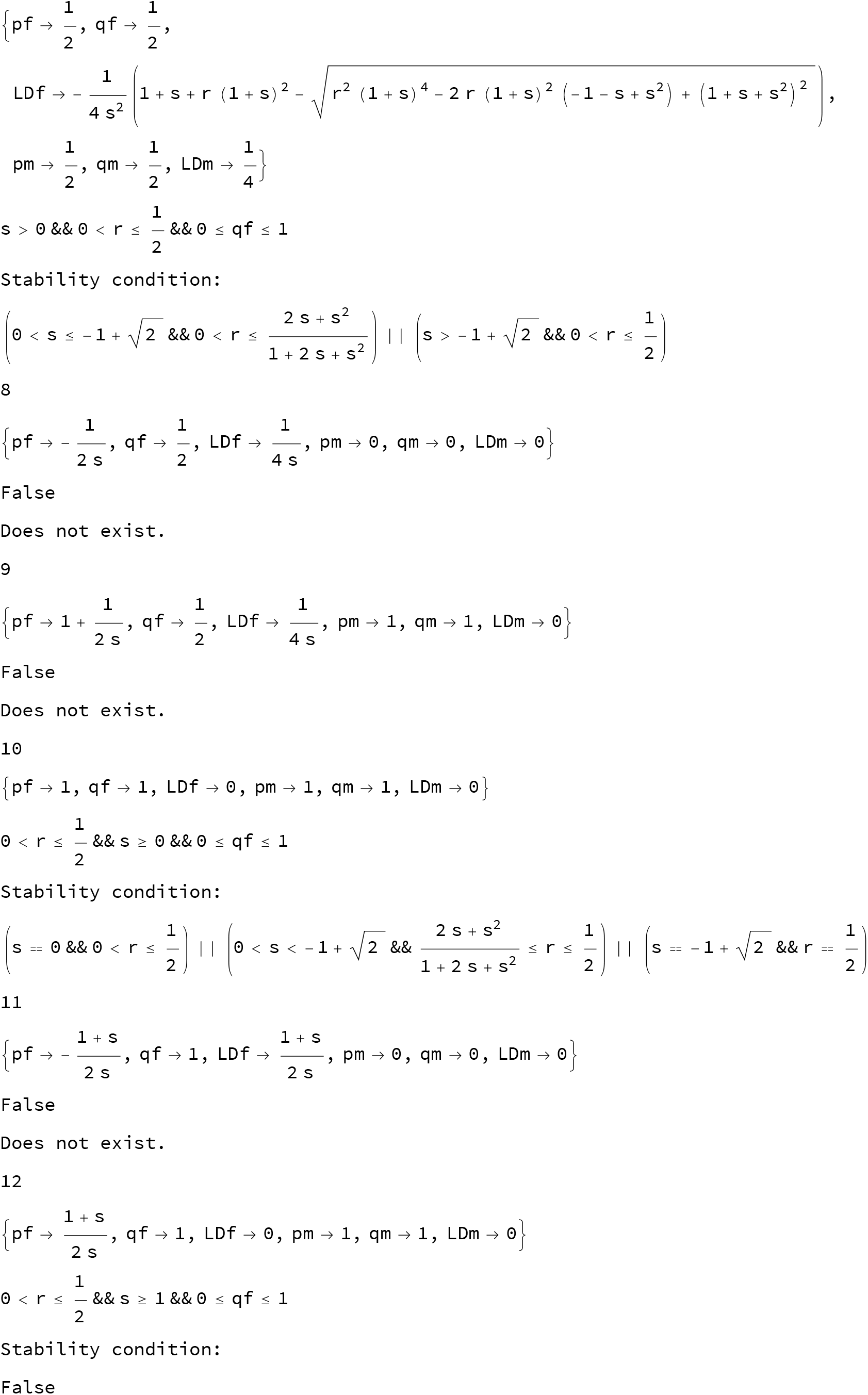

##### This is a plot of the solution for *γ*=1

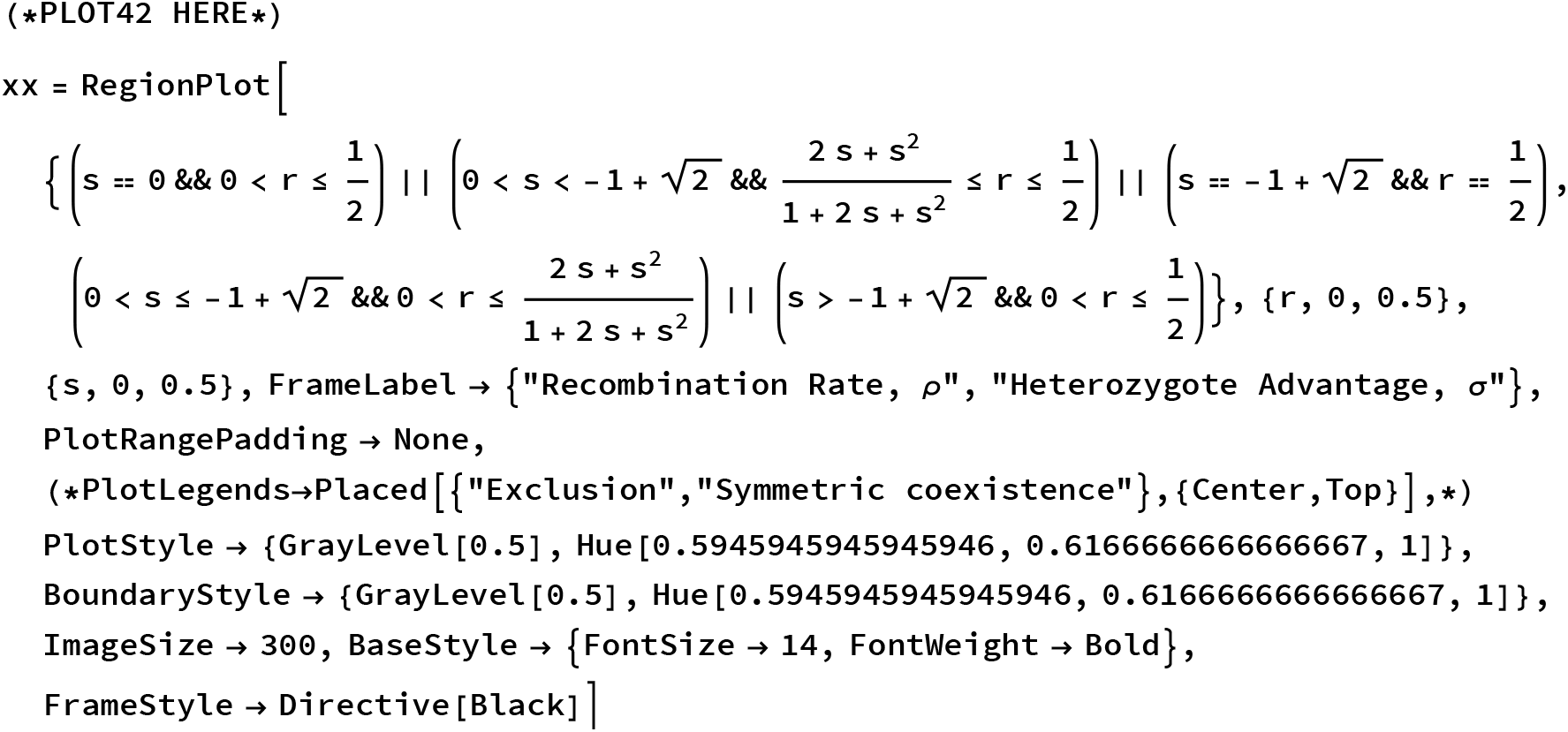

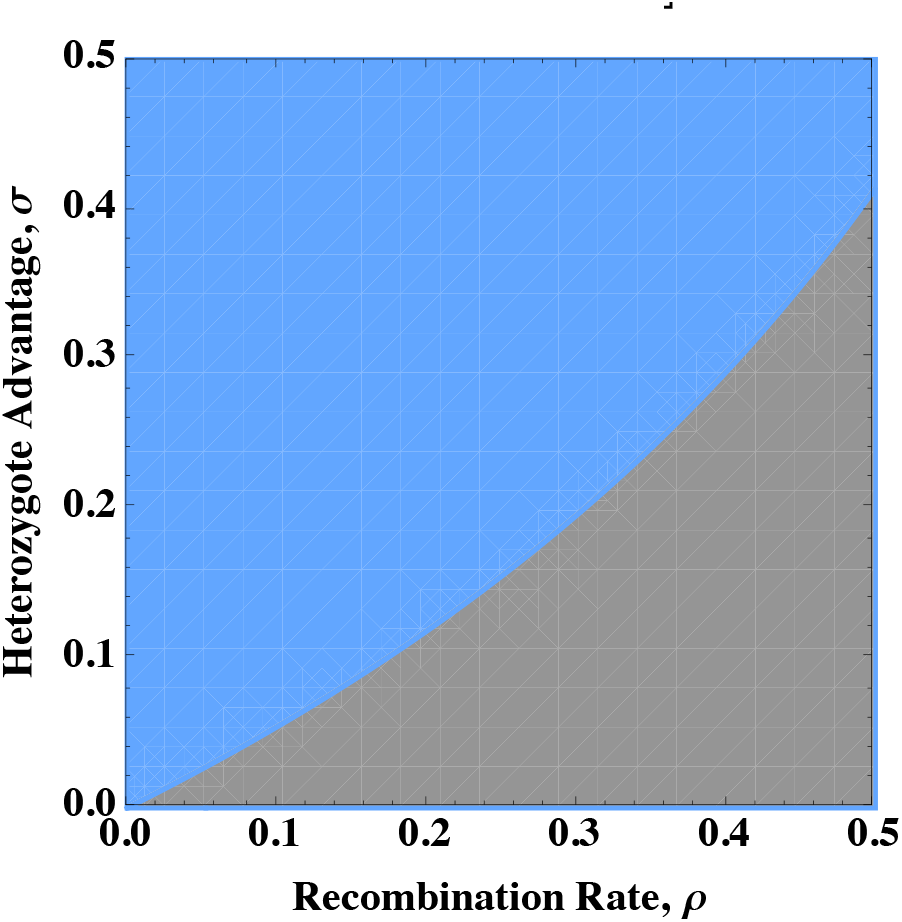

~~~
Export[ToString[NotebookDirectory[]] <> "analytic-limit-lethal-haplodiploid.pdf", xx]
~~~

#### Stability of monomorphic equilibria

This is the condition for local stability of exclusion. updated

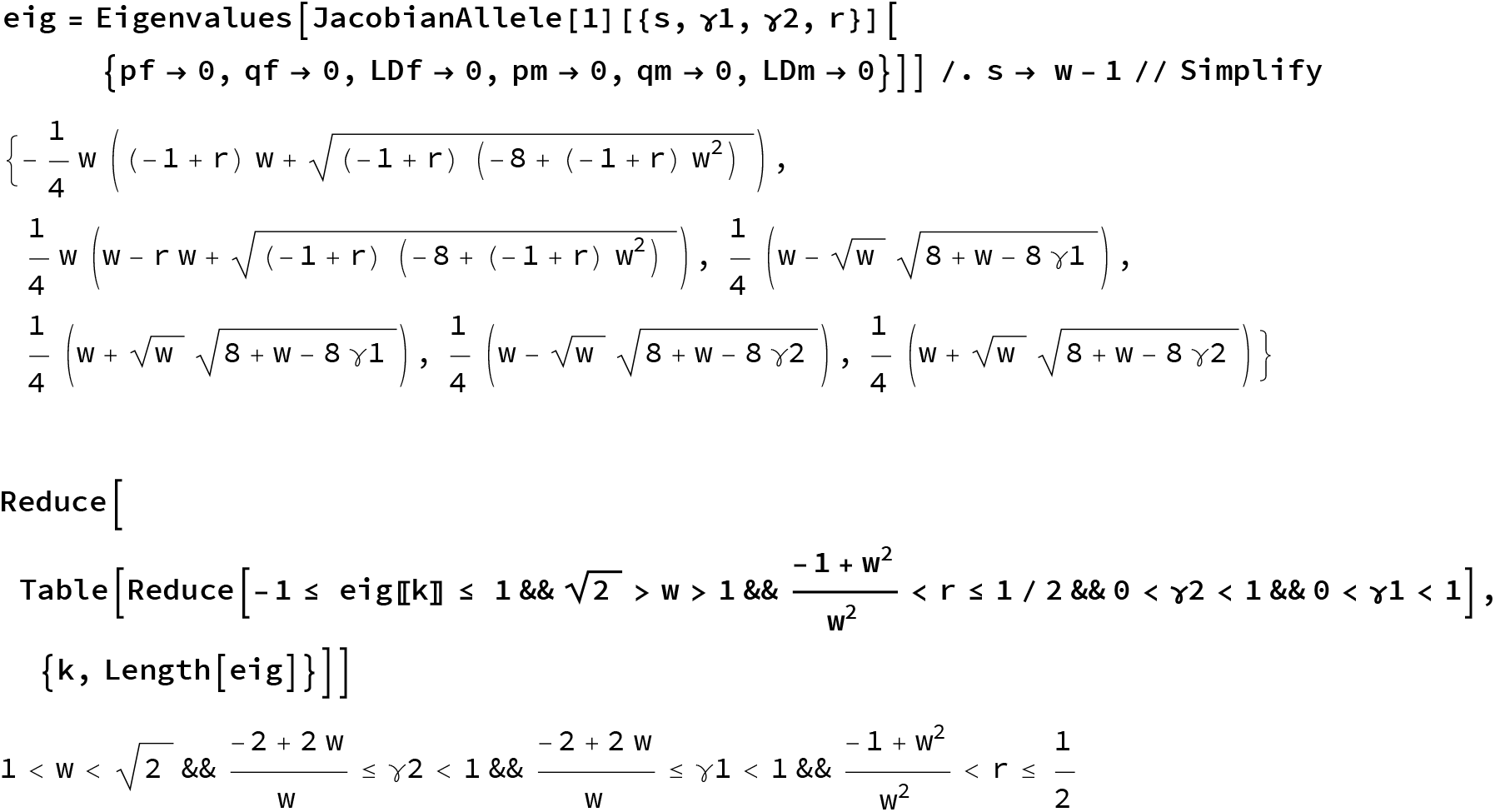

#### Plot solution

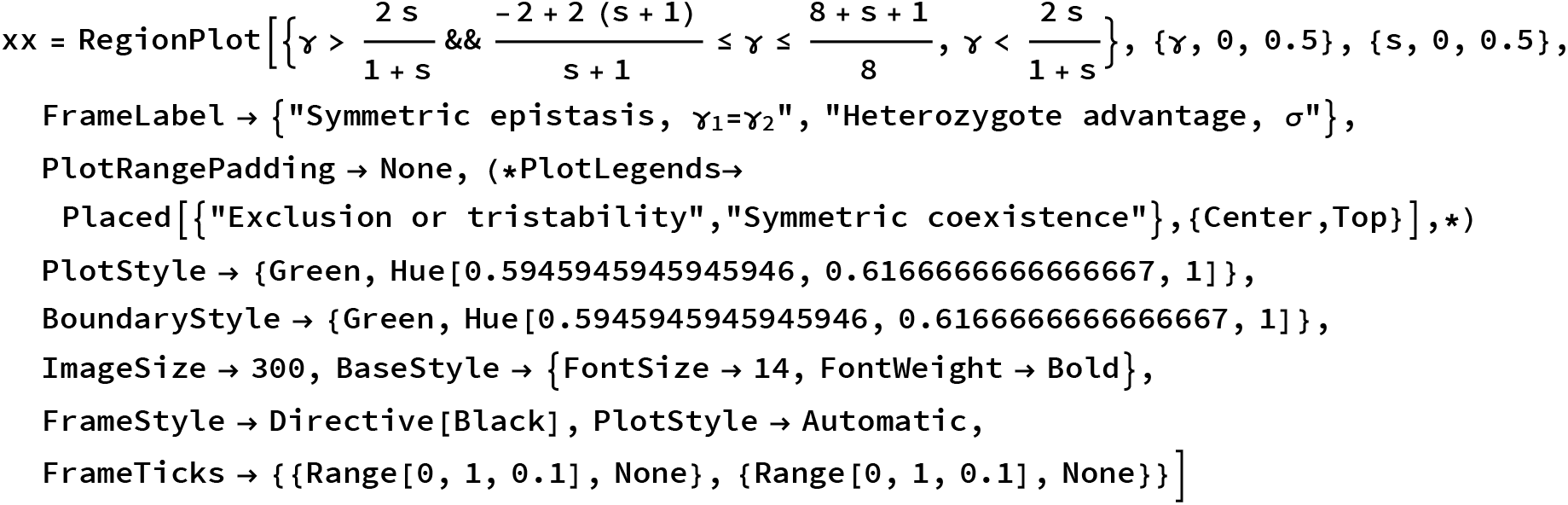
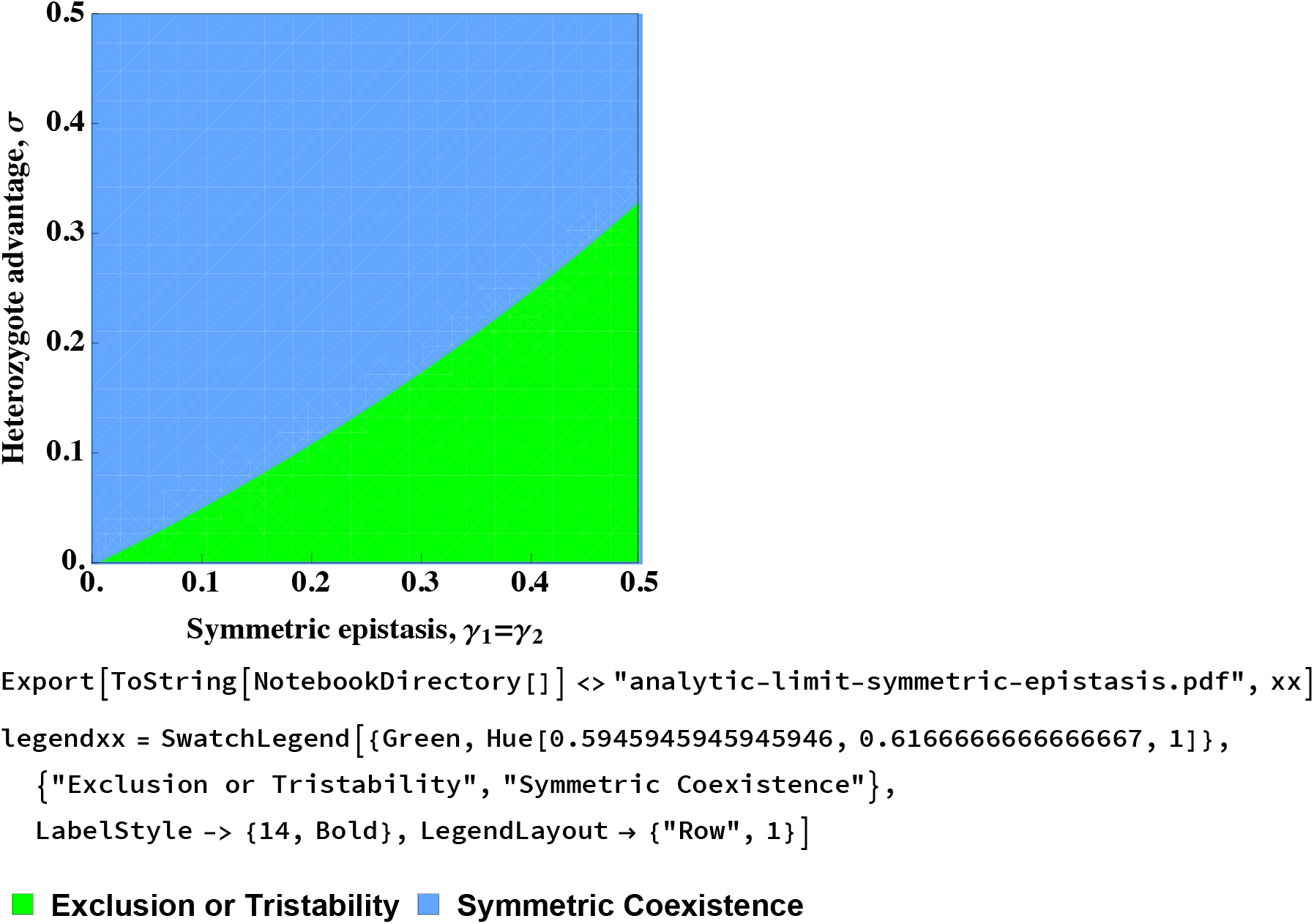

#### Equilibria and stability for γ=1 in diploids

Write out dynamics

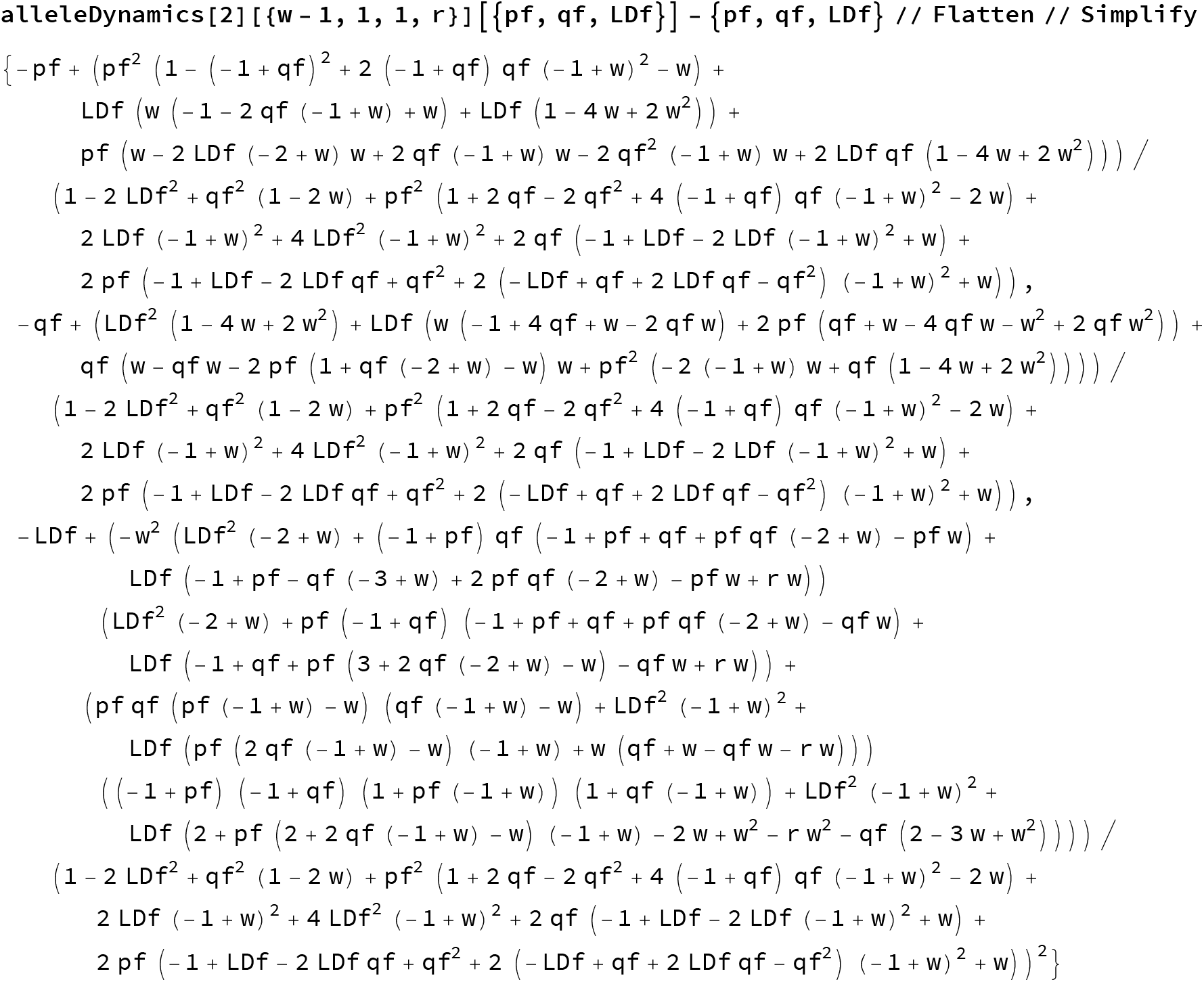

Try to find solutions:

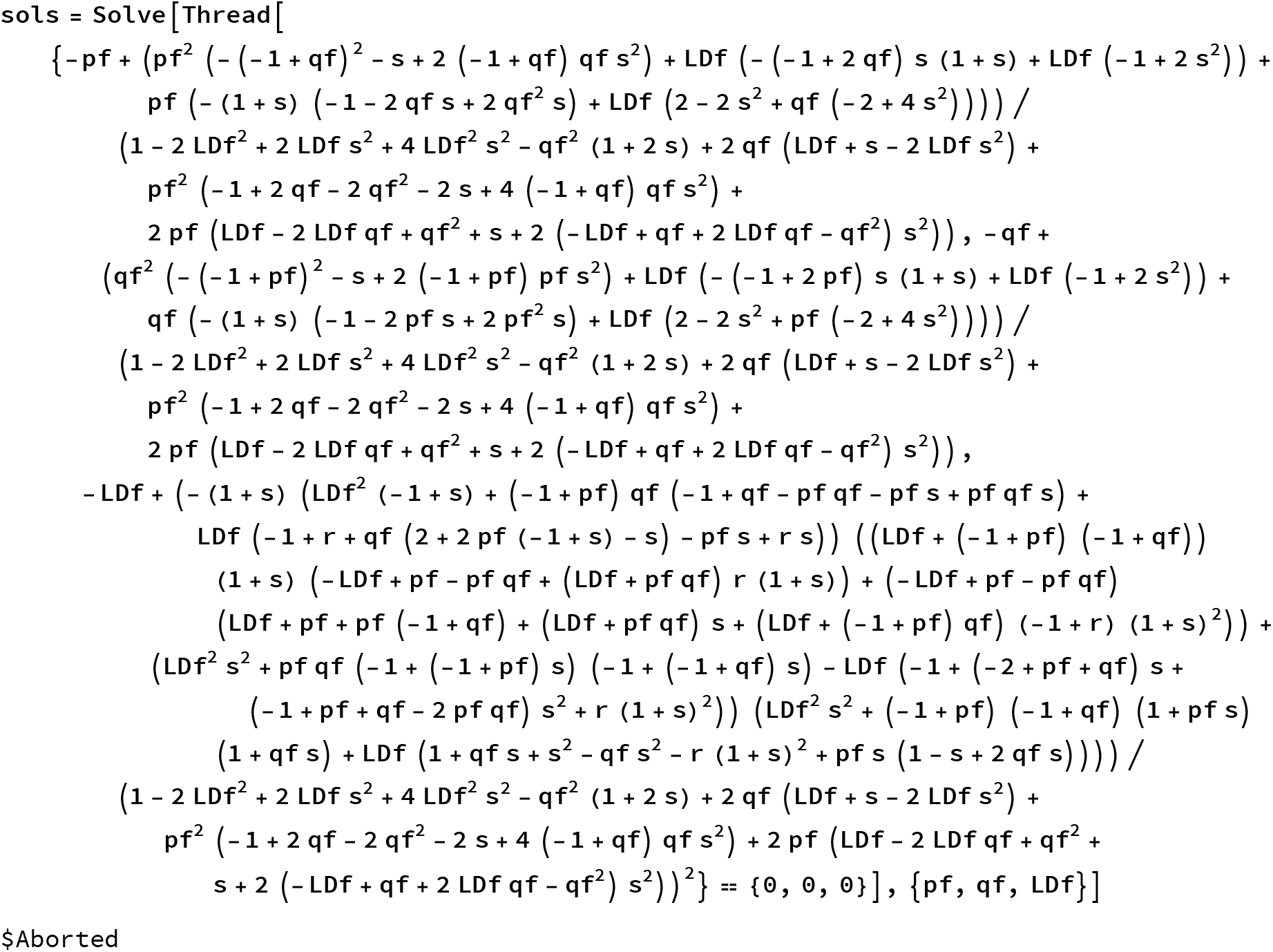

This is not solvable with Mathematica’s methods.

#### Try analysis of the internal equilibrium with *p_f_* = *q_f_*

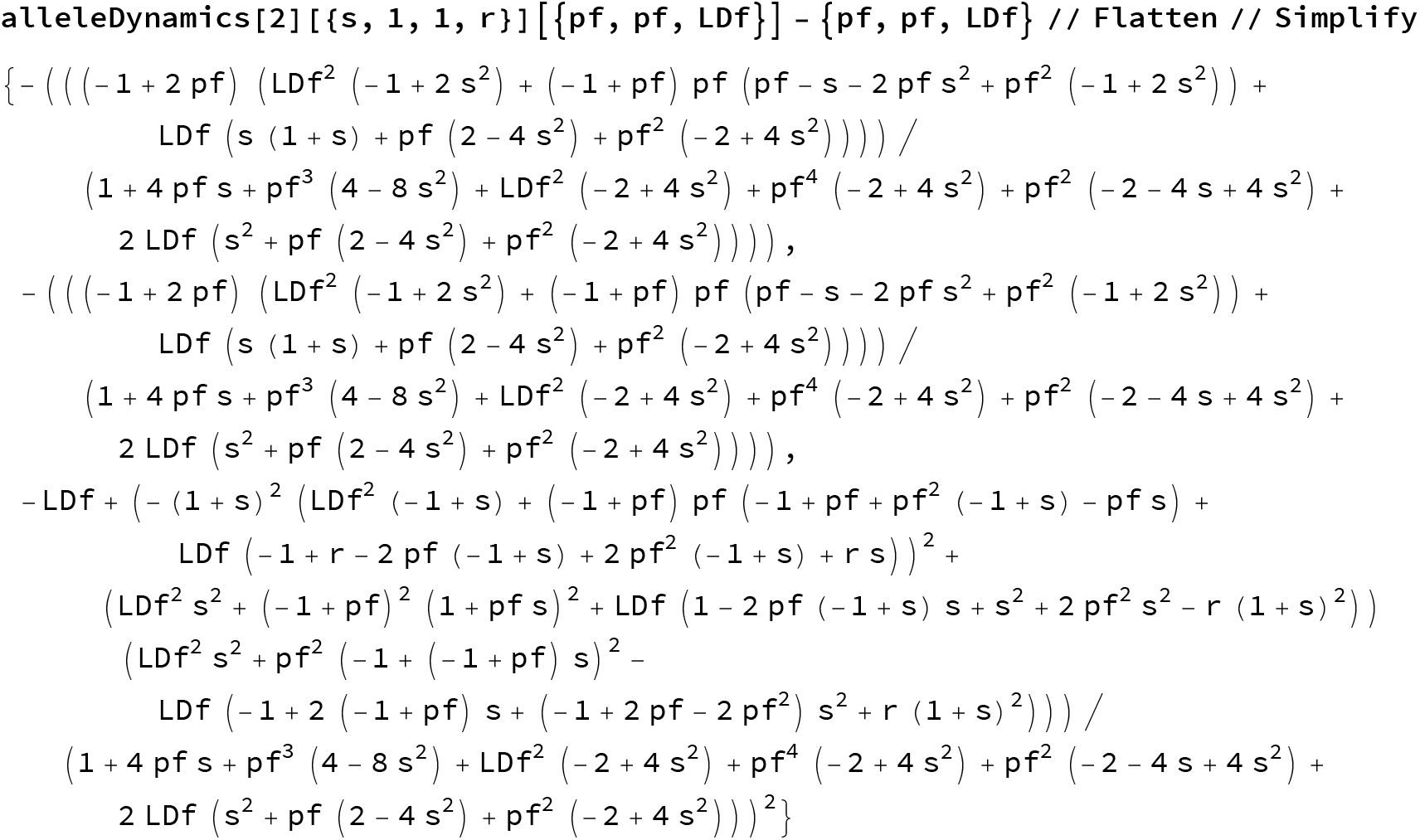
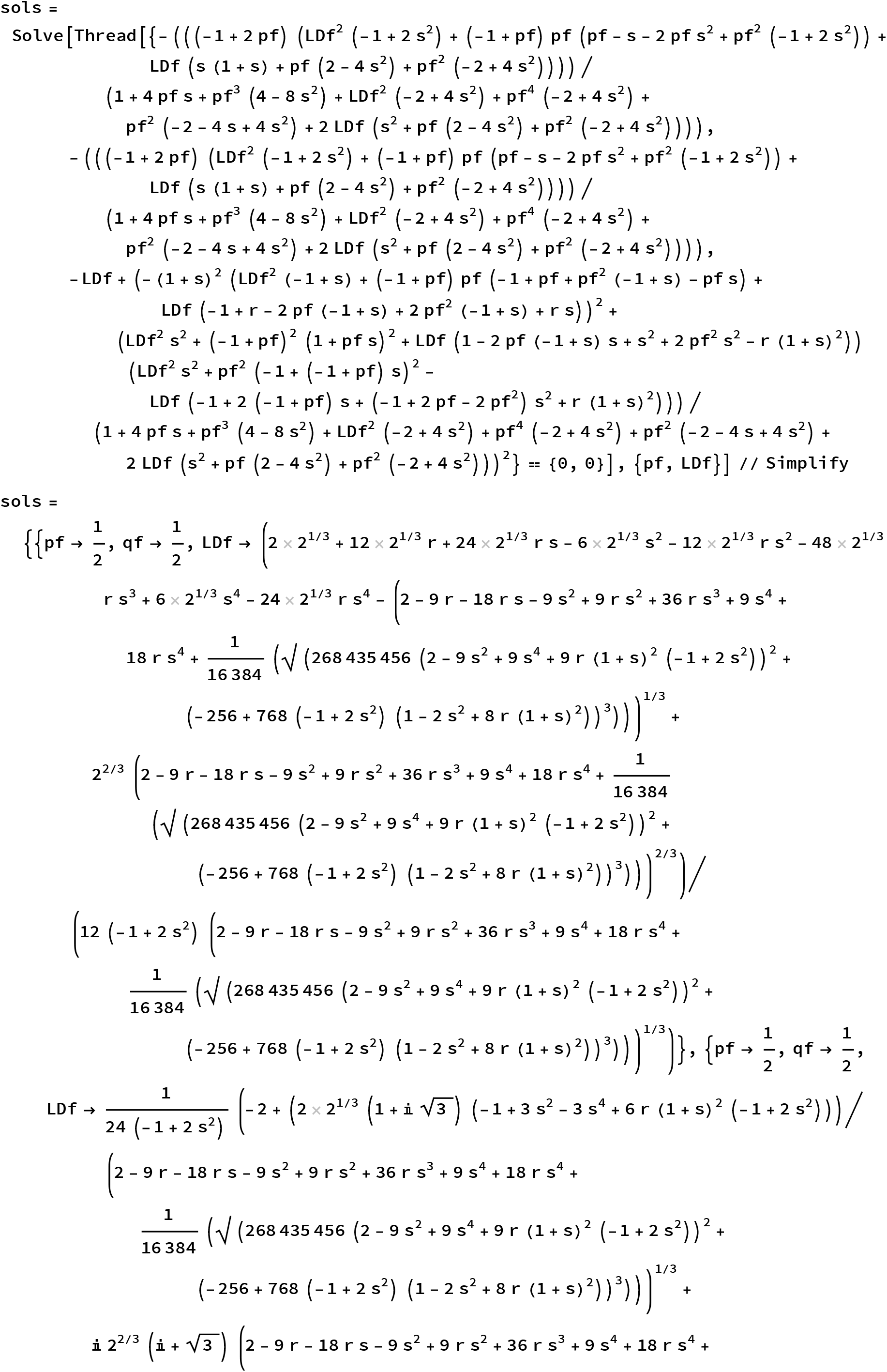
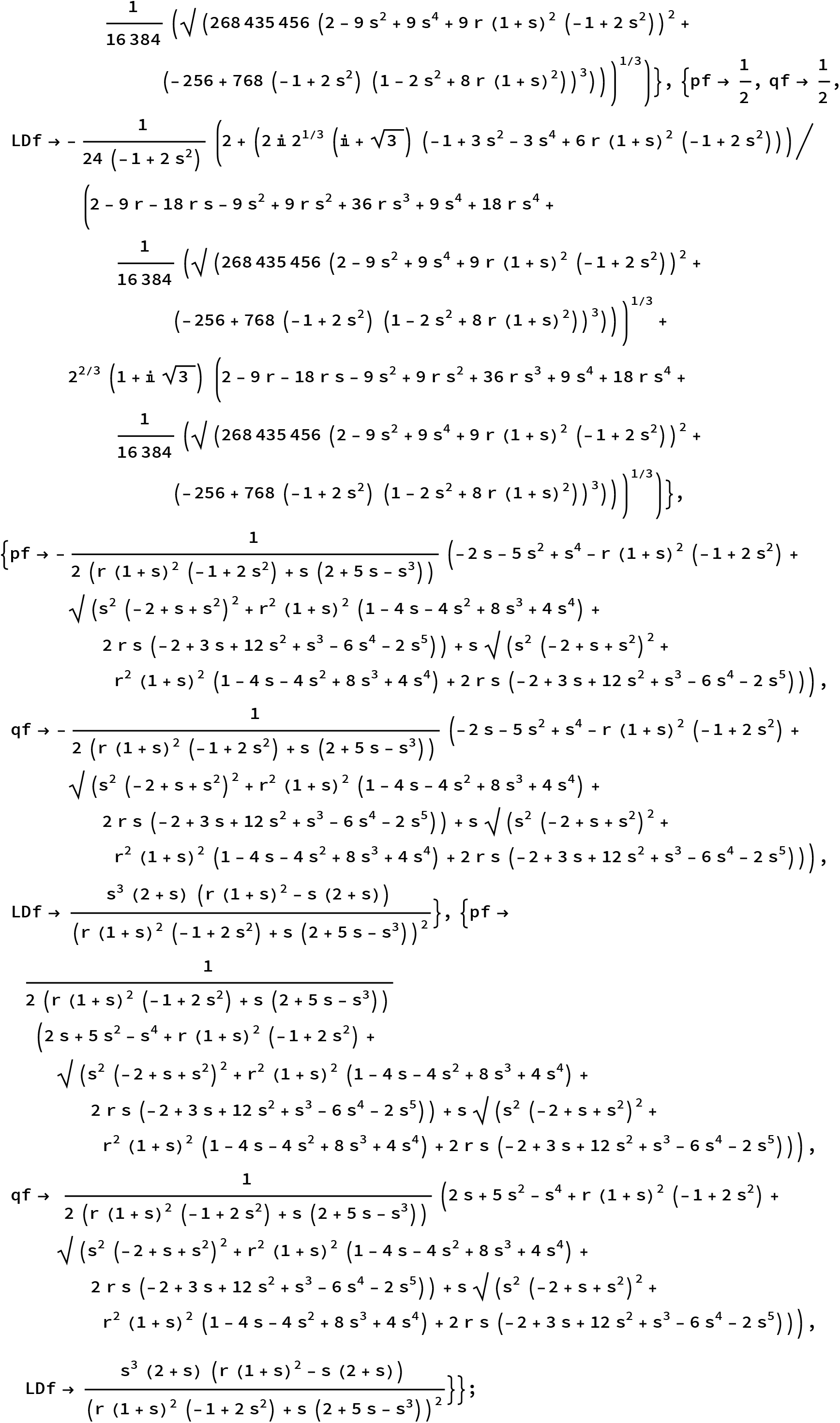

#### Check whether the last two yield the coordinates of the asymmetric-coexistence eq

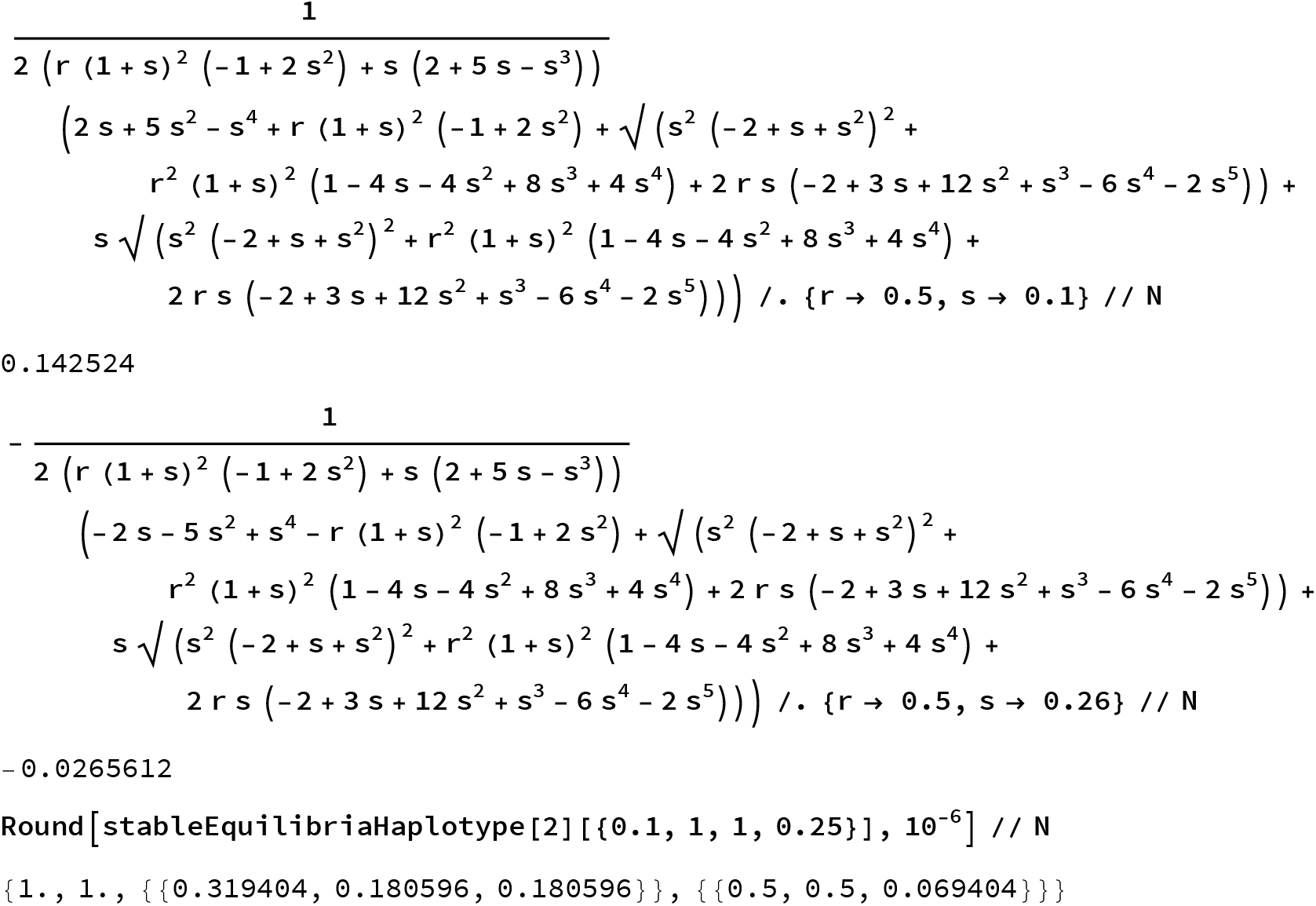

Yes, they do.

#### Try to do stability analysis - unsuccessful

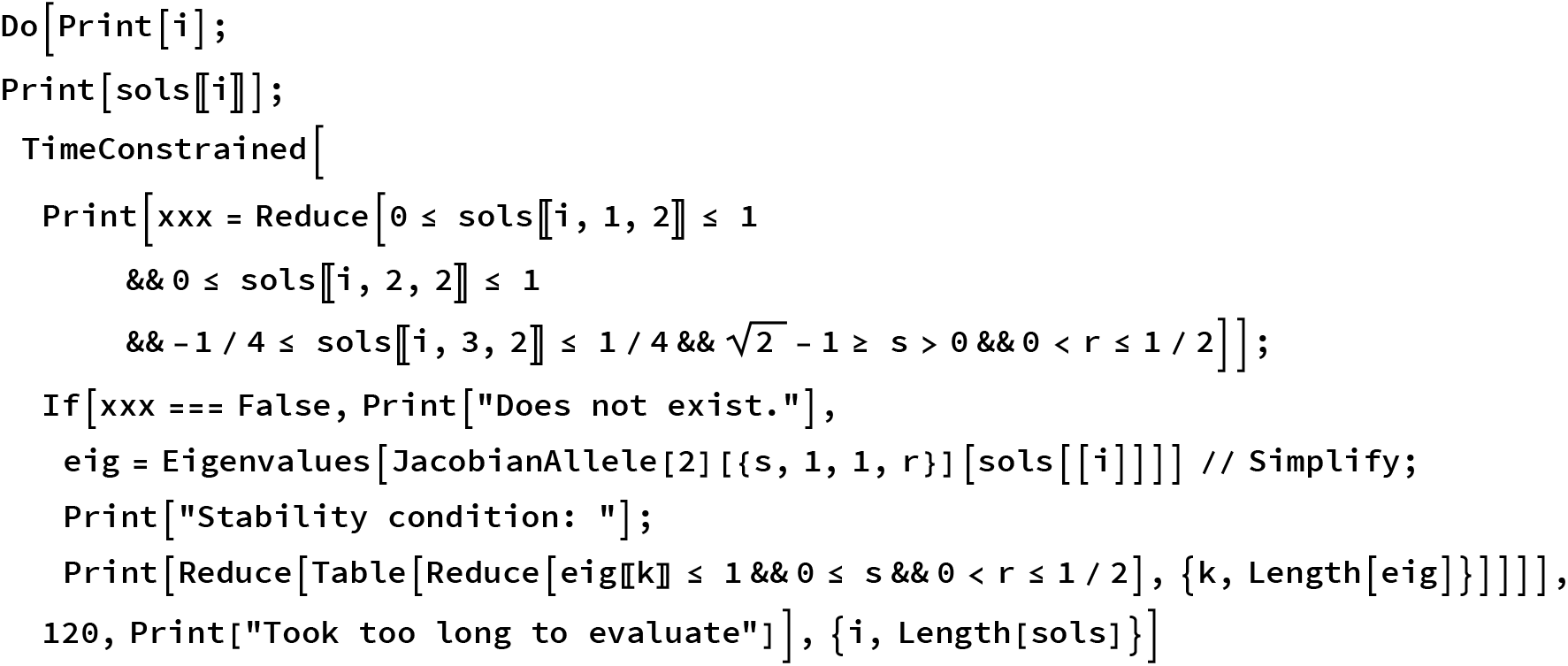

Check when the coordinates of the equilibrium pass 0 & 1; this could be a potential criterion.

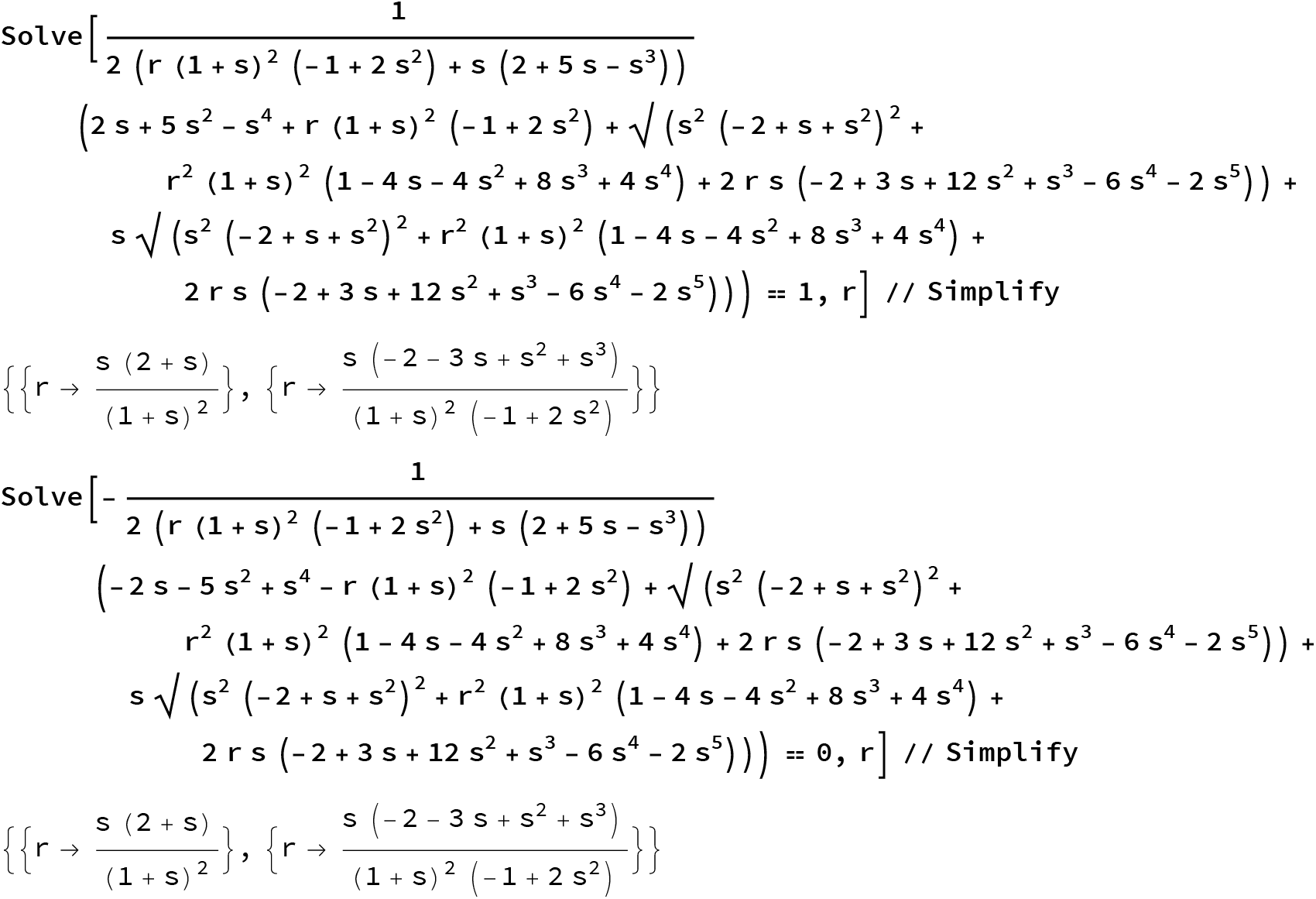

These are the criteria for existence of the “asymmetric coexistence” equilibrium.

#### Guess limit of stability from existence conditions above

Plot possible conditions and see which ones make most sense.

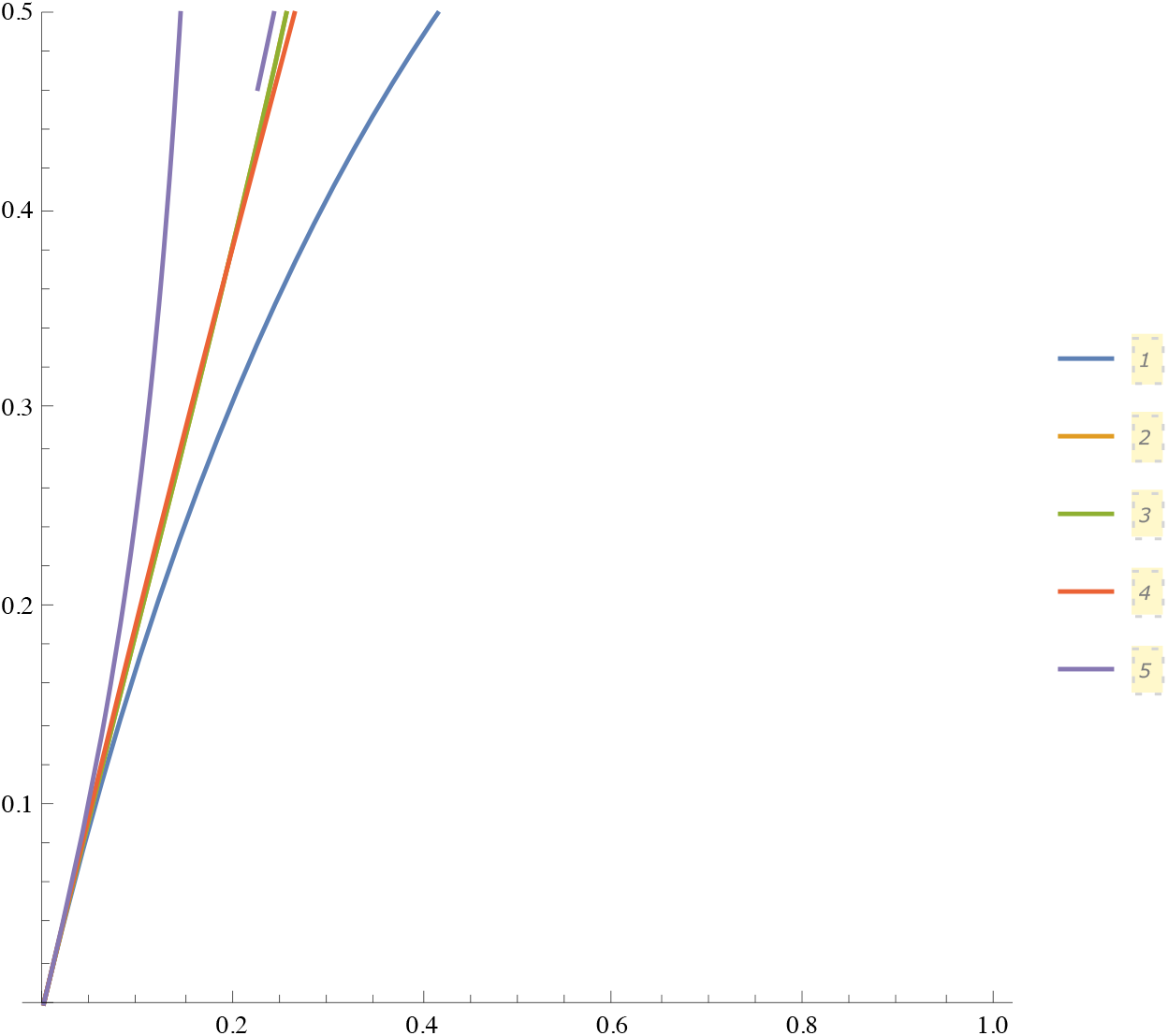
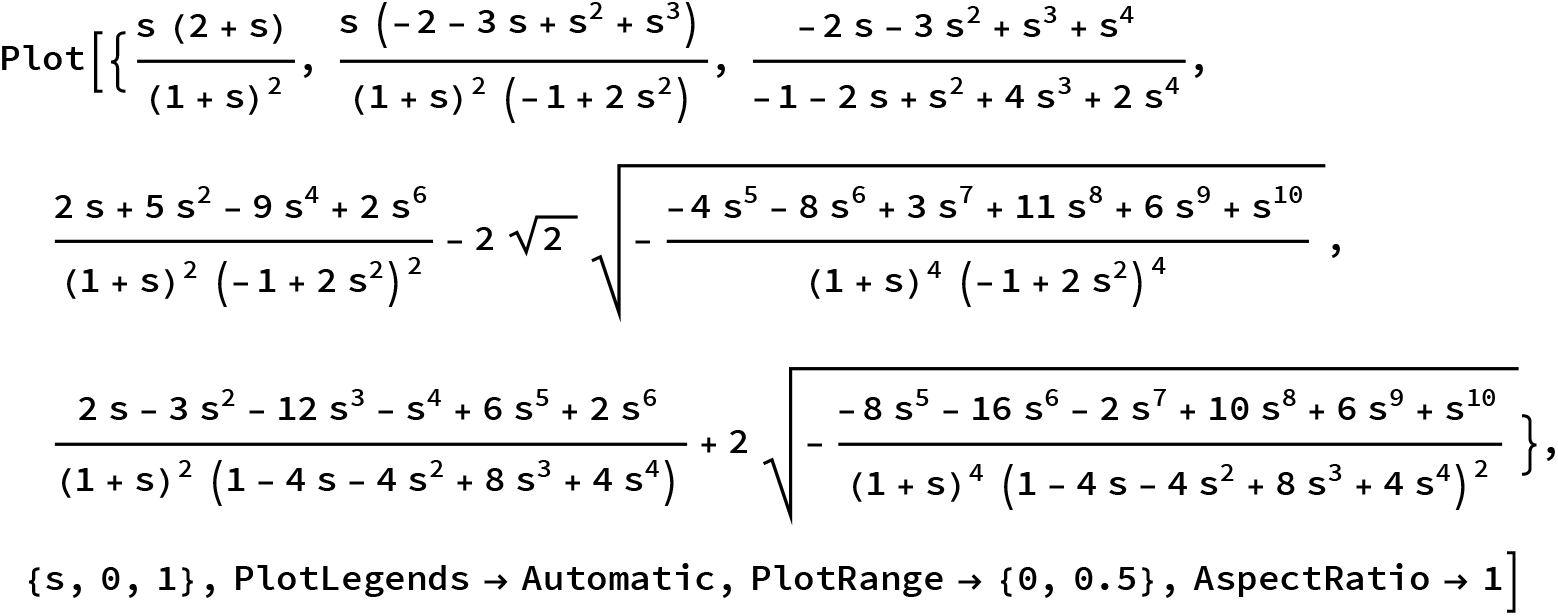

#### Plot solution

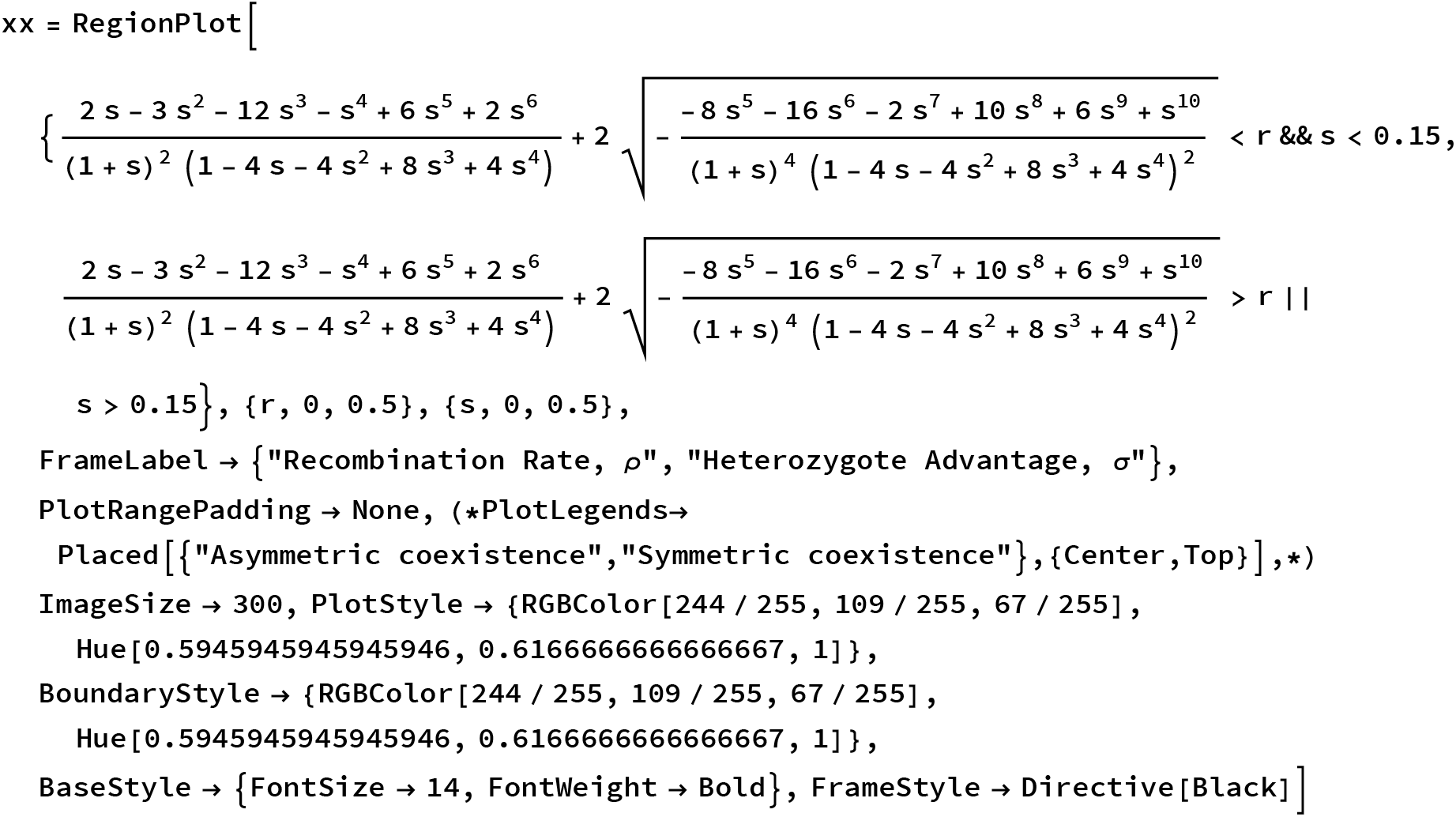
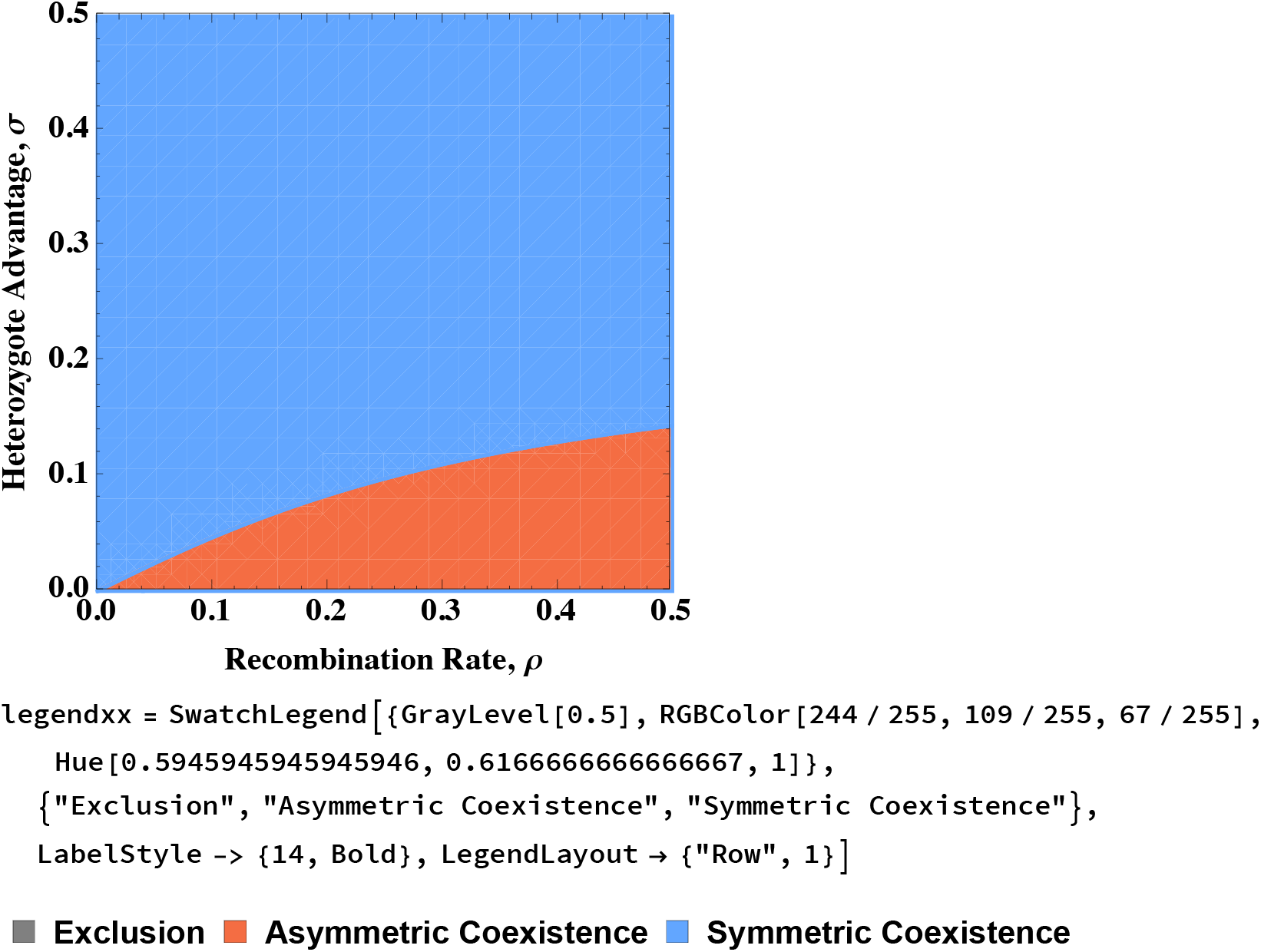

~~~
Export[ToString[NotebookDirectory[]] <> "analytic-limit-lethal-diploid.pdf", xx]
Export[ToString[NotebookDirectory[]] <> "analytic-limit-lethal-legend.pdf", legendxx]
/Users/nando/Documents/finnish_hybrid_ants/results/analytic-limit-lethal-diploid.pdf
/Users/nando/Documents/finnish_hybrid_ants/results/analytic-limit-lethal-legend.pdf
~~~

#### Coordinates of SLPs

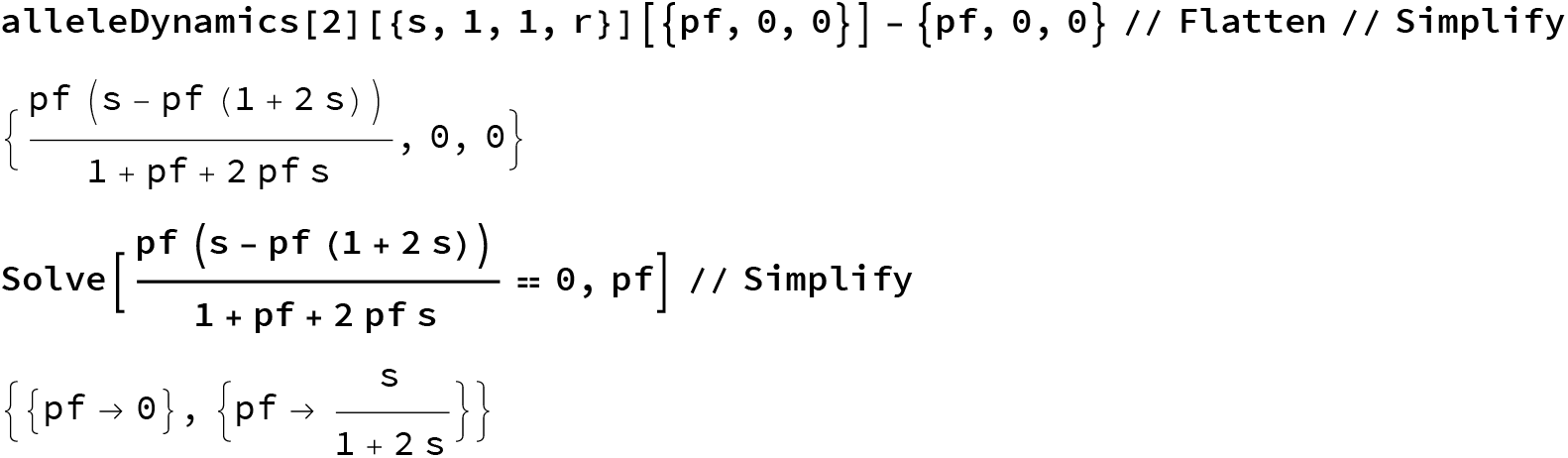

#### No stable monomorphic or SLP equilibria

These are all monomorphic & SLP equilibria: updated

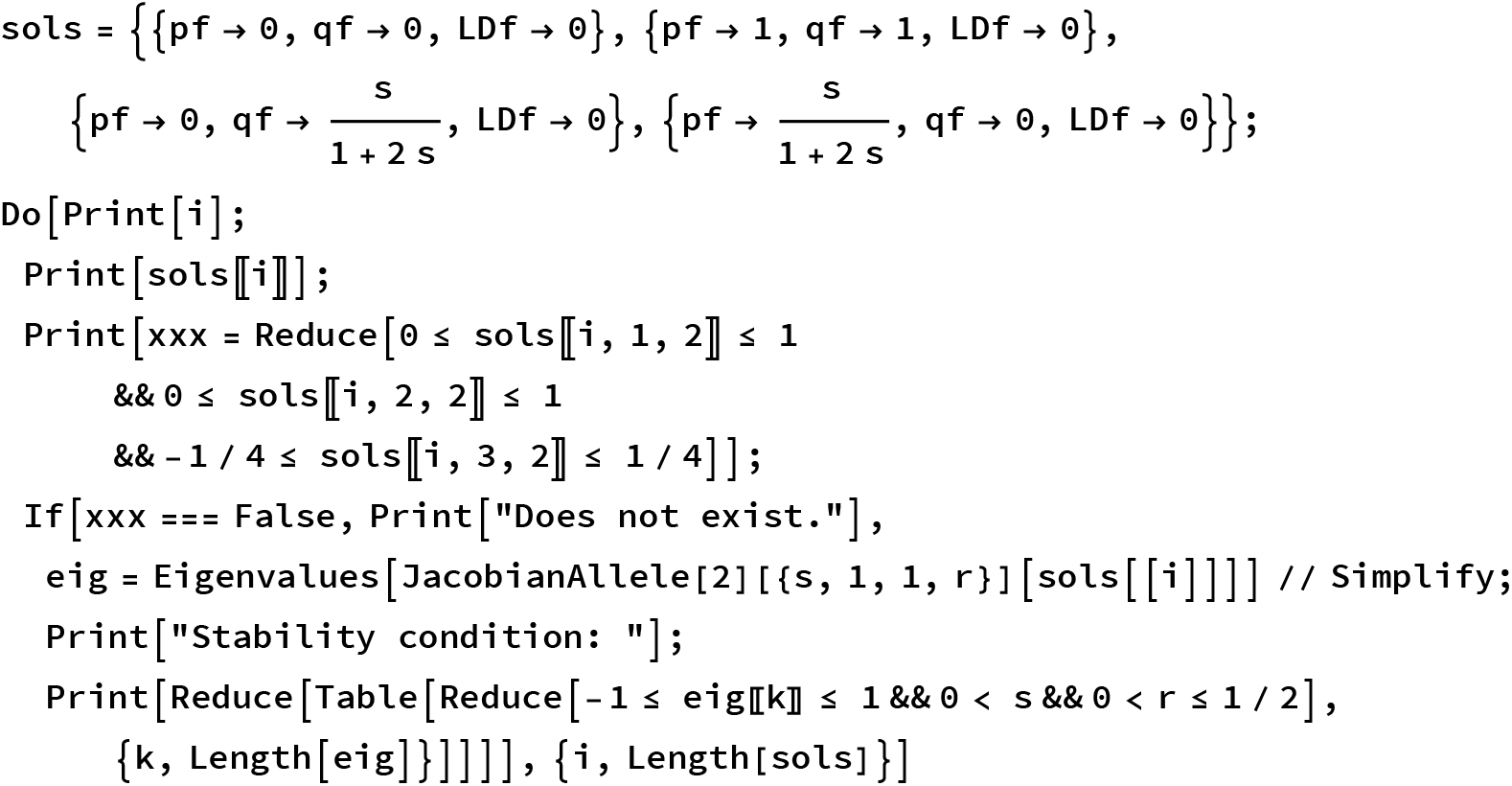
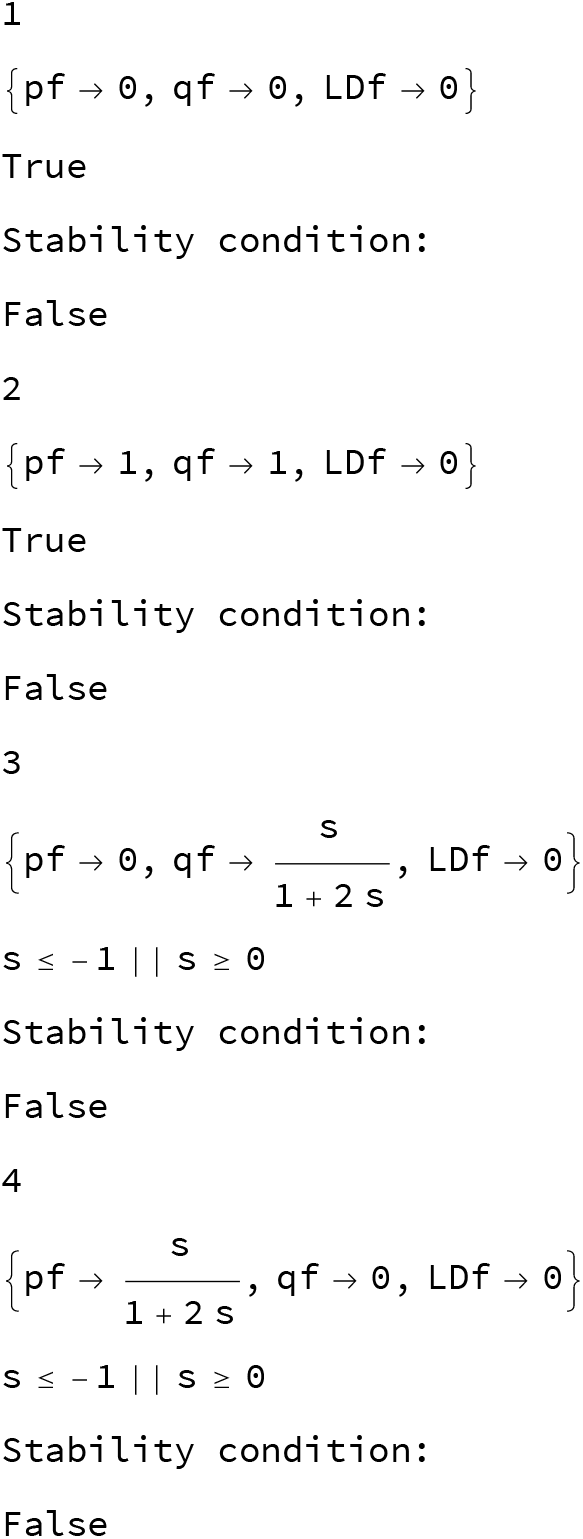

### Chapter 4: An extension to multiple loci

#### Main Functions

Function to generates gametes from diploid parents for four loci

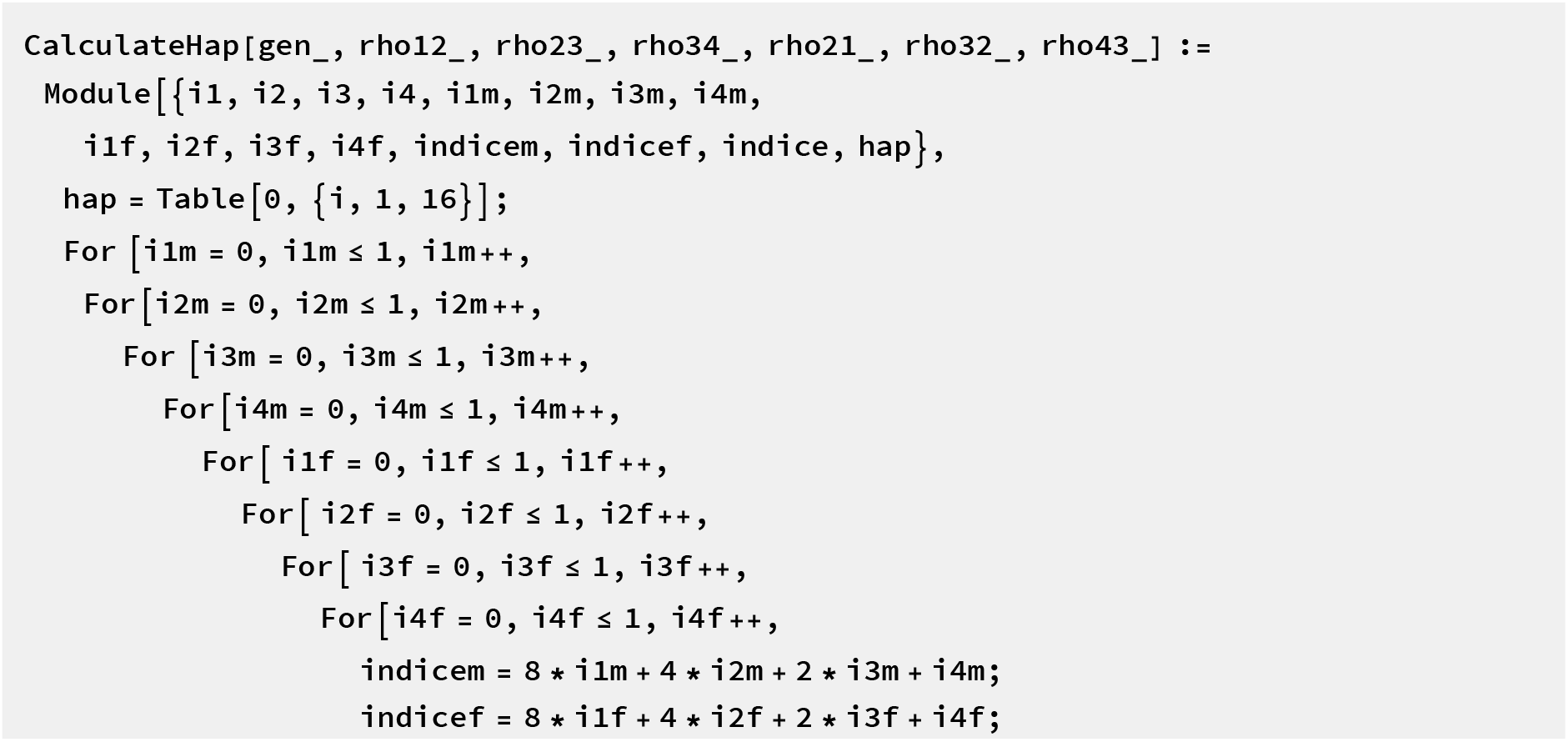
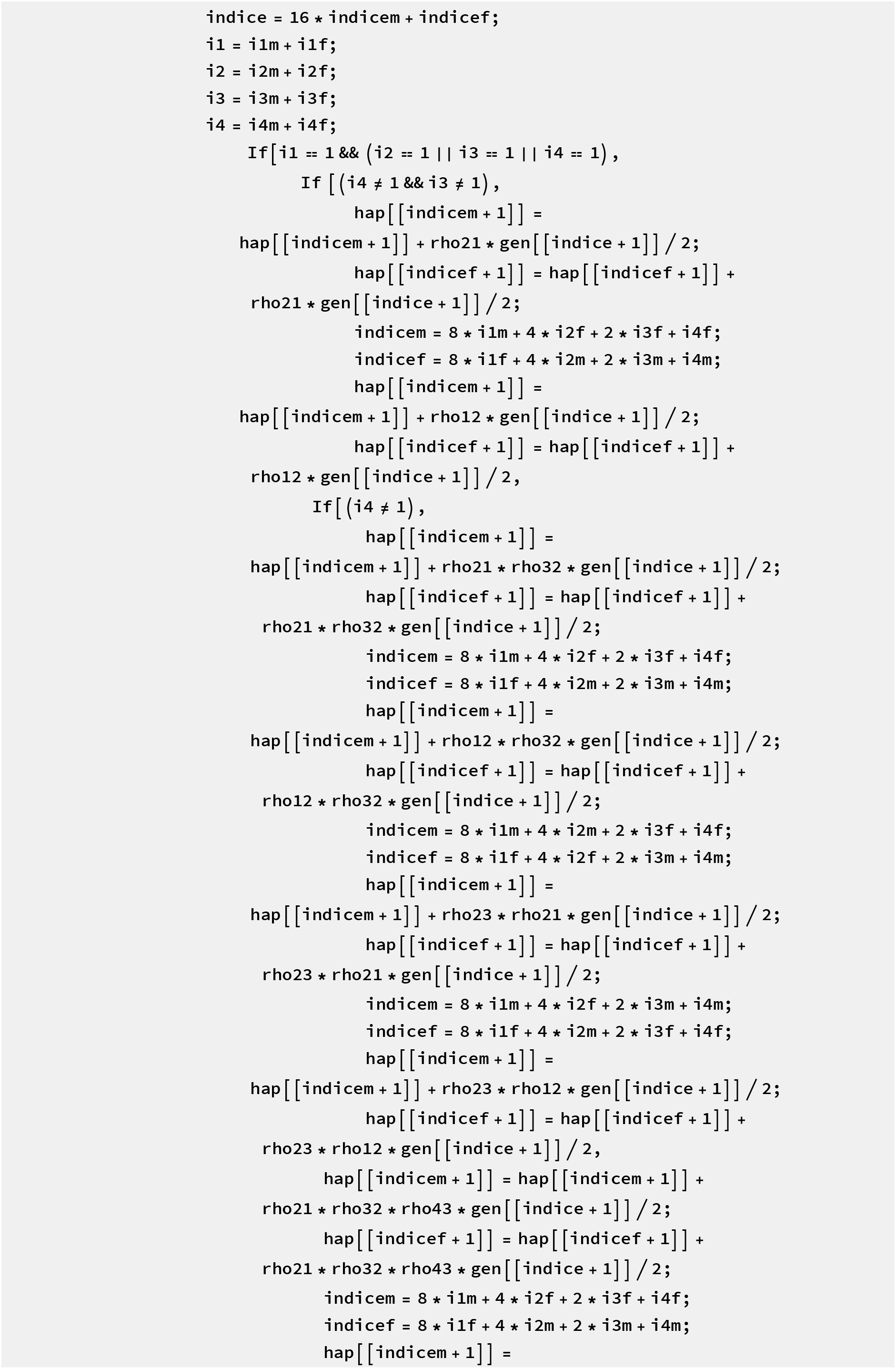
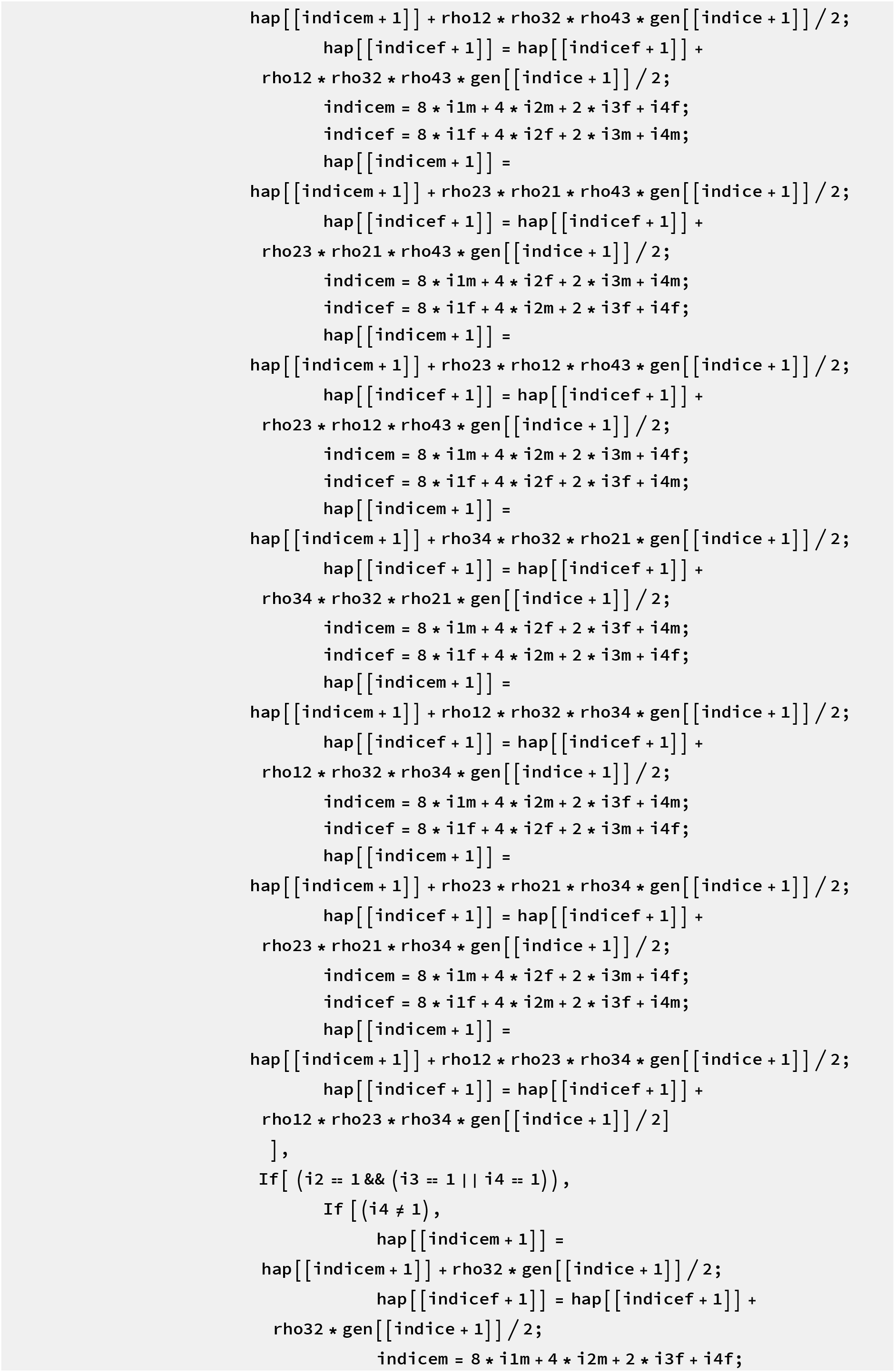
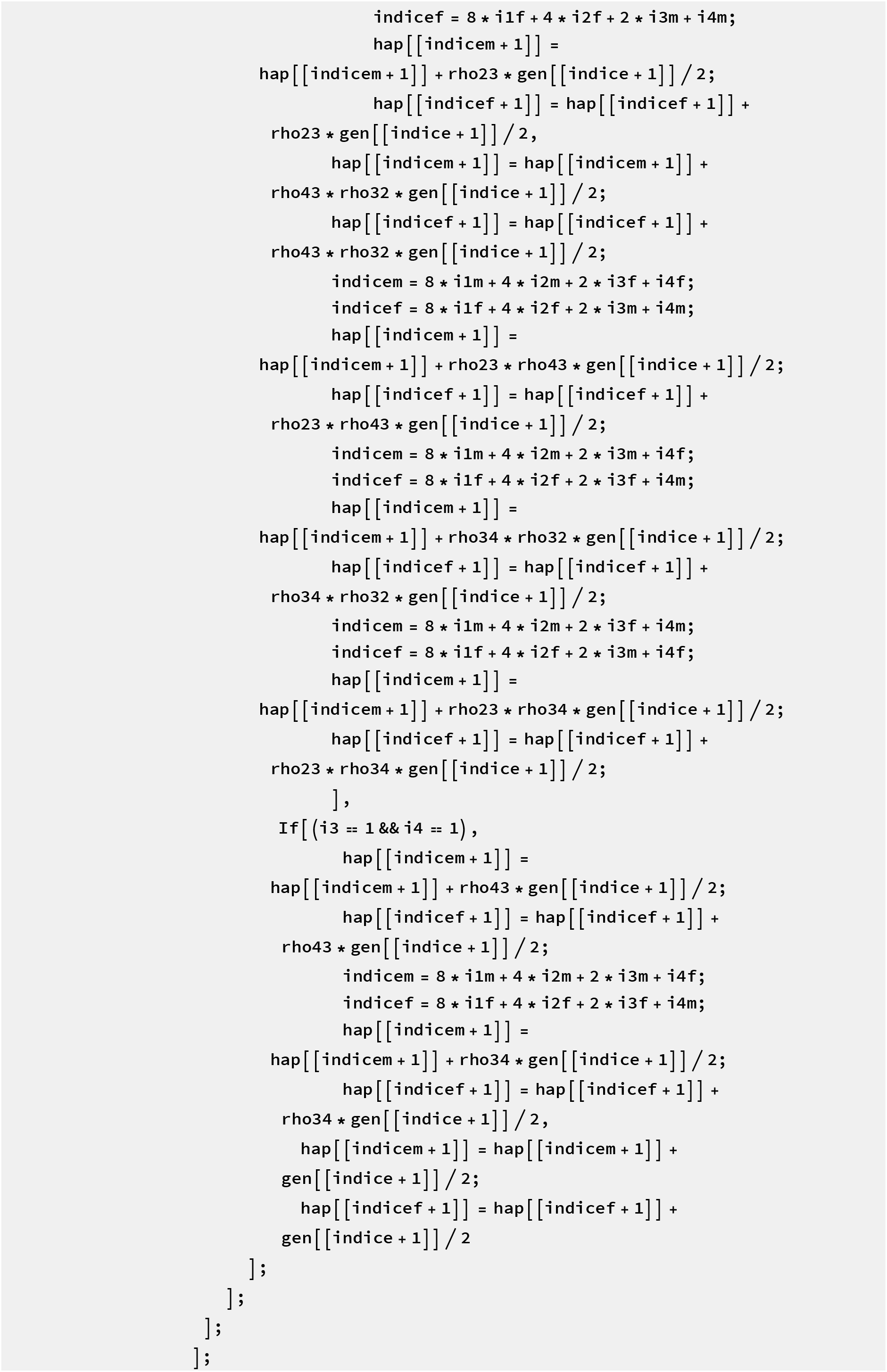
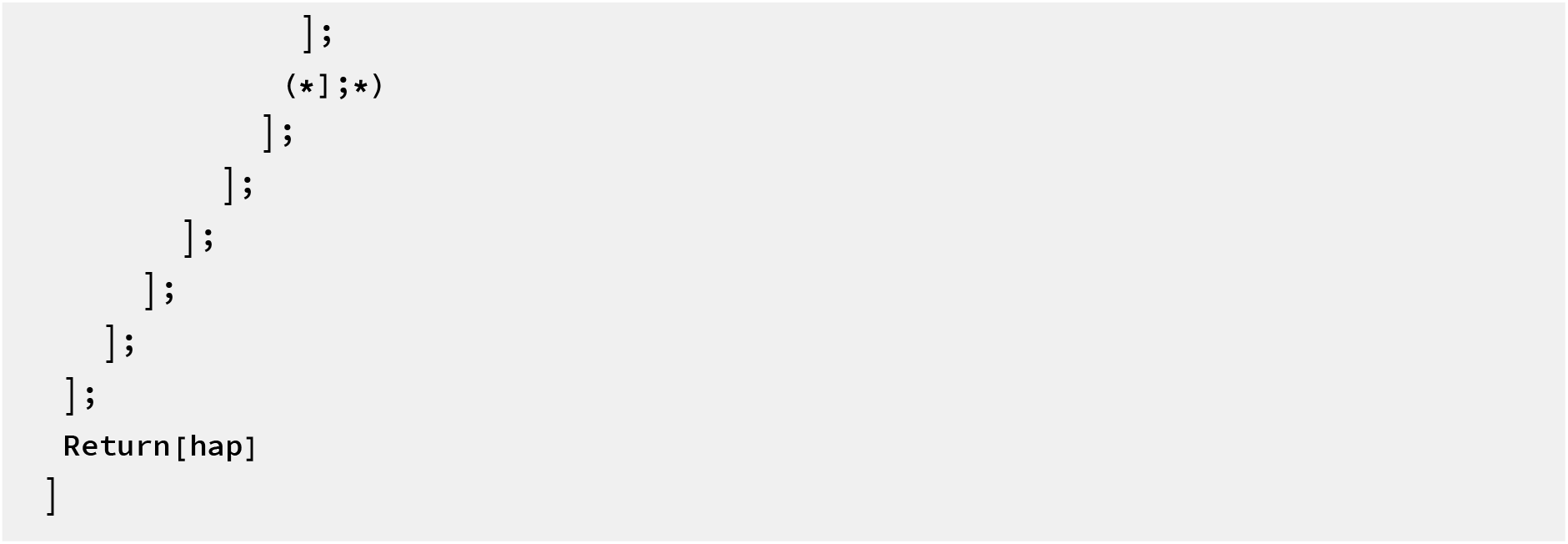

Define the dynamics of the system with fertilization, followed by selection and then recombination. Only tracks the haplotype frequncies. Takes arbitrary fitness table for males and females

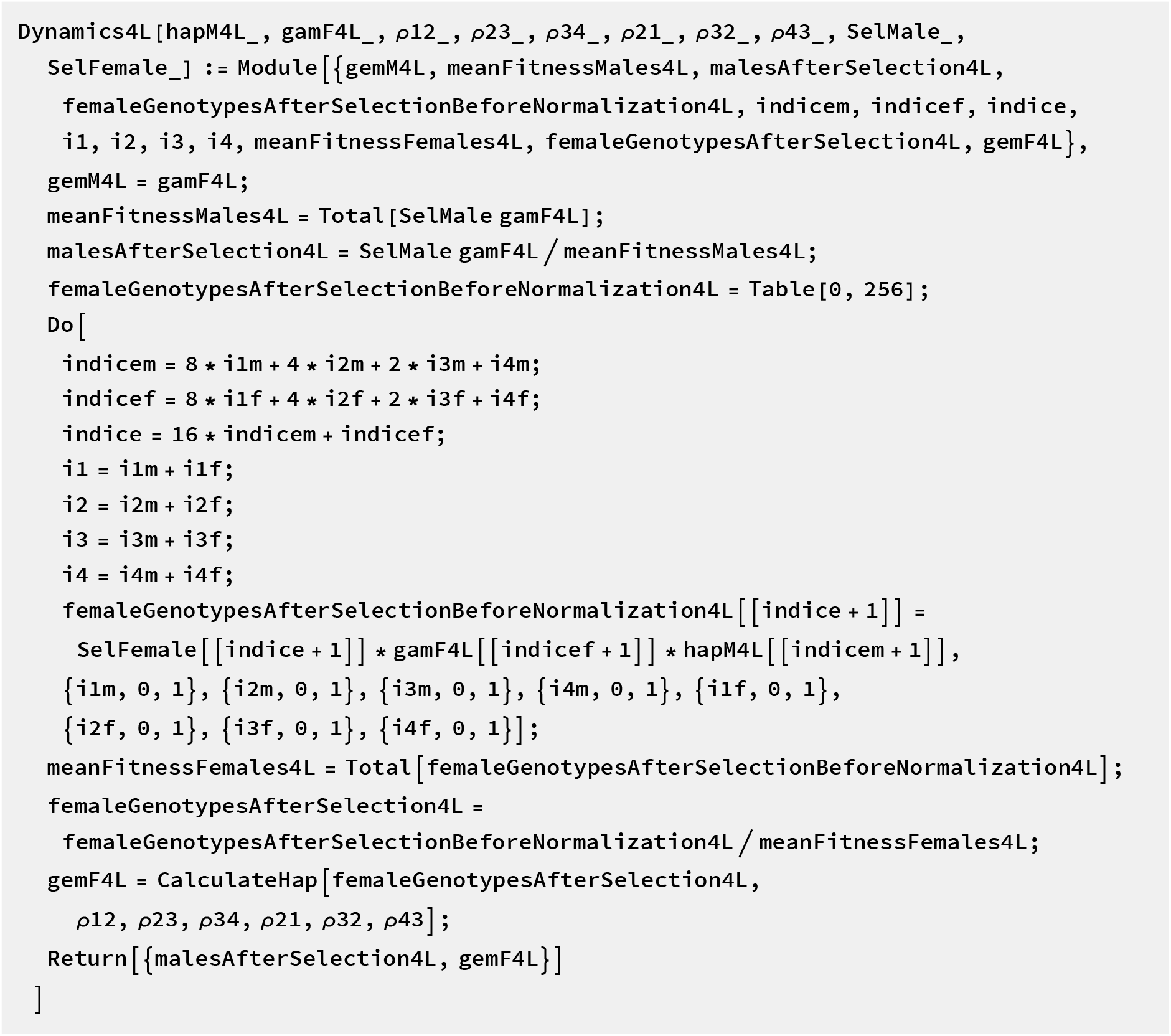

#### The pairwise case: 2 independent DMIs

Define the fitness table for males

~~~
selectionMatrixMales4L =
 Table[(1 - Boole[(i1 == 0 && i2 == 1)]* ϵ_“02”_)*(1 - Boole[(il == 1 && i2 == 0)]*ϵ_“02”_)
  (1 - Boole[(i3 == 0 && i4 == 1)]* η_“02”_)*(1 - Boole[(i3 == 1 && i4 == 0)]*η_“02”_),
 {i1, 0, 1}, {i3, 0, 1}, {i4, 0, 1}] // flatten
~~~

~~~
{1, 1 - *η*_02_, 1 - *η*_20_, 1, 1 - ∈_02_, (1 - ∈_02_)(1 - *η*_02_), (1 - ∈_02_)(1 - *η*_20_), 1 - ∈_02_,
 1 - ∈_20_, (1 -∈_20_)(1 - *η*_02_), (1 - ∈_20_)(1 - *η*_20_), 1 - ∈_20_, 1, 1 - *η*_02_, 1 - *η*_20_, 1}
~~~

Define the fitness table for females

~~~
selectionMatrixFemales4L = Table[0, 256];
Do[
 indicem = 8 * i1m +4 * i2m +2 * i3m + i4m;
 indicef =8 * i1f +4 * i2f +2 * i3f + i4f;
 indice =16 *indicem +indicef;
 i1 =i1m + i1f;
 i2 =i2m + i2f;
 i3 =i3m + i3f;
 i4 =i4m + i4f;
selectionMatrixFemales4L [[indice + 1]] = 1 *(1 + s1) ^^^(Boole [i1 == 1] + Boole [i2 == 1]) *
 (1 + s2) ^ (Boole[i3 == 1] + Boole[i4 == 1]) * (1 - Boole[i2 - i1 == 1] * ϵ_“01”_) *
(1 - Boole [i1 - i2 == 1] * ϵ_“10”_) * (1 - Boole[i1 == 0&&i2 == 2] *ϵ_“02”_) * (1 - Boole[i1 == 2 && i2 == 0] * ϵ_“20”_) * (1 - Boole [i1 == 1 &&
i2 == 1] * ϵ_“10”_) *
(1 - Boole [i4 - i3 == 1] * ϵ_“11”_) * (1 - Boole [i3 - i4 == 1] * ϵ_“10”_;) * (1 - Boole [i3 == 0 && i4 == 2] * ϵ_“02”_) *
(1 - Boole [i3 == 2&&i4 == 0] *ϵ_“20”_) * (1 - Boole [i3 == 1&&i4 == 1] *ϵ_“11”_), {i1m, 0, 1},
{i2m, 0, 1}, {i3m, 0, 1}, {i4m, 0, 1}, {i1f, 0, 1}, {i2f, 0, 1}, {i3f, 0, 1}, {i4f, 0, 1}]
~~~

~~~
hapM4L = {h1, h2, h3, h4, h5, h6, h7, h8, h9, h10, h11, h12, h13, h14, h15, h16};
~~~

~~~
gamF4L = {g1, g2, g3, g4, g5, g6, g7, g8, g9, g10, g11, g12, g13, g14, g15, g16};
~~~

Display in which order the haplotypes are calculated

~~~
Do [Print [i1, i2, i3, i4], {i1, 0, 1}, {i2, 0, 1}, {i3, 0, 1}, {i4, 0, 1}]
0000
0001
0010
0011
0100
0101
0110
0111
1000
1001
1010
1011
1100
1101
1110
1111
~~~

#### Test for the dynamics for the first pair of loci

Check if the dynamics obtained matches with the 2 locus case define previously

~~~
StartCondf = {g1, g2, g3, g4, 0, 0, 0, 0, 0, 0, 0, 0, 0, 0, 0, 0};
StartCondm = {hi, h2, h3, h4, 0, 0, 0, 0, 0, 0, 0, 0, 0, 0, 0, 0};

test = Dynamics4L[StartCondm, StartCondf, ρ12, ρ23, ρ34,
       ρ21, ρ32, ρ43, selectionMatrixMales4L, selectionMatrixFemales4L] /.
{*η*_“01”_ → 0, *η*_“10”_ → 0, *η*_“11”_ → 0} // FullSimplify // Together;

$Aborted

claudiaversion = haplotypeDynamicsX[1] [{s2, η_”02”_, η_“20”_, ρ34}] [
  {{g1, g2, g3, g4}, {h1, h2, h3, h4}}] // FullSimplify;

claudiaversion[[2]] - test[[1, 1 ;; 4]] // FullSimplify
{0, 0, 0, 0}
~~~

#### Test for the dynamics for the first pair of loci

Check if the dynamics obtained matches with the 2 locus case define previously

~~~
StartCondf = {g1, 0, 0, 0, g2, 0, 0, 0, g3, 0, 0, 0, g4, 0, 0, 0};
StartCondm = {h1, 0, 0, 0, h2, 0, 0, 0, h3, 0, 0, 0, h4, 0, 0, 0};
test = Dynamics4L[StartCondm, StartCondf, ρ12,
  ρ23, ρ34, selectionMatrixMales4L, selectionMatrixFemales4L] /.
{ϵ_“01”_ → 0, ϵ_“10”_ → 0, ϵ_“11”_ → 0} // FullSimplify // Together;
claudiaversion = haplotypeDynamicsX[1] [{s1, ϵ_“02”_, ϵ_“20”_, ρ12}] [
{{g1, g2, g3, g4}, {h1, h2, h3, h4}}] // FullSimplify;
claudiaversion[[2]] -
 {test[[1, 1]], test[[1, 5]], test[[1, 9]], test[[1, 13]]} // Simplify
{0, 0, 0, 0}
claudiaversion[[1]] -{test[[2, 1] ], test[[2, 5]], test[[2, 9]], test[[2, 13]]} // FullSimplify
 { 0, 0, 0, 0}
Dynamic4LGen = Dynamics4L[gamF4L, hapM4L, ρ12, ρ23, ρ34,
 ρ21, ρ32, ρ43, selectionMatrixMales4L, selectionMatrixFemales4L];
Jac4LGen = D[Dynamic4LGen, {{g1, g2, g3, g4, g5, g6, g7, g8, g9, g10, g11, g12, g13, g14,
g15, g16, h1, h2, h3, h4, h5, h6, h7, h8, h9, h10, h11, h12, h13, h14, h15, h16}}] ;
~~~

#### All heterozygote advantage are equal and epistasis is lethal

~~~
param = {s1 → s, s2 → s, ϵ_“01”_ → 0, ϵ_“10”_ → 0, ϵ_“02”_ → 1,
  ϵ_“20”_ -> 1, ϵ_“11”_ → 0, *η*_“01”_ → 0, *η*_“10”_ → 0, *η*_“02”_ → 1, *η*_“20”_ -> 1, *η*_“11”_ → 0};
SimplerDym = Dynamic4LGen /. param;
~~~

Define the Jacobian Matrix

~~~
Jac4LGen =
 D [Flatten [SimplerDym], {{g1, g2, g3, g4, g5, g6, g7, g8, g9, g10, g11, g12,    g13, g14, g15, g16, h1, h2, h3, h4, h5, h6, h7, h8, h9, h10, h11, h12, h13, h14, h15, h16}}] ;
~~~

First case exclusion: 0000 haplotype

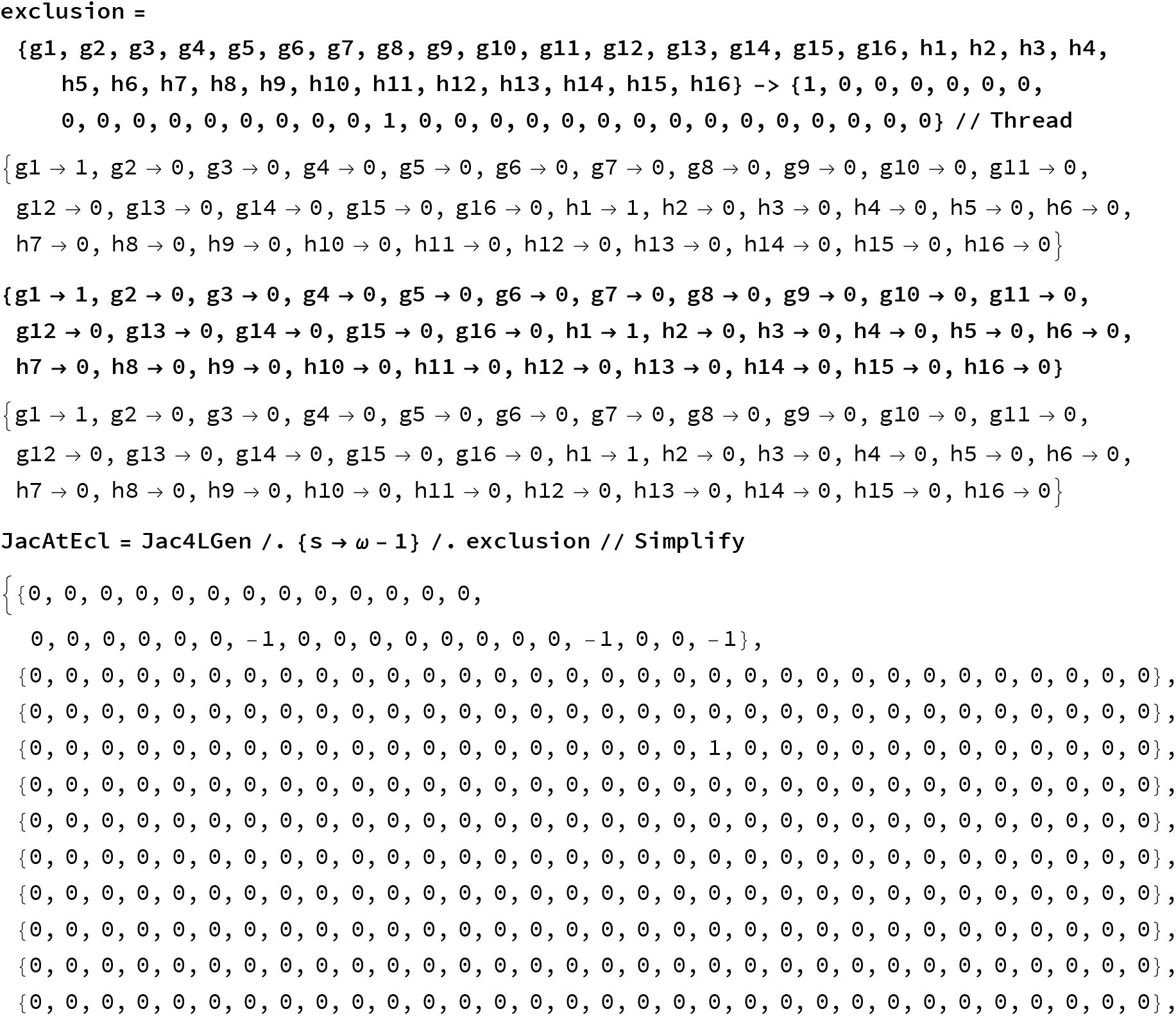
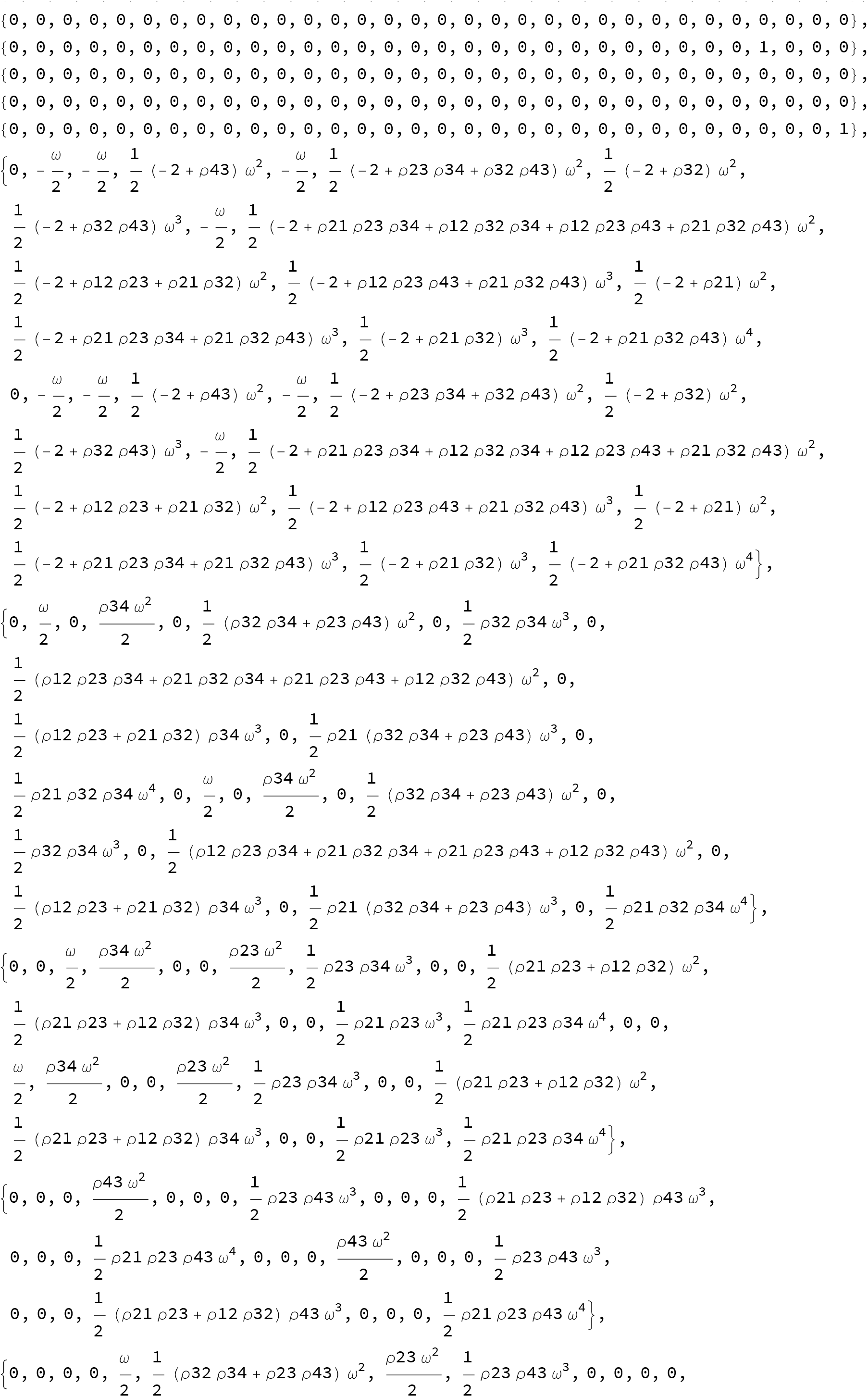
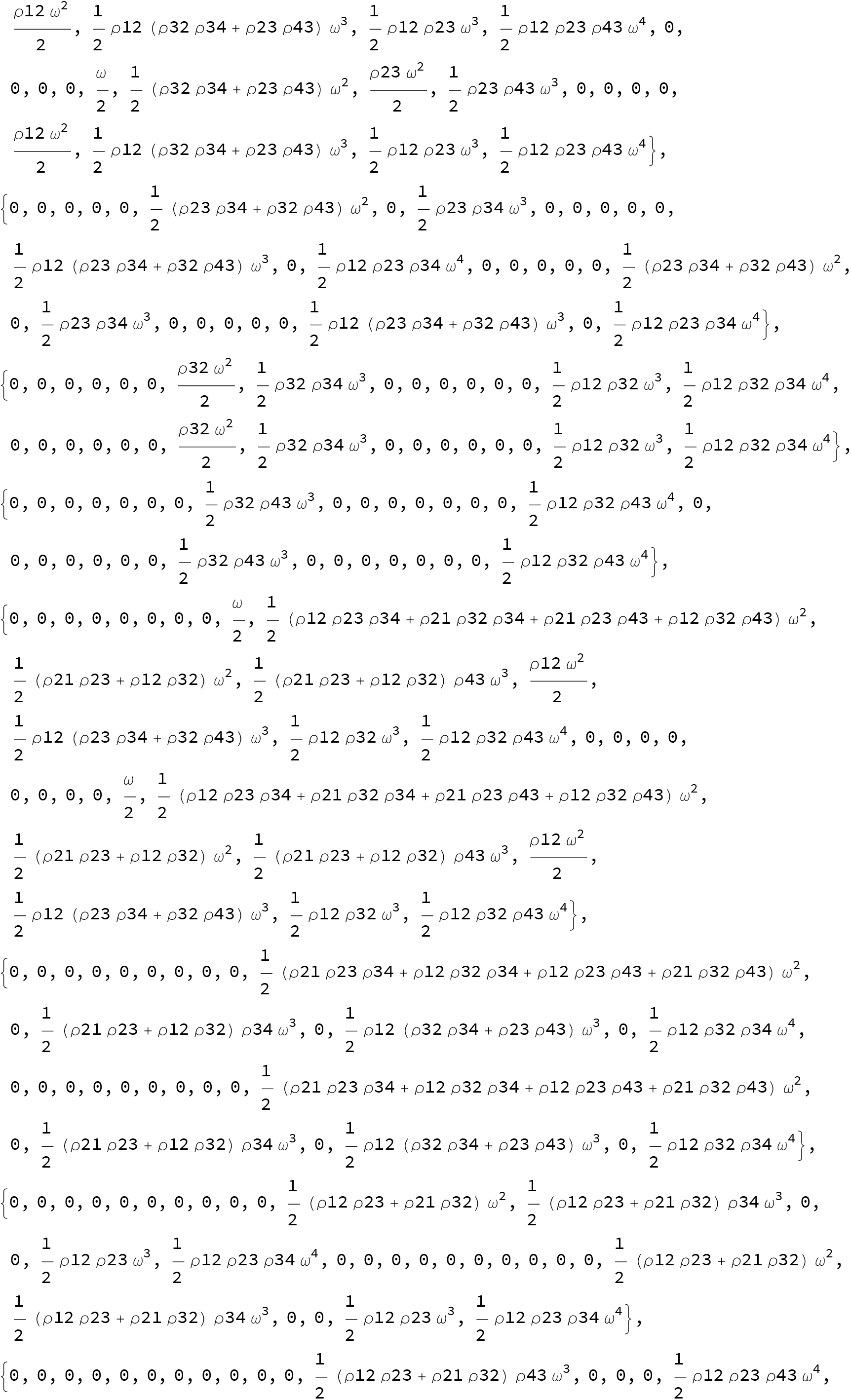
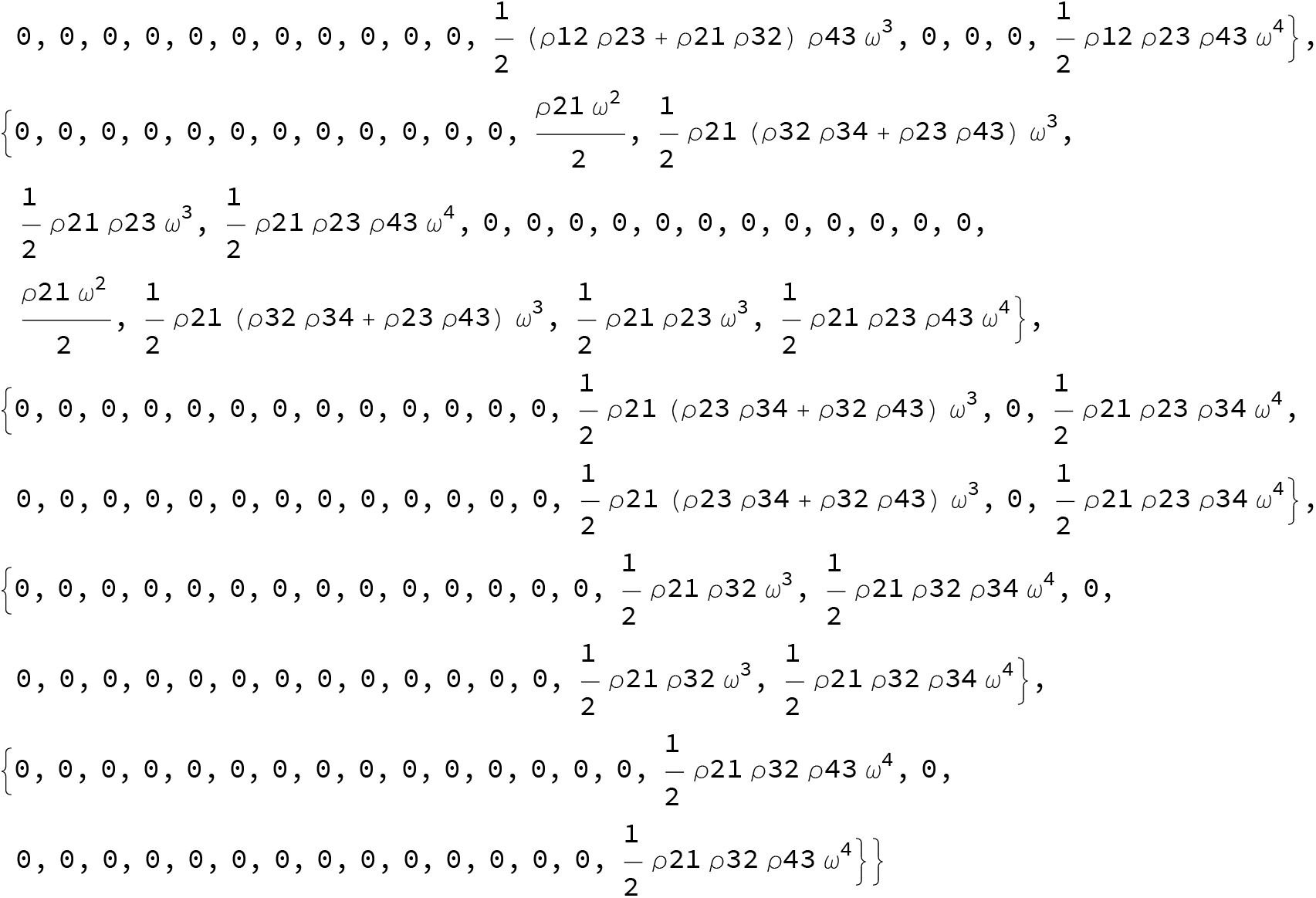

JacAtEcl // Simplify // MatrixForm

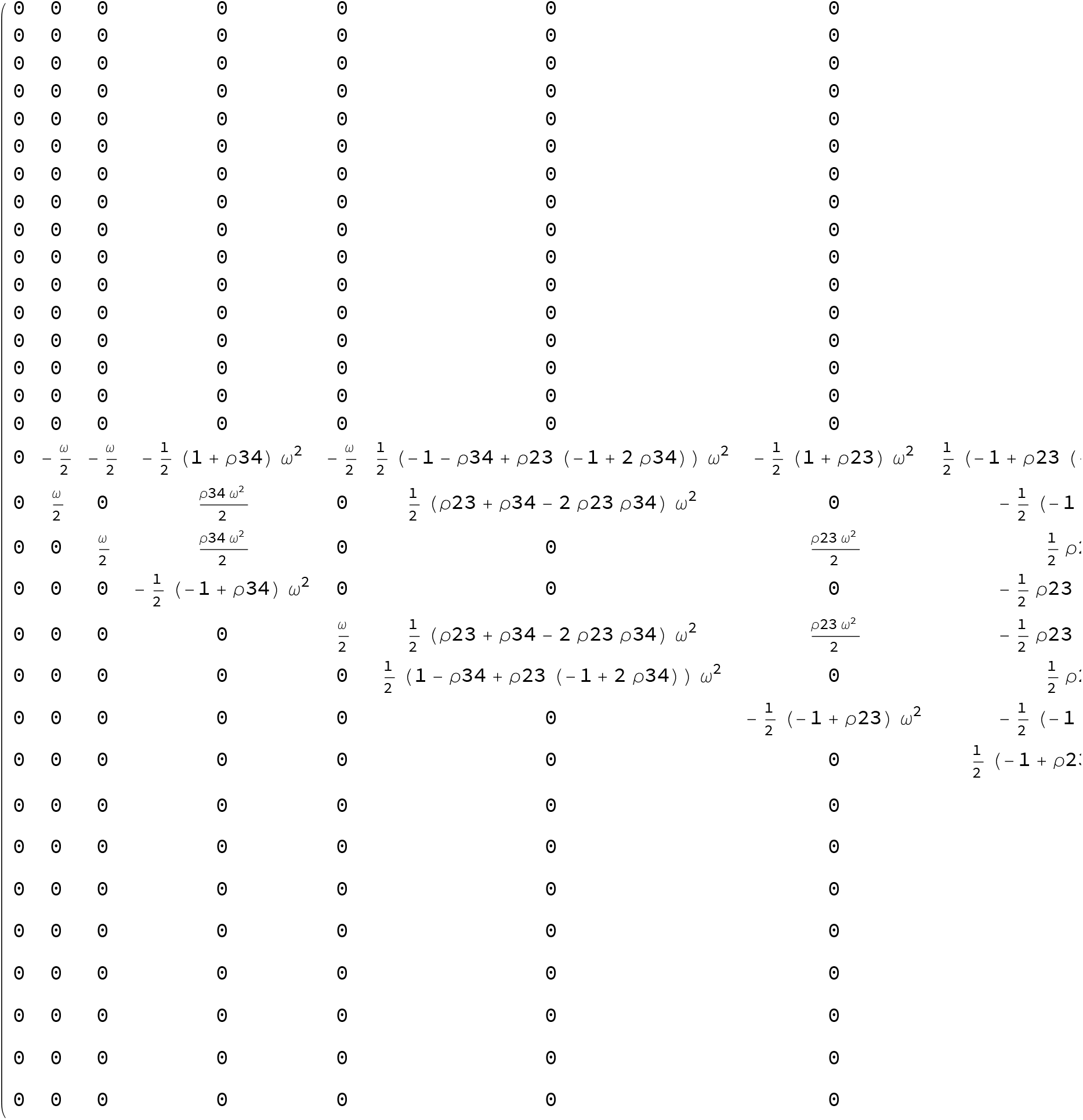

Calculate the Eigenvalues

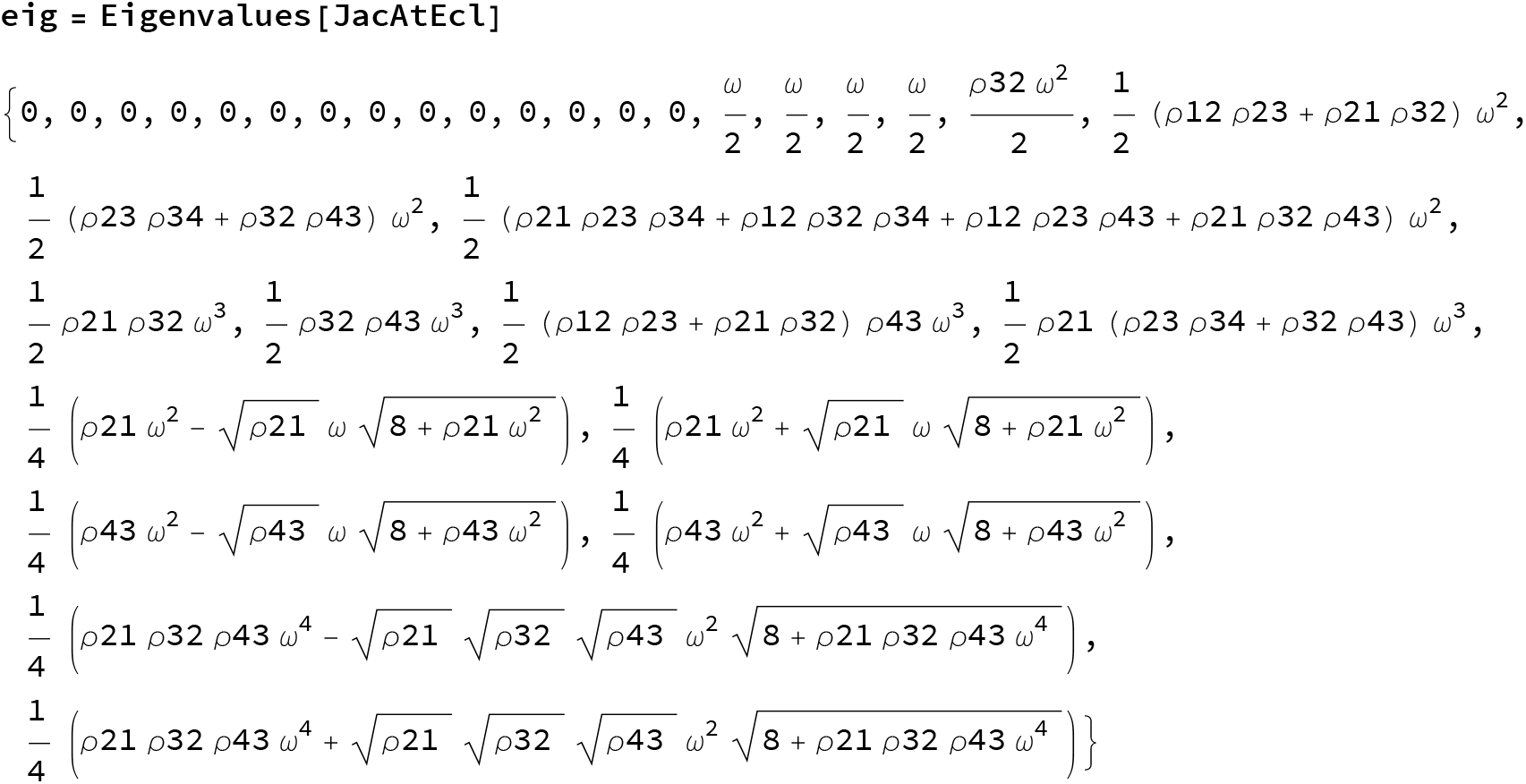

Finally replace *ρ*21 by 1-*ρ*12. This is done only now to avoid Mathematica using 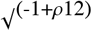 as an intermediate step

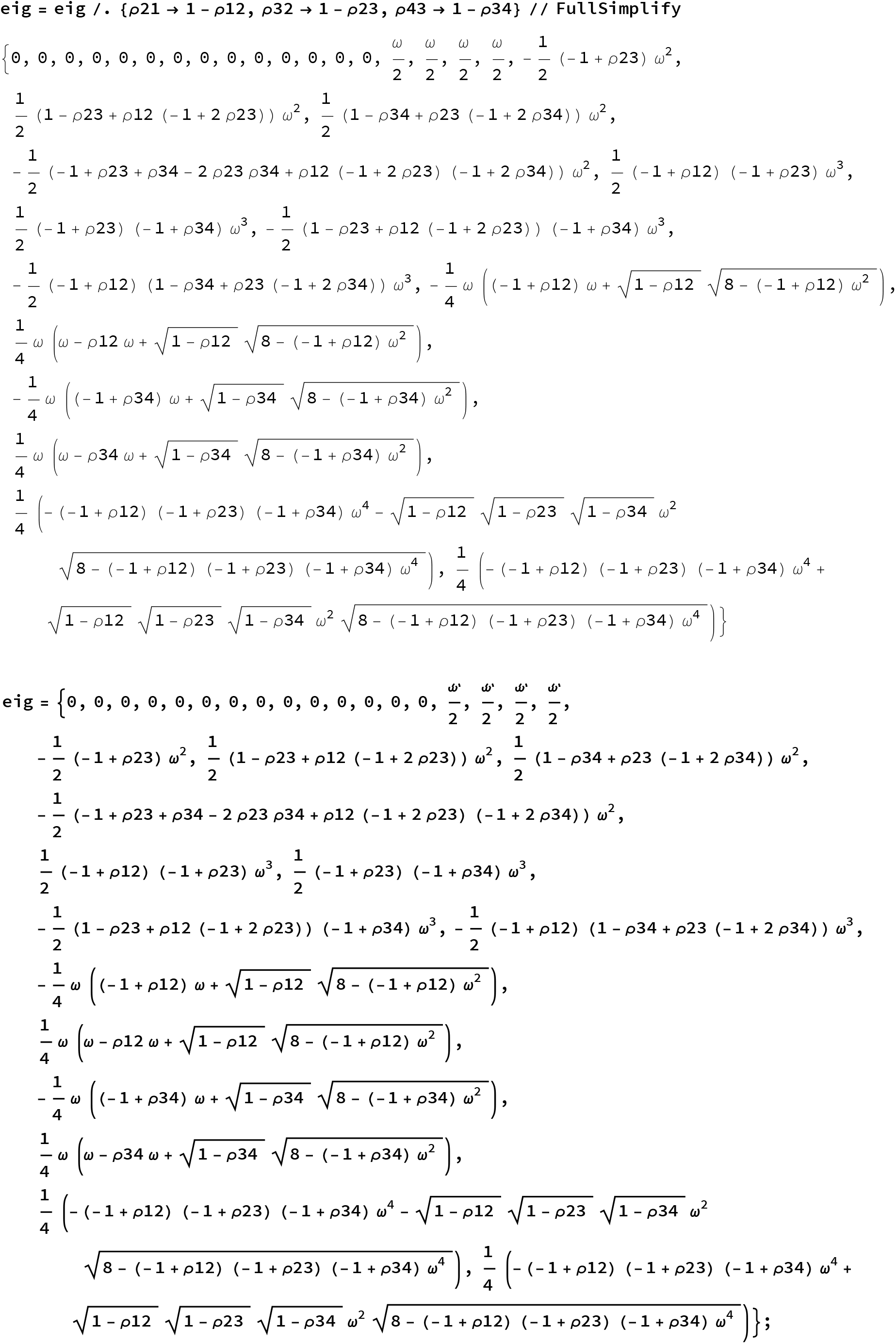

If all loci are on different chromosomes

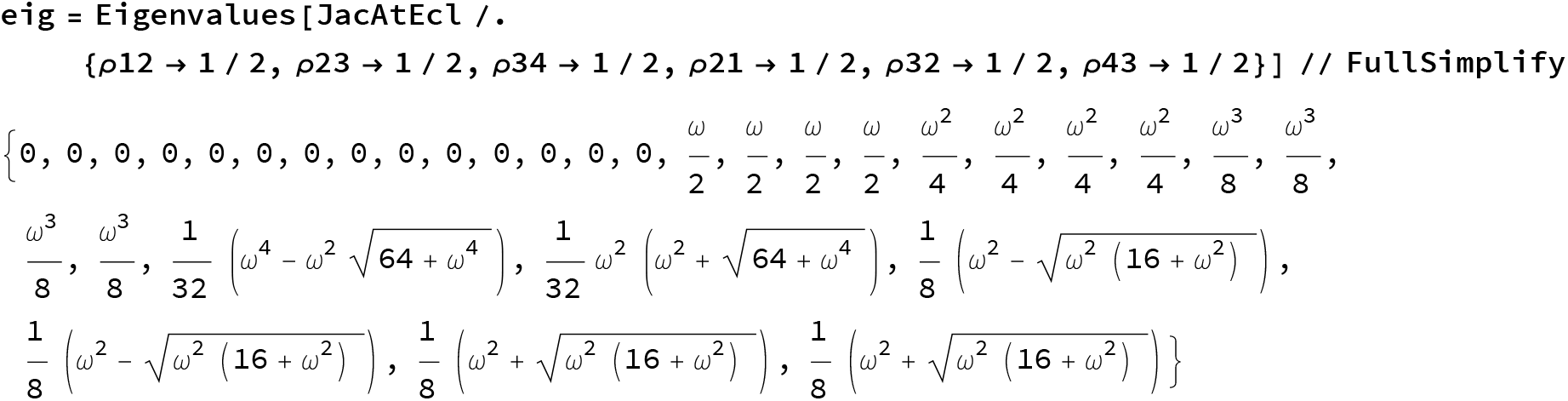

Case 1 Independent pairs of loci

For simplicity we first check that all Eigenvalues are smaller than 1:

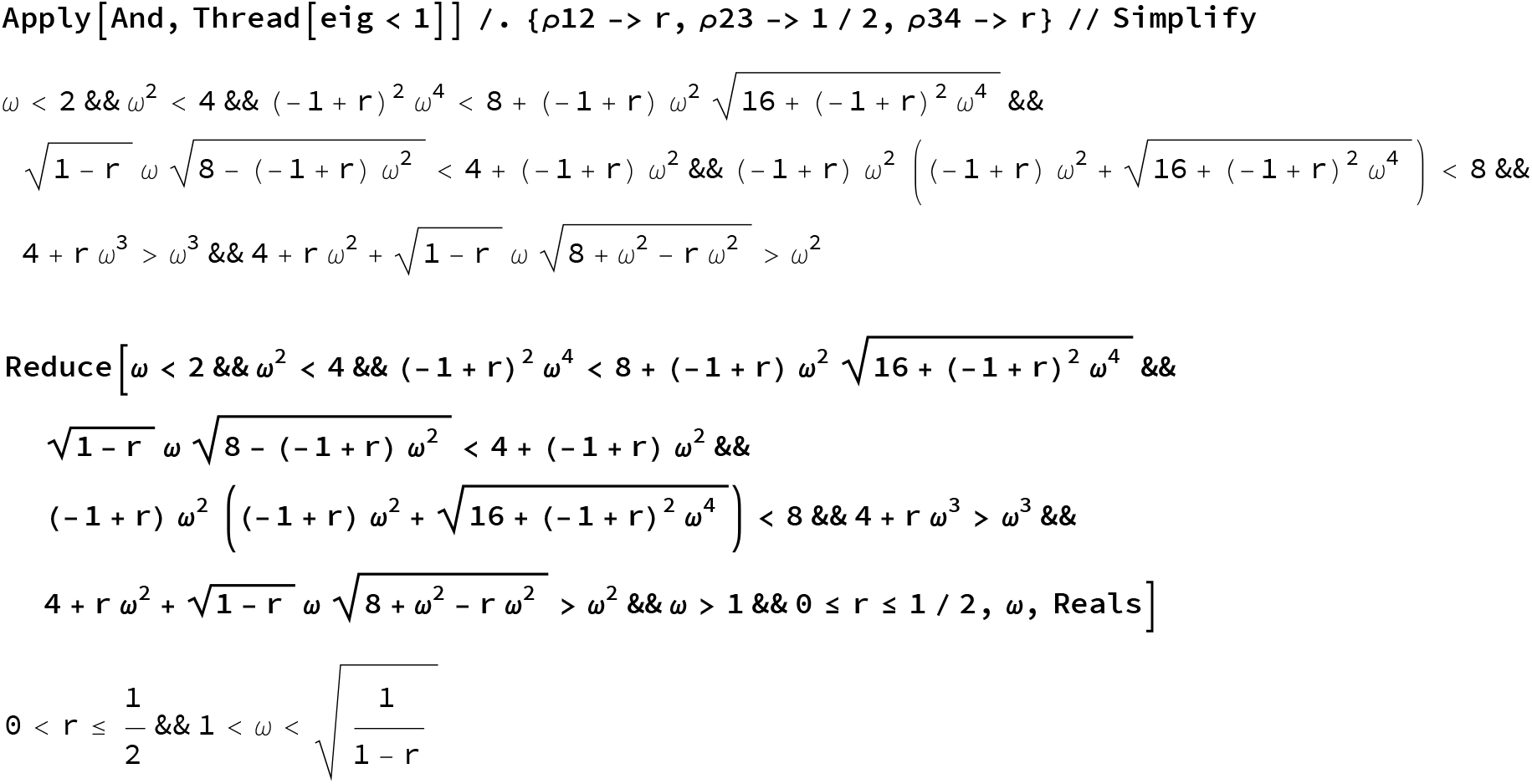

Then we check that all Eigenvalues are larger than - 1 :

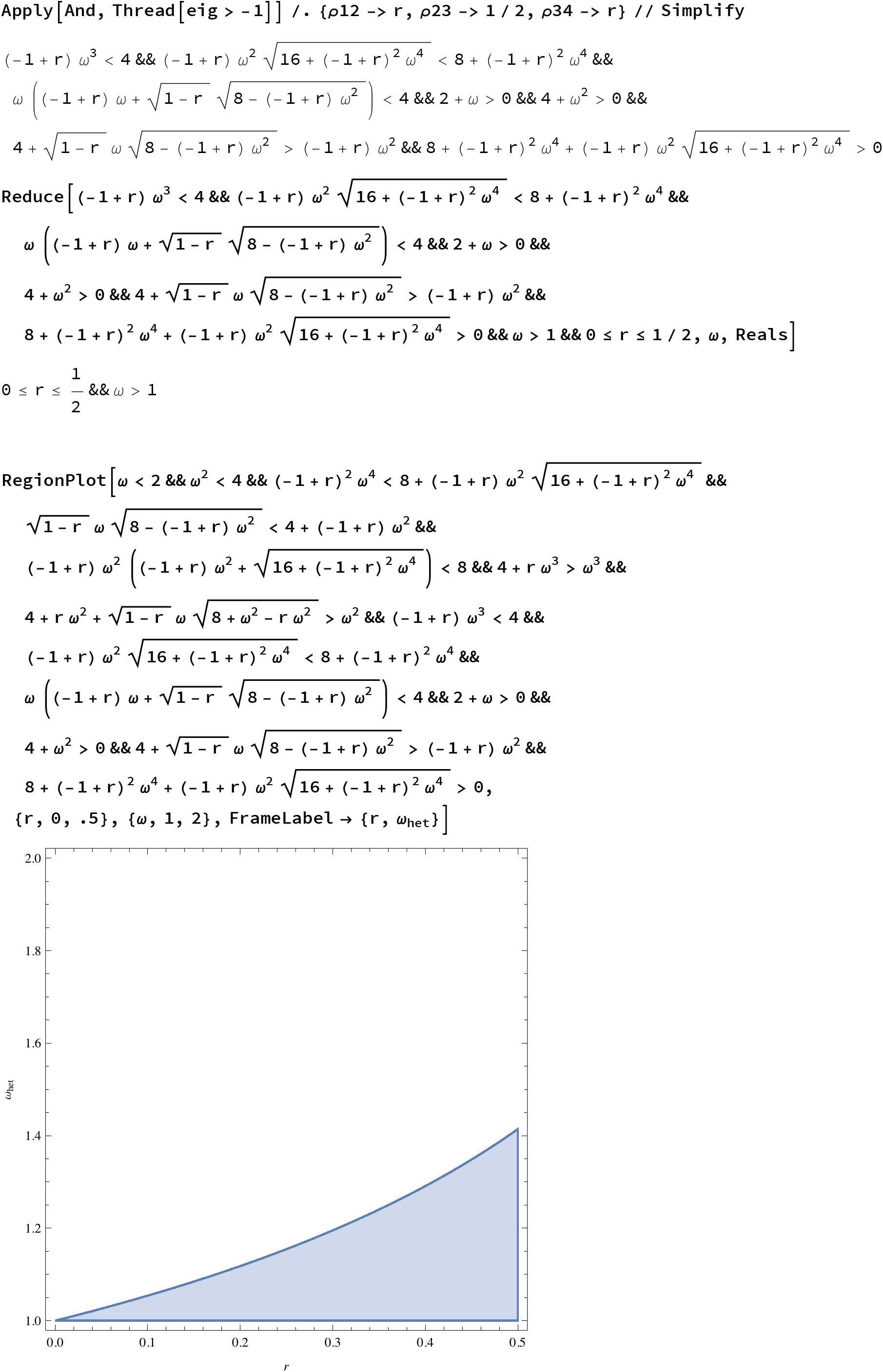

Case 2 the second and third loci are in TL (r->0)

For simplicity we first check that all Eigenvalues are smaller than 1:

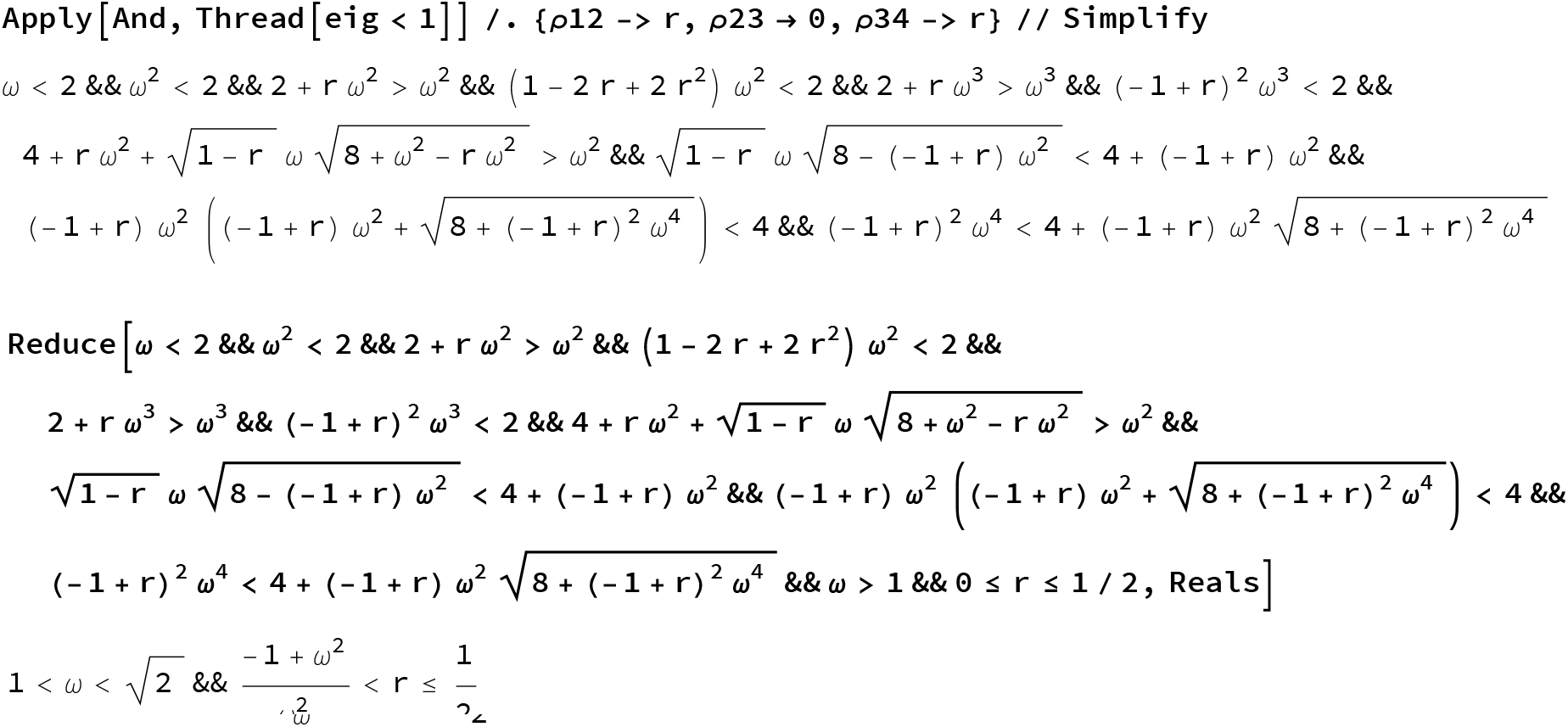

Then we check that all Eigenvalues are larger than - 1 :

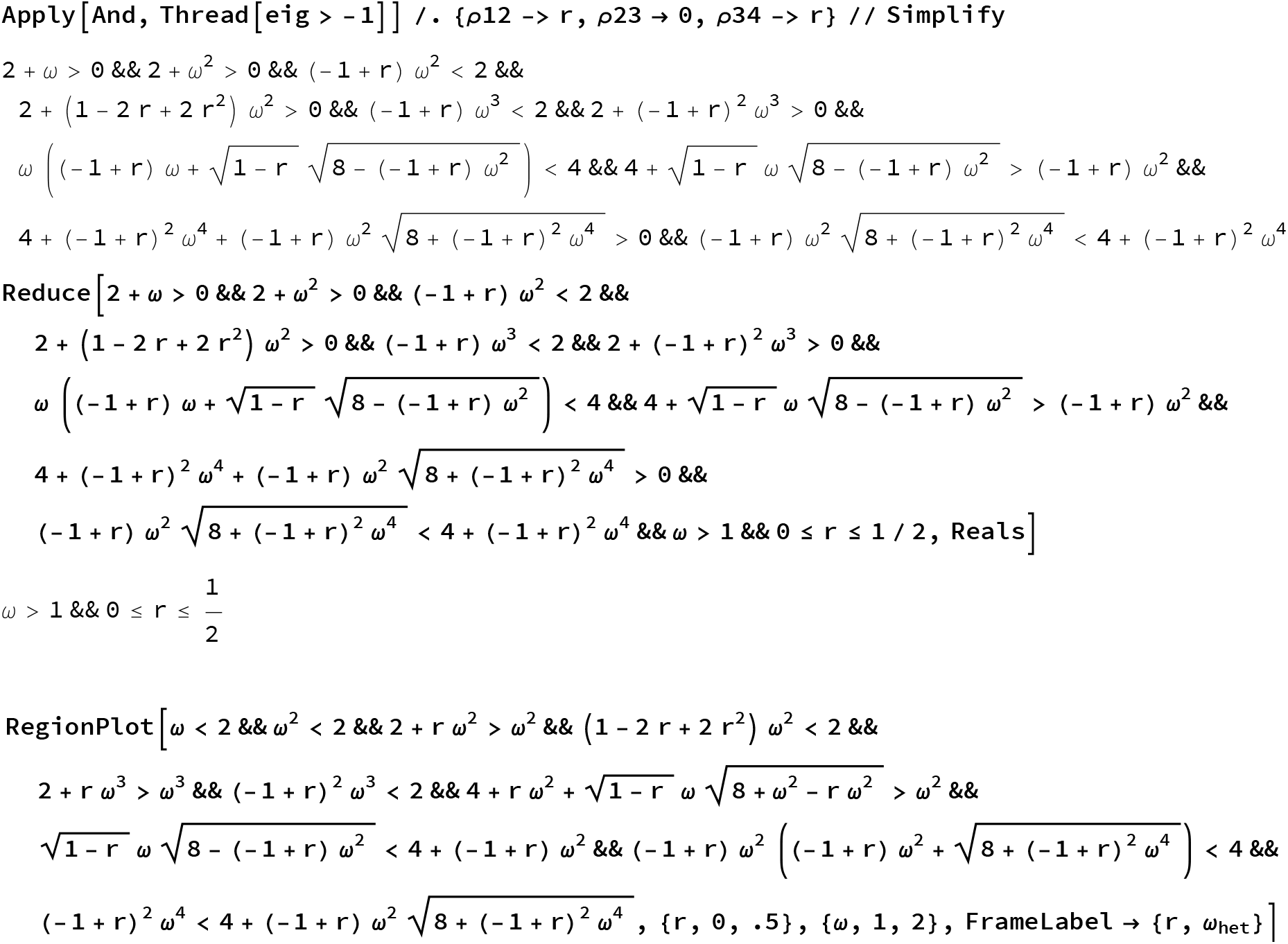
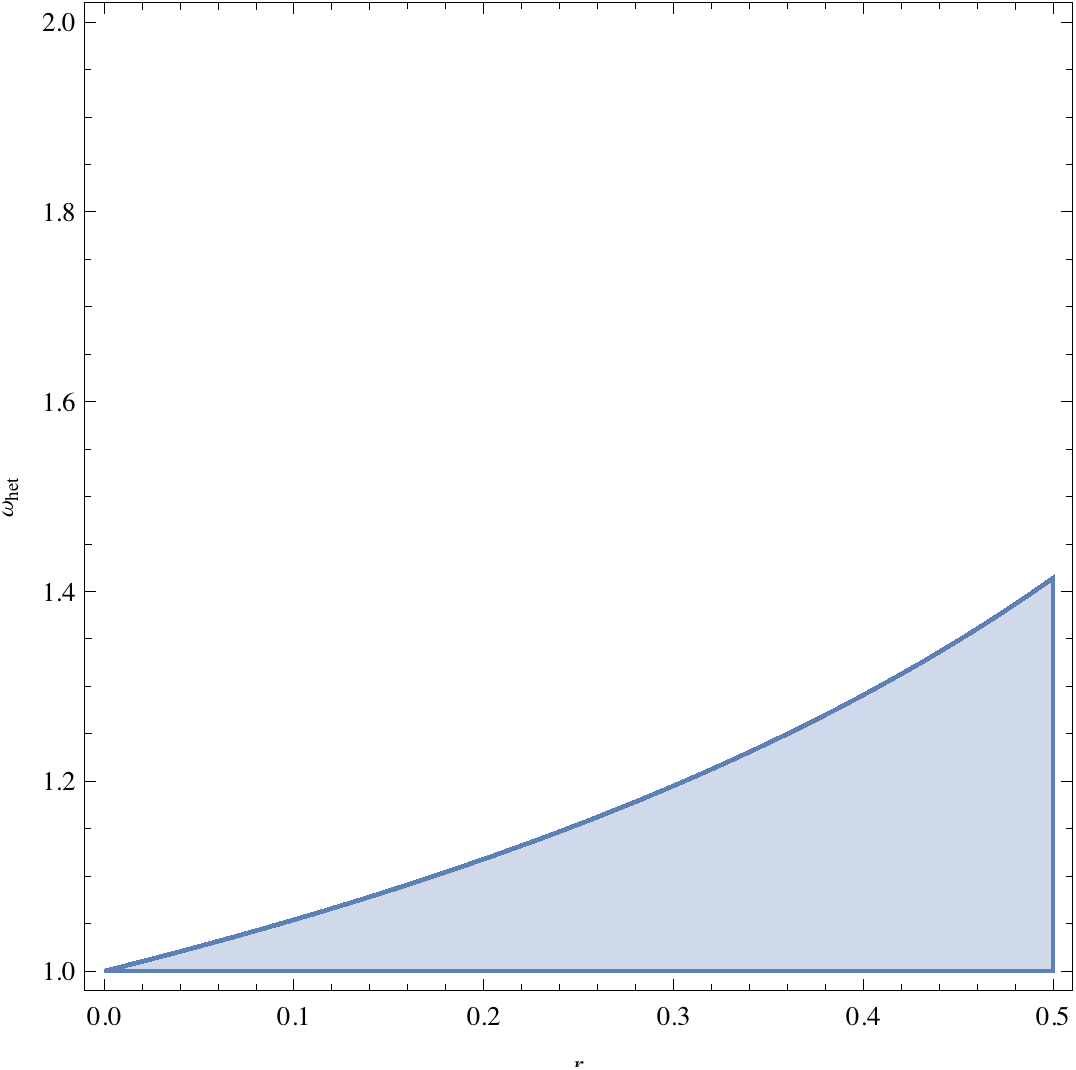

Case 3 : General case

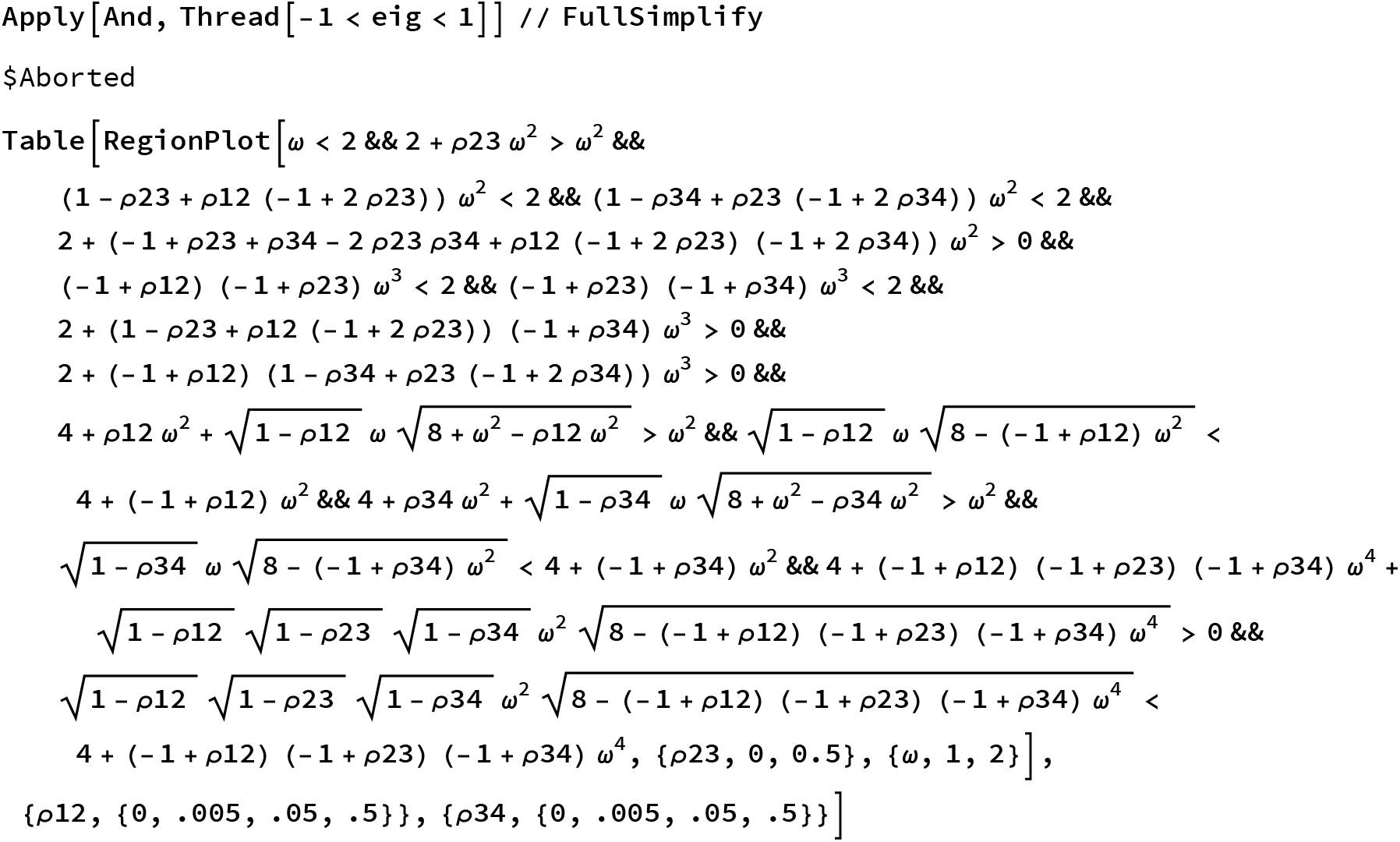
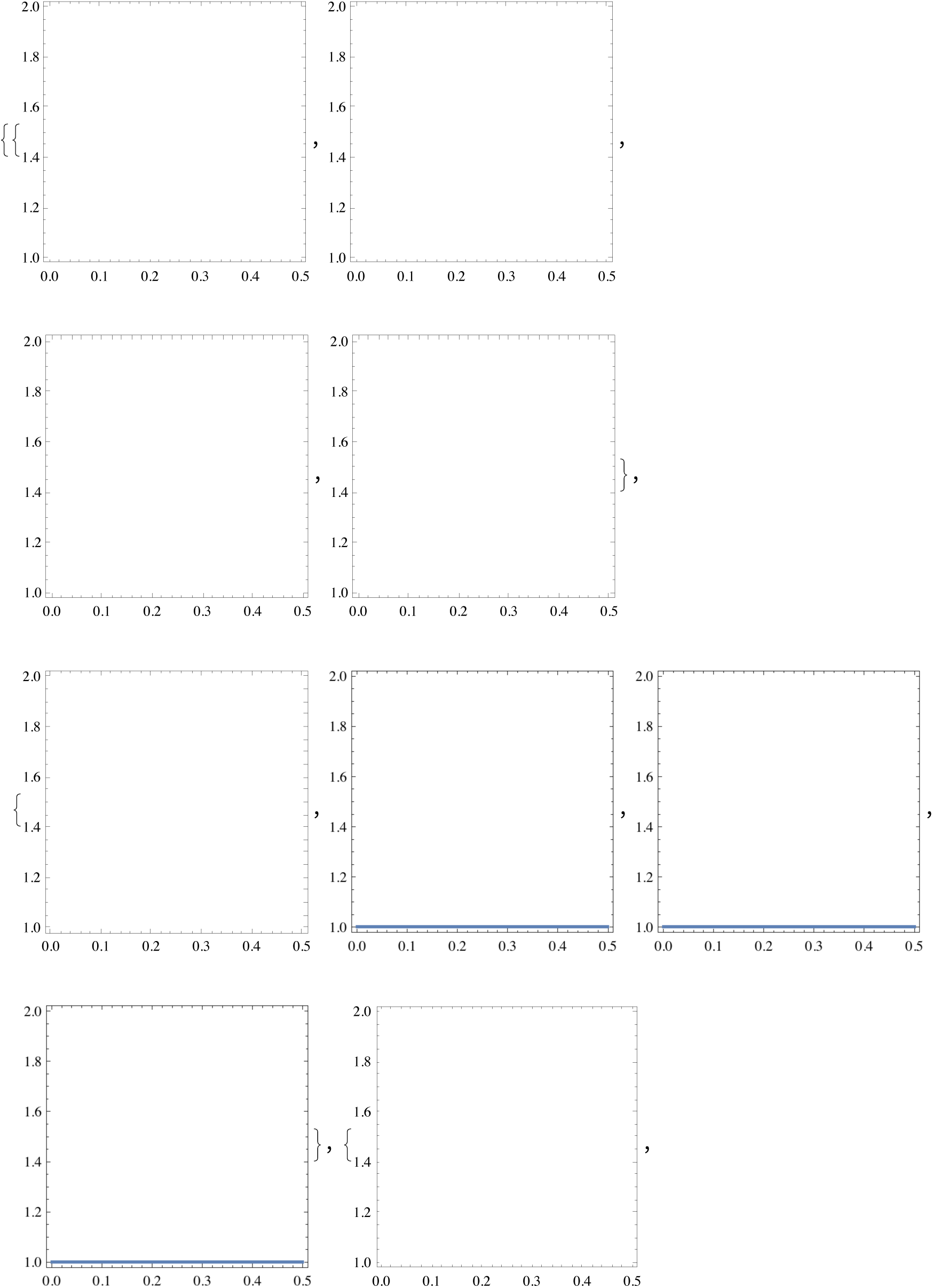
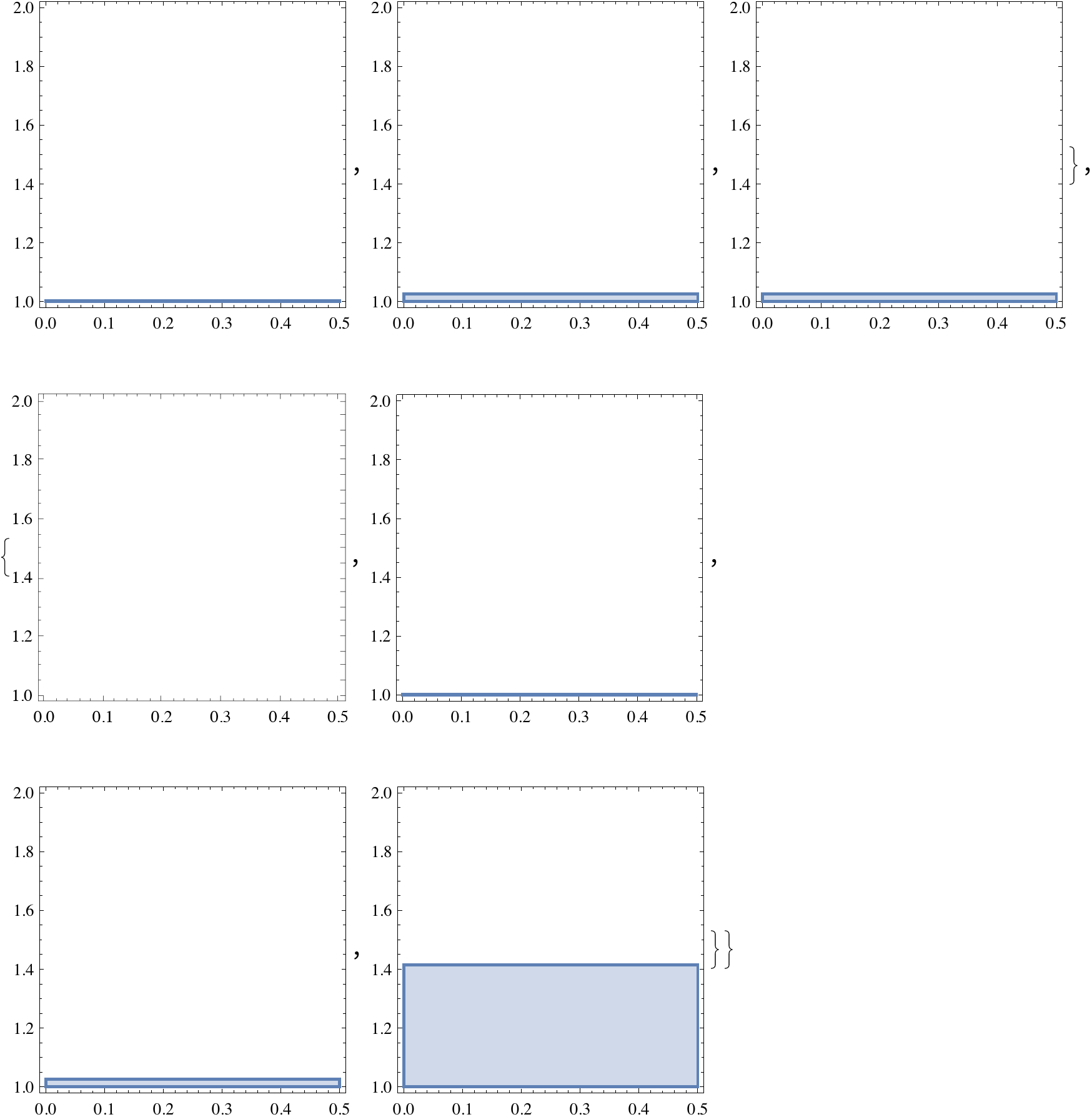
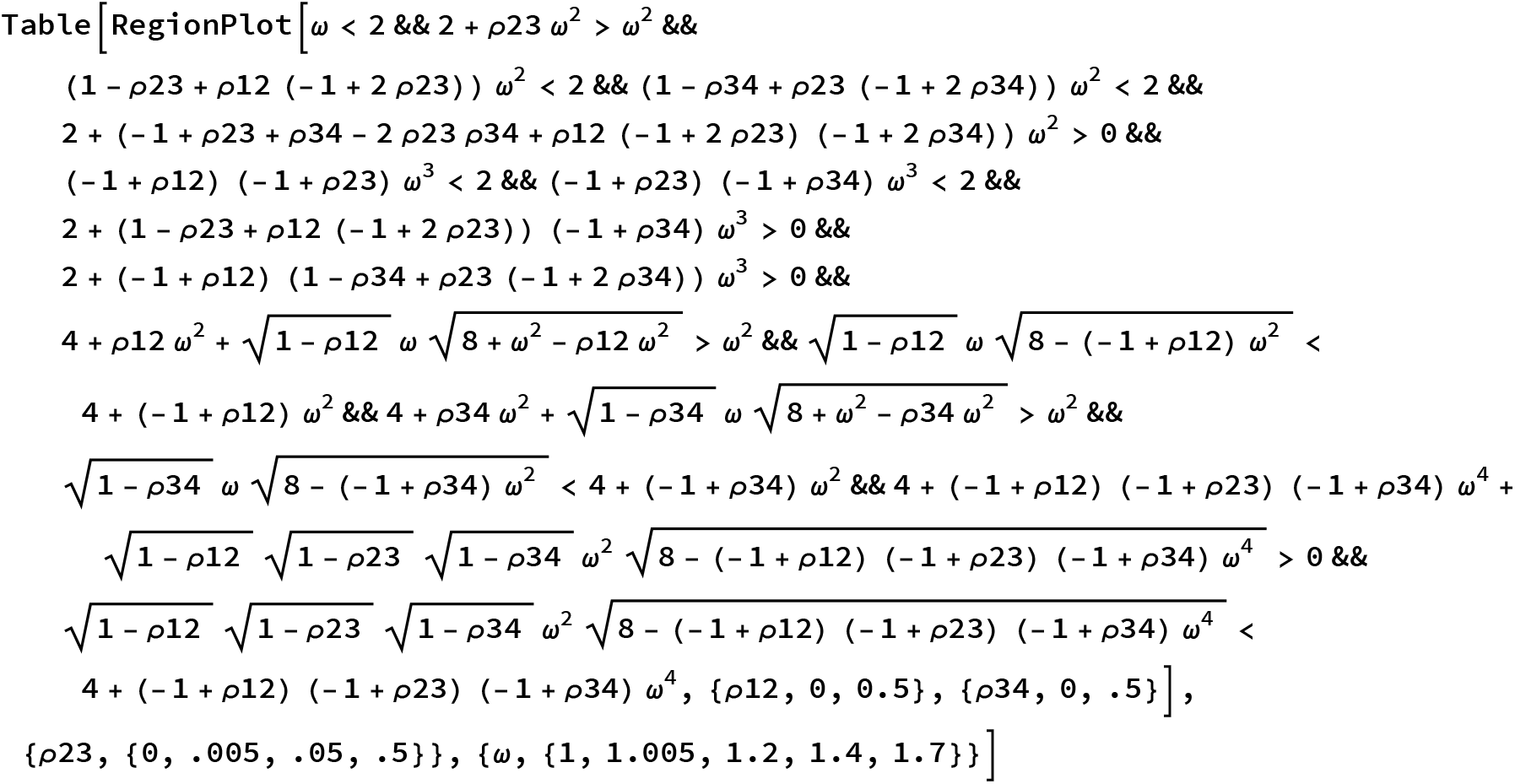
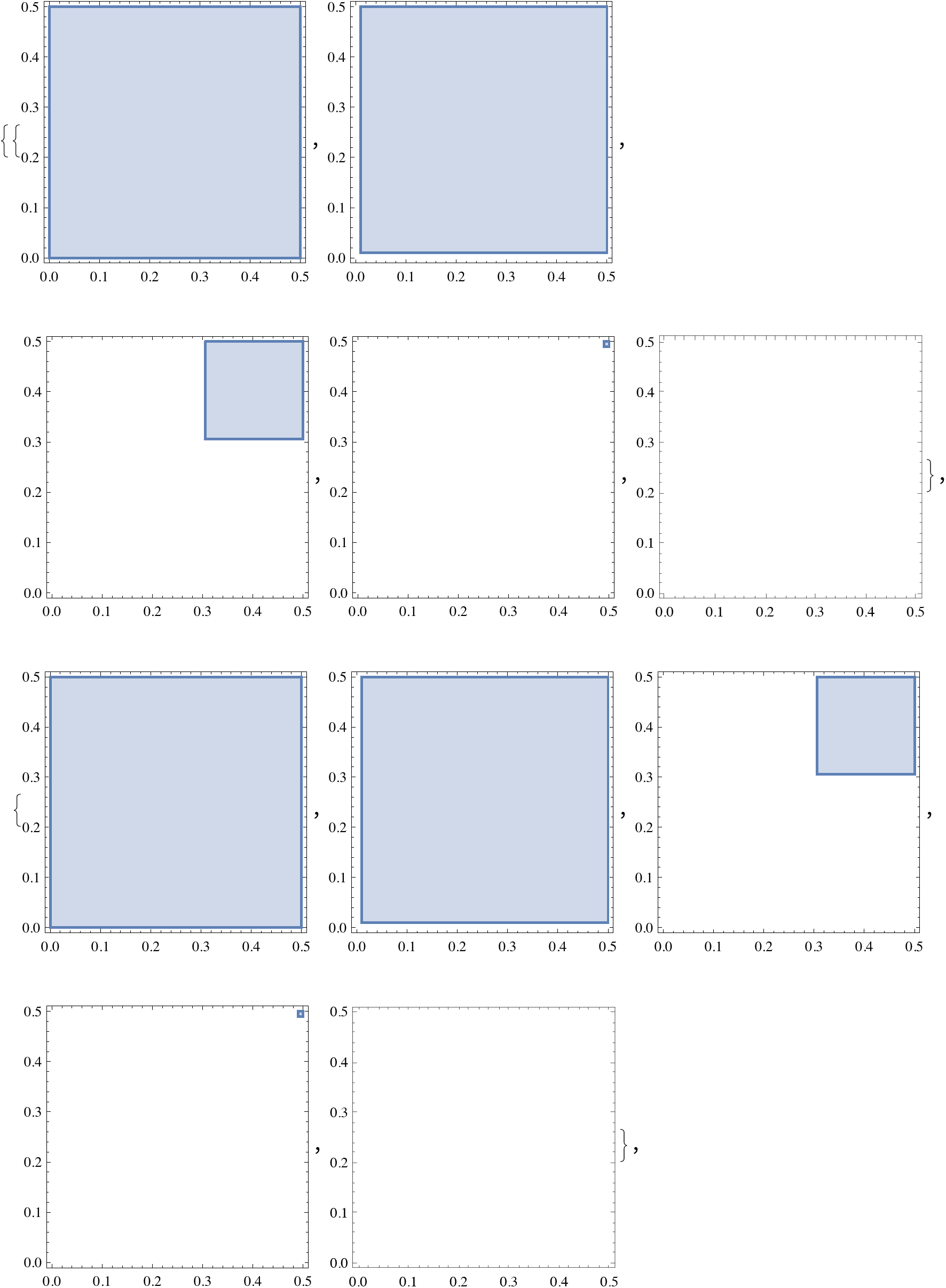
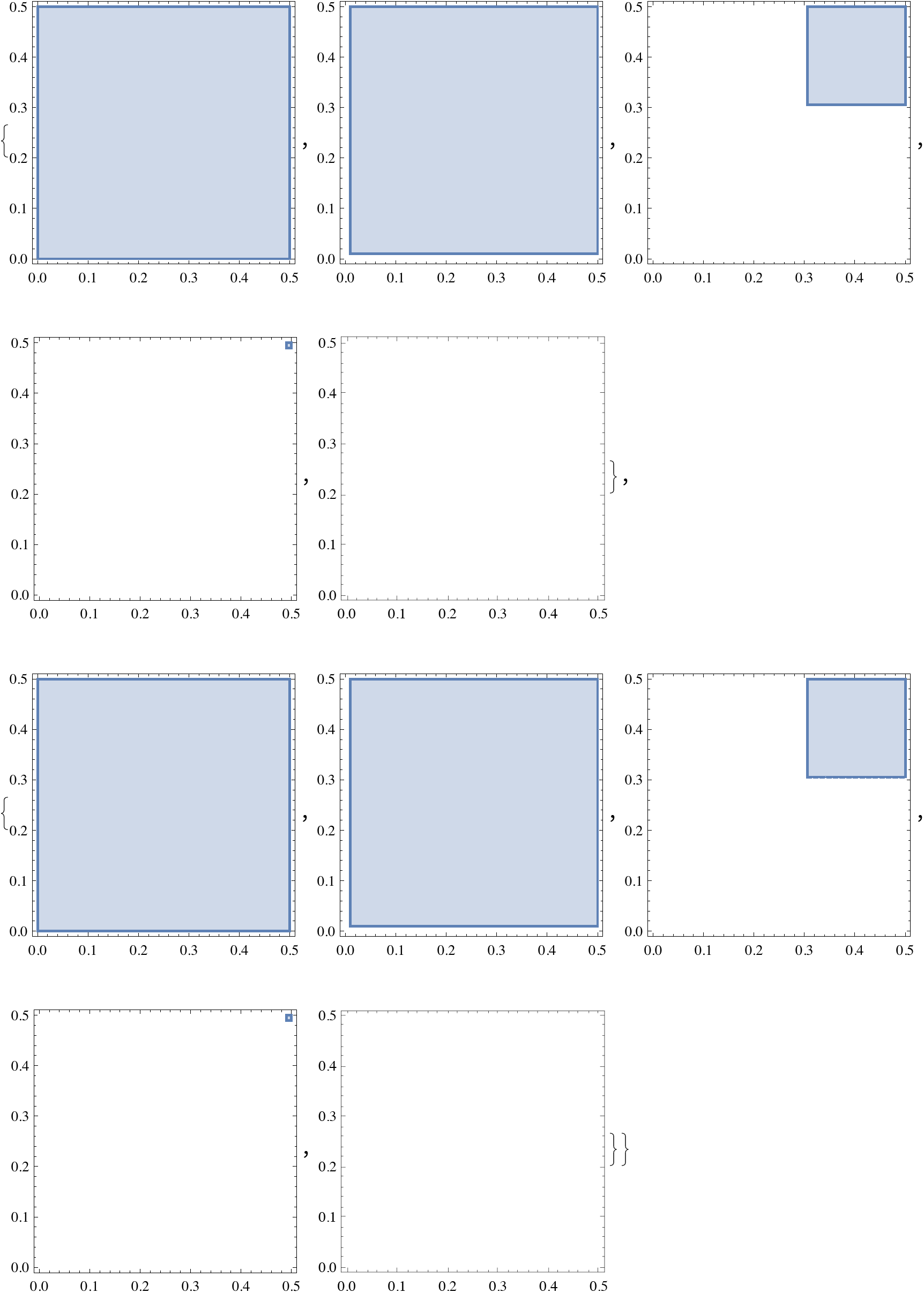

Conditions for local stability of the exclusion case for arbitrary recombination rate

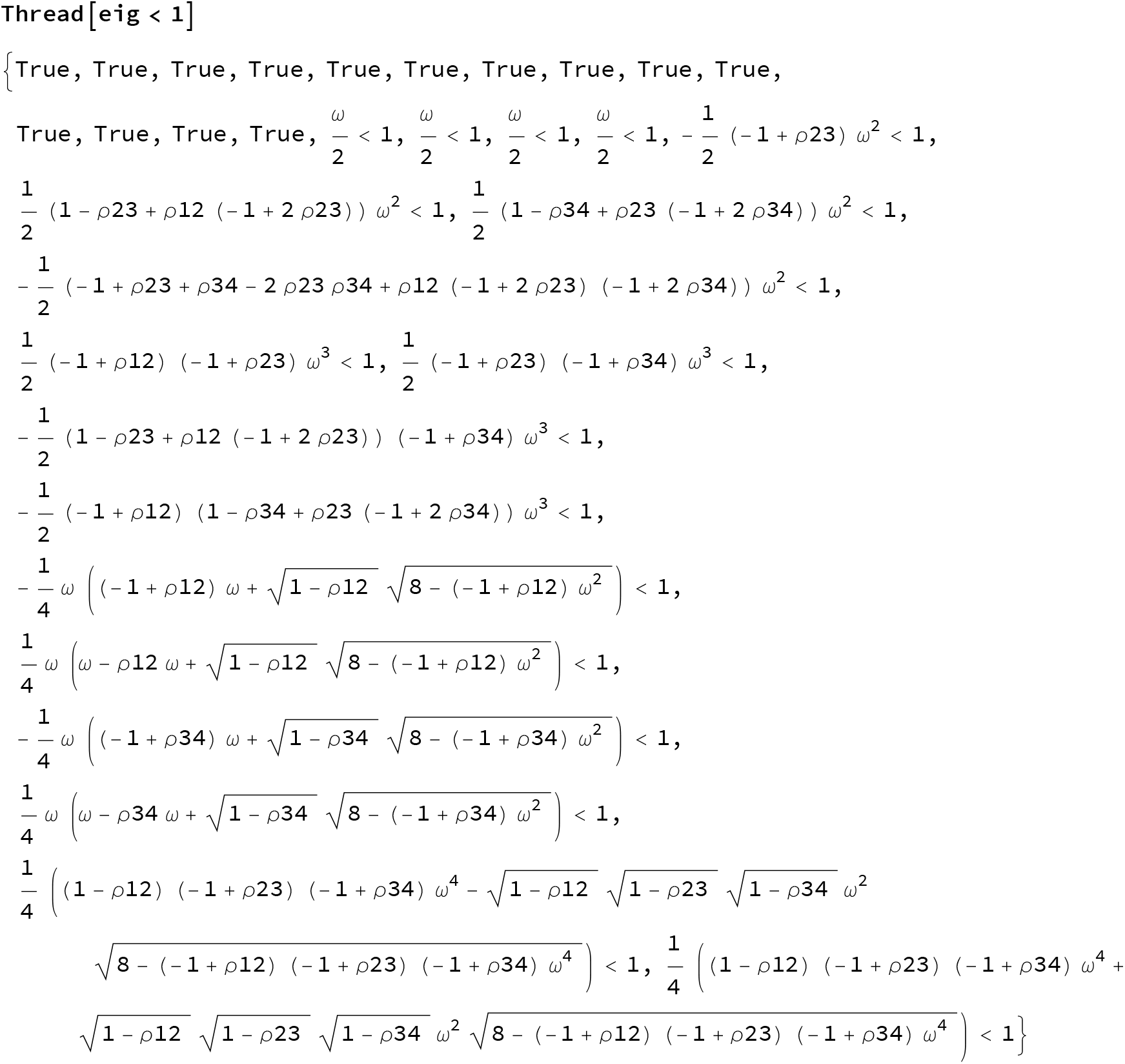
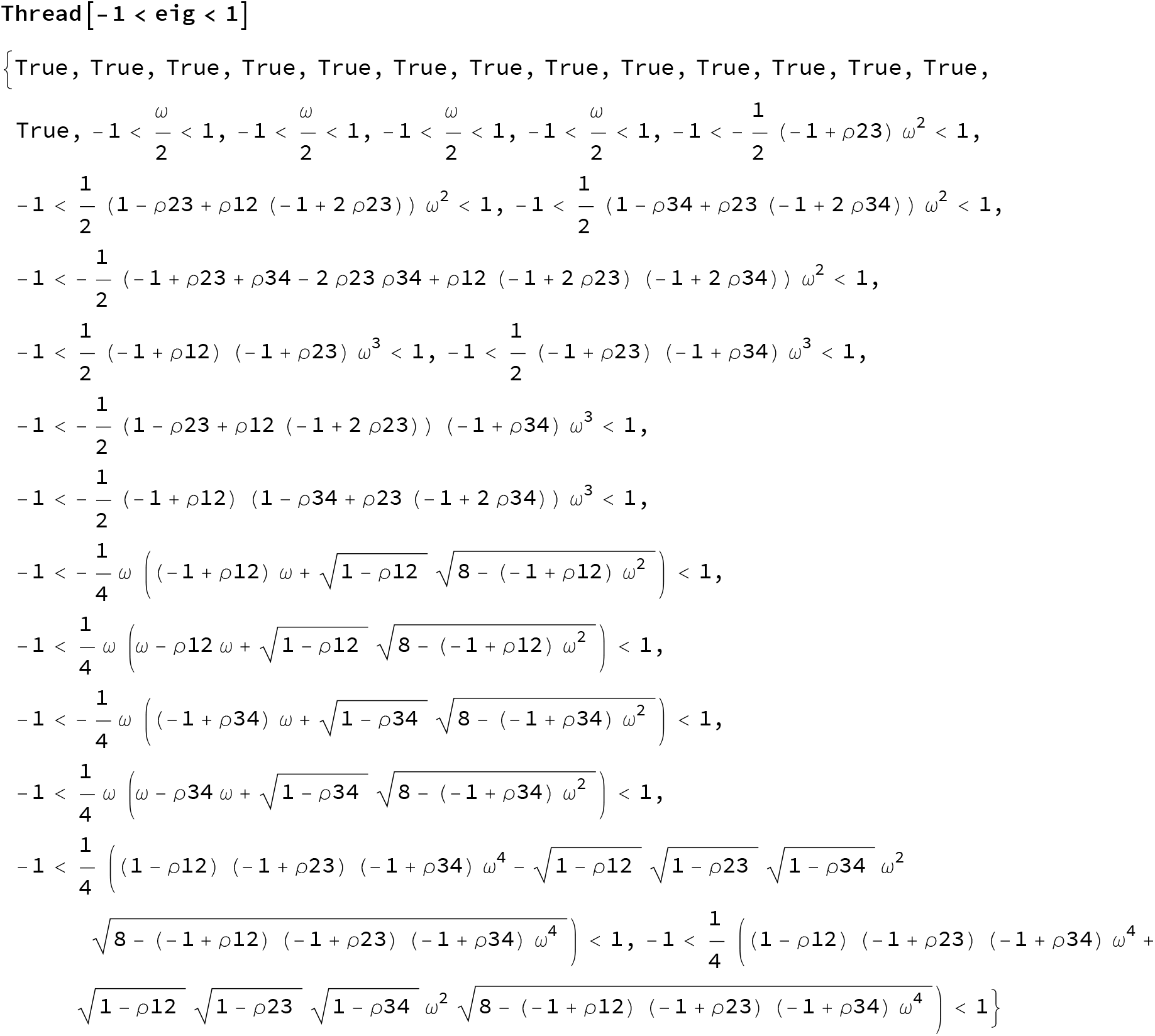

Below we check each eigenvalues one by one and infer the boundary on ω

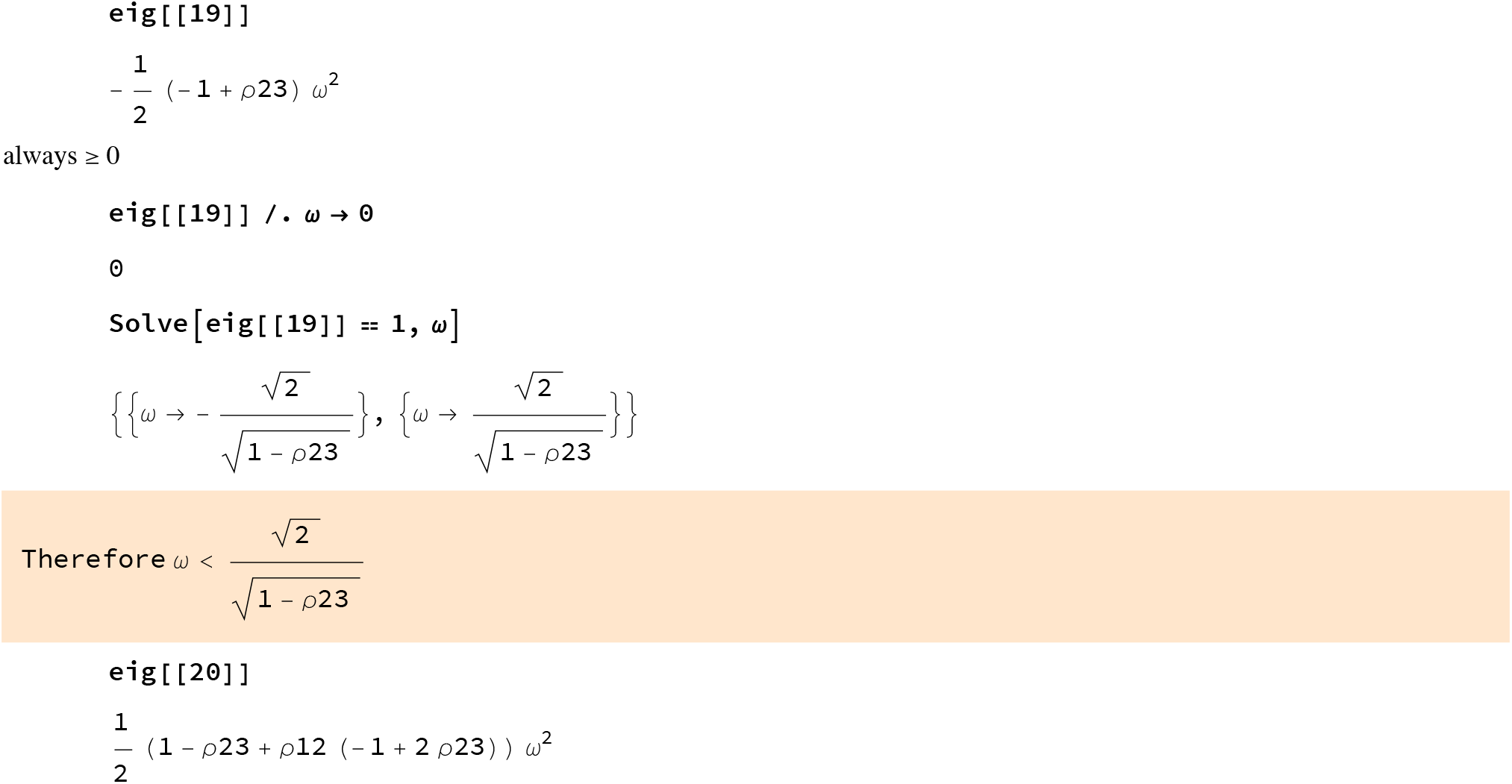
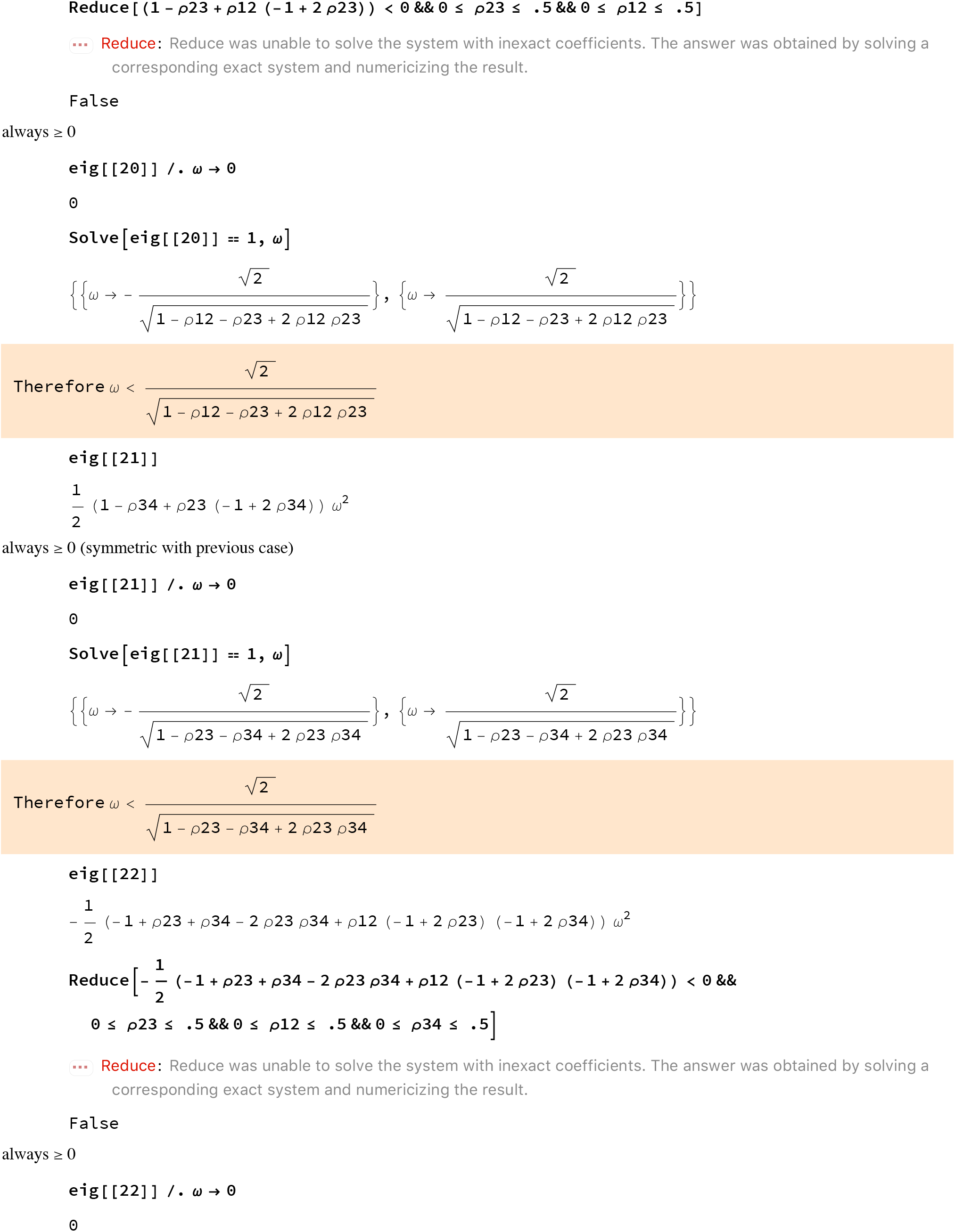
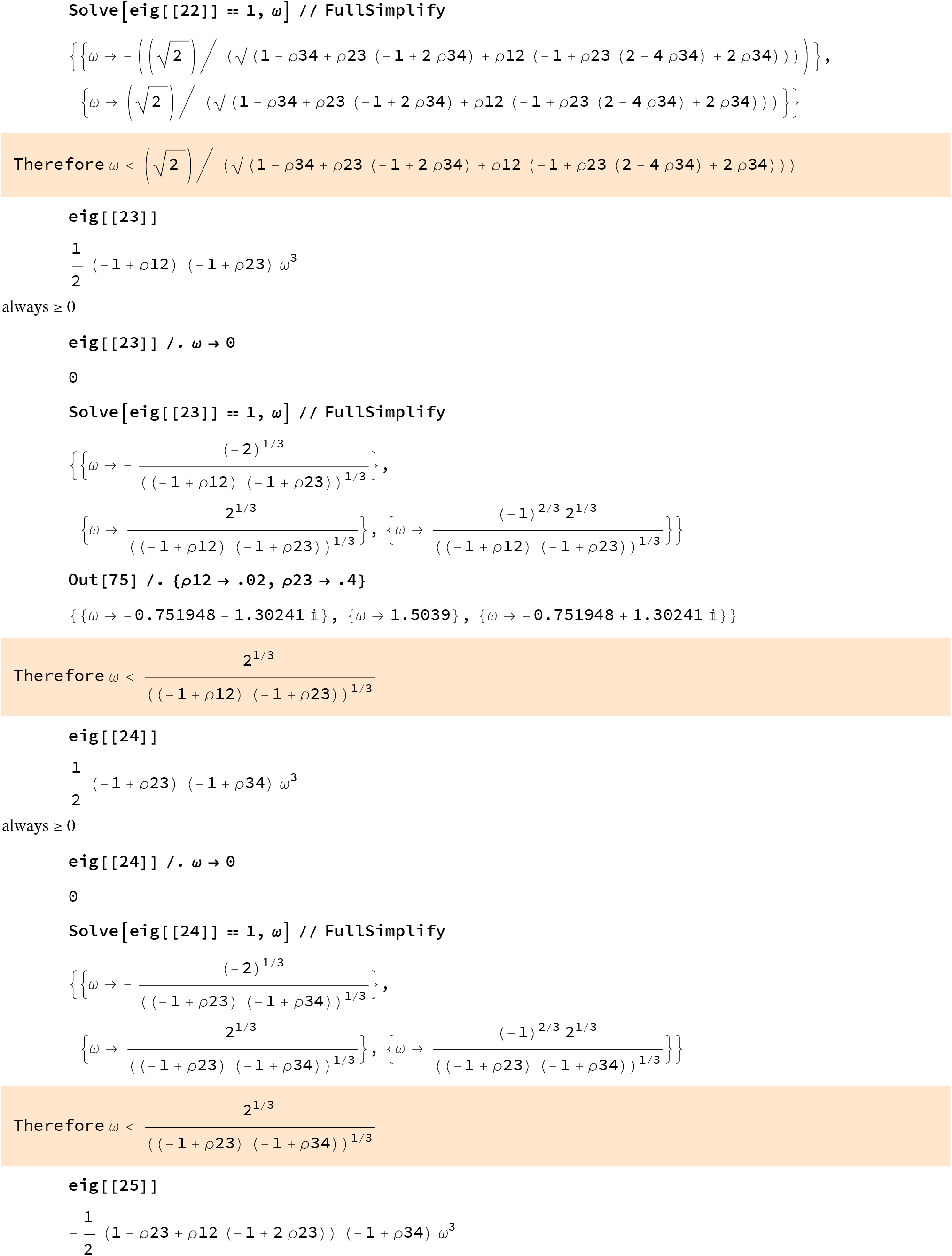
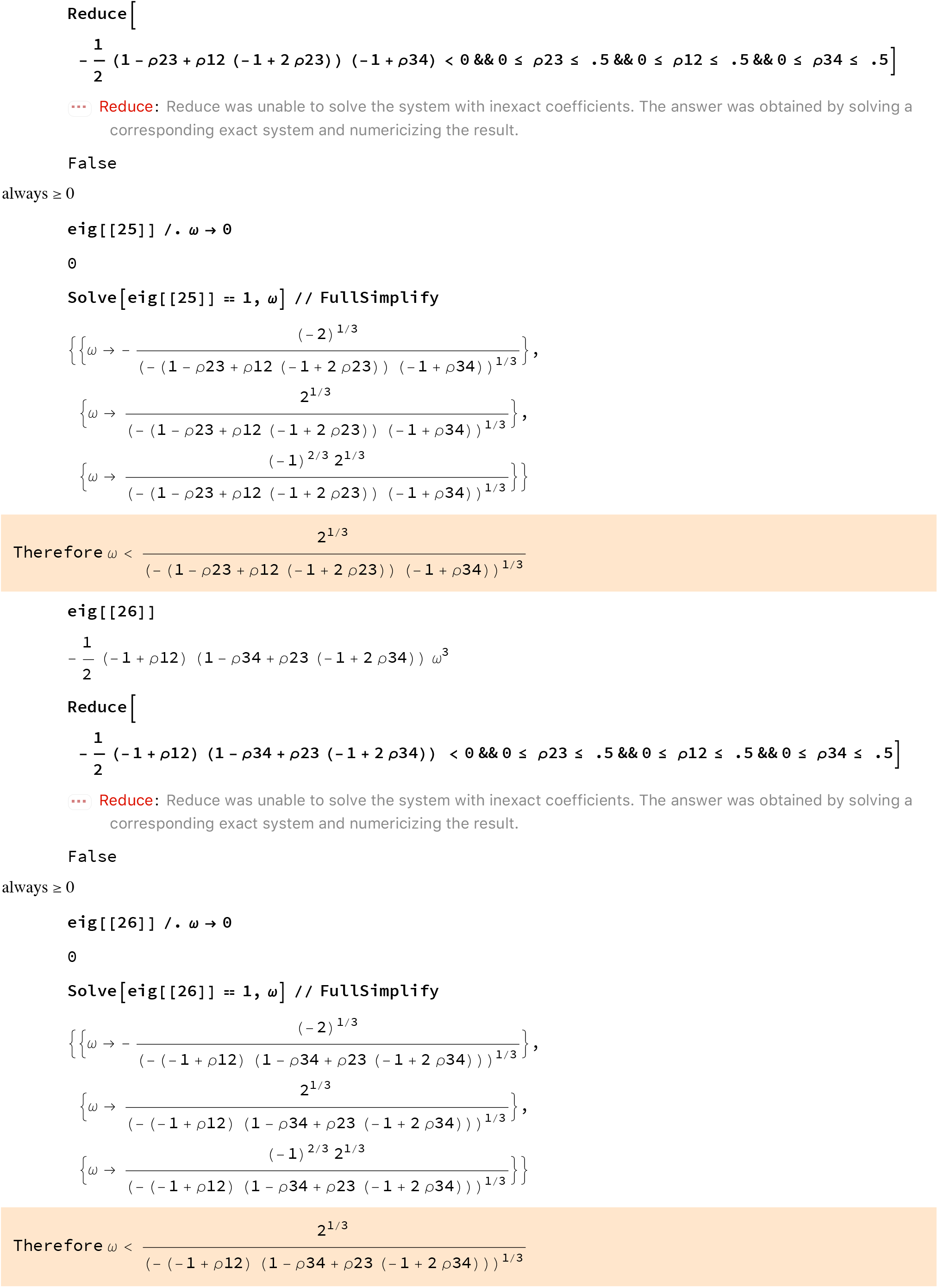
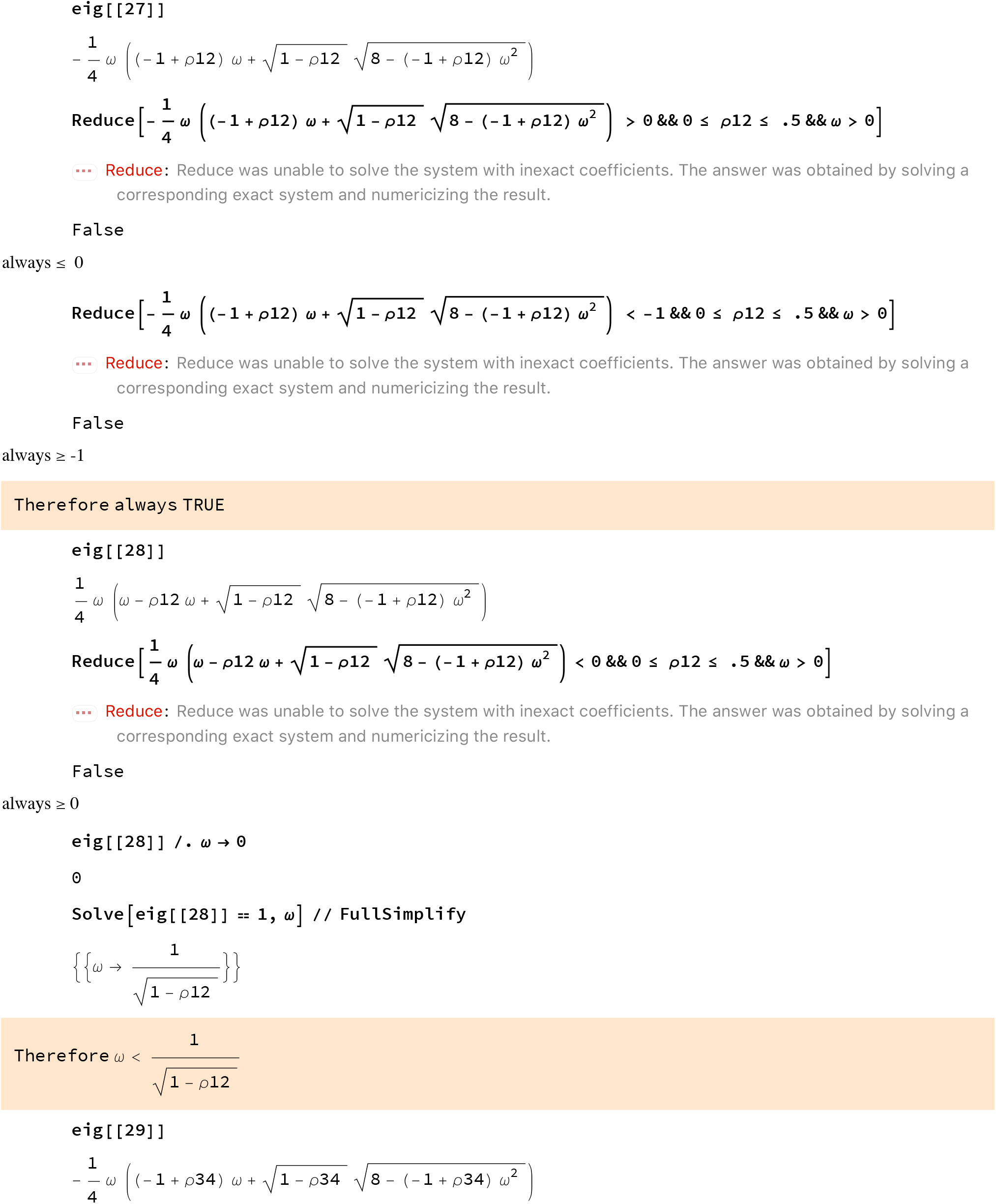

per symmetry with eig[[27]],≥ -1 and always <=0

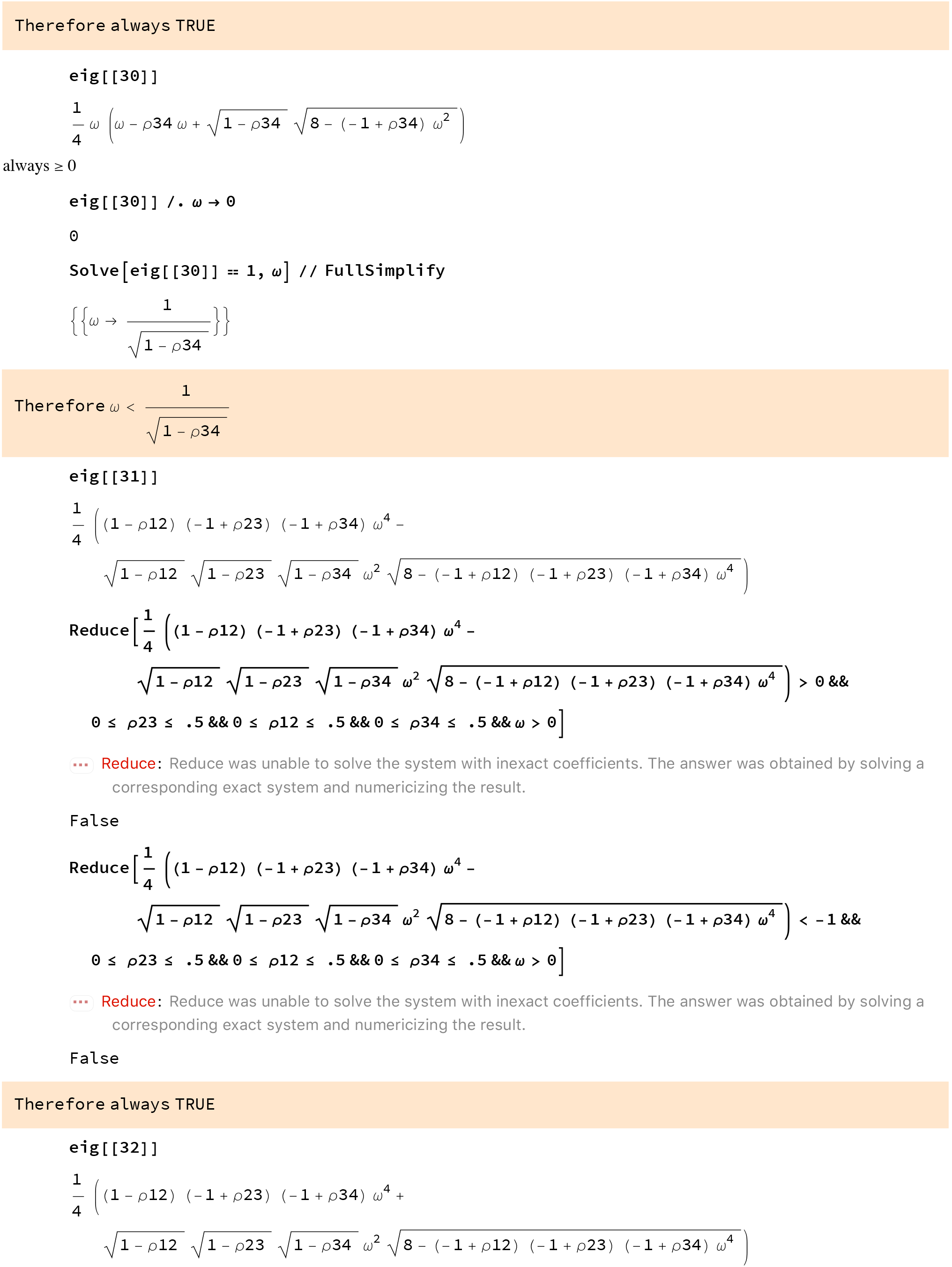
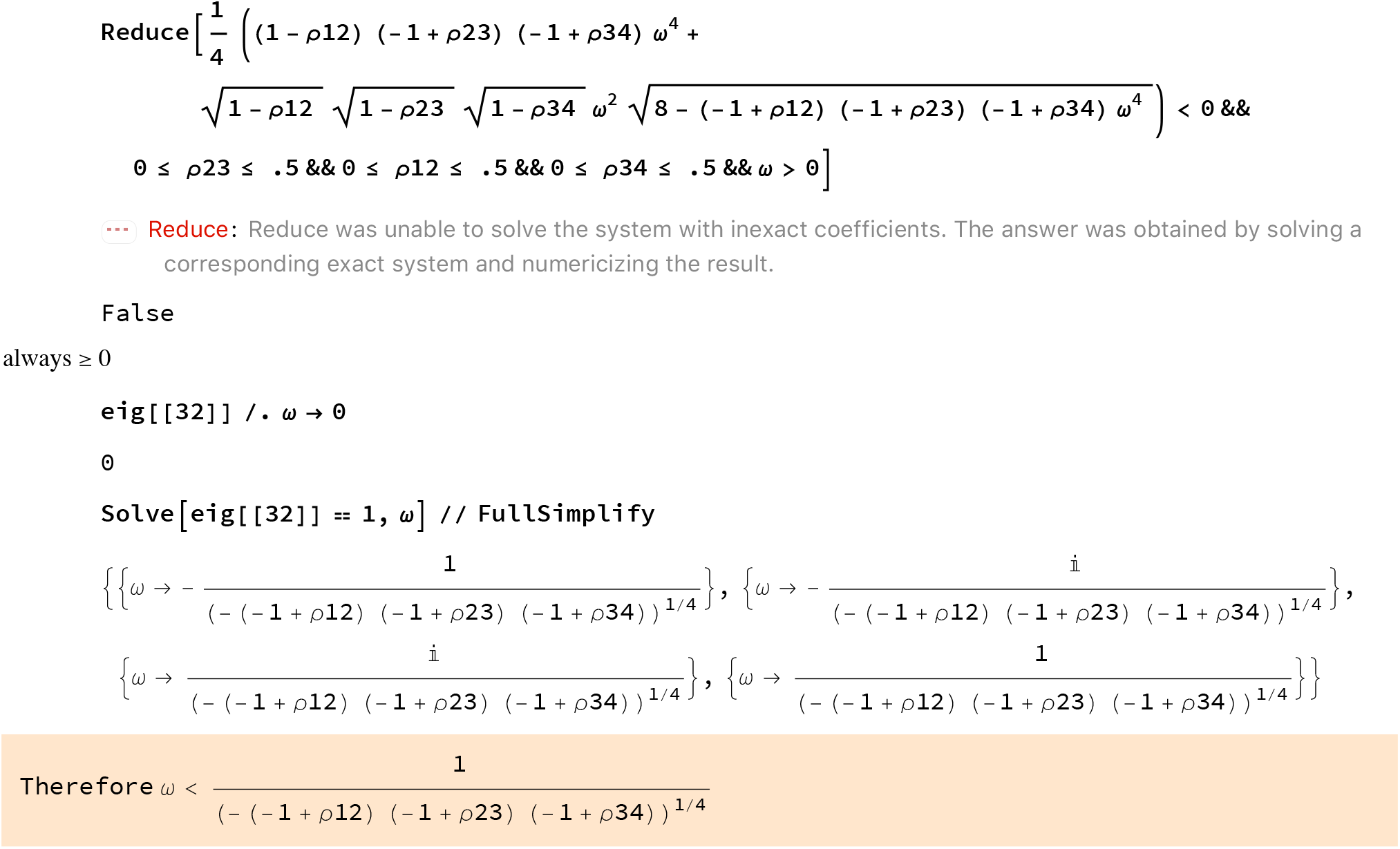

By combining each individual condition, we obtain:

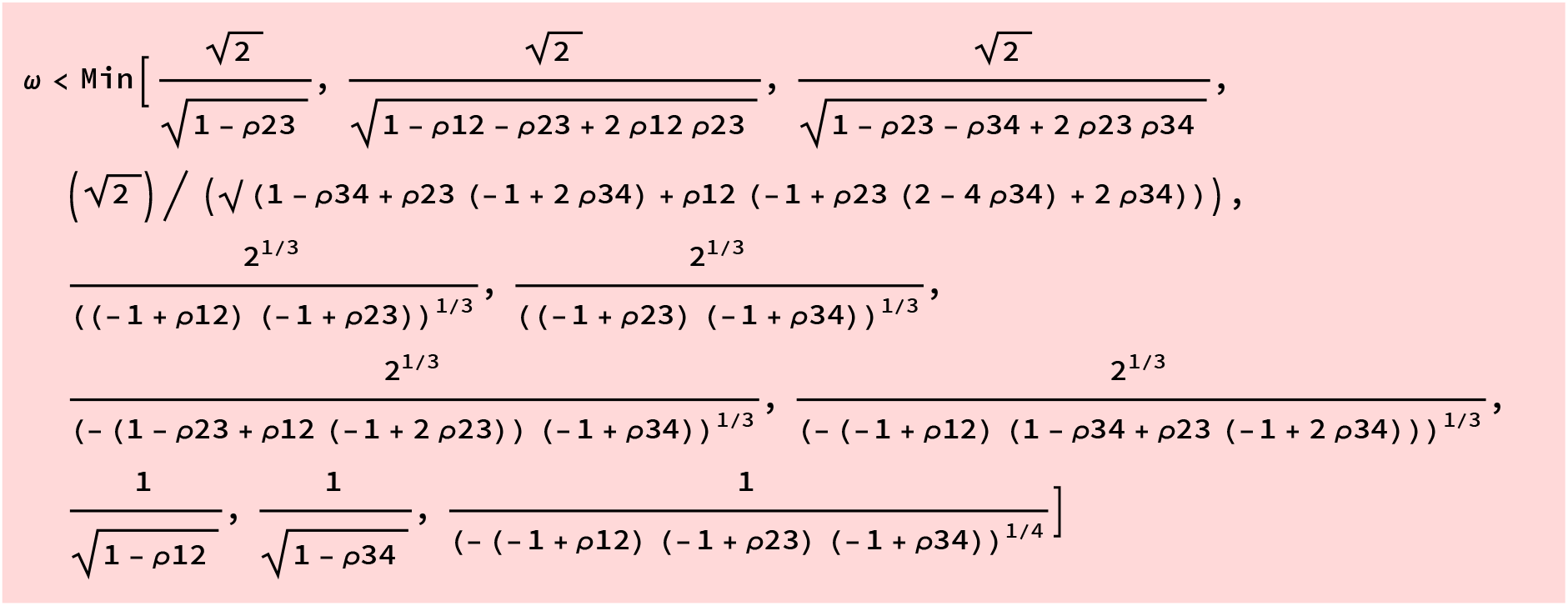
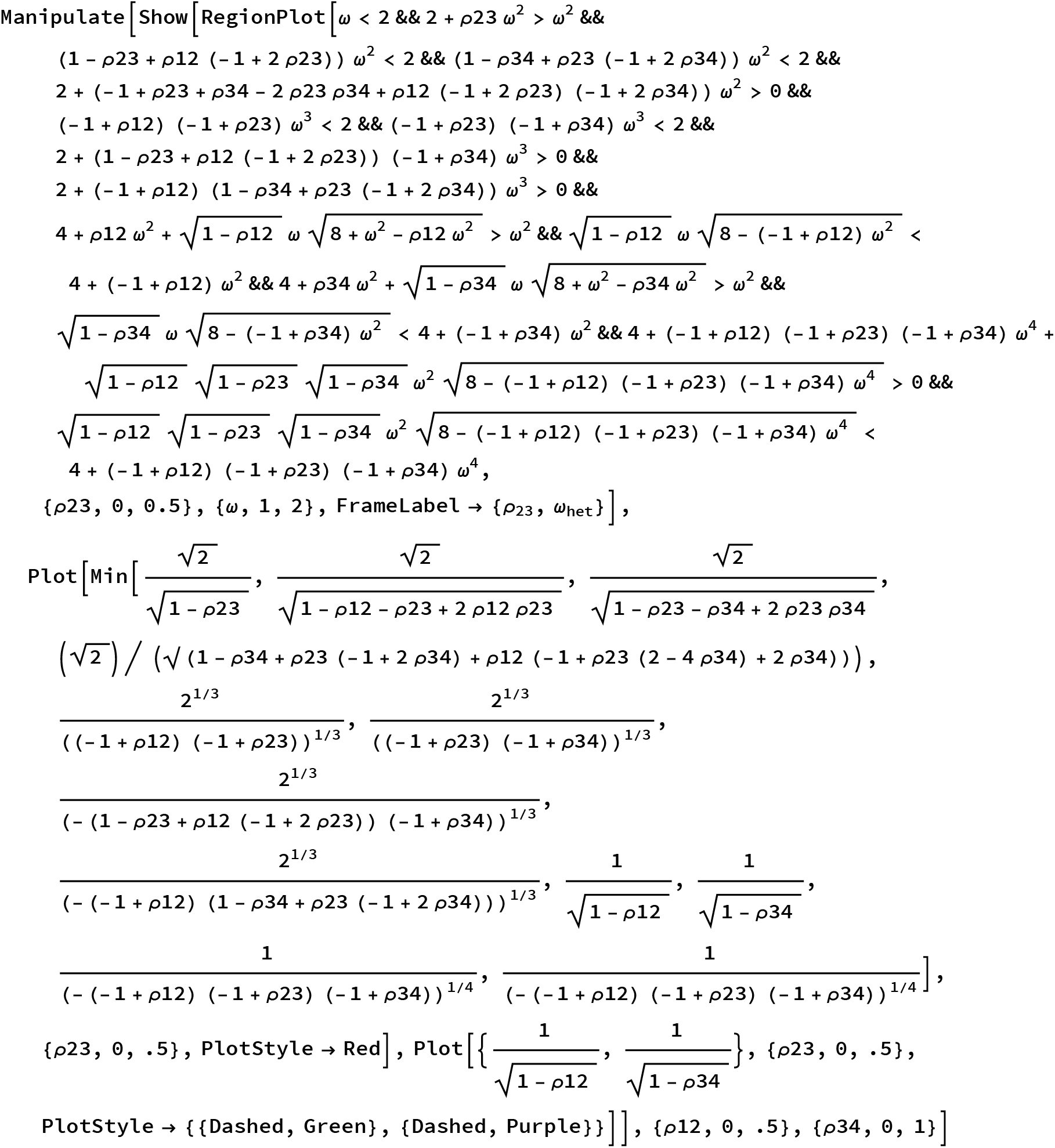
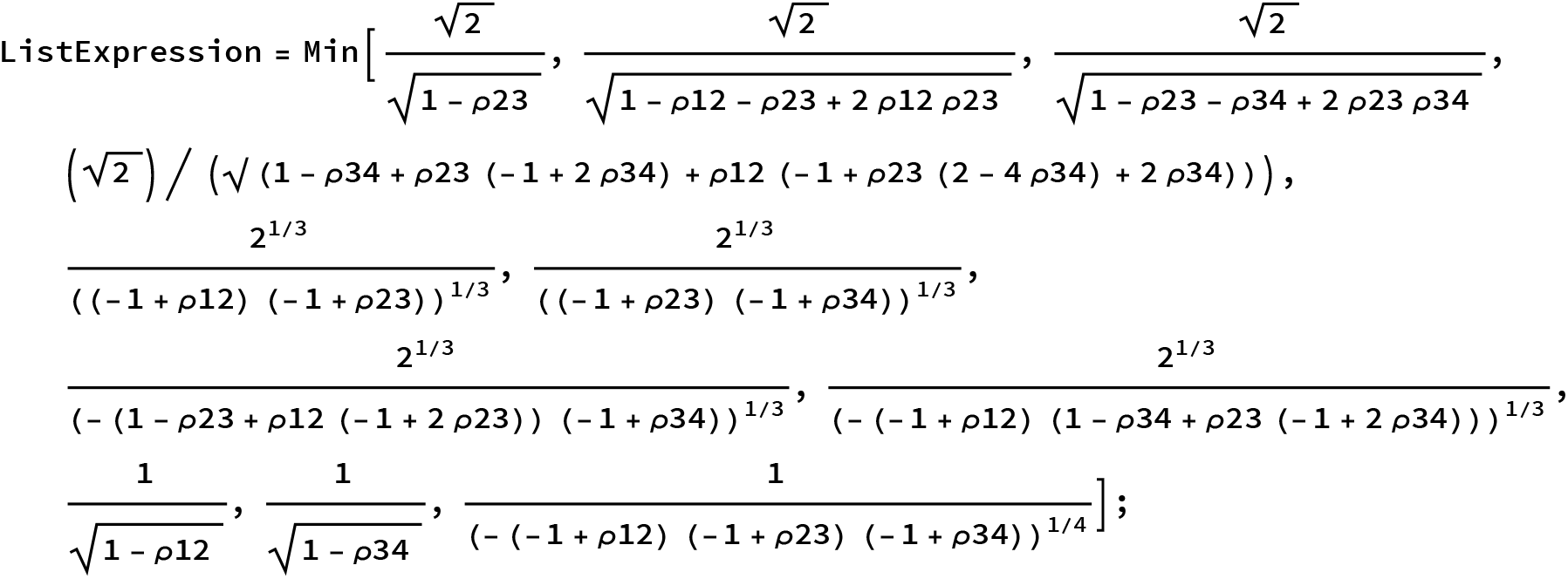
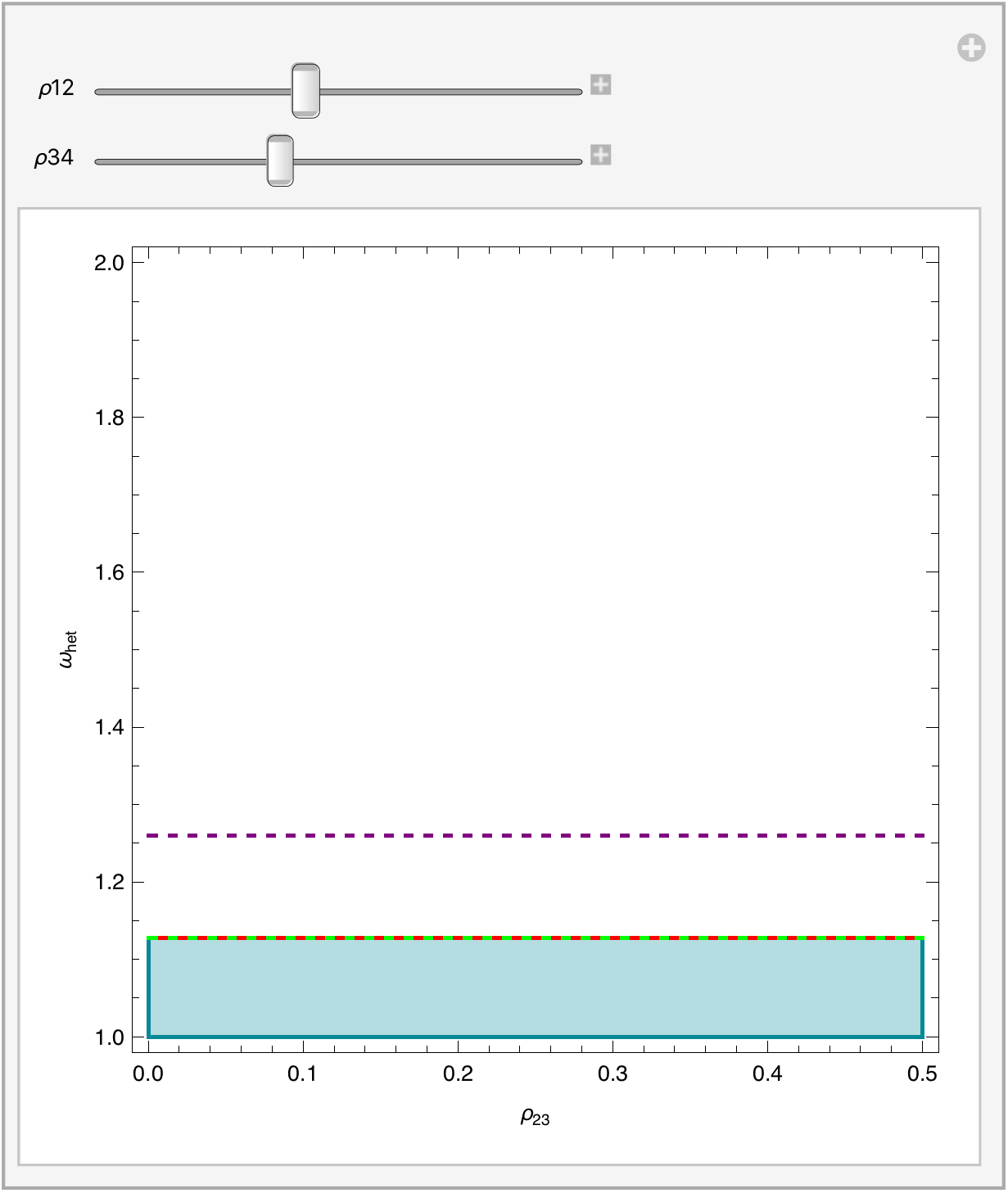

The previous expression can be reduce to two elements:

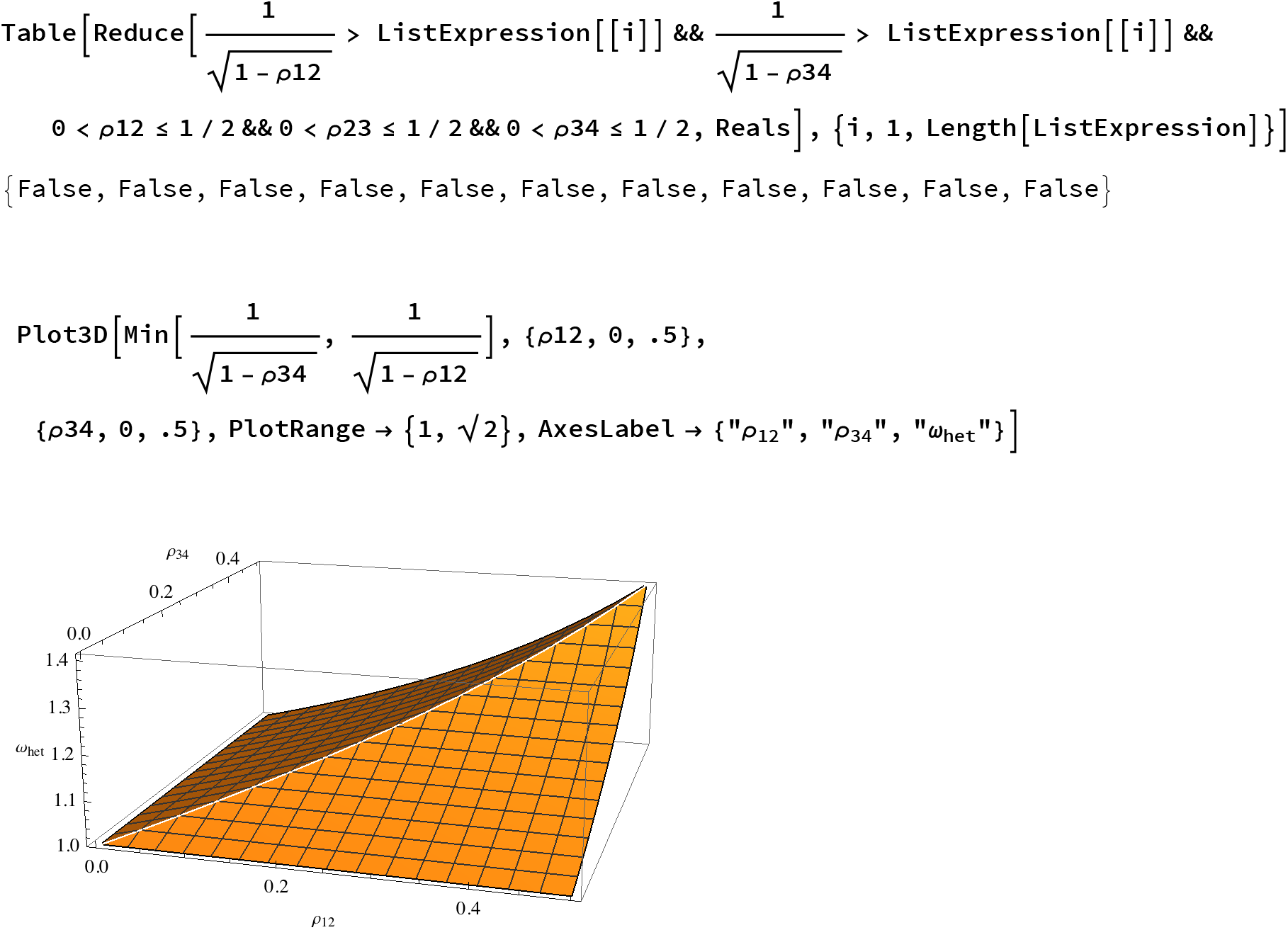

#### The network case: all loci interacts with each other (epistasis is here lethal)

Define the male fitness table

~~~
selectionMatrixMales4Lall ϵ = Table[0 +Boole.i1 == i2 == i3 == i4,
  {i1, 0, 1}, {i3, 0, 1}, {i4, 0, 1}] // Flatten
~~~

~~~
{1, 0, 0, 0, 0, 0, 0, 0, 0, 0, 0, 0, 0, 0, 0, 1}
~~~

Define the female fitness table

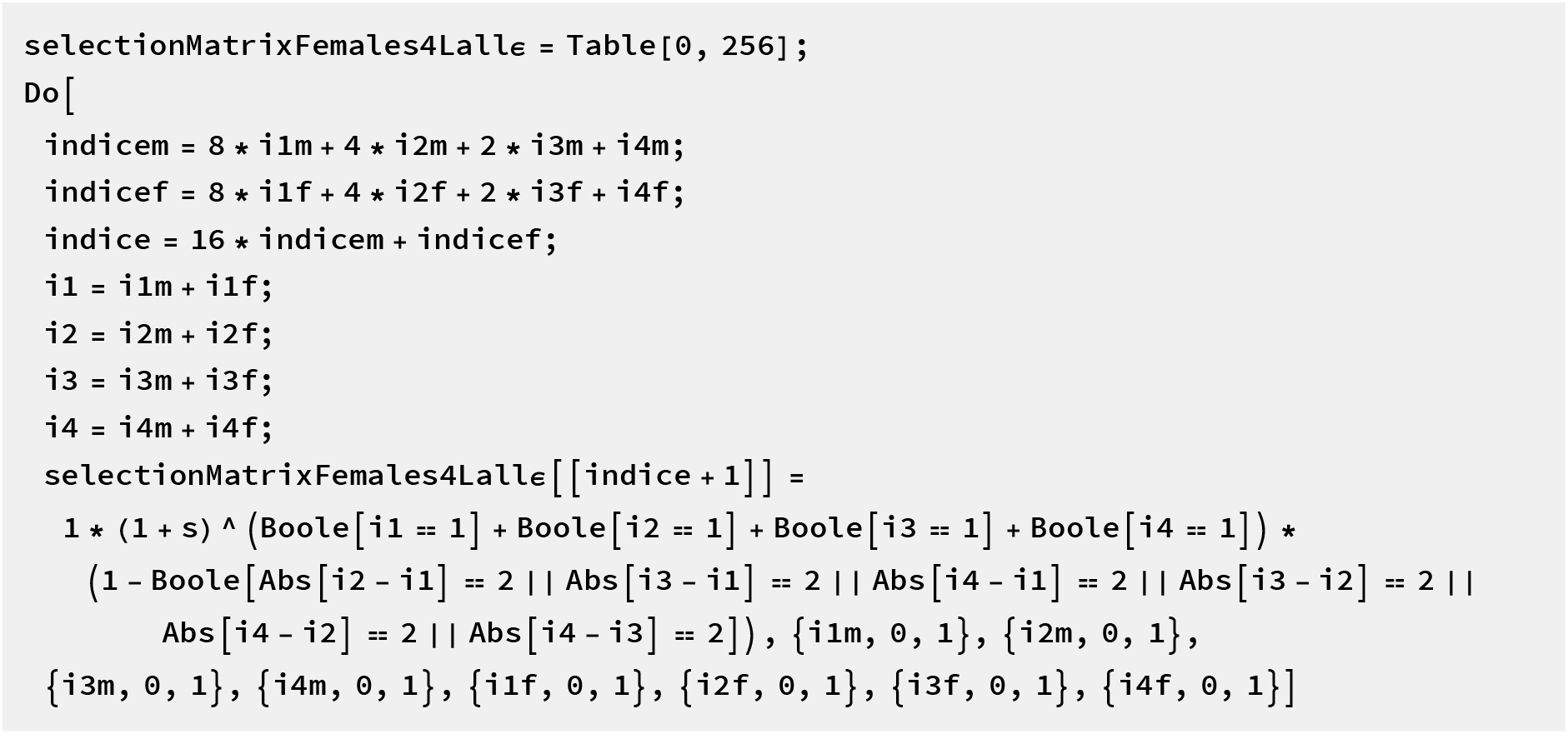
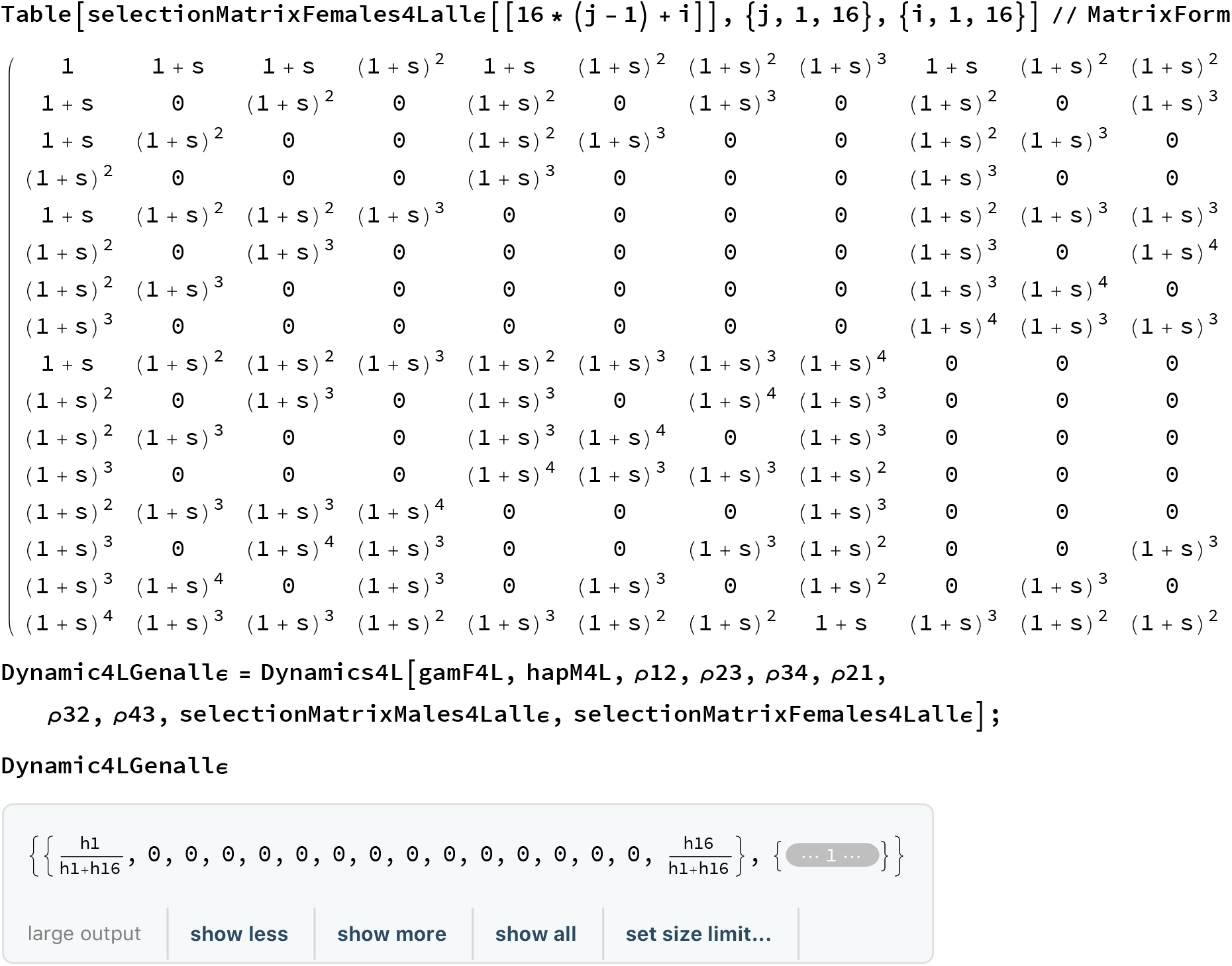

Define the Jacobian matrix

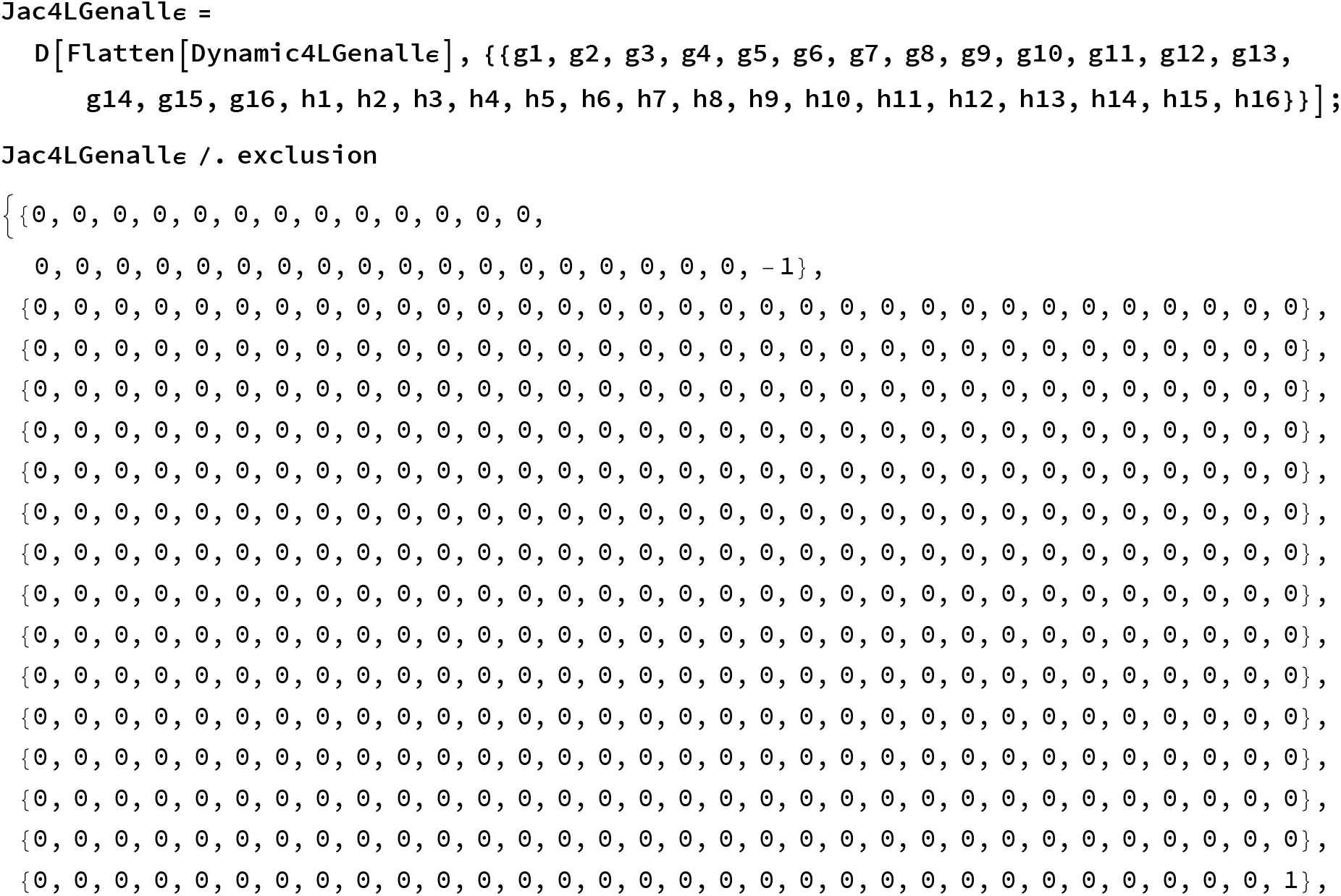
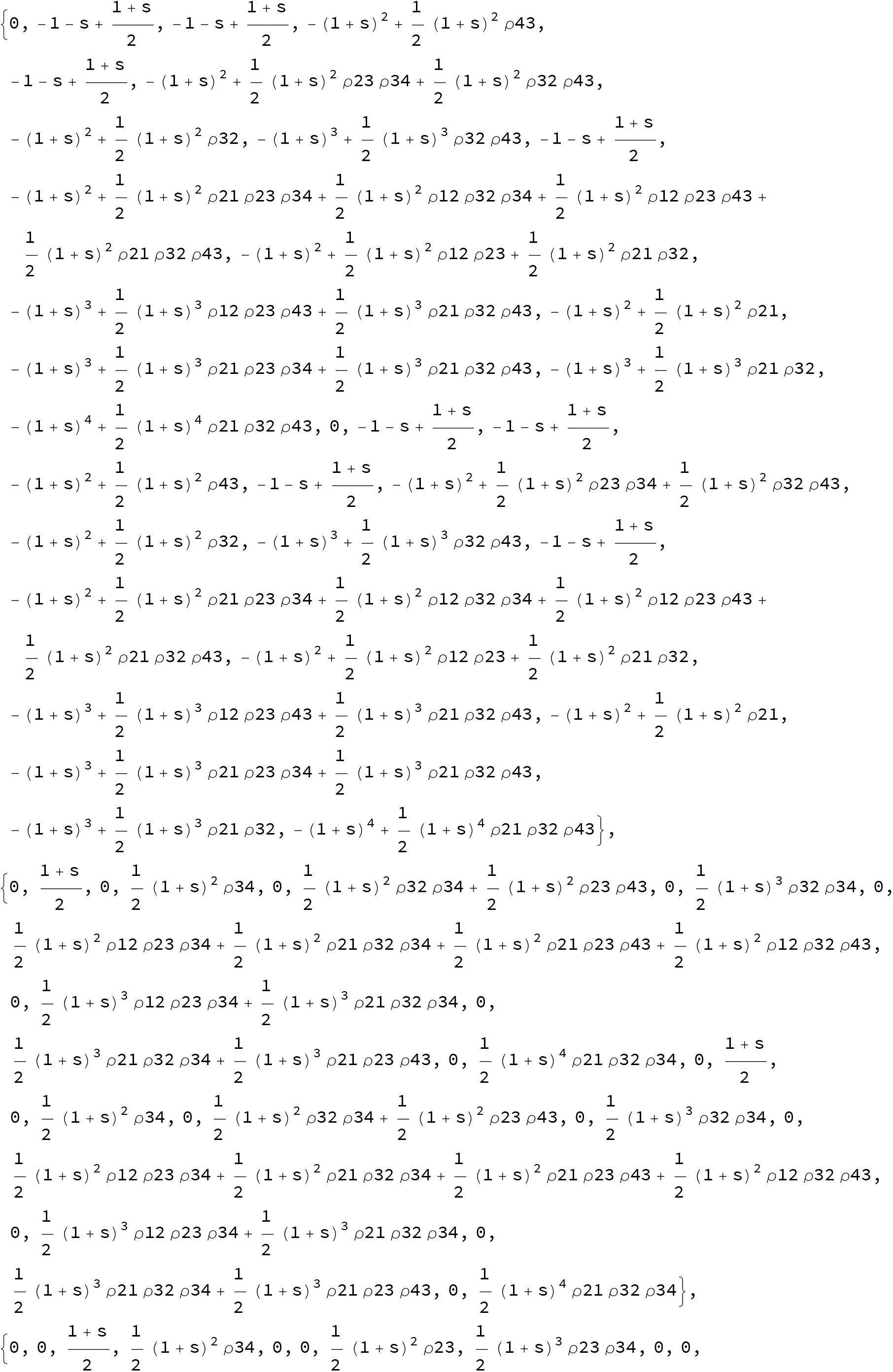
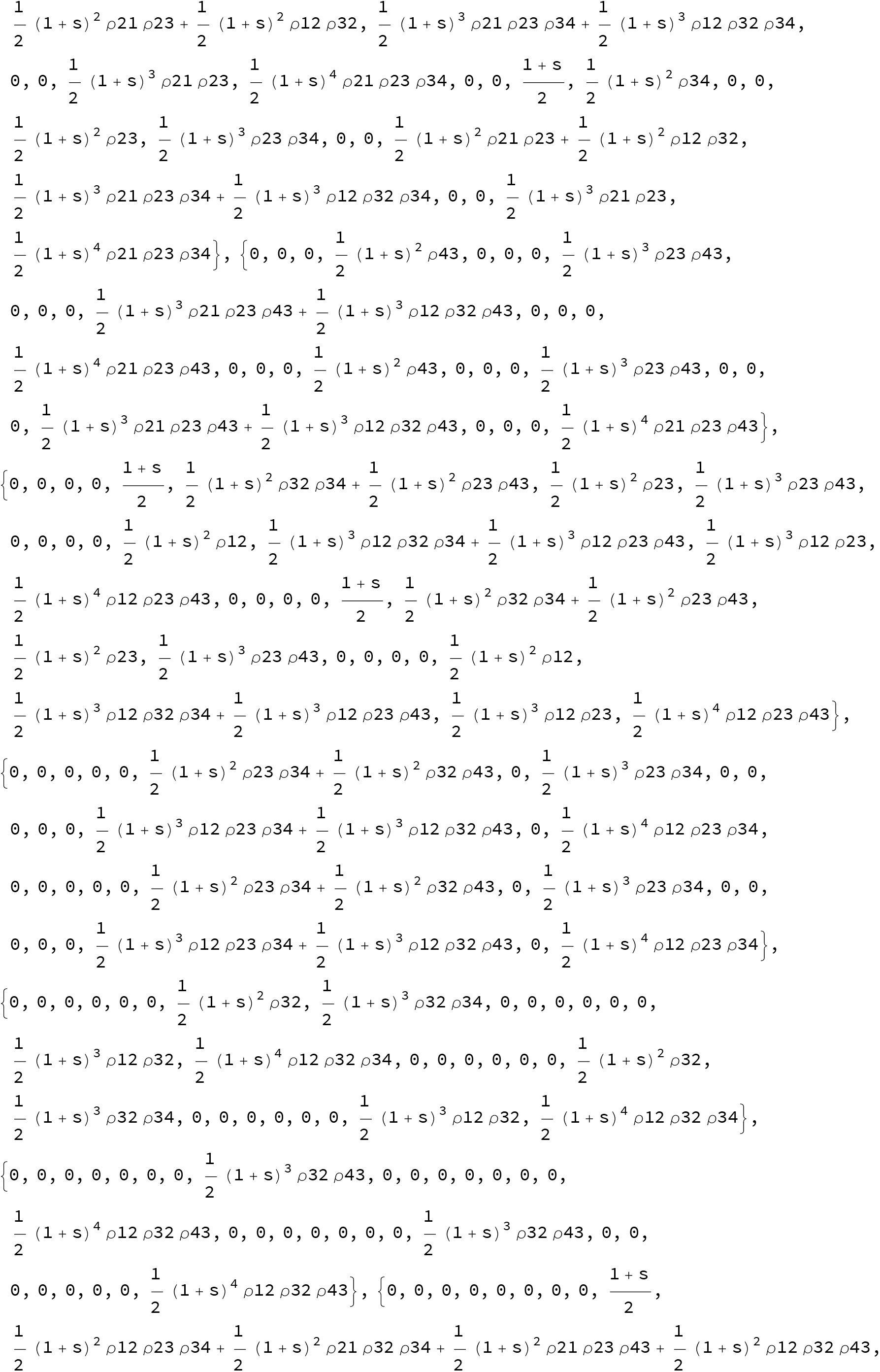
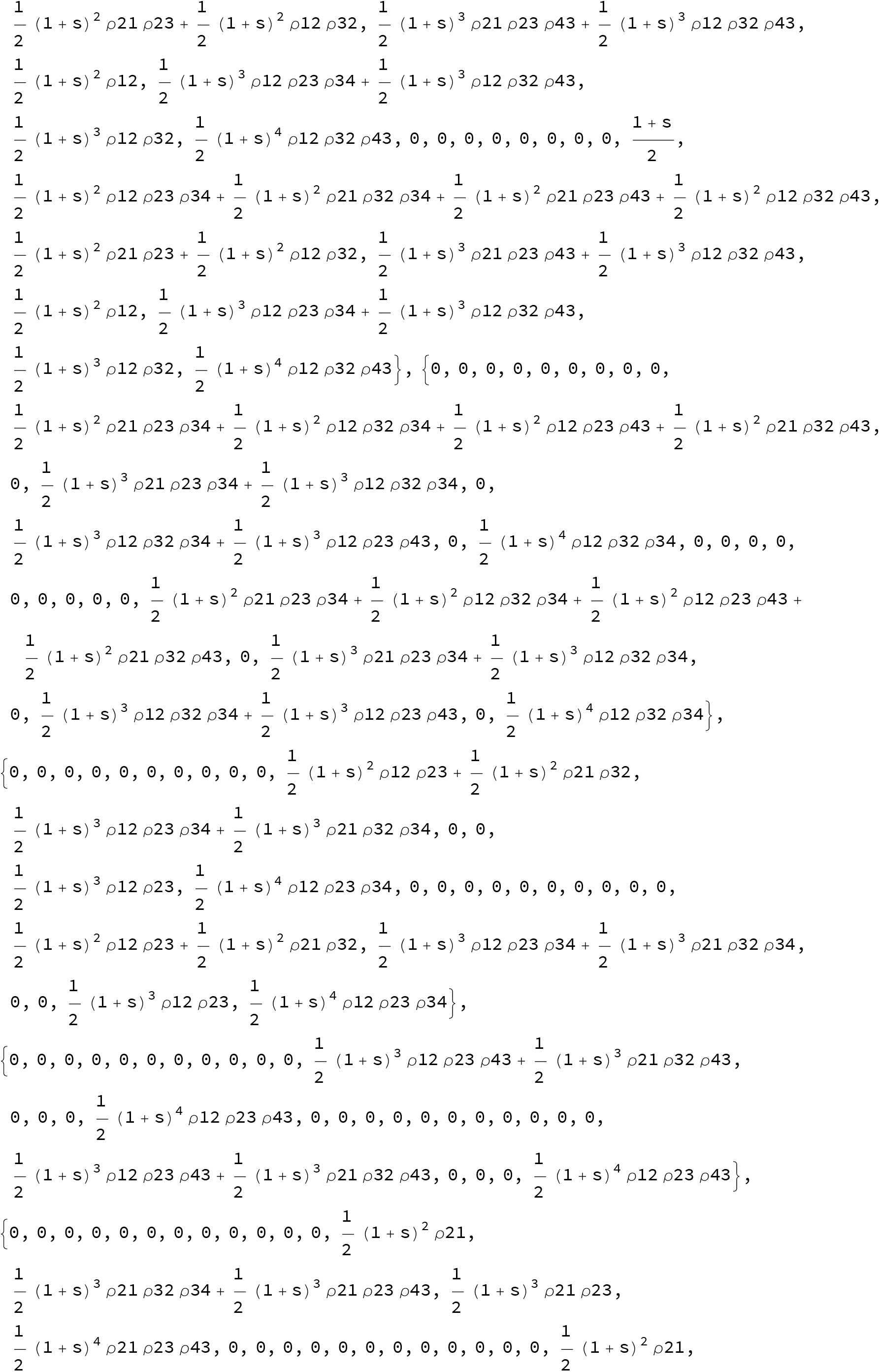
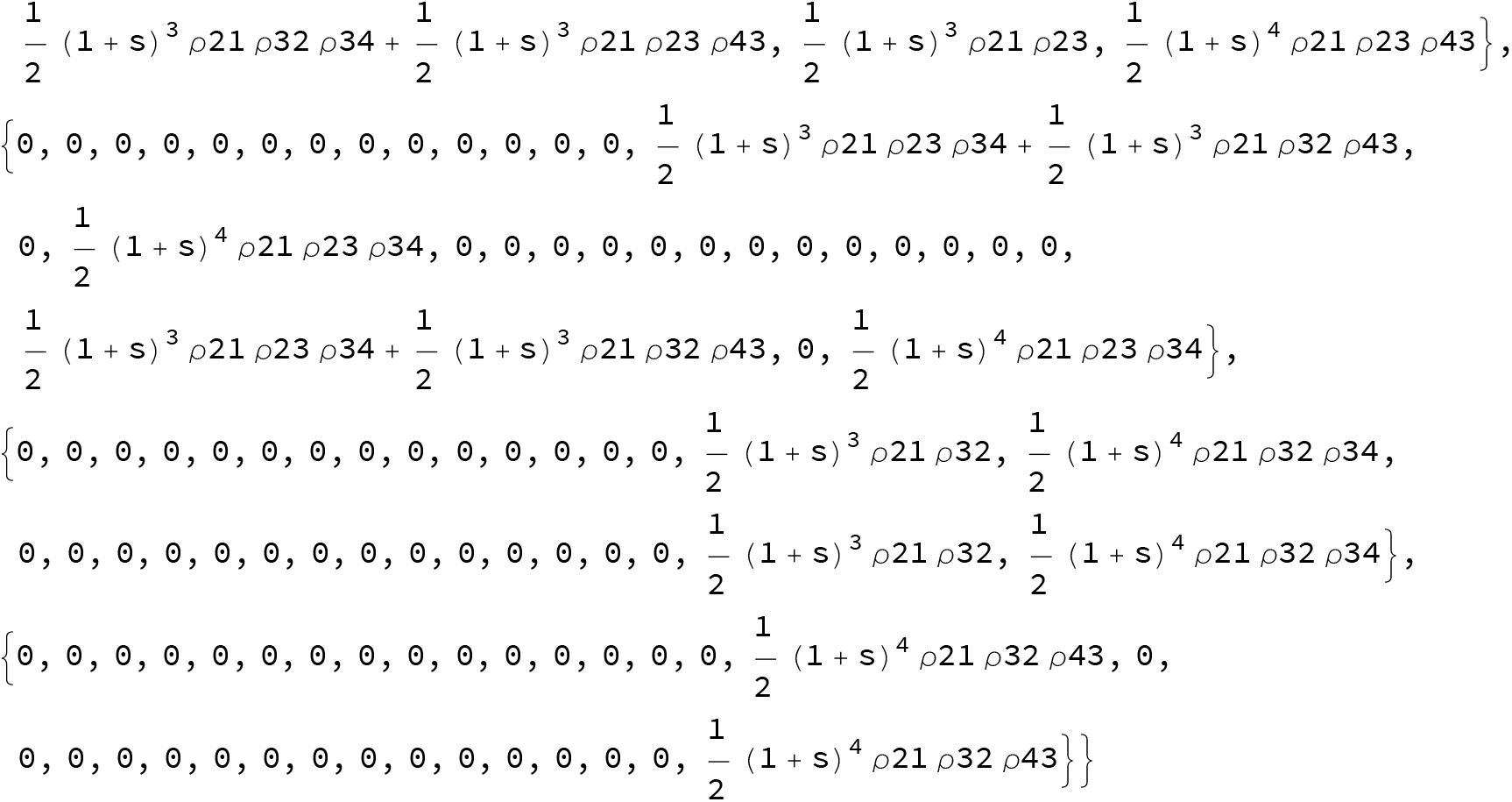

Define the Eigenvalues, with using again *ρ*21=1-*ρ*12 to avoid unwanted simplification by Mathematica

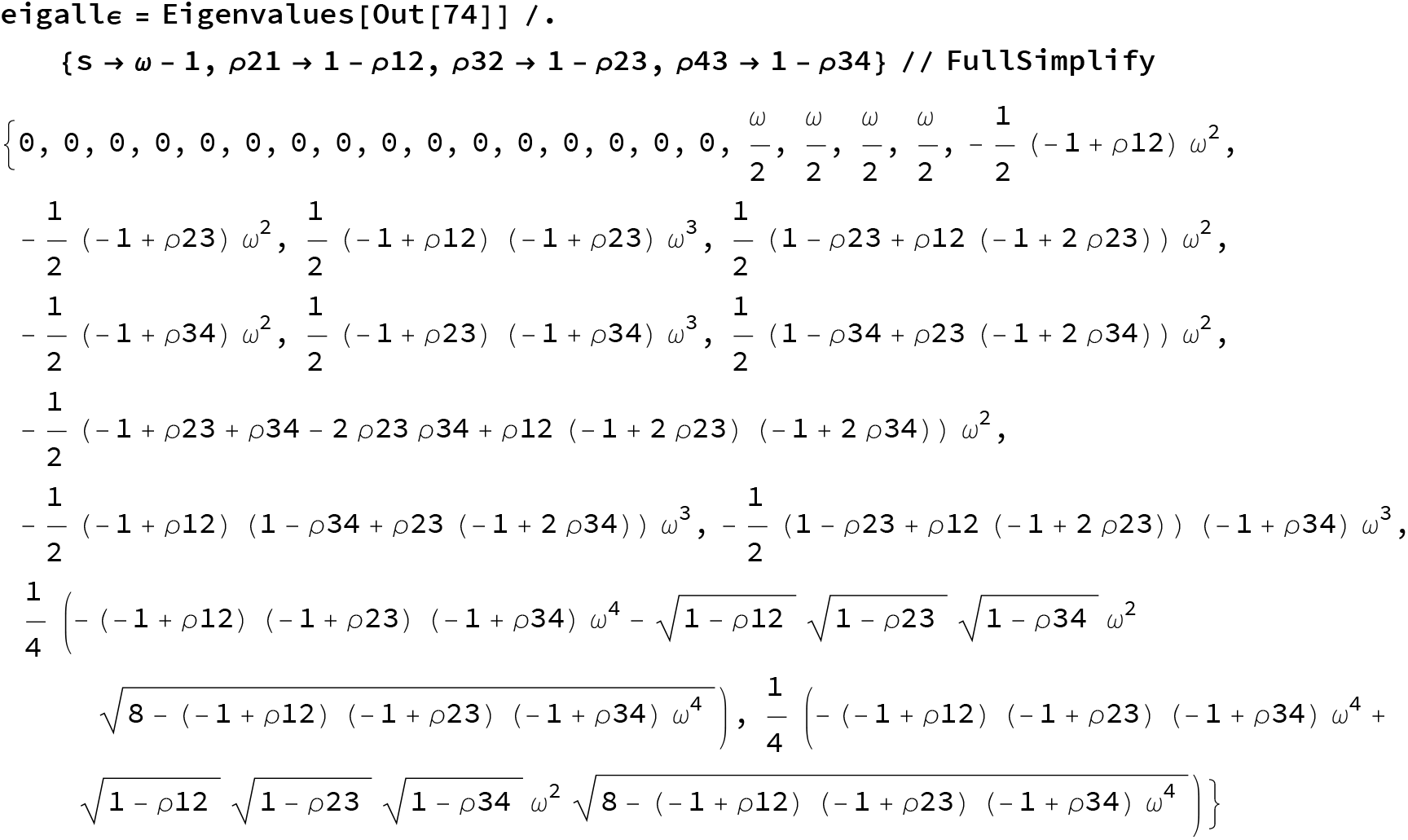
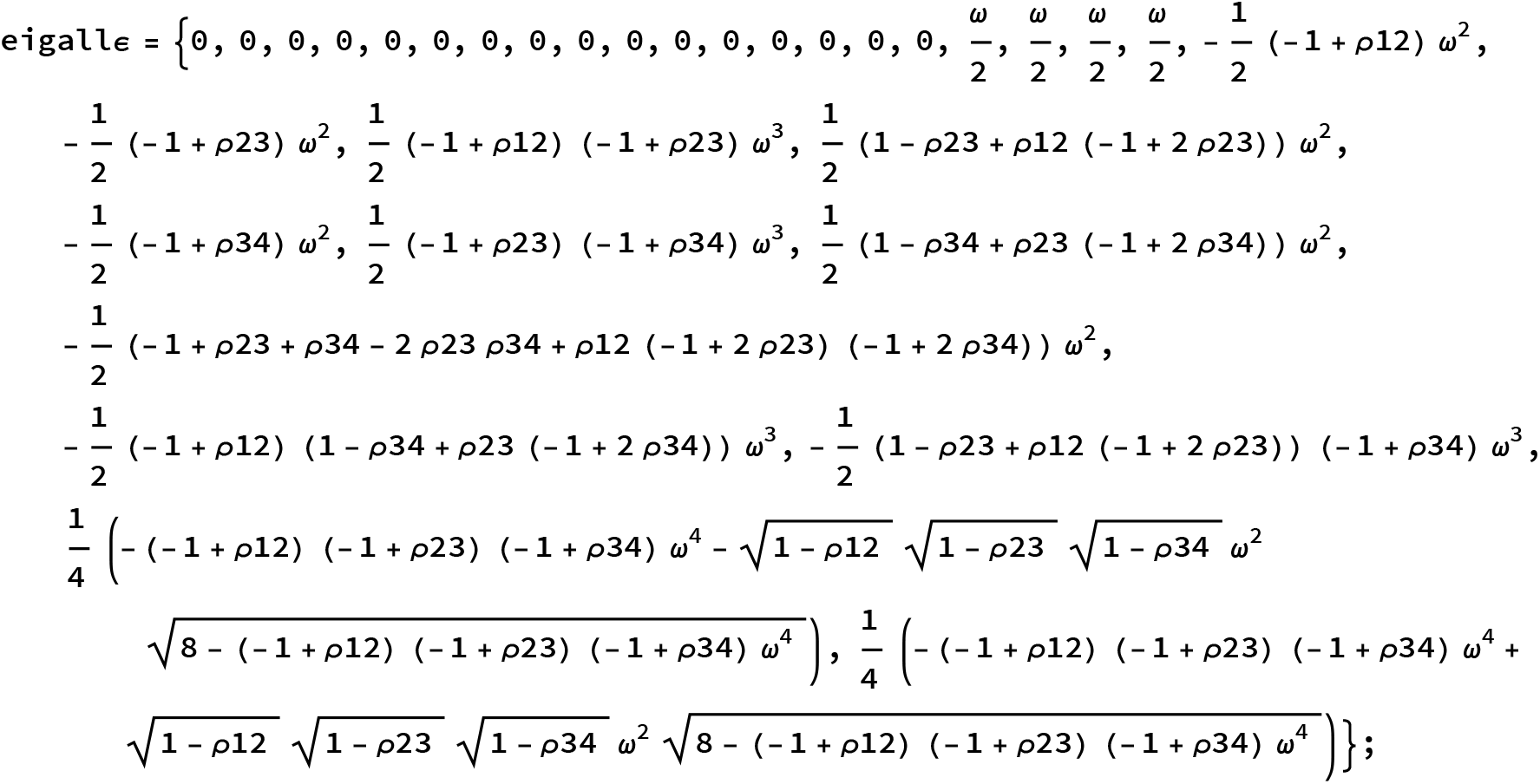

Case 1 Equidistant loci

For free recombination

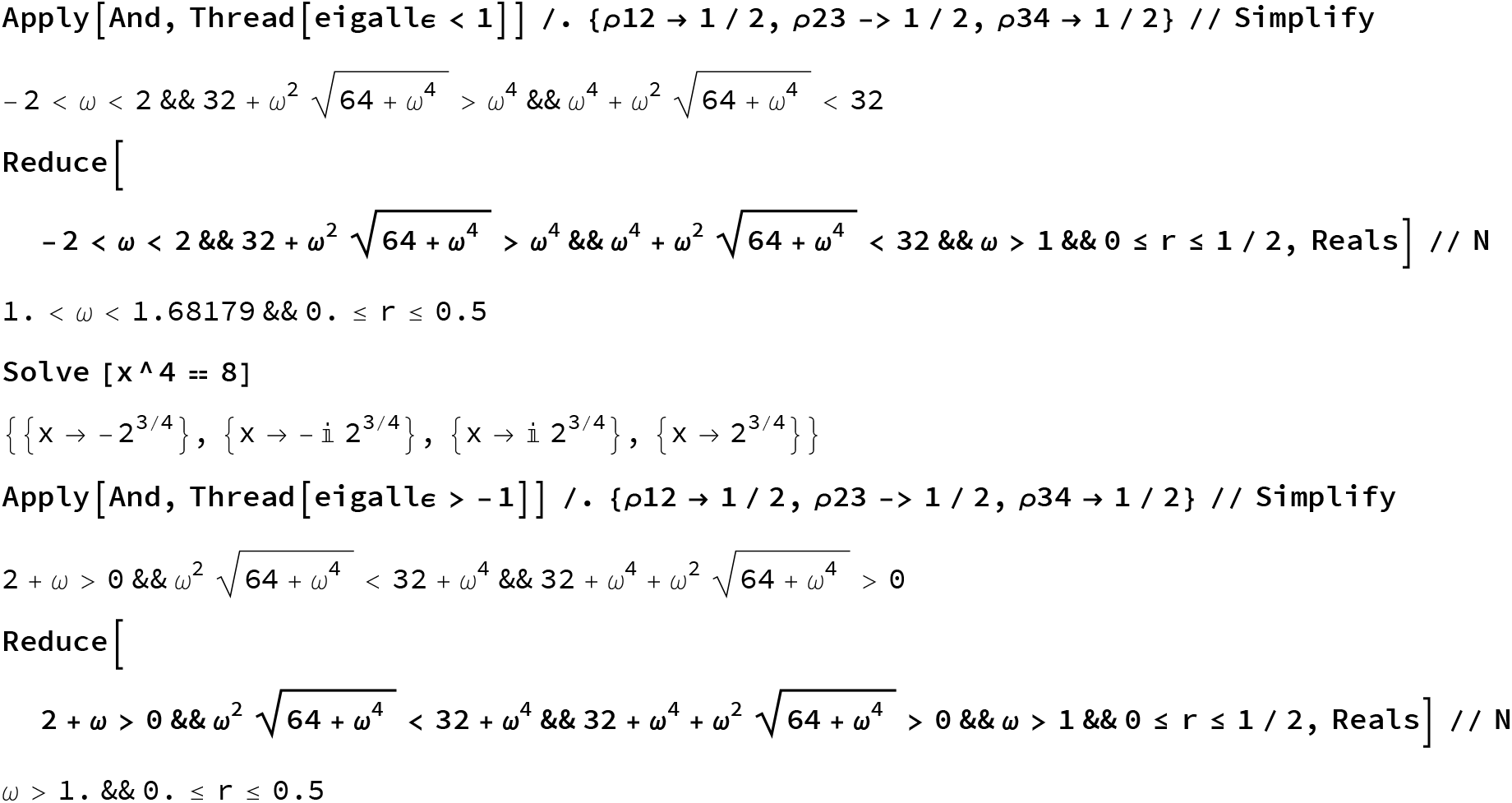

For arbitrary recombination

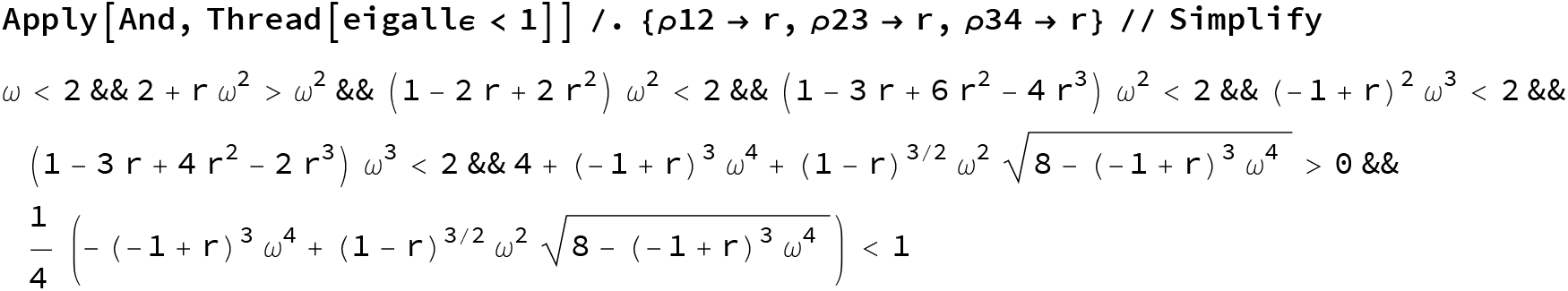
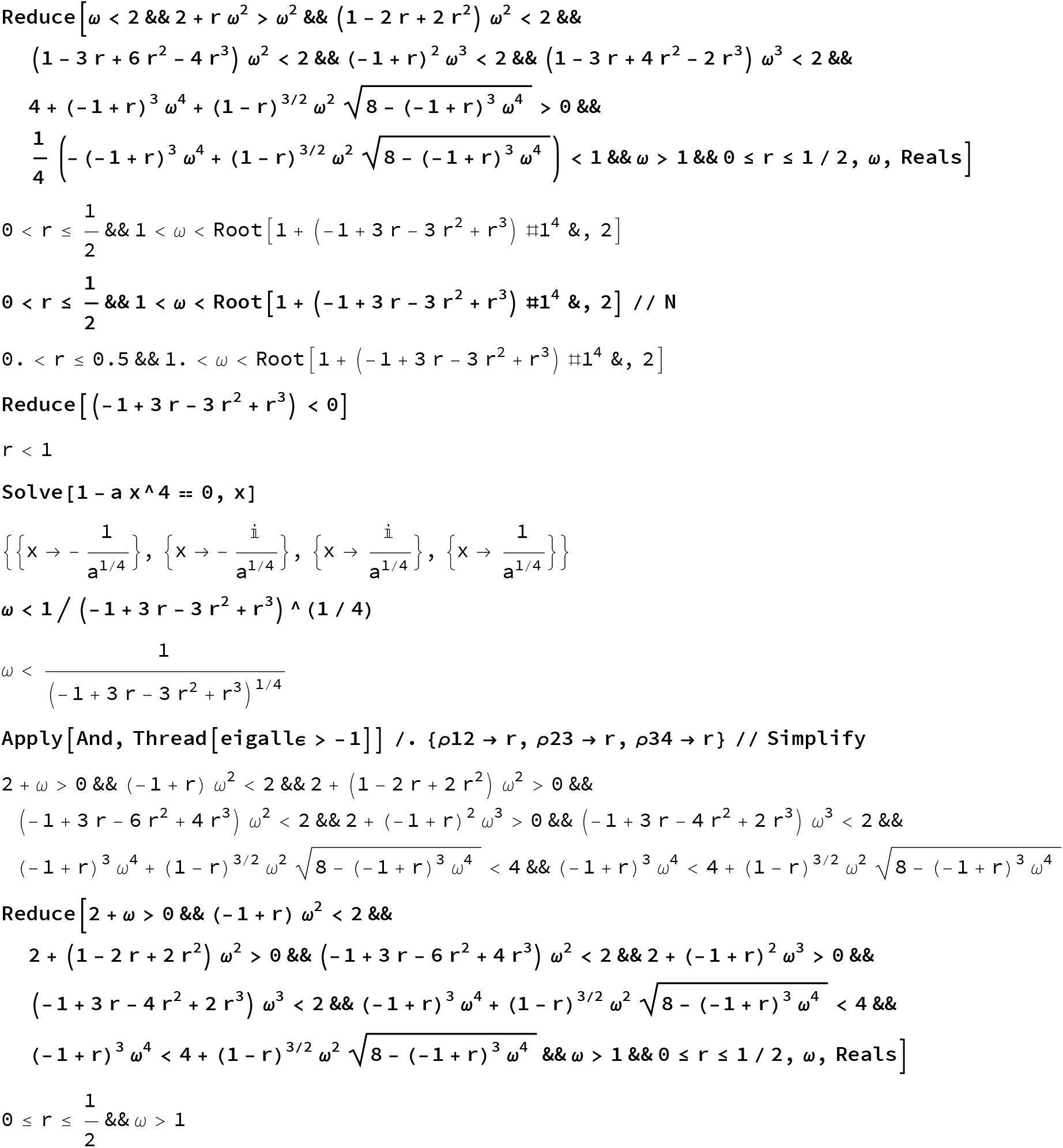
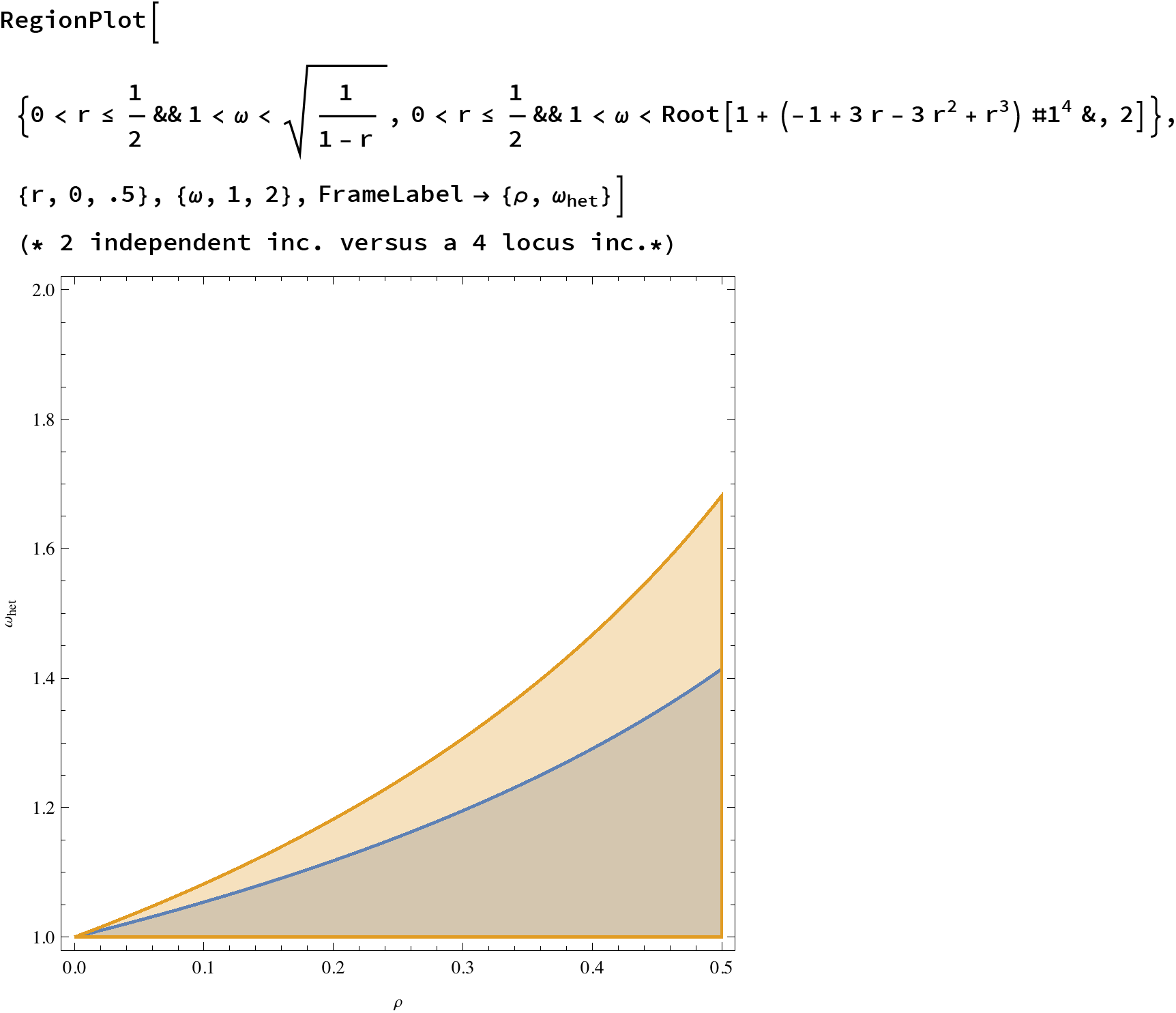
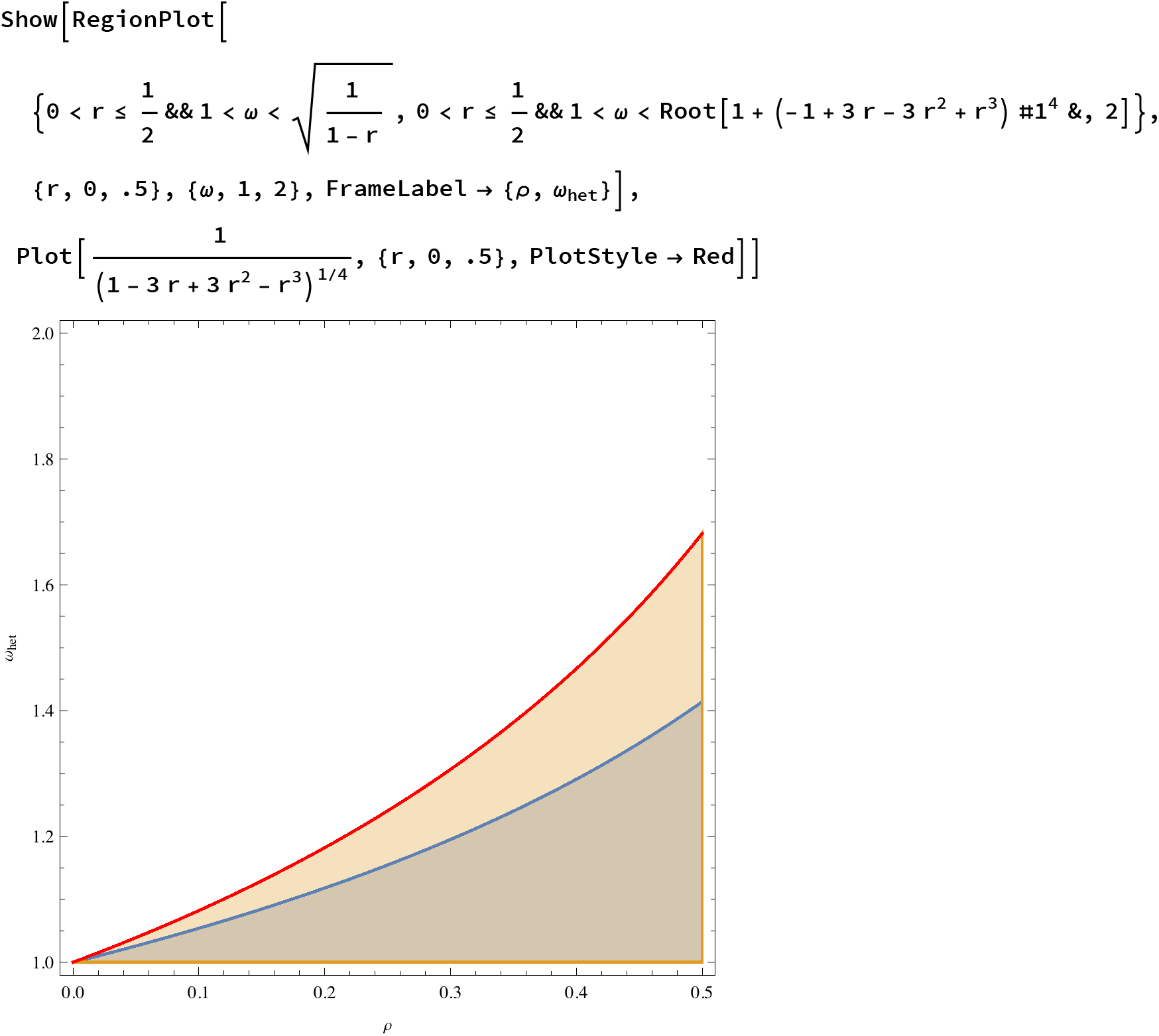

Case 2: 2 Loci per choromosome

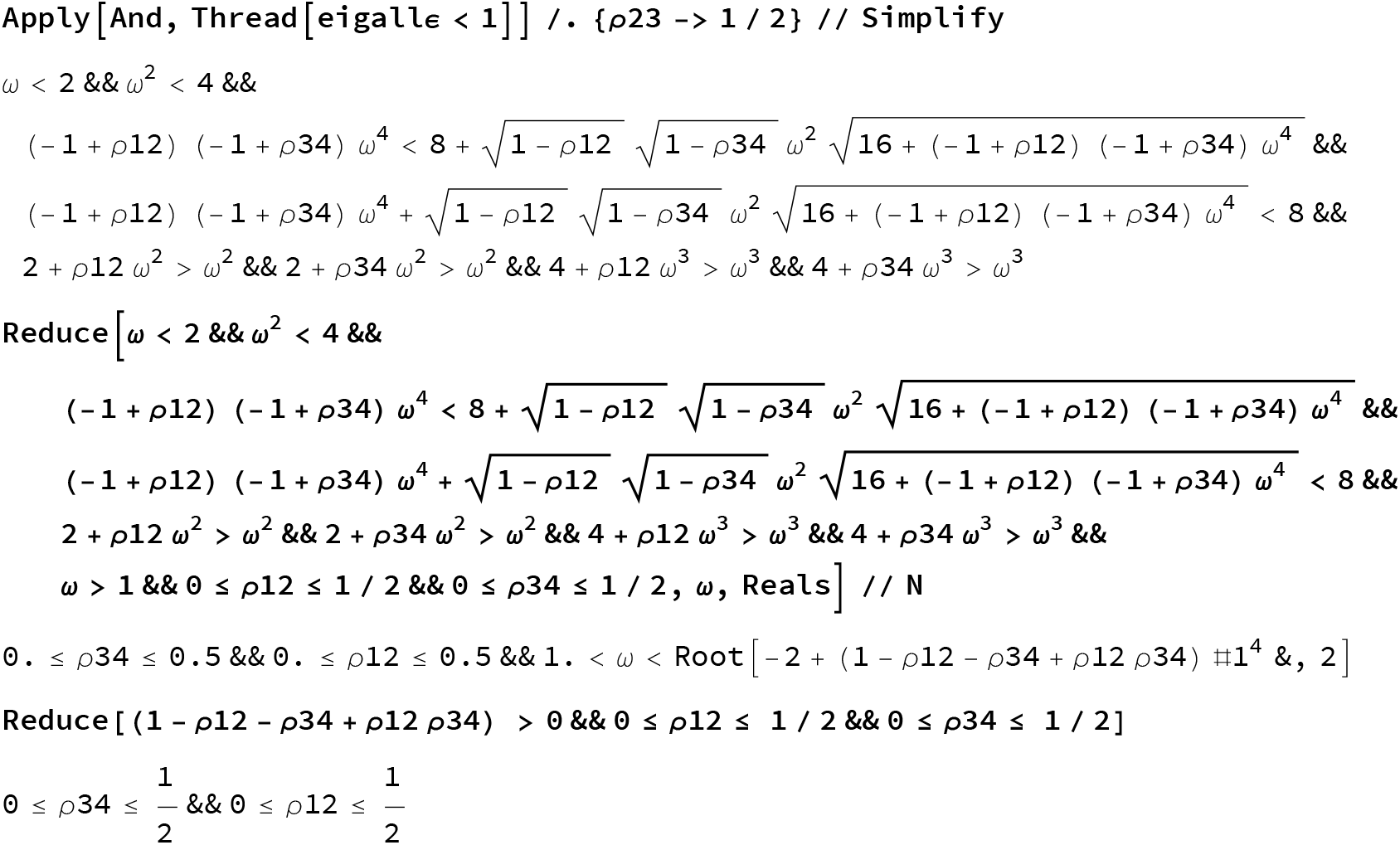
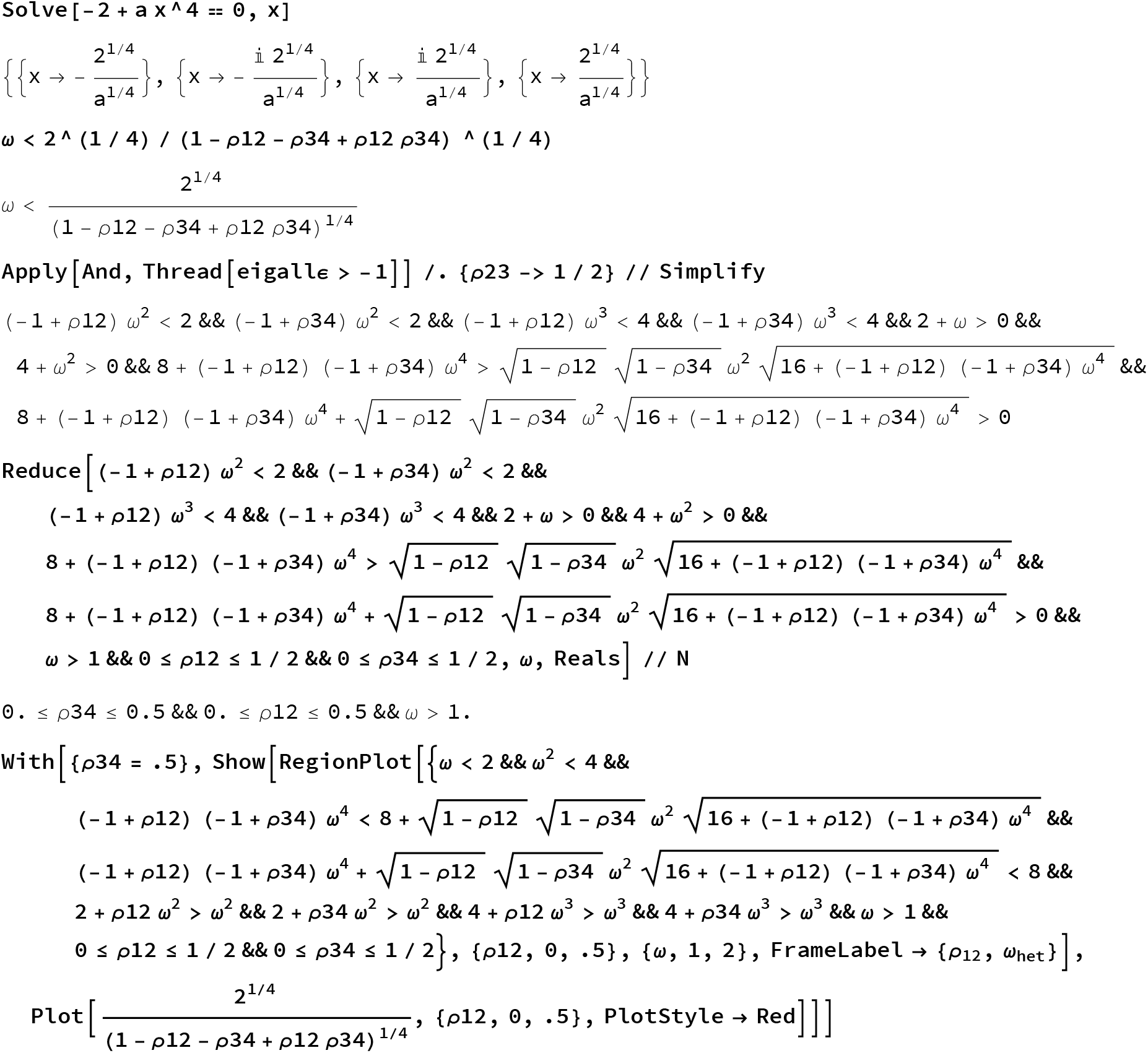
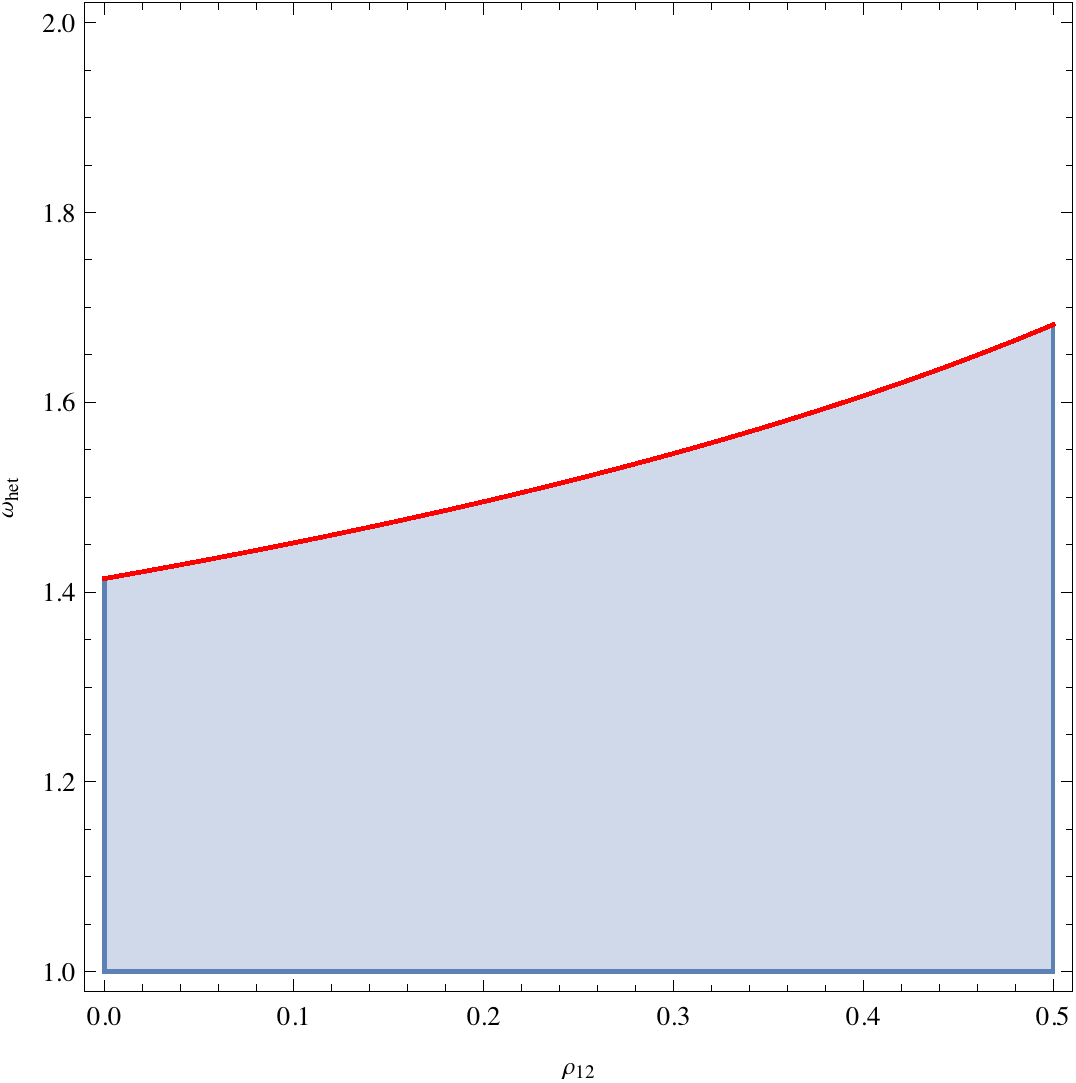

Case 3: Arbitrary recombination rate

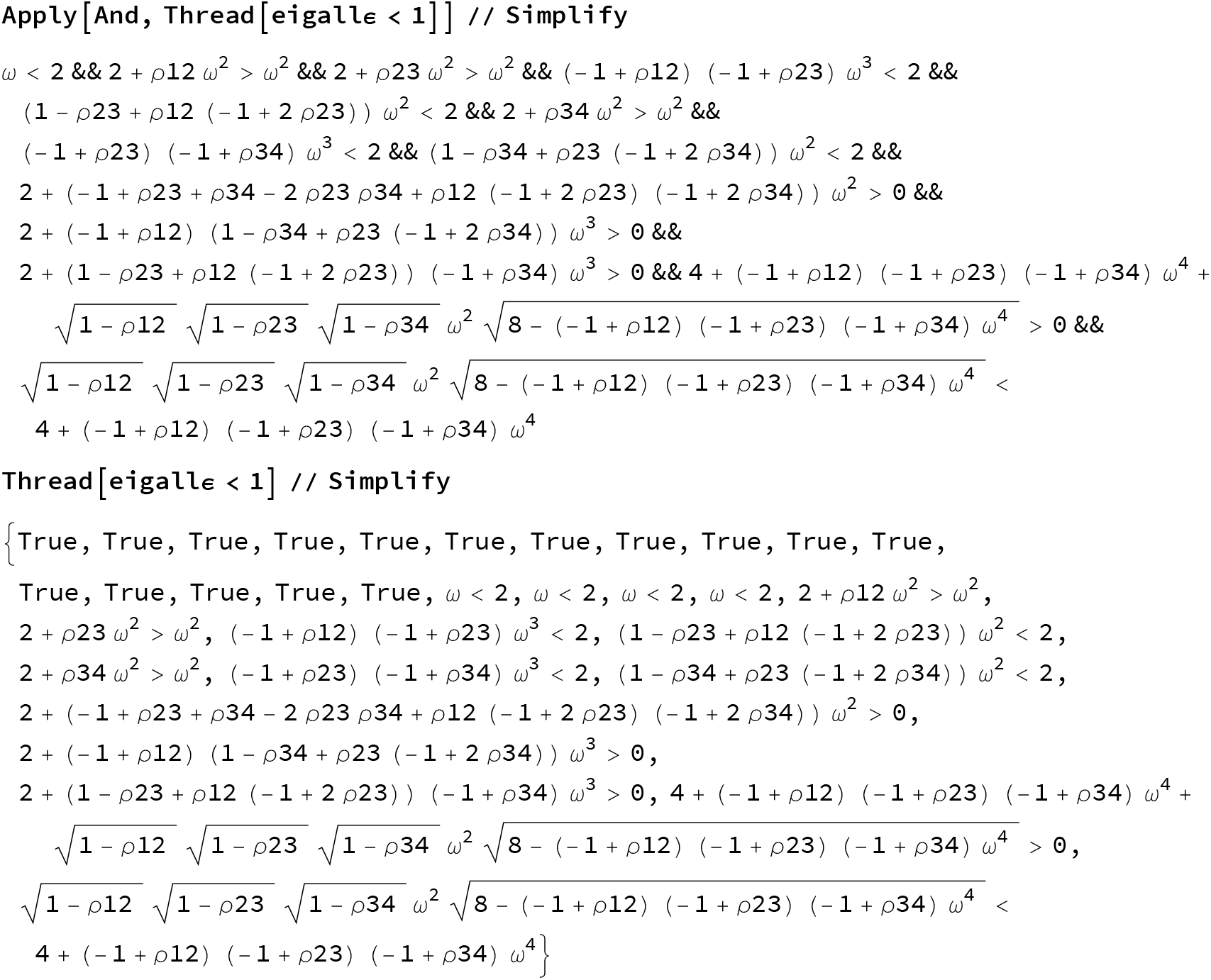

Define the conditions for every single eigenvalues for the stability of the exclusion case

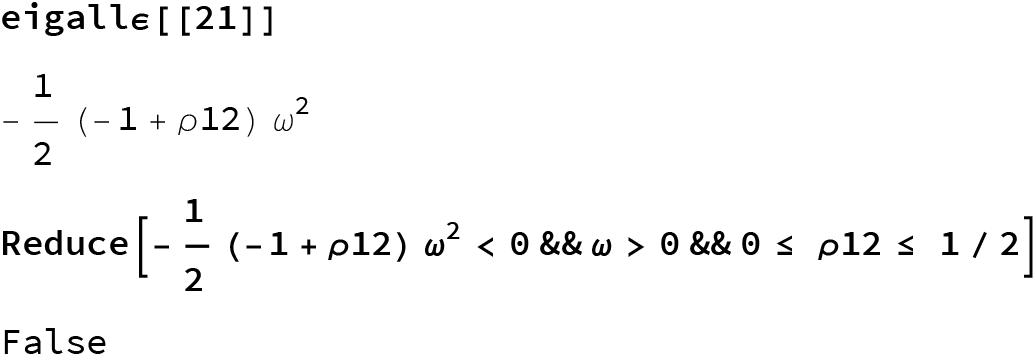

Always ≥ 0

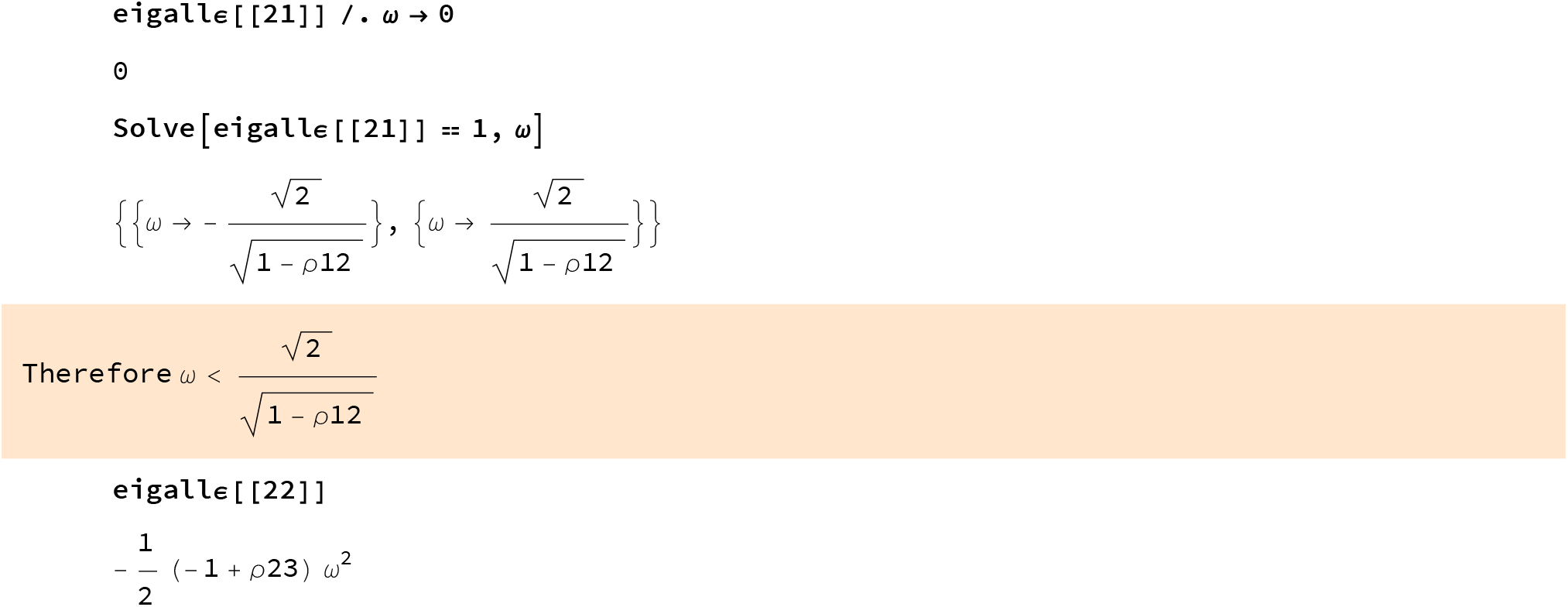

Always ≥ 0

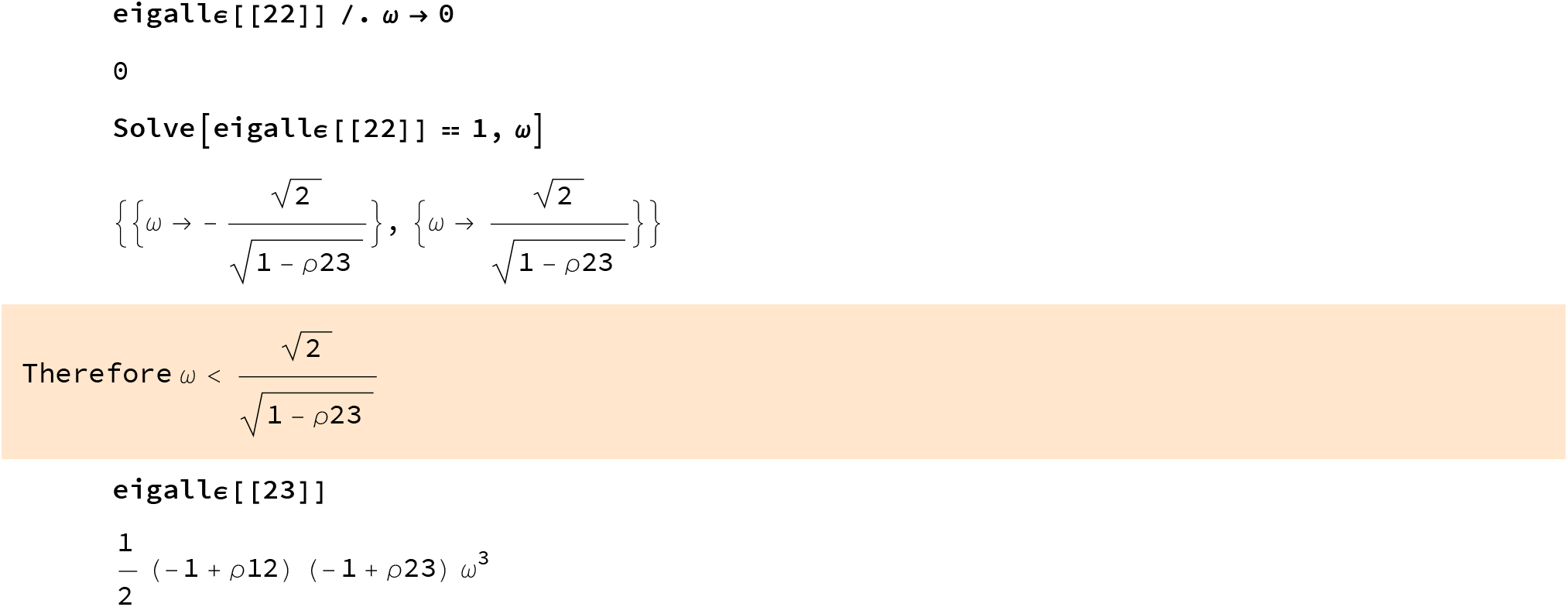

Always ≥ 0

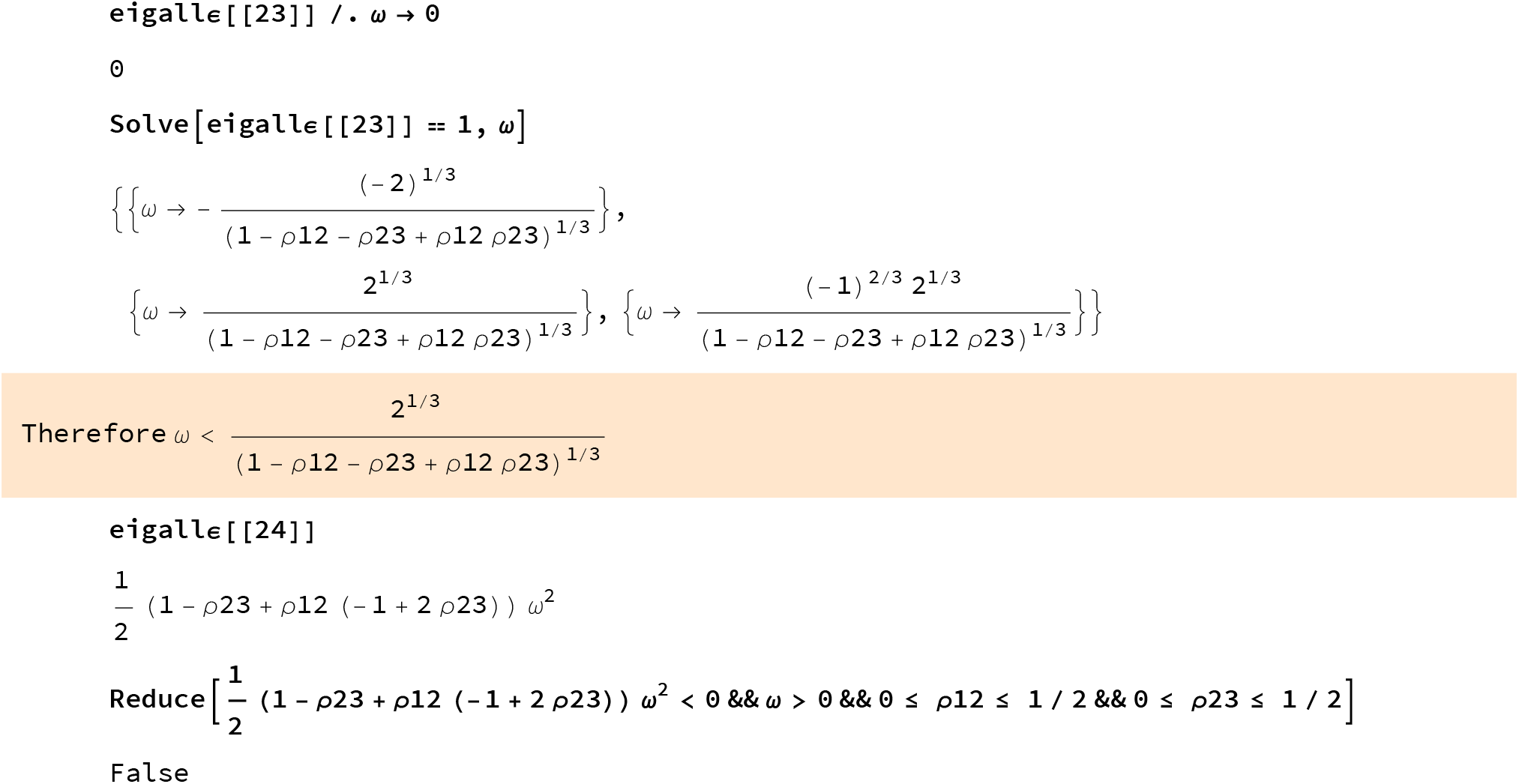

Always ≥ 0

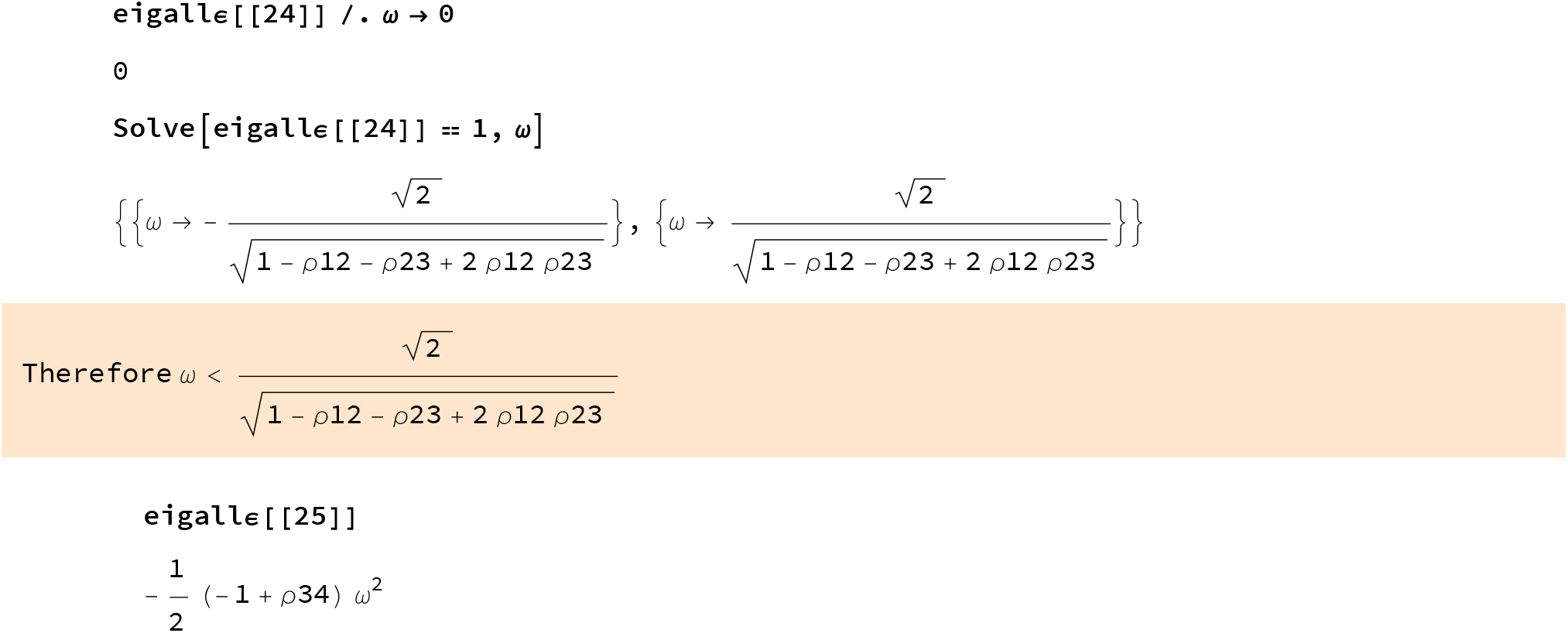

Always ≥ 0

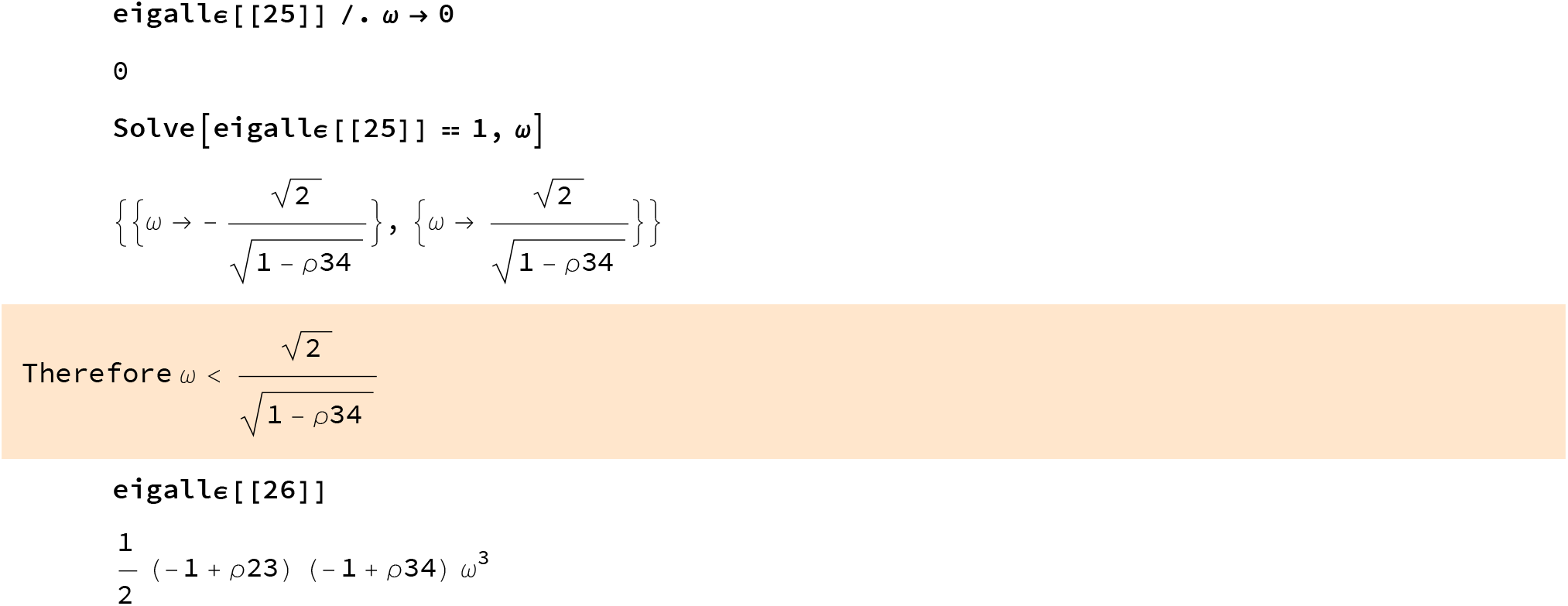

Always ≥ 0

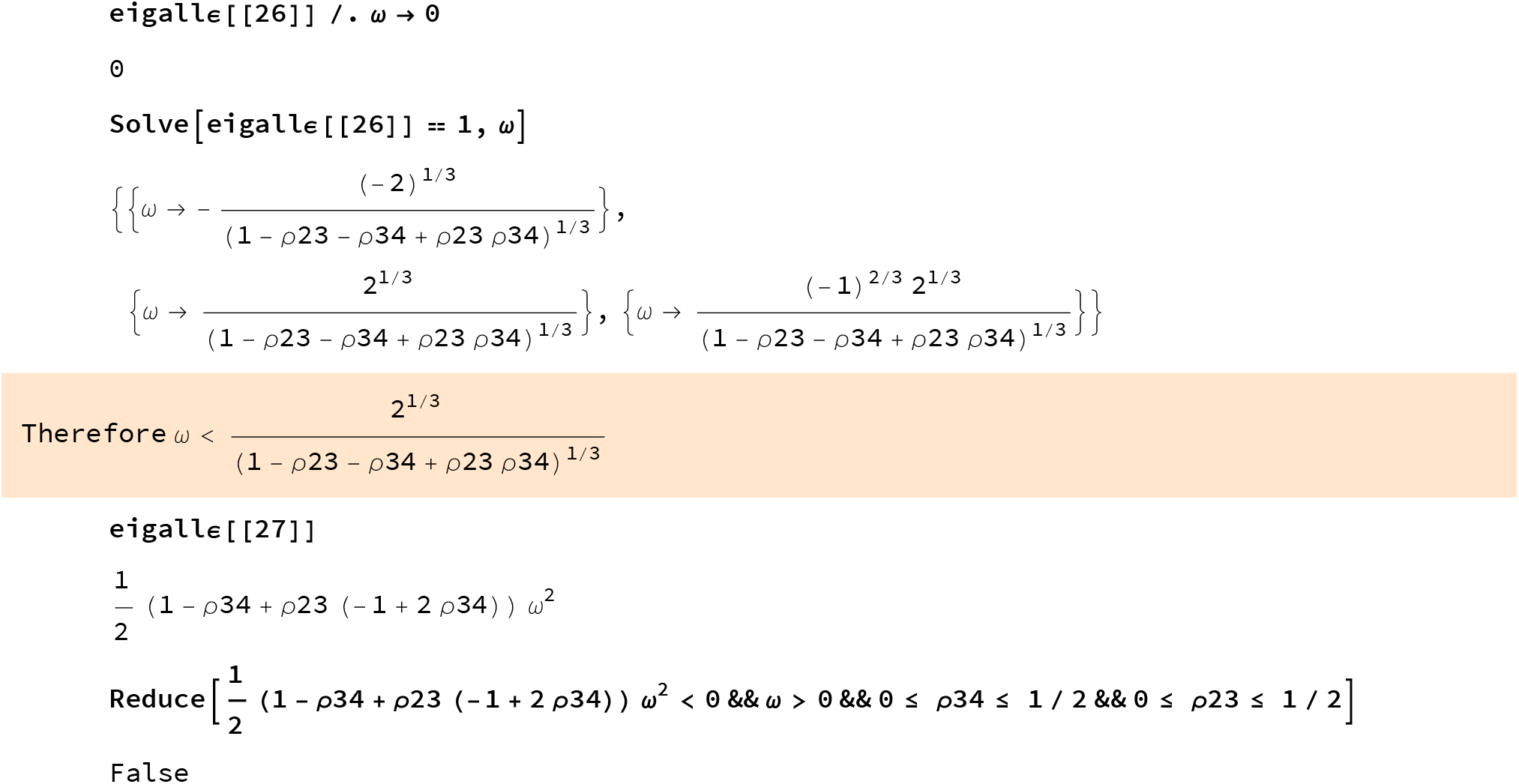

Always ≥ 0

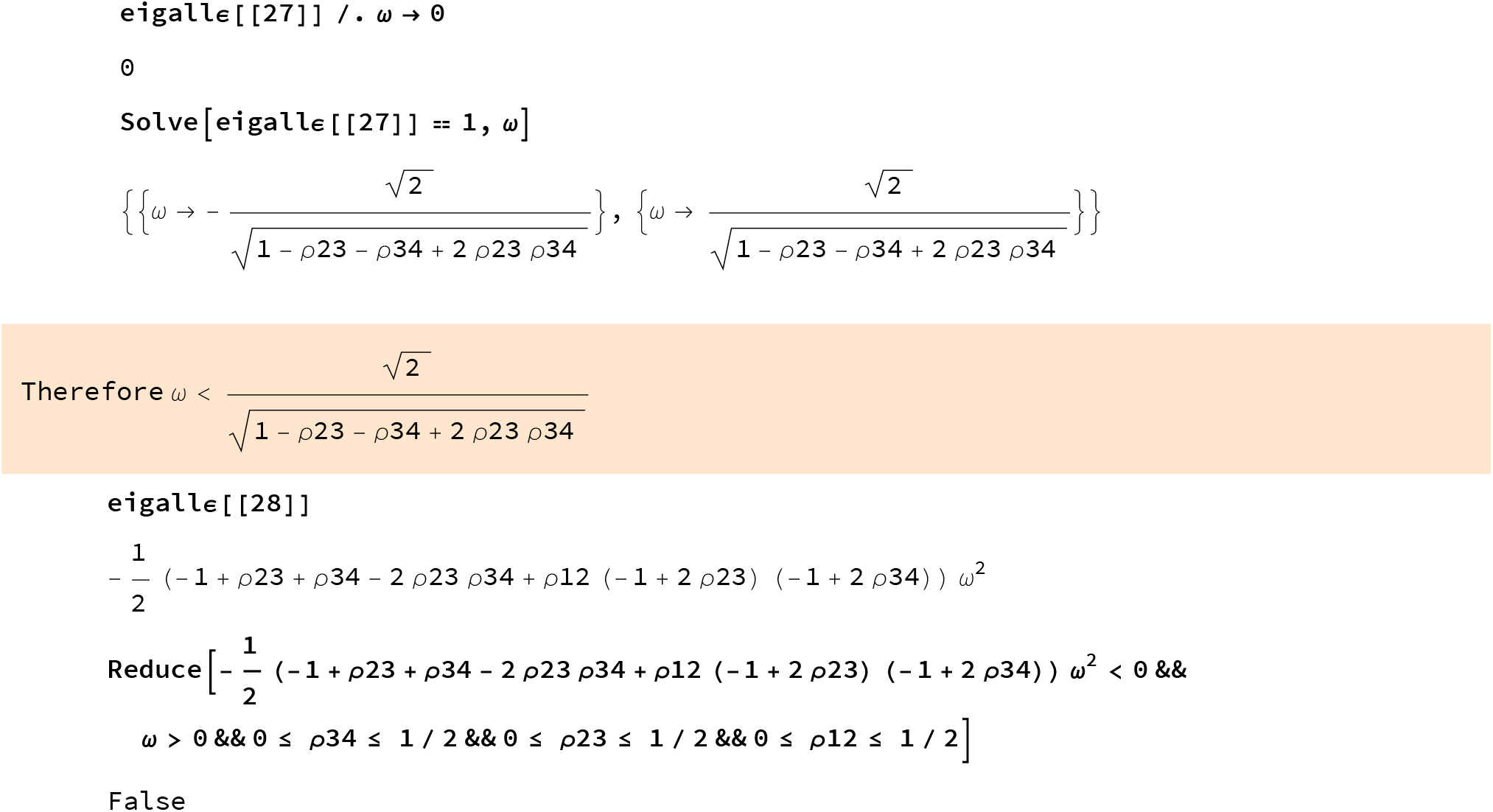

Always ≥ 0

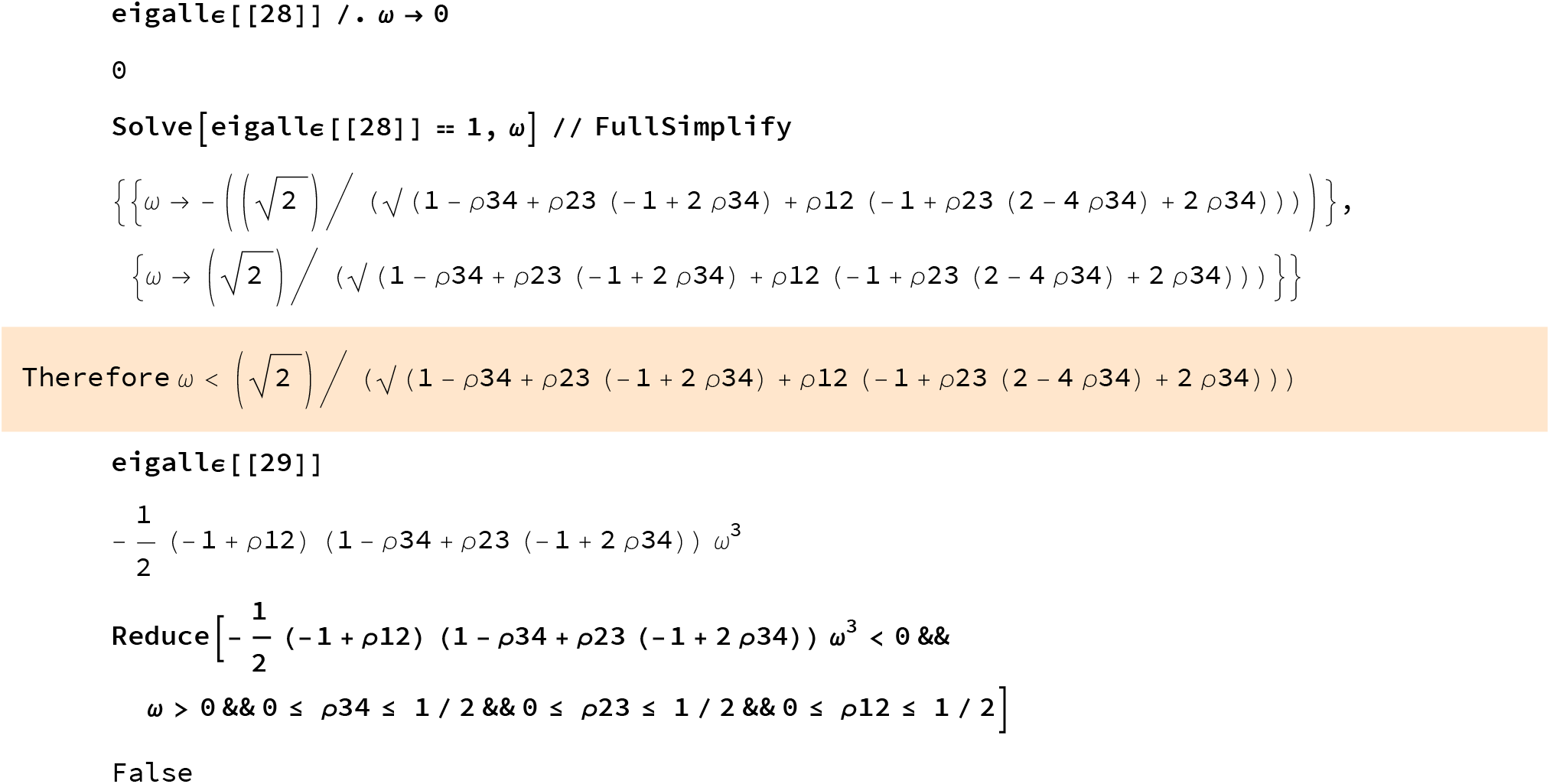

Always ≥ 0

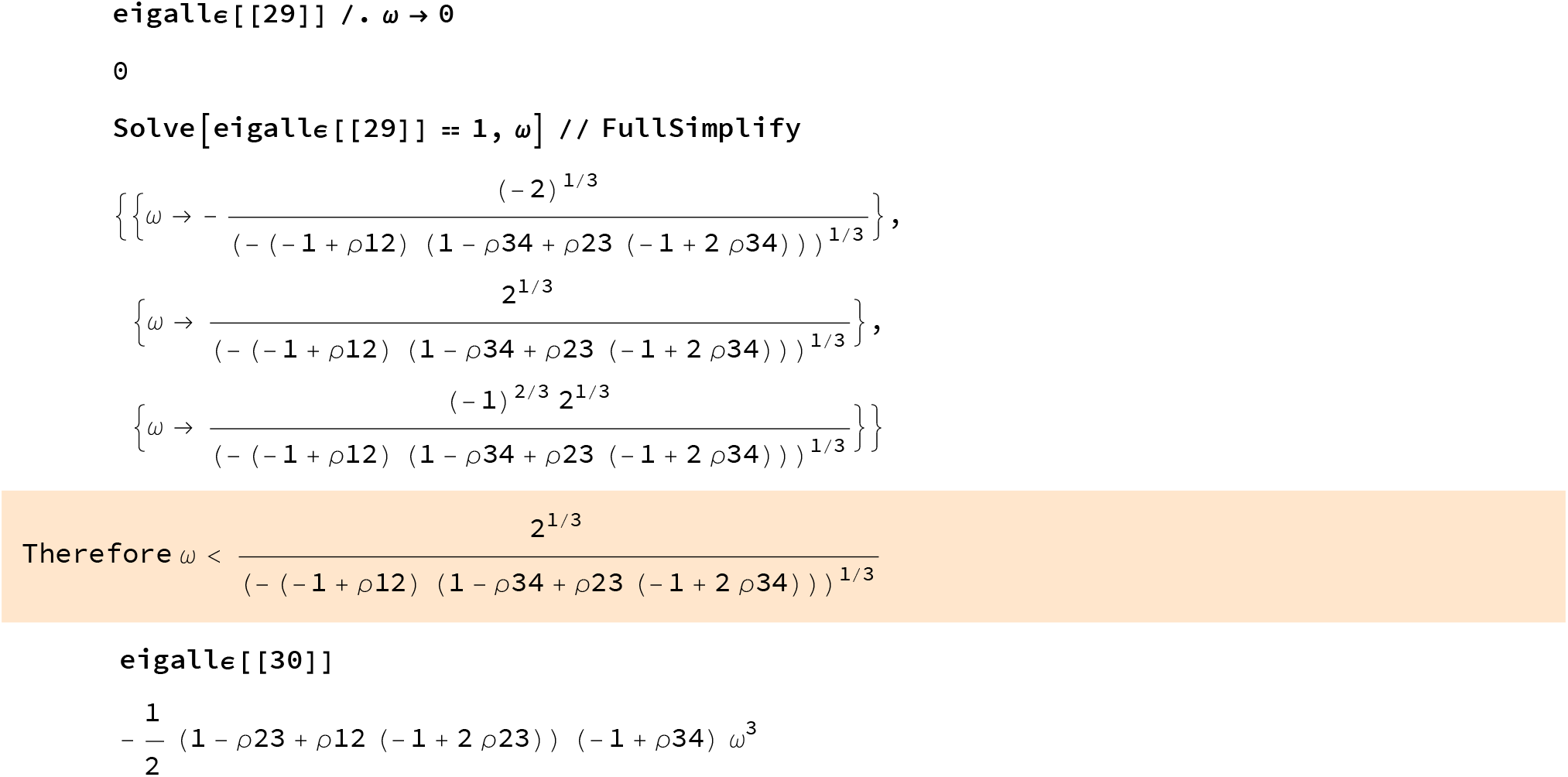

Always ≥ 0 per symmetry with the previous case

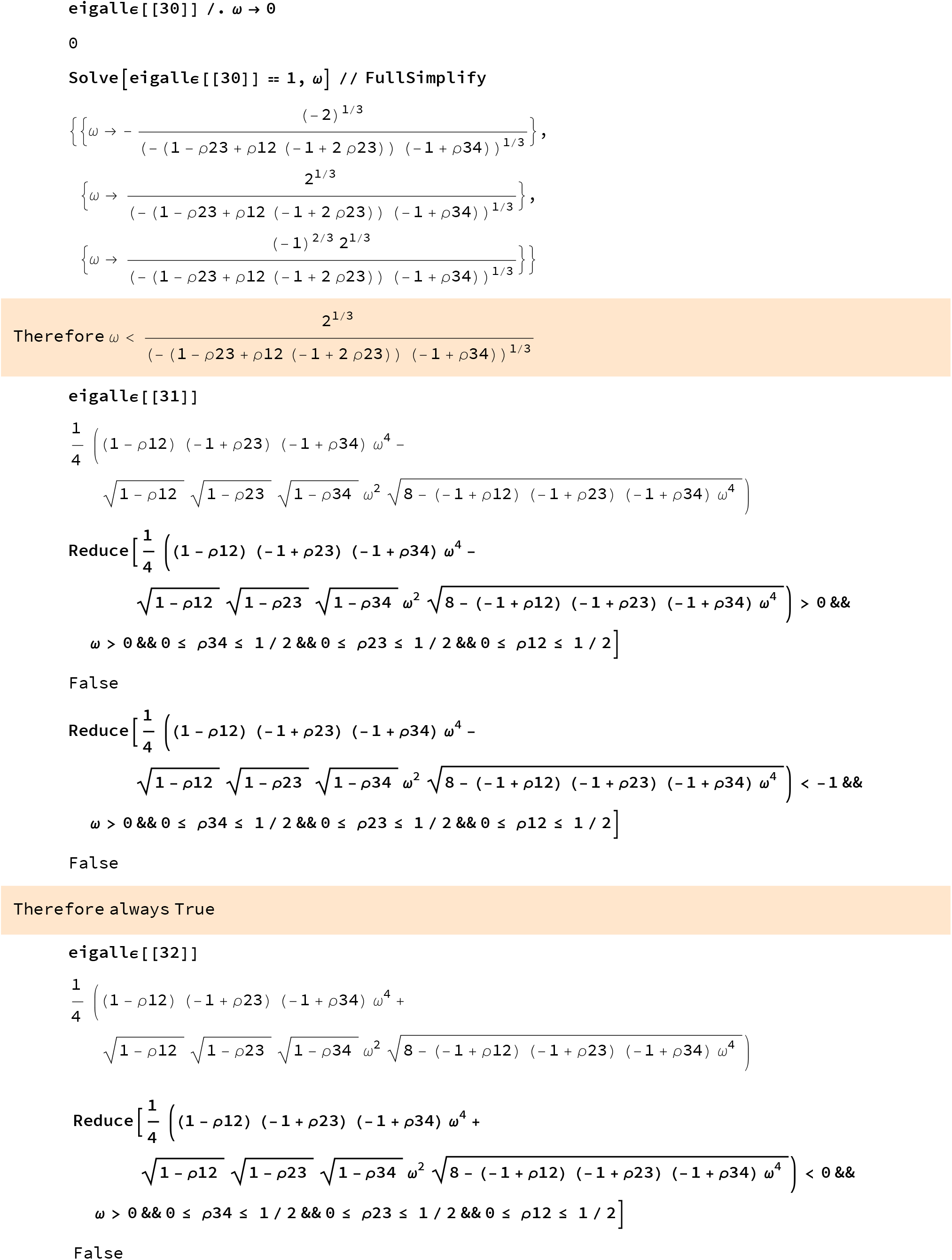

Always ≥ 0

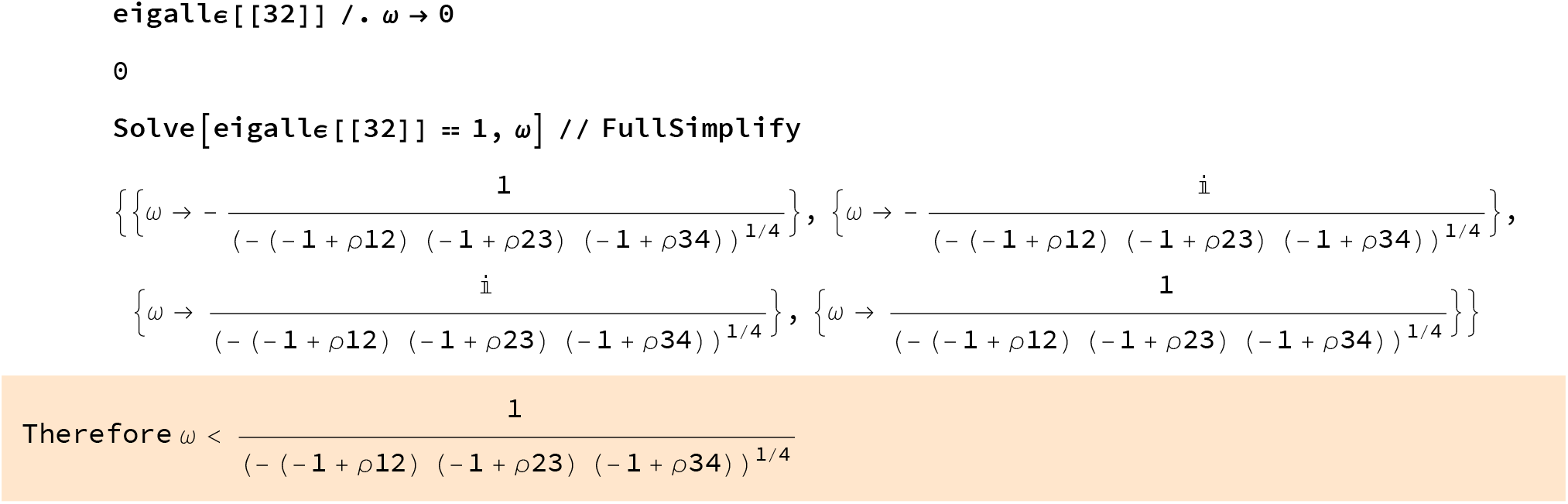

Put all the conditions together

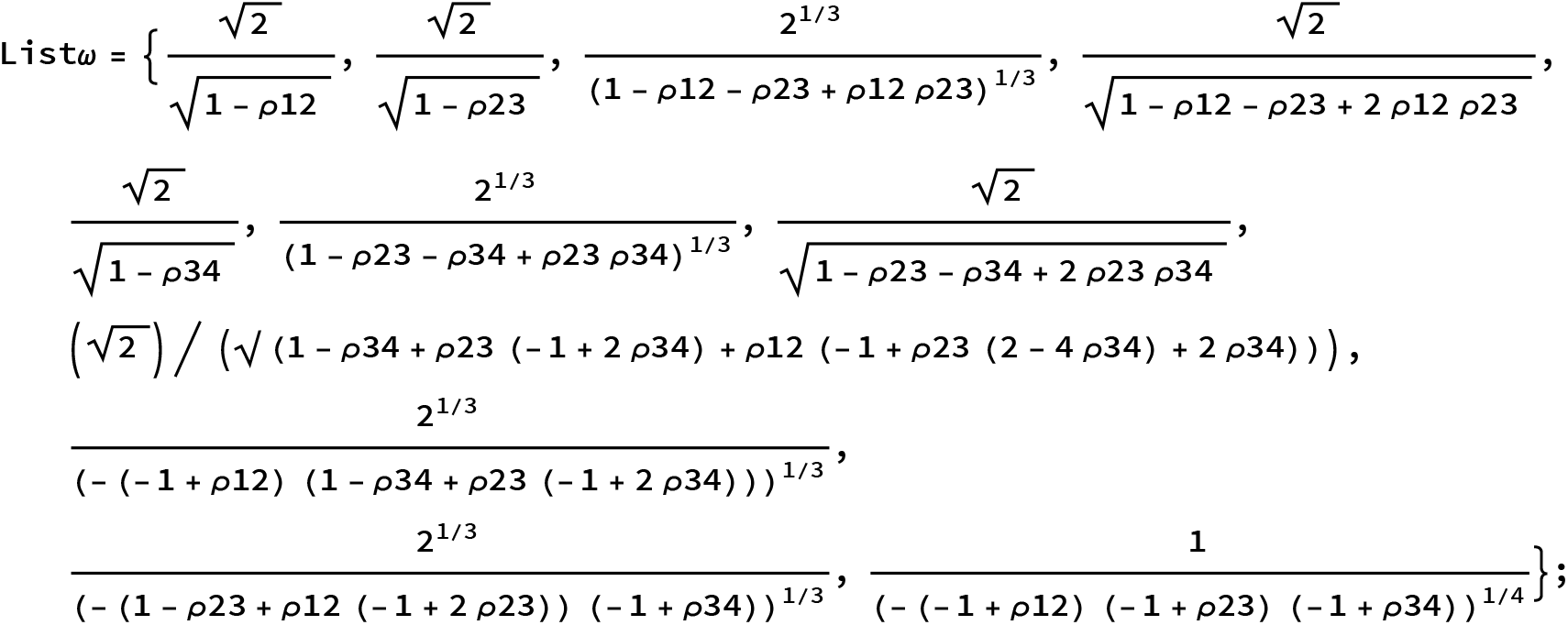
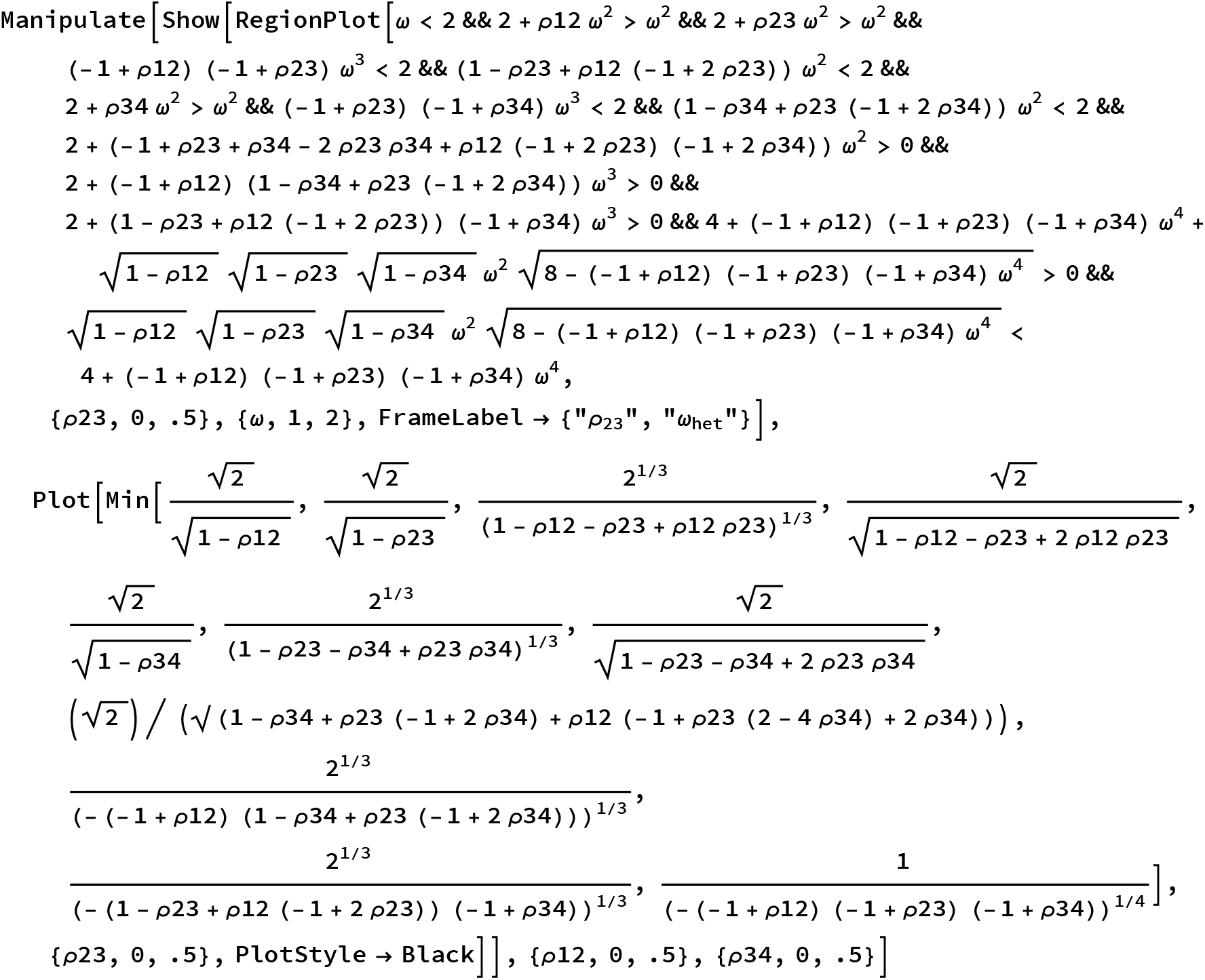
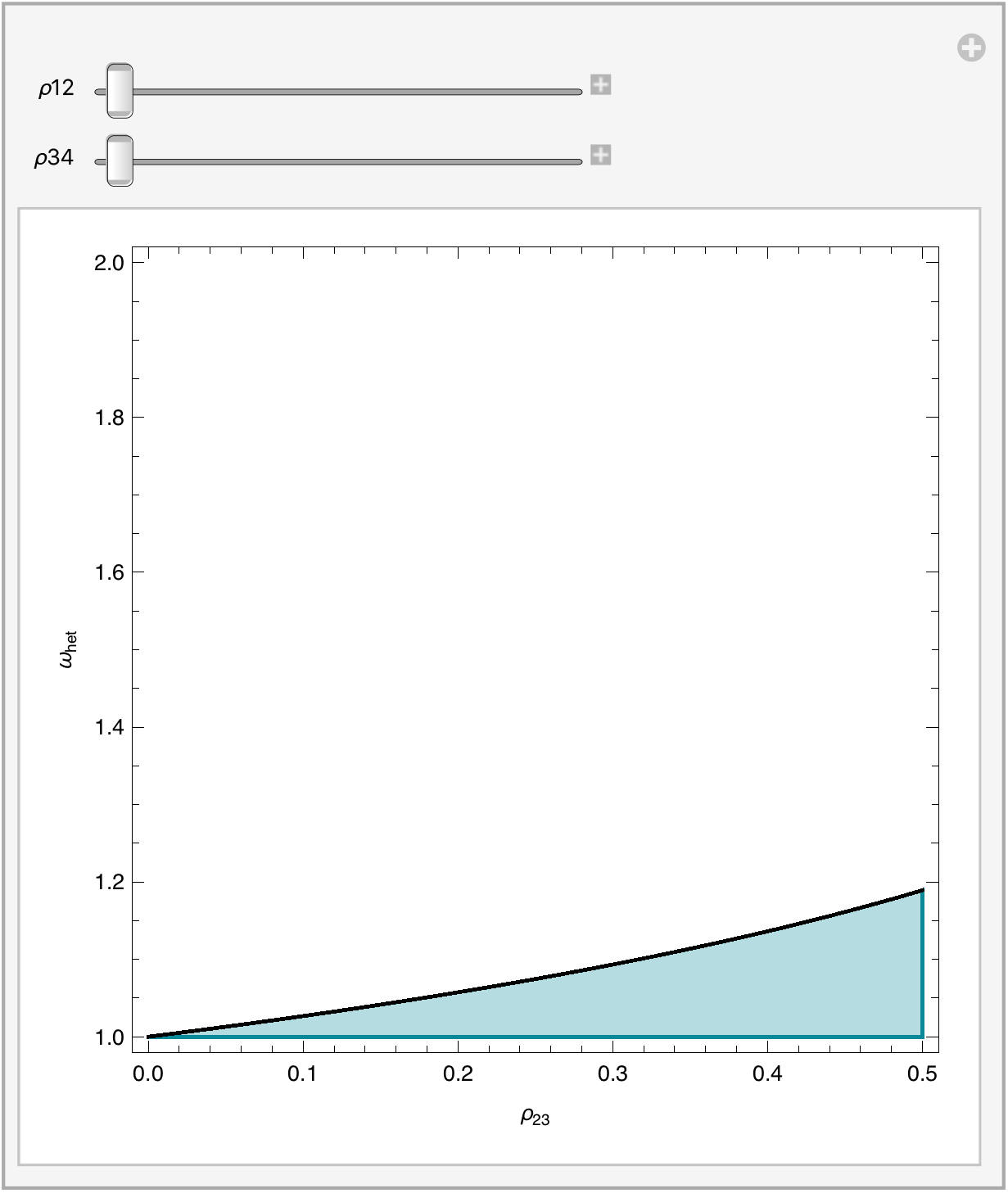
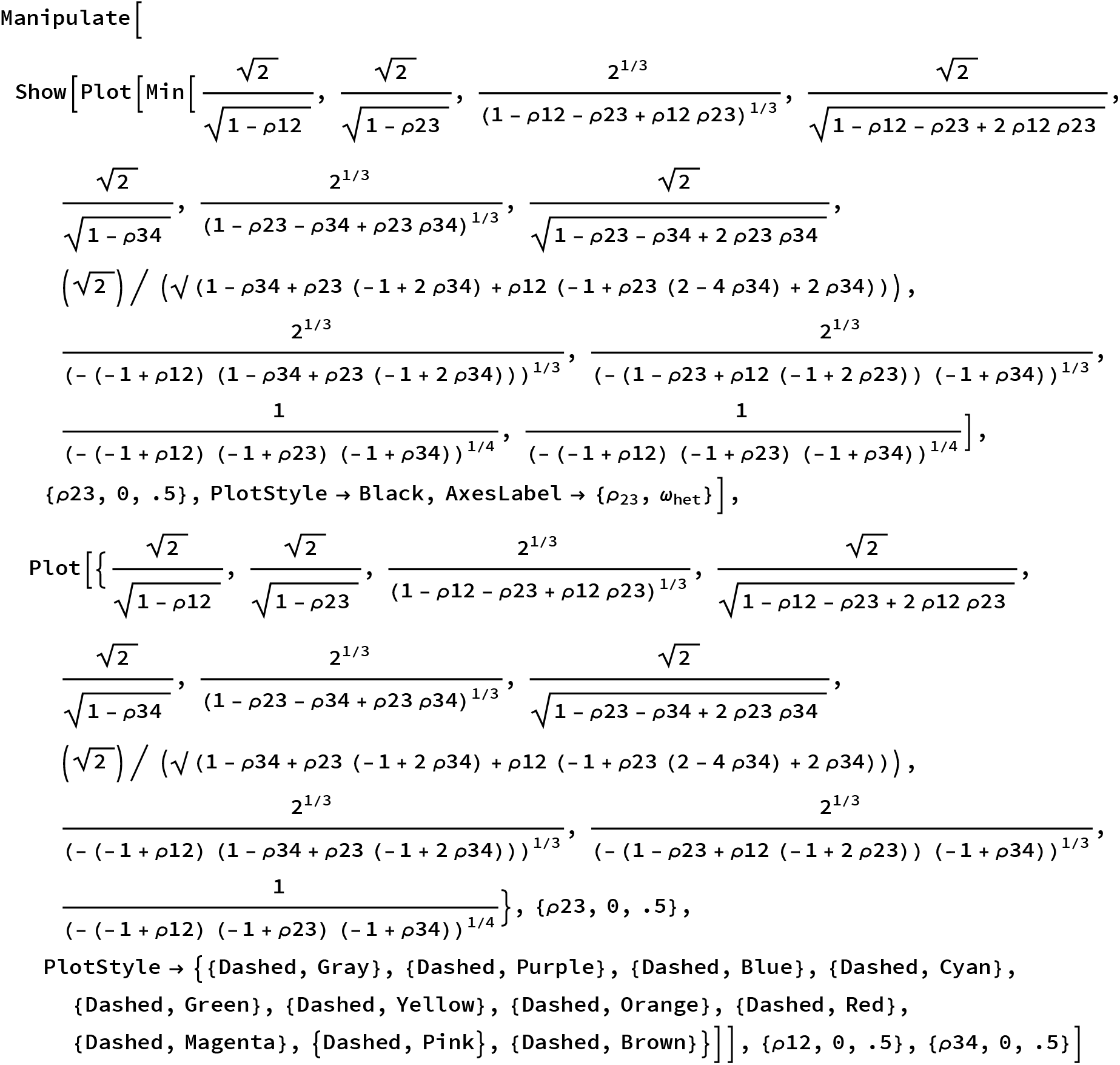
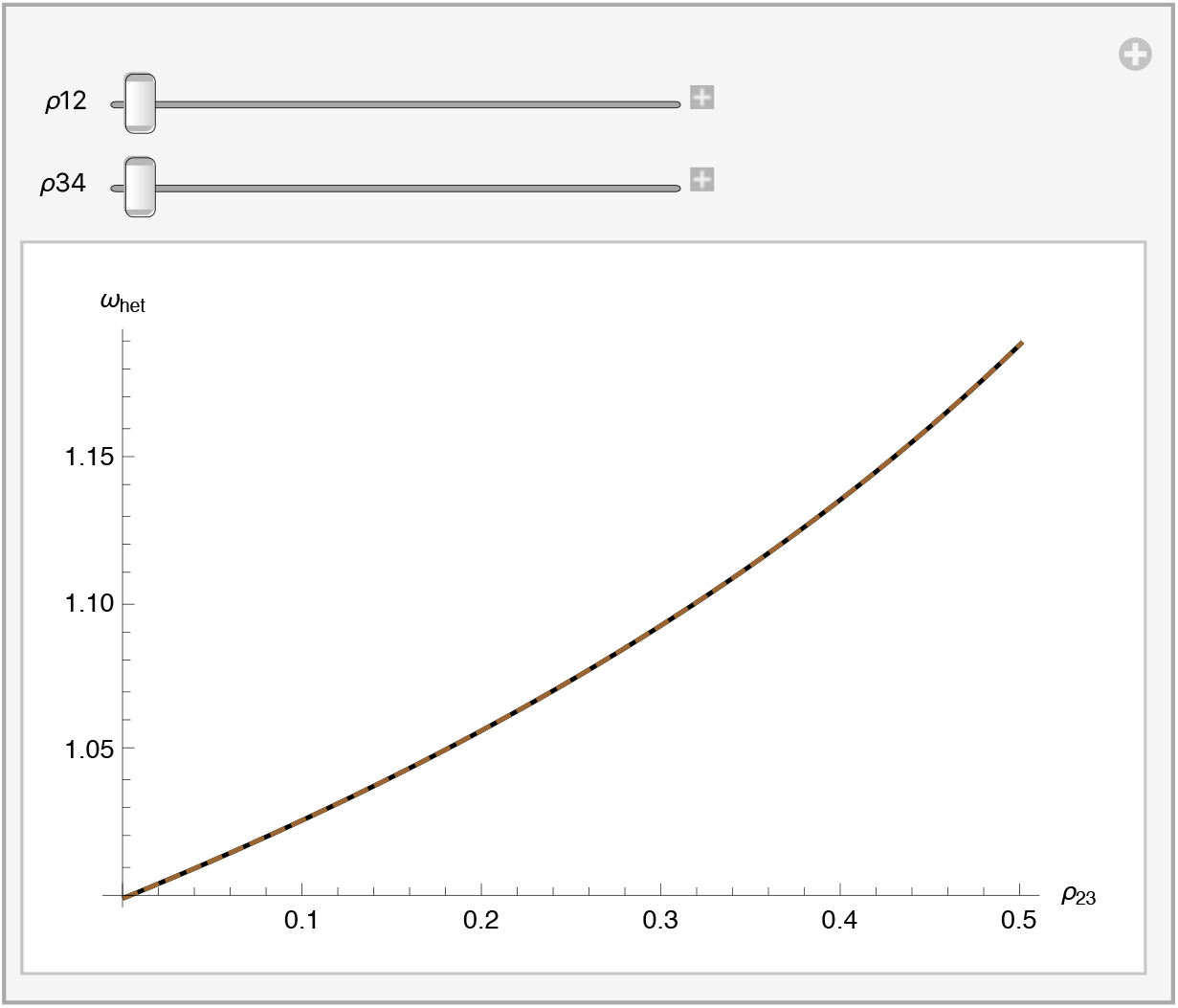

Simplify the conditions:

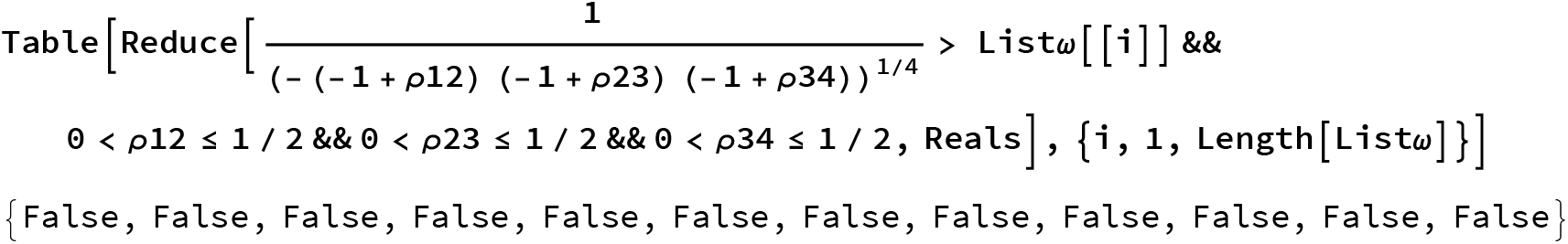

Extrapolate for n loci and shows that for n from to 2 to 10 that the more loci that more stable exclusion is

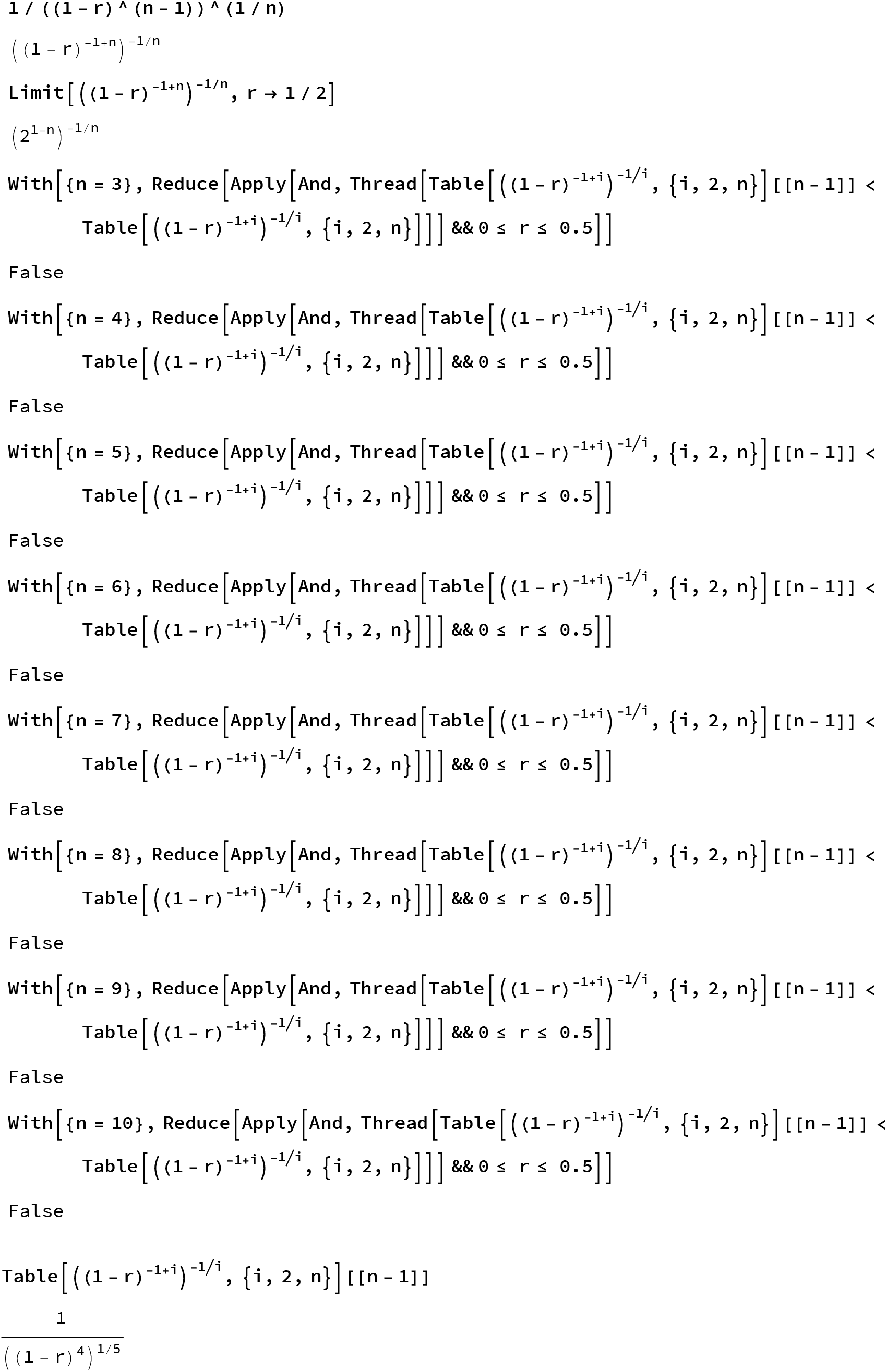

#### Plot of solution for multiple loci

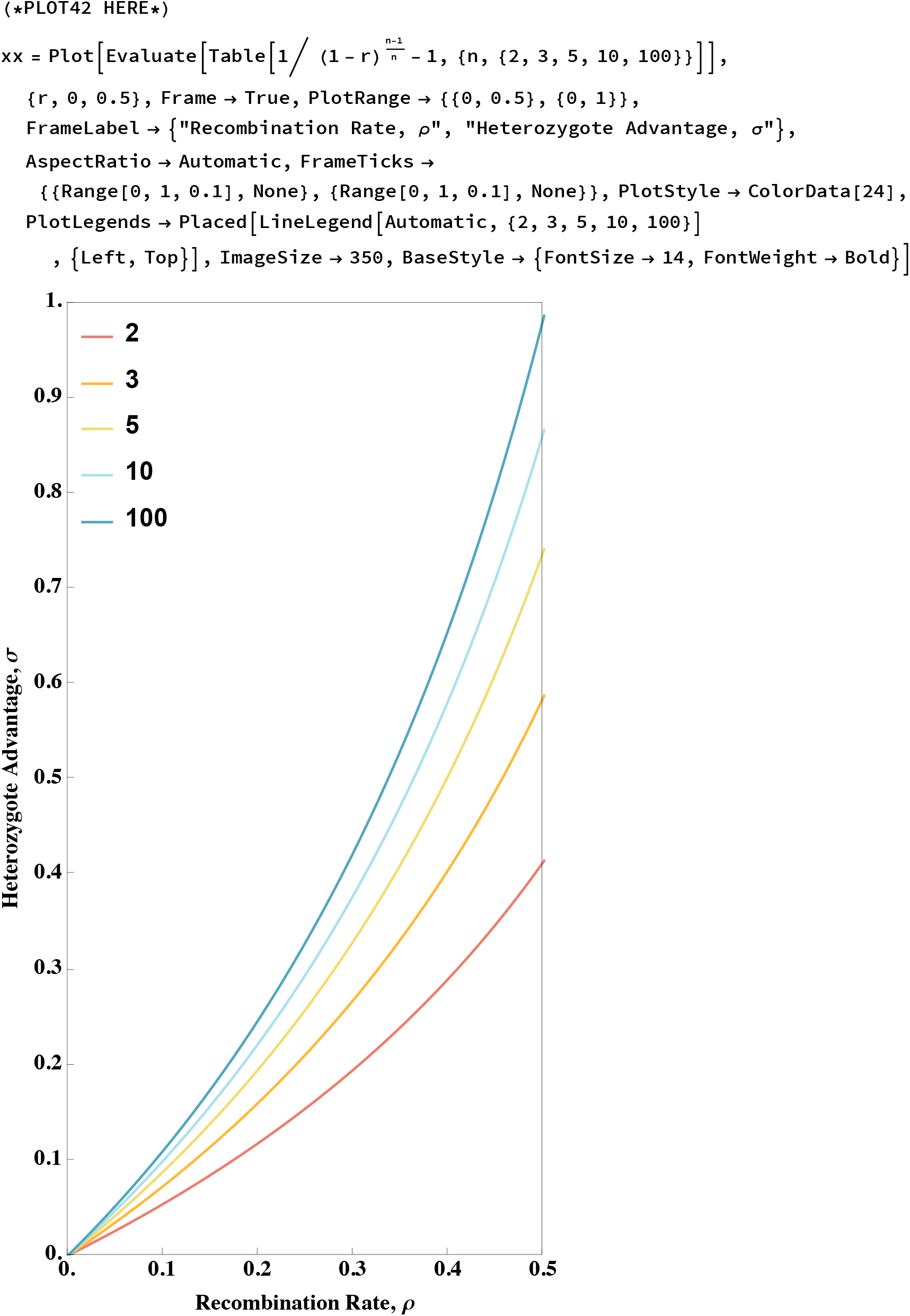

~~~
**Export[ToString[NotebookDirectory[]] <>
   "analytic-limit-lethal-haplodiploid-many-loci.pdf", xx]**
/Users/nando/Documents/finnish_hybrid_ants/results/analytic-limit-lethal-haplodiploid-many-loci.pdf
~~~

### Chapter 5: Simulations with (and without) preference

#### Code to run the simulations

For convenience in tracking convergence trajectories (but incredible annoyance in LD calculations), I summarized the 16 female genotypes into 7 classes in this order:

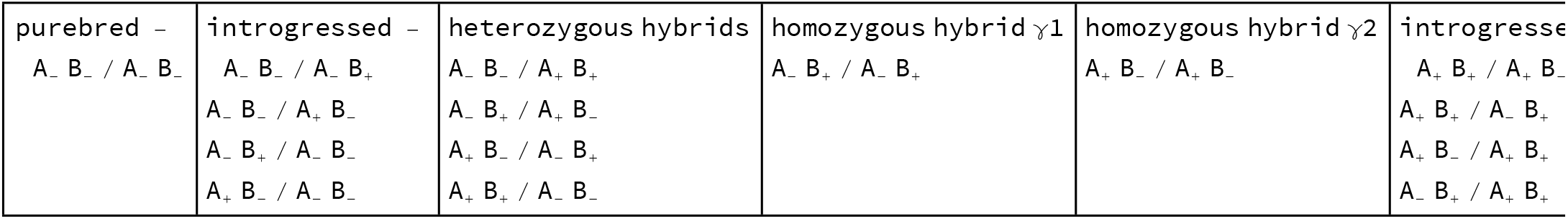

Each class has either a different fitness but cannot be assigned to one of the two parental subpopulations (e.g. homozygous hybrids) or has the same fitness as another class but is assigned to a different parental subpopulation (e.g. introgressed types).

~~~
**fem7CategoriesPosition =
fem7CategoriesPosi tion =
  {{1}, {2, 3, 5, 9}, {4, 7, 10, 13}, {6}, {11}, {8, 12, 14, 15}, {16}};
(* I determined the category of each position in the genotypes vector by hand…
 Yeah, this is rather dirty. *)
fem7Catfreq [gen_] : =
 Table [Total [gen [ [1] ][ [fem7CategoriesPosition [ [k] ]]]], {k, 1, 7}] // N**
~~~

This is the workhorse function for the simulations.

Parameter combinations are inputted first as a list in this order: {α, σ, γ1, γ2, p}

Then the starting genotype frequencies are given as a list of two lists, female genotype frequencies (16 values [0,1] summing to 1) followed by male genotype frequencies (16 values [0,1] summing to 1). Here is an example input:

{{0.4,0,0,0,0,0,0,0,0,0,0,0,0,0,0,0.6},{0.4,0,0,0.6}}

The output is to several global variables:

**convergence** returns 1 if simulation successfully converged after 100000 generations and 0 otherwise;

**finalfrequencies** and **finalfrequenciesAfterSelection** return a list of 11 genotype frequencies (first 7 correspond to female categories listed above and last 4 correspond to male haplotype frequencies) either before or after selection, respectively;

**results** and **resultsAfterSelection** return a table of genotype frequencies for every 10th generation either before selection or after selection, respectively (it is used to track trajectories to convergence);

**generation** returns the generation when convergence was reached (or 100000 if simulation failed to converge).

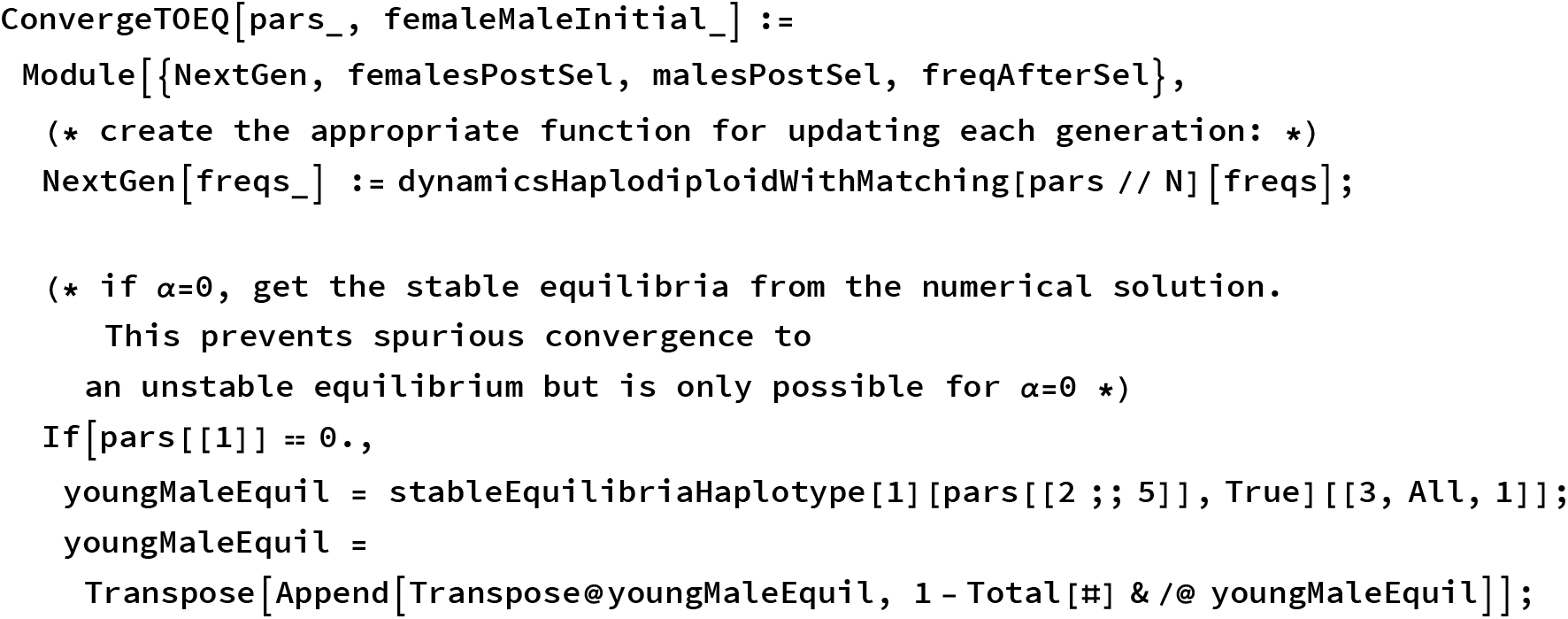
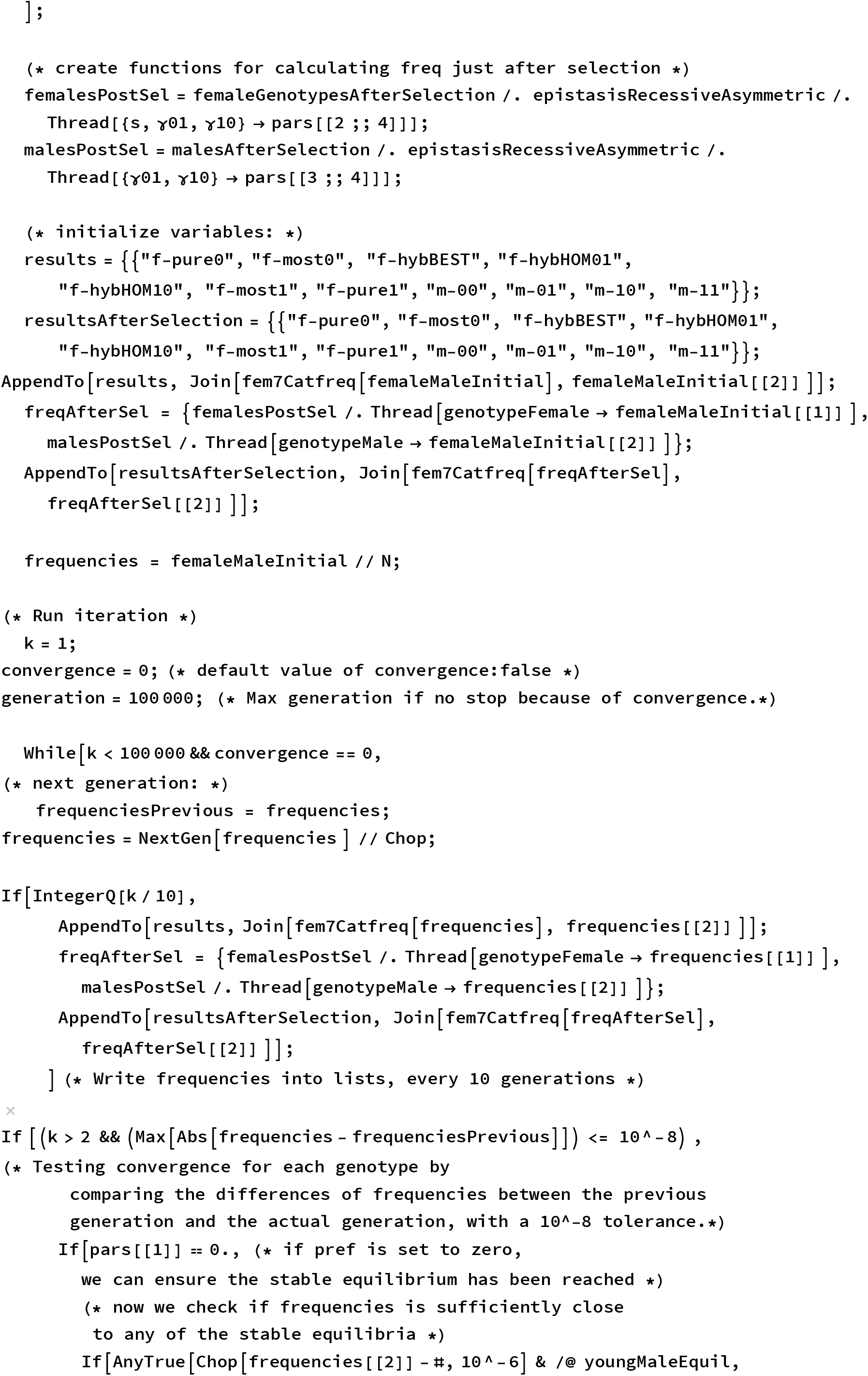
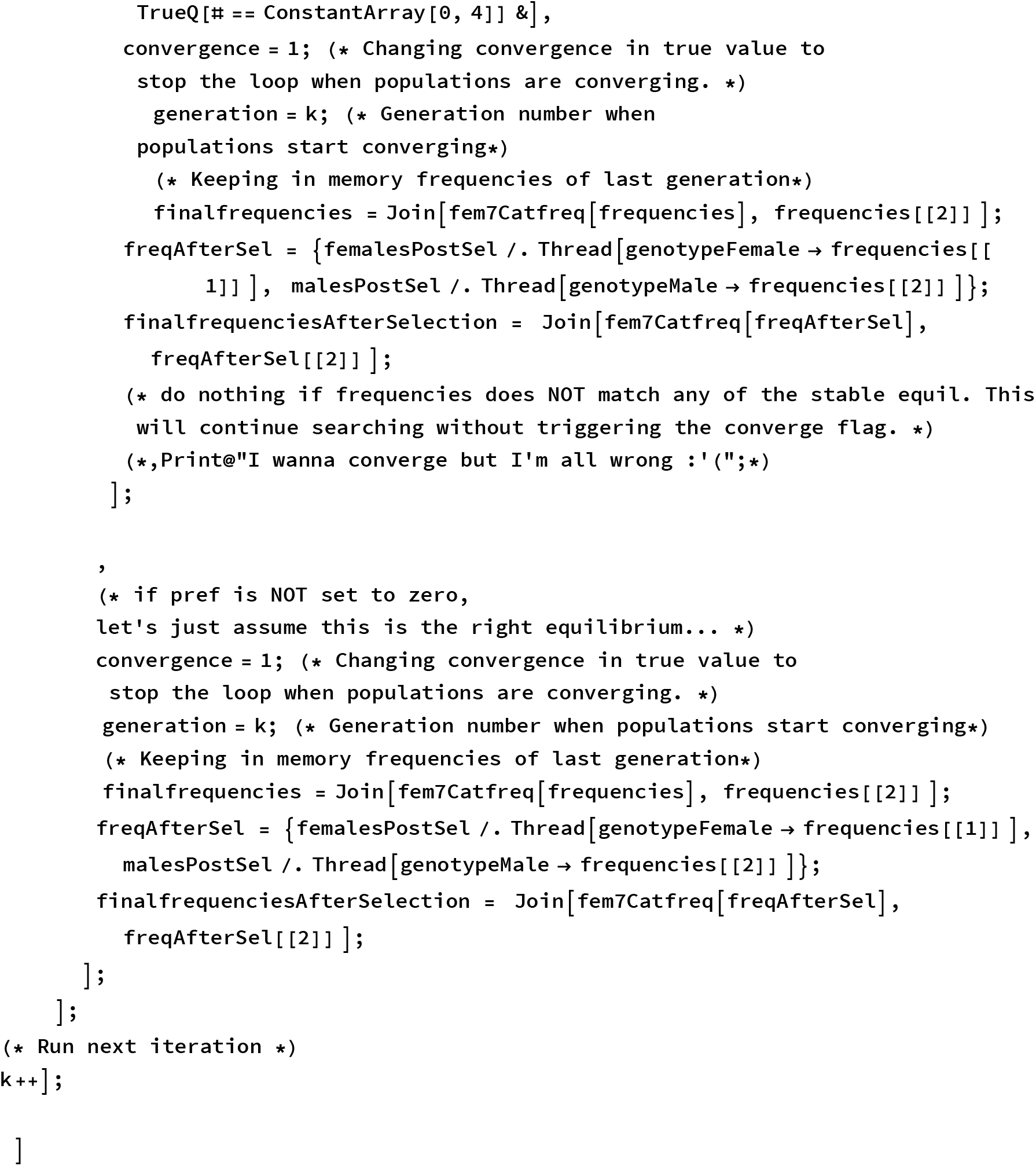

And here is the function to run simulations of the diploid model.

In this case the initial genotype frequencies are specified as just one list (16 values [0,1] summing to 1); here is an example:

{0.4,0,0,0,0,0,0,0,0,0,0,0,0,0,0,0.6}

?

Here are two functions that allow you to analyze the output from simulations with preference:

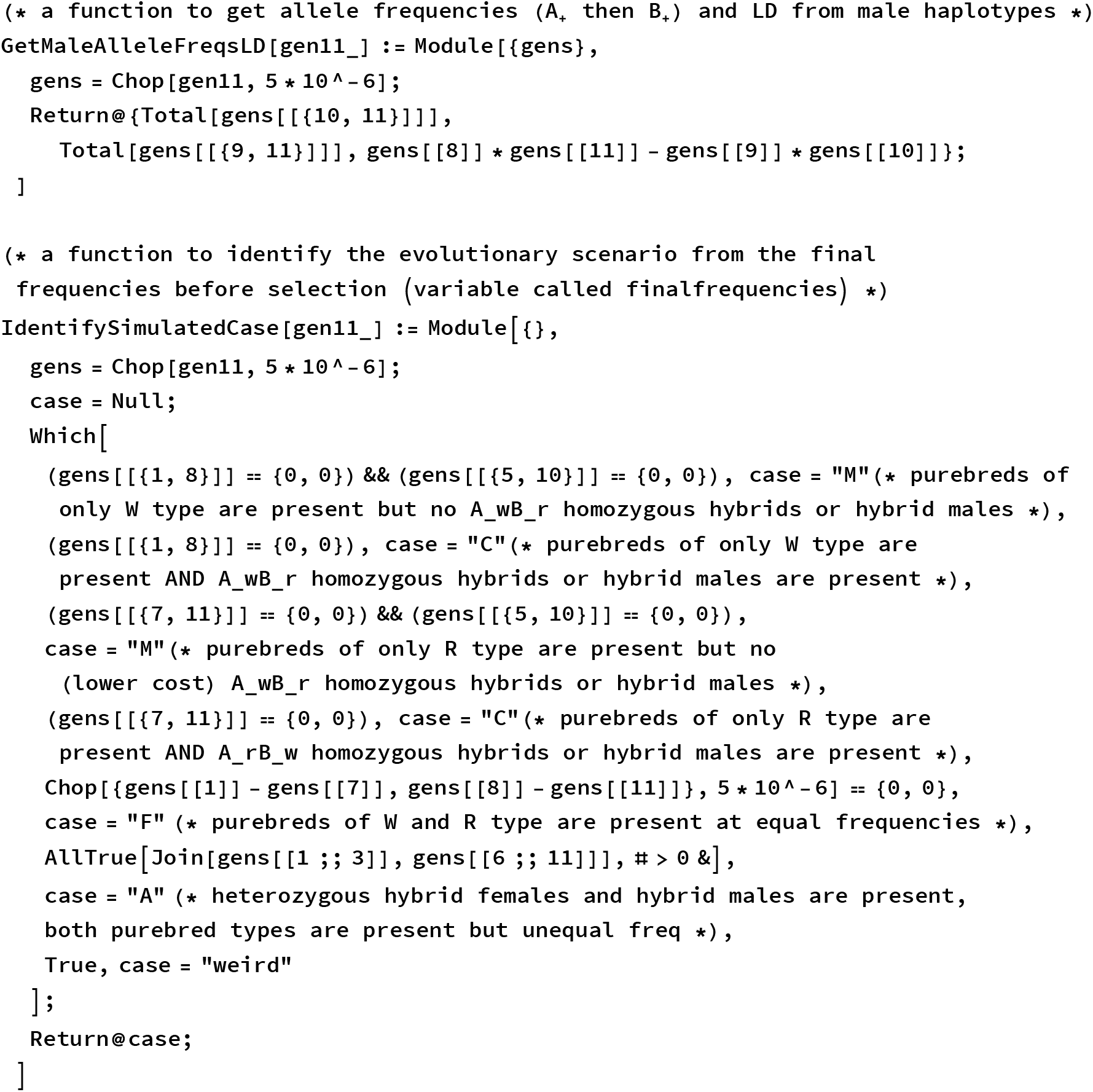

Note that you need to multiply the output from **IdentifySimulatedCase** with the boolean output from **ConvergeTOEQ** that indicates whether convergence was reached for the simulation.

#### Basin of attraction

The phase plane diagrams given in the manuscript show the basin of attraction. See Chapter 2 above for plotting the phase plane diagrams. The basin of attraction was done by simulation below.

This function finds the index of the stable equilibrium (from **stableEquilibriaHaplotype**) that is converged to for a transect along the secondary contact line:

~~~
SecContactBasin[pars_, pqVals_ (* a list from (0,1) avoiding 0.5 *)] :=
 Module[-init},
  convergeOUT = {};
  genOUT = {};
  eqIndexOUT = {};
(* pick initial conditions along y=x secondary contact *)
Do [
 (* get initial genotypes *)
 init = GetGenotypesfromHaps[GetHapsfromAlleles[pq, pq]];
 (* run simulation *)
 ConvergeTOEQ[Join[{0.}, pars], init];
 (*
 TimeConstrained[ConvergeTOEQ[Join[{0.},pars], init],1,”Fail!”] (* increase the
  time if the function gives errors about “Part 1 of {} does not exist. “ *)
*)
(* keep info on whether convergence happened *)
 AppendTo[convergeOUT, convergence];
 AppendTo[genOUT, k];
 (* keep index of converged equilibrium *)
 AppendTo[eqIndexOUT,
  Position[Chop[finalfrequencies[[8;; 11]] -#, 10Λ-6] & /@ youngMaleEquil,
   ConstantArray [0,4]][[1,1]]];
, {pq, pqVals}];
]

 (* here is the complete list of pqVals I used for analysis: *)
 pqRange = Subdivide[0.01, 0.98, 11] (* first 2 values picked to avoid symmetry *)
~~~

#### Grid plots of different parameter combinations

Create grid plots without preference:

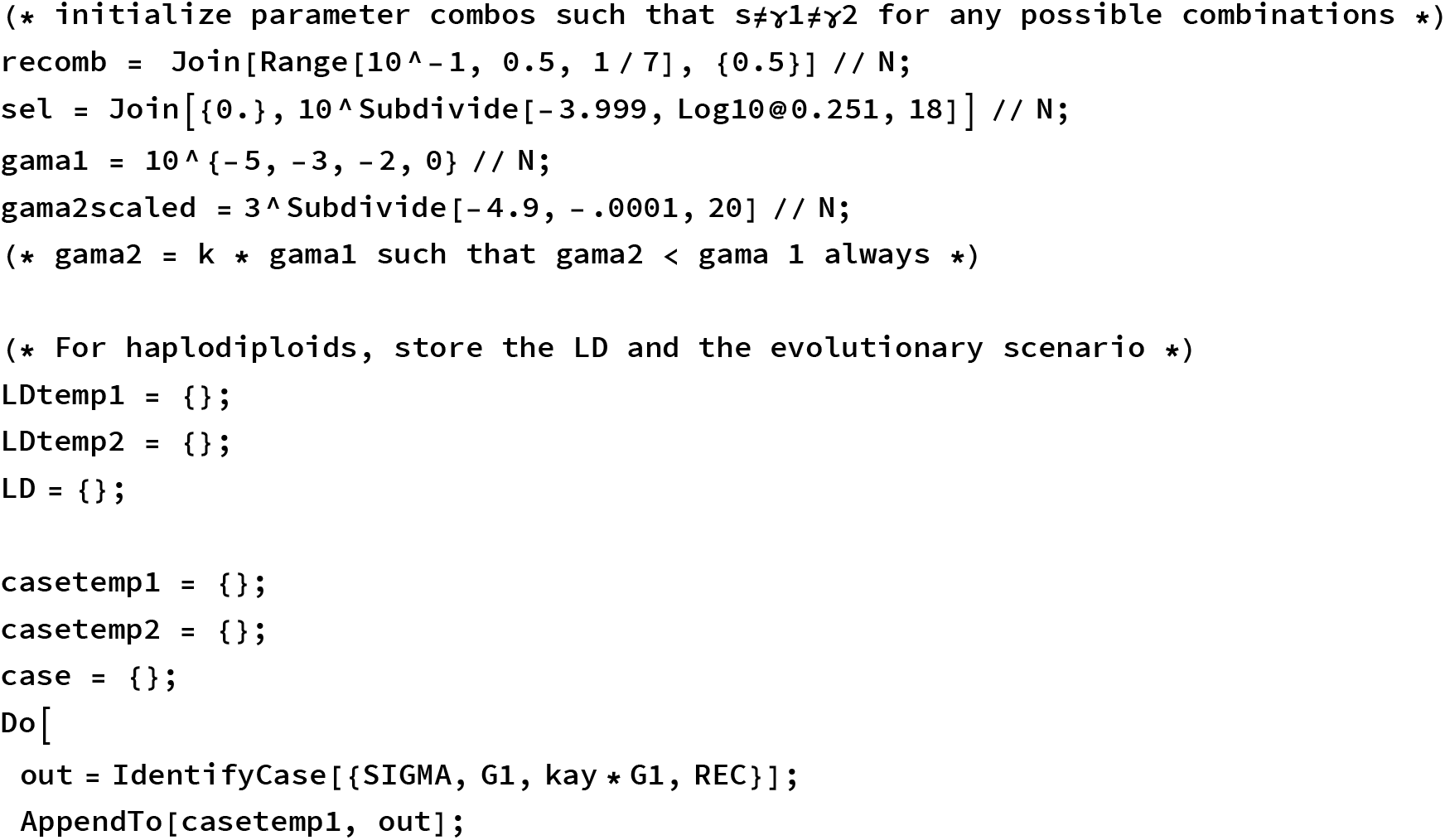
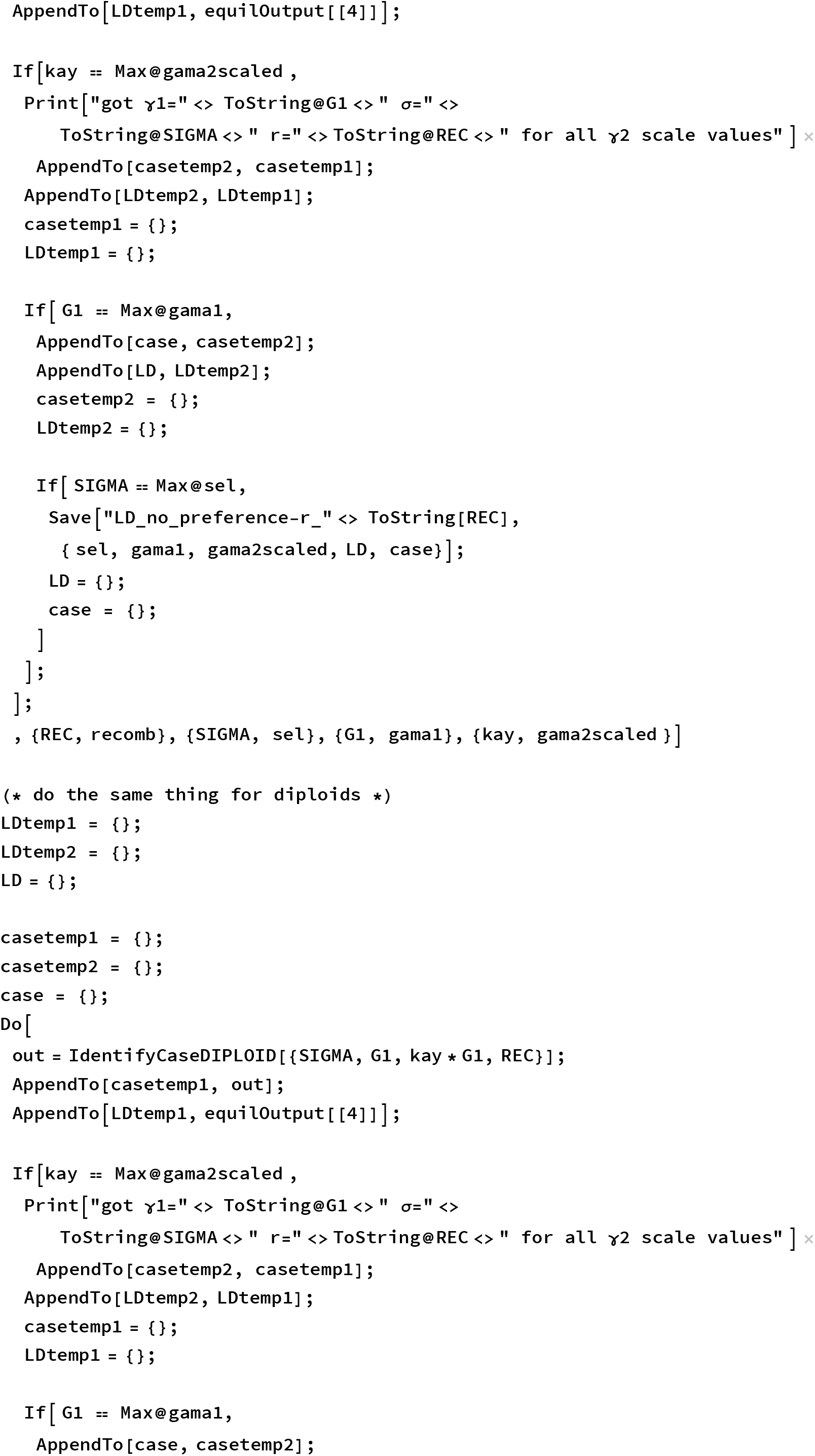
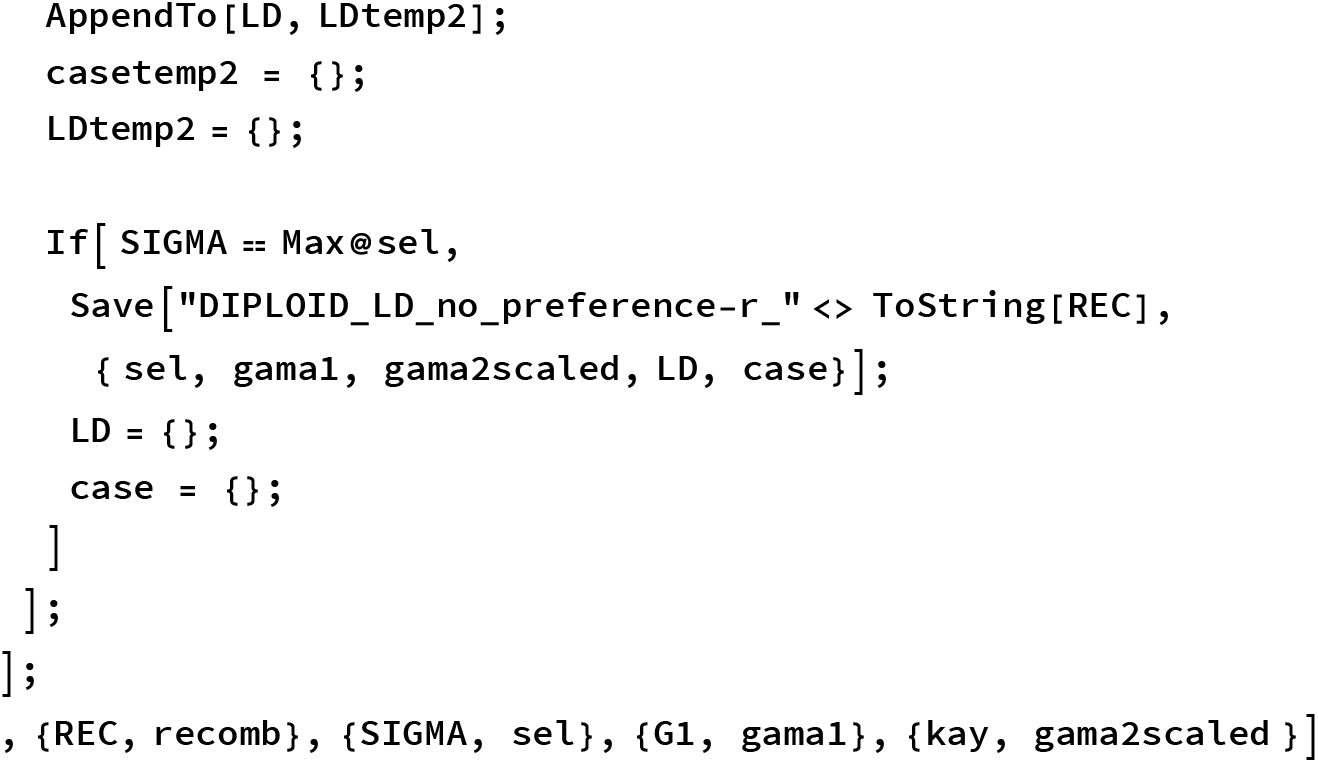

Here is a function to get LD/Dmax from the list of 3 values that I stored for all of the LD’s: {freq of allele w at locus A, freq of allele w at

~~~
**locus B, LD}
LDoverDmax[vals_] :=
Quiet[vals[[3]] / (Min[1 - vals[[1]], 1 - vals[[2]]] * Min[vals[[1]], vals[[2]]])]
**
~~~

### Chapter 6: Data fitting

First, here are the estimated values for the male frequencies in the observed data and the purebred female frequencies:

~~~
**
(*male frequencies before selection: *)
maleRpureBS = 0.0412;
maleRhybridBS = 0.0618;
maleWhybridBS = 0.26013;
maleWpureBS = 0.63687;

(*male frequencies after selection: *)
maleRpureAS = 0.1022;
maleRhybridAS = 0.0008110;
maleWhybridAS = 0.06928;
maleWpureAS = 0.8277;

(*female frequencies before selection:
  “intro” refers to females inferred to be purebred
 based on having NO loci heterozygous for introgressed alleles
  “diag” refers to females inferred to be purebred based on having
 MORE THAN ZERO loci homozygous for diagnostic alleles *)
femaleRintroBS = 0.02163;
femaleWintroBS = 0.51129;
femaleRdiagBS = 0.08343;
femaleWdiagBS = 0.31395;

(*female frequencies after selection: *)
femaleRintroAS = 0.;
femaleWintroAS = 0.4238;
femaleRdiagAS = 0.002142;
femaleWdiagAS = 0.1889;**
~~~

#### Data fitting to the model without preference

Two functions to calculate the sum of squared differences (SSD) between the model and data. The input is a list of two lists, first before selection and second after selection. Each of the two lists gives the 11 genotype frequencies (first 7 correspond to female categories listed above and last 4 correspond to male haplotype frequencies).

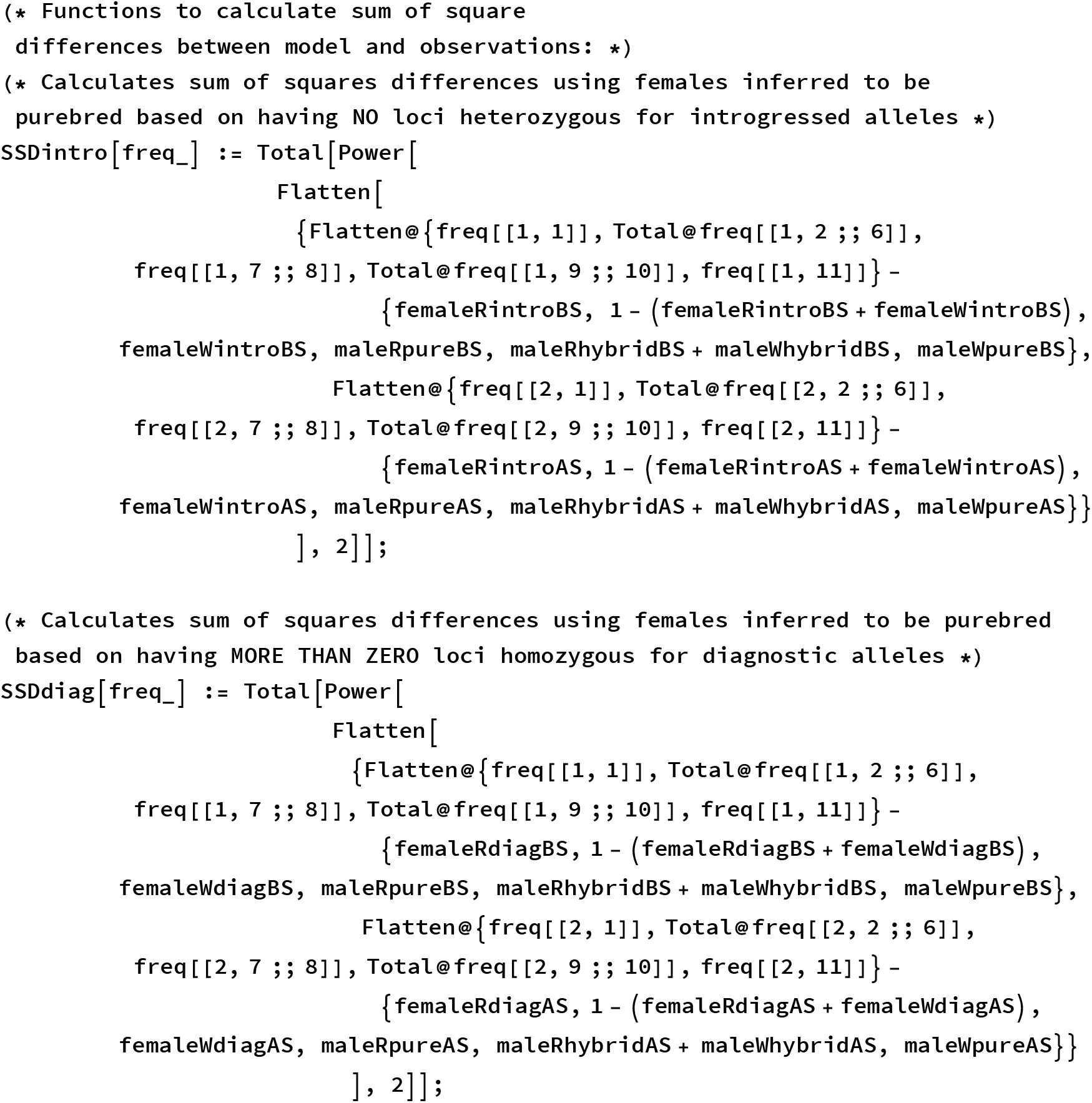

Calculate the SSD for a lot of parameter values and save the results to output:

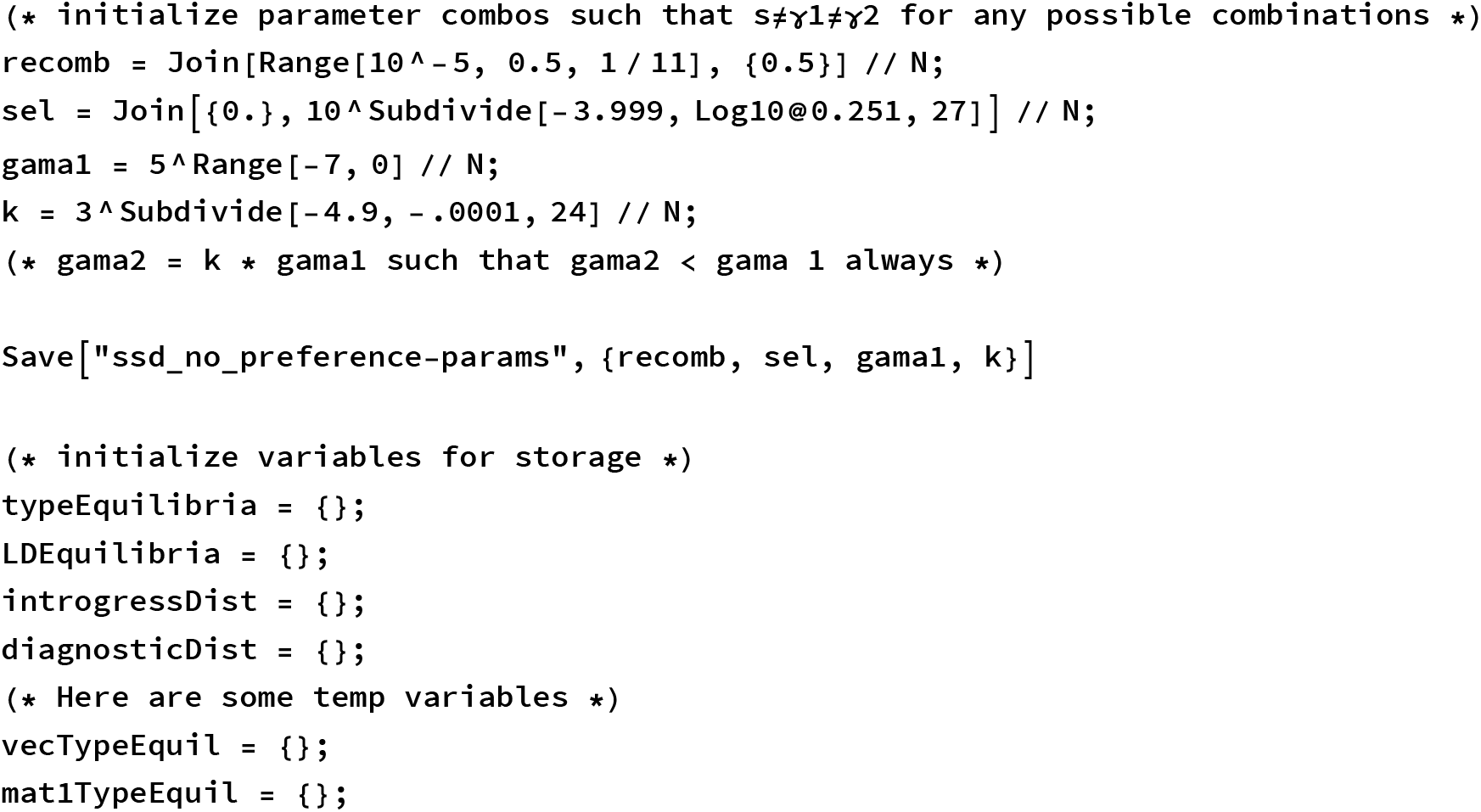
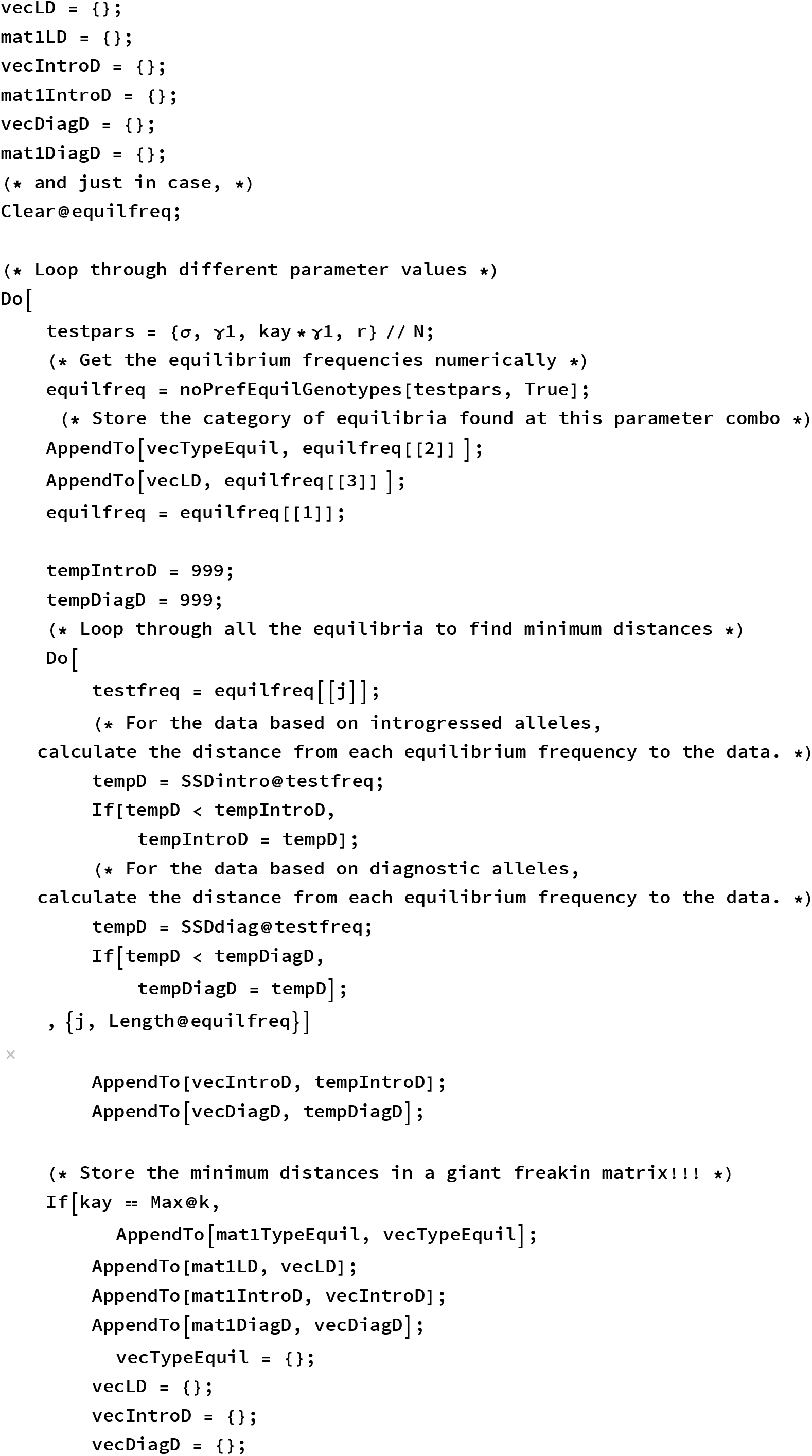
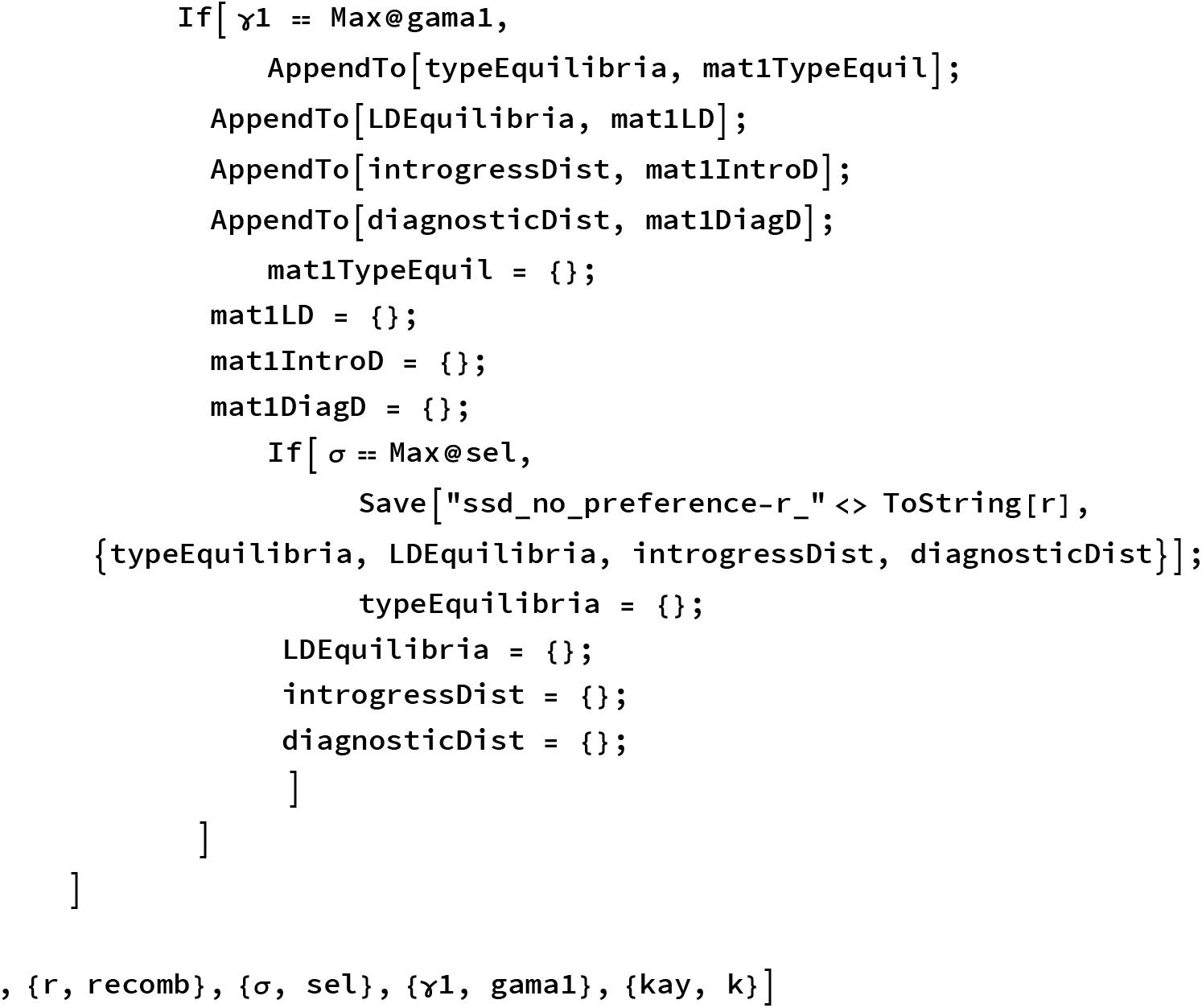

Finally pick the top fitting models using the following (rather arbitrary) method:

~~~
(* read in the data from multiple files *)
Get[ToString[NotebookDirectory[]] <> “ssd_no_preference-params”];
~~~

~~~
typeEquilibriaTEMP = {};
LDEquilibriaTEMP = {};
introgressDistTEMP = {};
diagnosticDistTEMP = {};
Do [
 Get[ToString[NotebookDirectory[]] <> “ssd_no_preference-r_” <> ToString[r]];
 AppendTo[typeEquilibriaTEMP, typeEquilibria];
 AppendTo[LDEquilibriaTEMP, LDEquilibria];
 AppendTo[introgressDistTEMP, introgressDist];
 AppendTo[diagnosticDistTEMP, diagnosticDist];
, {r, recomb}]
~~~

~~~
typeEquilibria = typeEquilibriaTEMP;
LDEquilibria = LDEquilibriaTEMP;
introgressDist = introgressDistTEMP;
diagnosticDist = diagnosticDistTEMP;
Remove[typeEquilibriaTEMP, LDEquilibriaTEMP,
 introgressDistTEMP, diagnosticDistTEMP ]
~~~

~~~
(* visualize the ranked model fits. The steps you
 see on the plot correspond to scenarios like exclusion and
 symmetric coexistence which have exactly equal (and bad) SSD *)
ListPlot[Sort@Flatten[introgressDist],
 FrameLabel →{“Ranked Model Fits”, “Sum of Squares Distance to Data”}, Frame → True]
Length@Select[Flatten[introgressDist], # < 0.095 &]
Length @ Select [Flatten [introgressDist], # < 0.094 &]
(* I’m totally eyeballing this,
so take what look like the top values. These correspond
 to the best values as per my entirely arbitrary cut-off:D *)
topIntro = DeleteDuplicates@Flatten[
  Position[introgressDist, #] & /@ Sort[Flatten[introgressDist]][[1;; 124]], 1];
(* need to delete duplicates because position gets ALL indices
 that have exactly the same SSD *)
~~~

~~~
(* Throw out the values that converge to CF because I found out below that the
 single locus polymorphism is that one that is being preferred and we already know
 that for this tristable param combo the SLP can only be reached under secondary
 contact when p and q are very very small/large.We think that it’s more likely
 for the secondary contact to have been closer to 50-50 for some reason … *)
topIntro = Delete[topIntro, Position[Map[EquilType, topIntro], “CF”]];
outcomesIntro = Map[EquilType, topIntro]
Length@topIntro
~~~

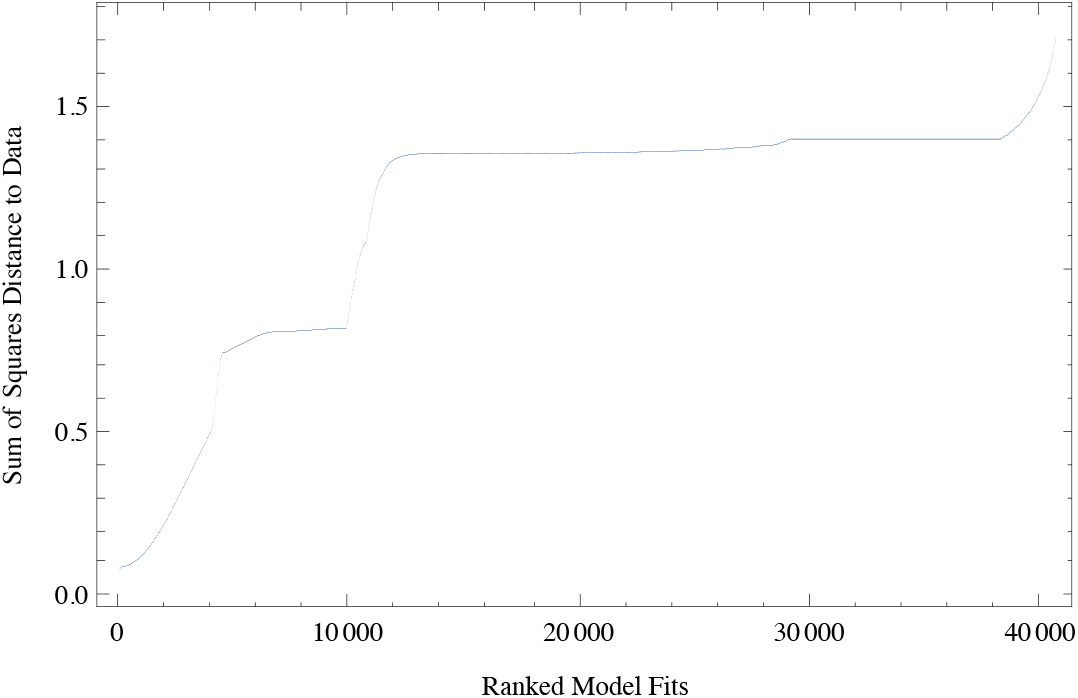

218

124

~~~
{A, A, C, A, C, C, C, A, A, A, A, A, A, A, A, A, A, A, A, C, C, C, C, C, C, C, C, C, C, C, C, C,
C, C, C, C, C, C, C, A, A, A, C, A, A, A, C, C, C, A, C, C, C, C, C, A, A, A, A, A, C, C,
C, C, C, C, A, C, C, C, C, C, C, C, C, C, C, C, C, C, C, C, C, C, C, C, C, C, C, C, C, A,
A, A, A, A, A, C, C, C, C, C, C, A, C, C, C, C, C, C, C, C, C, C, C, C, C, C, C, C, C, C}
~~~

122

Repeat the same method as above for the diagnostic data (you need to load the data by running the cells above first).

~~~
ListPlot[Sort@Flatten[diagnosticDist],
 FrameLabel →”Ranked Model Fits”, “Sum of Squares Distance to Data”}, Frame → True]
Length@Select[Flatten[diagnosticDist], # < 0.24 &]
Length@Select[Flatten[diagnosticDist], # < 0.25 &]
topDiag = DeleteDuplicates@Flatten[
  Position[diagnosticDist, #] & /@ Sort[Flatten[diagnosticDist] ][[ 1;; 167]], 1];
~~~

~~~
(* Throw out the values that converge to CF because I found out below that
 the single locus polymorphism is that one that is being preferred and we
 already know that for this tristable param combo the SLP can only be reached
 under secondary contact when p and q are very very small/large.We think
 that it’s more likely for p and q to have some intermediate value. *)
 topDiag = Delete[topDiag, Position[Map[EquilType, topDiag], “CF”]];
outcomesDiag = Map[EquilType, topDiag]
Length@topDiag
~~~

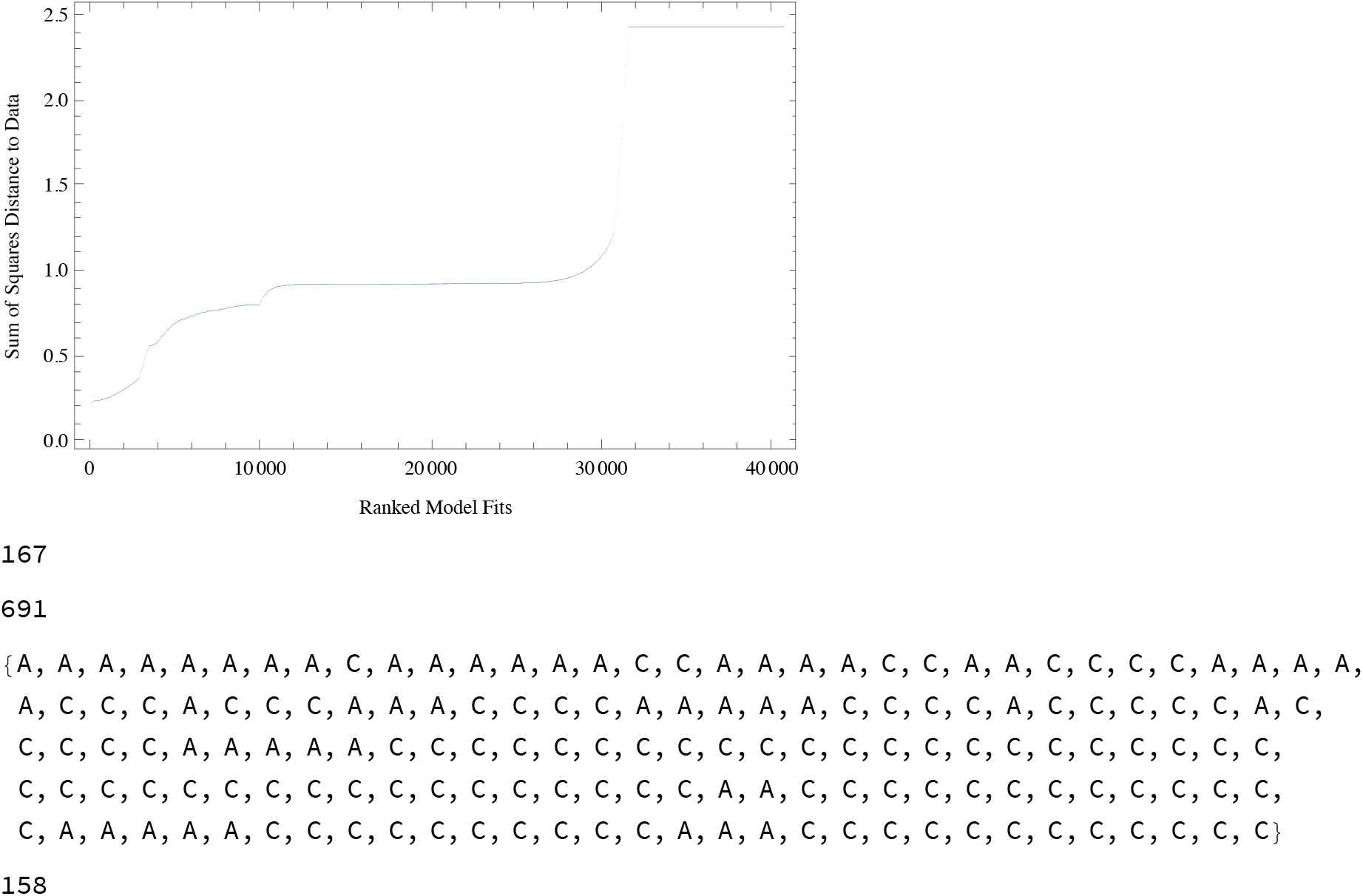

All of the analysis is straightforward from here…

However, maybe it’s helpful if I provide an example of how I got the model predictions. I’m showing the method just for the data with diagnostic alleles but the same thing was done exactly for introgressed alleles.

~~~
(* classify the outcomes *)
Tally@outcomesDiag

(* all top models have 2 equilibria. Figure
 out the index of the preferred equilibrium *)
cfEqIndex = {};
bestFitFreq = {};
Do [
 equilfreq = noPrefEquilGenotypes[GetParamVals@topDiag[[i]] ];
 indexPref = Position[SSDdiag[#] & /@ equilfreq, Min[SSDdiag[#] & /@ equilfreq]];
 AppendTo[cfEqIndex, indexPref];
 AppendTo[bestFitFreq, Flatten[equilfreq[[Flatten@indexPref]], 1] ]
, {i, Length@topDiag}]
Flatten@cfEqIndex

(* Given the indices above,it is ALWAYS the 1st equilibrium that is being preferred.
 Plot the predicted values: *)
DistributionChart[Transpose@bestFitFreq[[All, 1] ],
 ChartLabels -> {“00”, “most0”, “F1”, “01”, “10”, “most1”, “11”,
 “00”, “01”, “10”, “11”}, PlotLabel → “before selection”, ChartElementFunction → “HistogramDensity”]
~~~

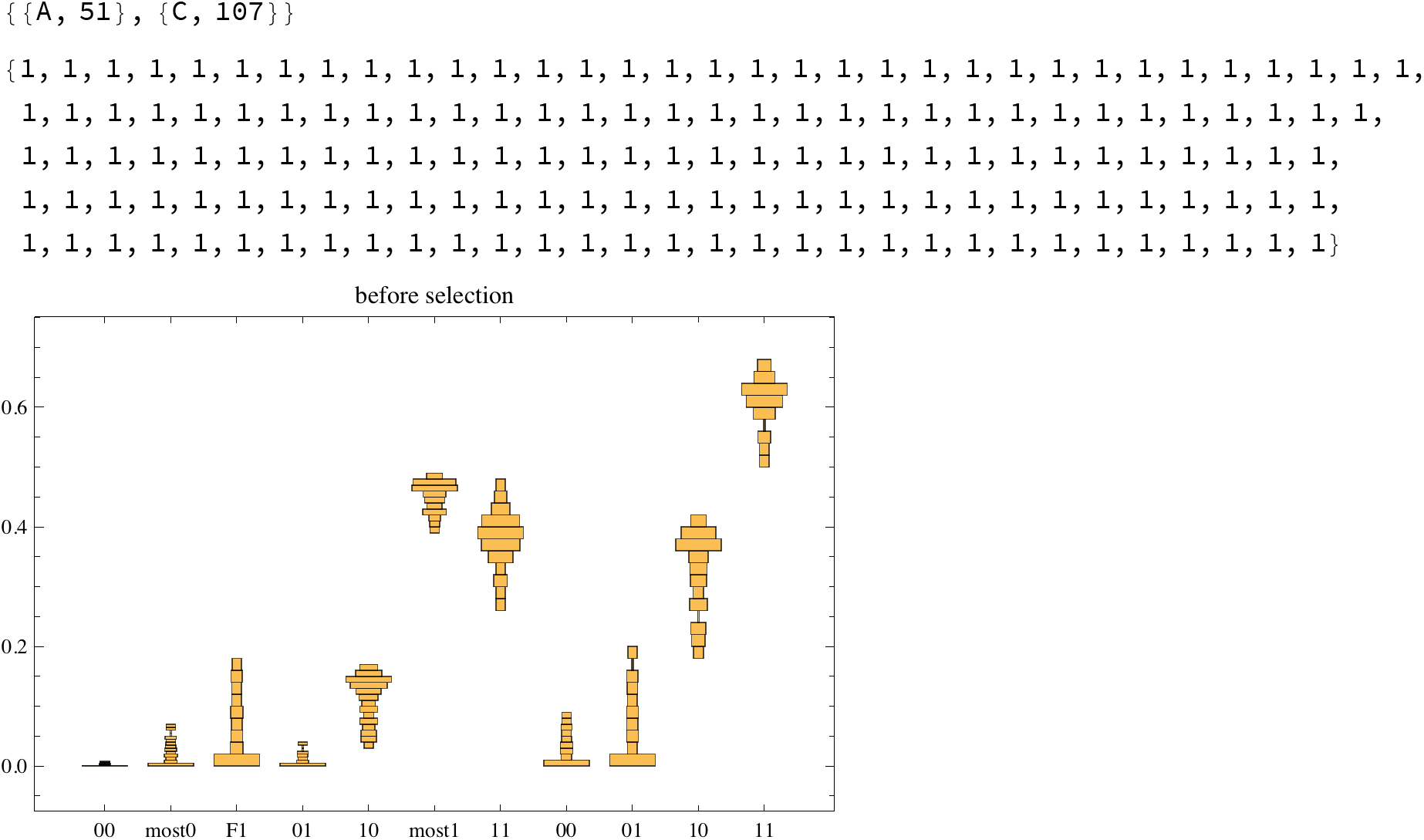

#### Data fitting to the model with preference

The data fitting to the model with preference was done by analogous methodology as above. See below.

Please note that these simulations took about *TWO WEEKS* to run…

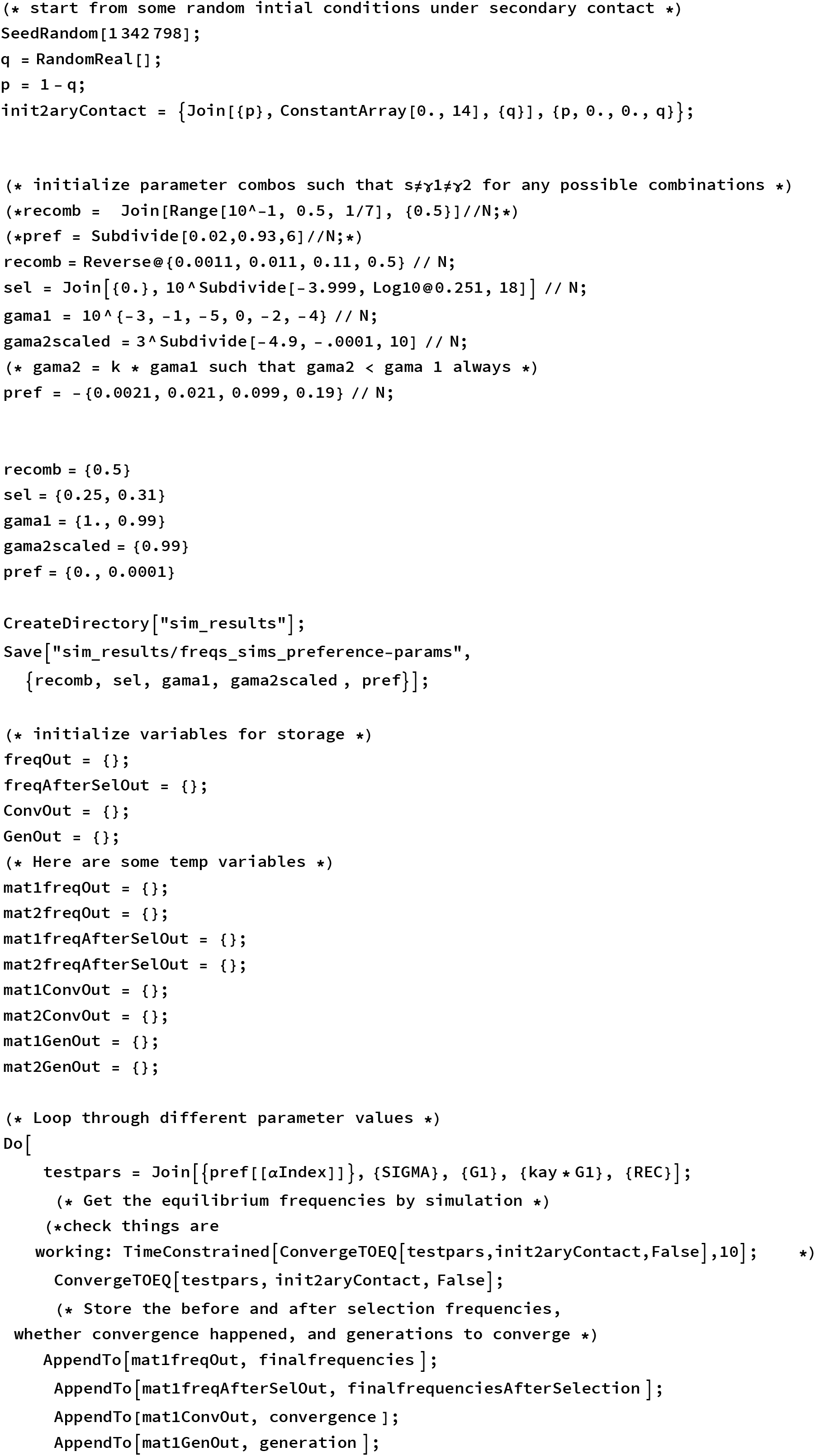
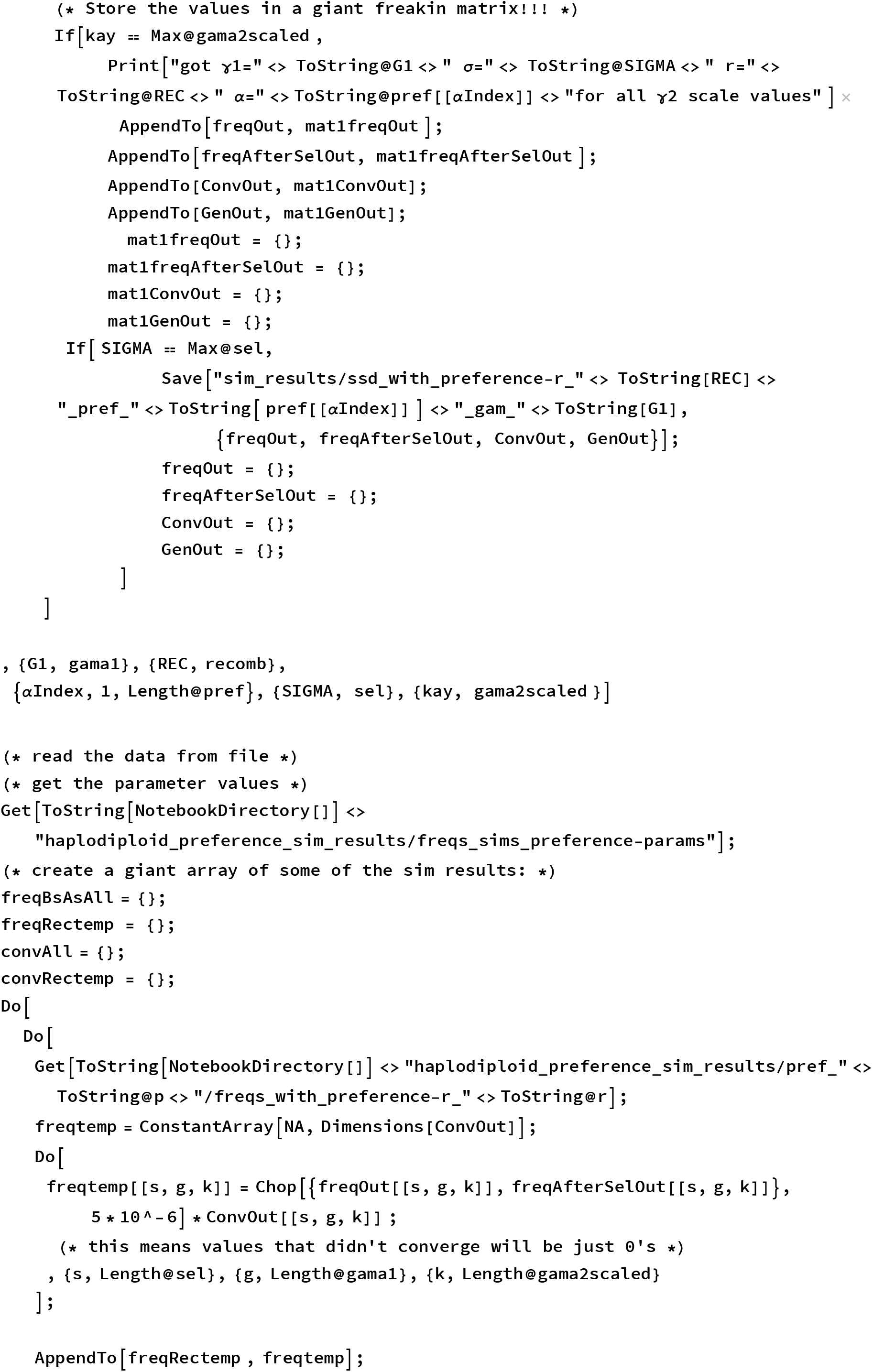
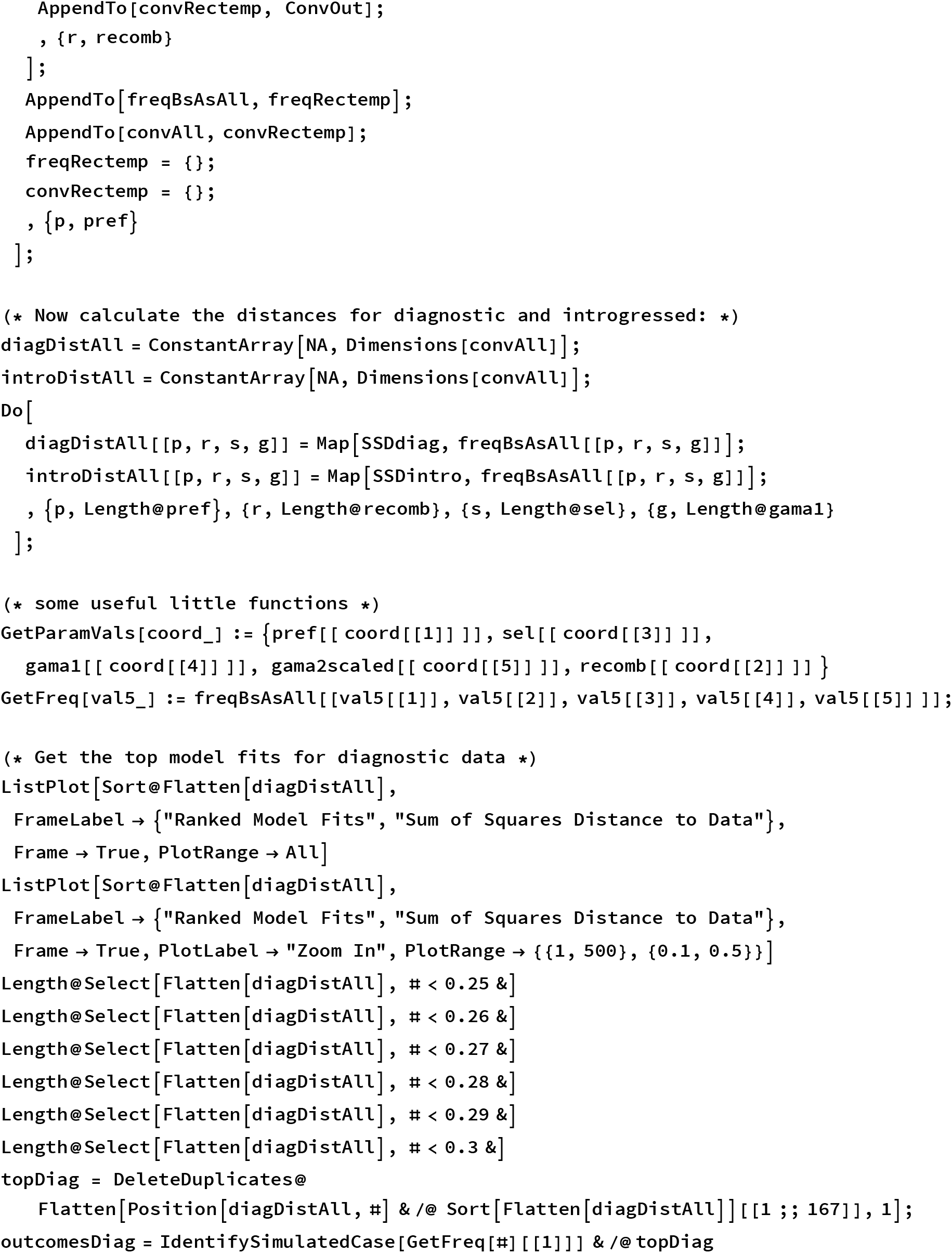
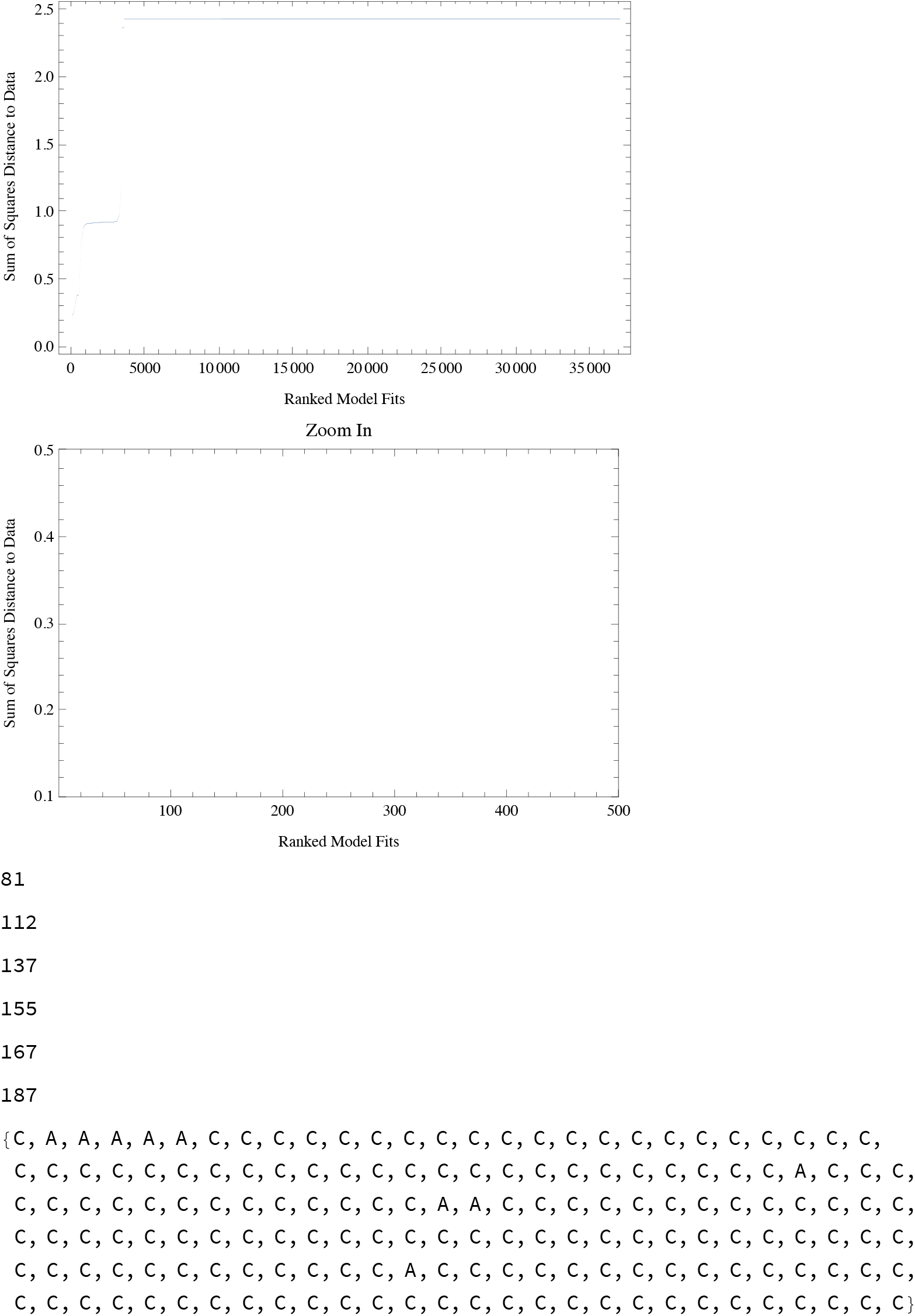

Repeat the same method as above for the diagnostic data (you need to load the data by running the cells above first).

~~~
ListPlot[Sort@Flatten[introDistAll],
 FrameLabel → {“Ranked Model Fits”, “Sum of Squares Distance to Data”},
 Frame → True, PlotRange → All]
ListPlot[Sort@Flatten[introDistAll],
 FrameLabel → {“Ranked Model Fits”, “Sum of Squares Distance to Data”},
 Frame → True, PlotLabel → “Zoom In”, PlotRange → {{1, 300}, {0., 0.3}}]
Length@Select[Flatten[introDistAll], # < 0.14 &]
Length@Select[Flatten[introDistAll], # < 0.15 &]
Length@Select[Flatten[introDistAll], # < 0.16 &]
Length@Select[Flatten[introDistAll], # < 0.17 &]
topIntro = DeleteDuplicates@
 Flatten[Position[introDistAll, #] & /@ Sort[Flatten[introDistAll] ][[1;; 115]], 1];
IdentifySimulatedCase[GetFreq[#][[1]]] & /@ topIntro
(* get rid of the weird one *) topIntro = Delete[topIntro,
 Position[IdentifySimulatedCase[GetFreq[#][[1]]] & /@ topIntro, “weird”]];
outcomesIntro = IdentifySimulatedCase[GetFreq[#][[1]]] & /@ topIntro
~~~

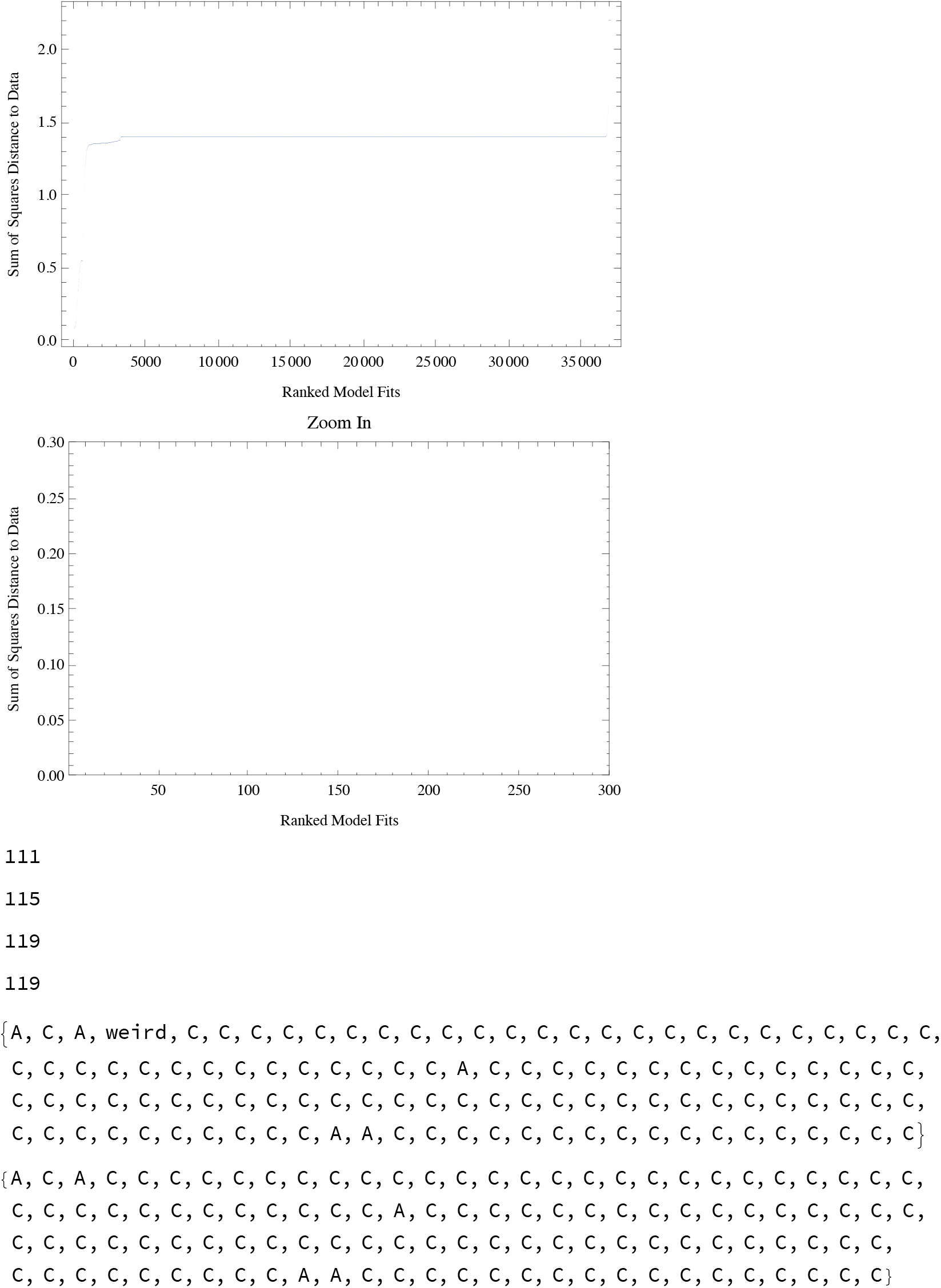

## Bibliography

Abbott, J., Nordén, A., and Hansson, B. (2017). Sex chromosome evolution: Historical insights and future perspectives. Proceedings of the Royal Society of London B: Biological Sciences, 284(1854):20162806.

Abbott, R., Albach, D., Ansell, S., Arntzen, J. W., Baird, S. J. E., Bierne, N., Boughman, J., Brelsford, A., Buerkle, C. A., Buggs, R., Butlin, R. K., Dieckmann, U., Eroukhmanoff, F., Grill, A., Cahan, S. H., Hermansen, J. S., Hewitt, G., Hudson, A. G., Jiggins, C., Jones, J., Keller, B., Marczewski, T., Mallet, J., Martinez-Rodriguez, P., Möst, M., Mullen, S., Nichols, R., Nolte, A. W., Parisod, C., Pfennig, K., Rice, A. M., Ritchie, M. G., Seifert, B., Smadja, C. M., Stelkens, R., Szymura, J. M., Väinölä, R., Wolf, J. B. W., and Zinner, D. (2013). Hybridization and speciation. Journal of Evolutionary Biology, 26(2):229–246.

Albert, A. and Otto, S. (2005). Sexual selection can resolve sex-linked sexual antagonism. Science, 310(5745):119–121.

Arnqvist, G. and Rowe, L. (2005). Sexual conflict.

Bateson, W. (1909). Heredity and variation in modern lights. Darwin and modern science, pages 85–101.

Beresford, J., Elias, M., Pluckrose, L., Sundström, L., Butlin, R., Pamilo, P., and Kulmuni, J. (2017). Widespread hybridization within mound-building wood ants in Southern Finland results in cytonuclear mismatches and potential for sex-specific hybrid breakdown. Molecular Ecology, 26(15):4013–26.

Bernardes, J., Stelkens, R., and Greig, D. (2017). Heterosis in hybrids within and between yeast species. Journal of Evolutionary Biology, 30(3):538–548.

Buerkle, C. A., Morris, R. J., Asmussen, M. A., and Rieseberg, L. H. (2000). The likelihood of homoploid hybrid speciation. Heredity, 84(4):441–451.

Bürger, R. (2000). The mathematical theory of selection, recombination, and mutation, volume 228. Wiley Chichester.

Butlin, R. K. and Ritchie, M. G. (2013). Pulling together or pulling apart: hybridization in theory and practice. Journal of Evolutionary Biology, 26(2):294–298.

Charlesworth, B., Coyne, J., and Barton, N. (1987). The relative rates of evolution of sex chromosomes and autosomes. The American Naturalist, 130(1):113–146.

Chen, C., Zhiguo, E., and Lin, H.-X. (2016). Evolution and molecular control of hybrid incompatibility in plants. Frontiers in Plant Science, 7:1208.

Chen, Z. (2013). Genomic and epigenetic insights into the molecular bases of heterosis. Nature Review Genetics, 14(7):471–482.

Corbett-Detig, R. B., Zhou, J., Clark, A. G., Hartl, D. L., and Ayroles, J. F. (2013). Genetic incompatibilities are widespread within species. Nature, 504(7478):135–7.

Crozier, R. and Pamilo, P. (1996). Evolution of social insect colonies: Sex allocation and kin selection. Oxford University Press, Oxford, UK.

De Cara, M., Barton, N., and Kirkpatrick, M. (2008). A model for the evolution of assortative mating. The American Naturalist, 171(5):580–596.

de la Filia, A., Bain, S., and Ross, L. (2015). Haplodiploidy and the reproductive ecology of arthropods. Current Opinion in Insect Science, 9:36–43.

Dieckmann, U. and Doebeli, M. (1999). On the origin of species by sympatric speciation. Nature, 400(6742):354–357.

Dobzhansky, T. (1936). Studies on hybrid sterility. II. Localization of sterility factors in *Drosophila pseudoobscura* hybrids. Genetics, 21(2):113.

Evans, J., Shearman, D., and Oldroyd, B. (2004). Molecular basis of sex determination in haplodiploids. Trends in Ecology and Evolution, 19(1):1–3.

Fraïsse, C., Elderfield, J., and Welch, J. (2014). The genetics of speciation: are complex incompatibilities easier to evolve? Journal of Evolutionary Biology, 27(4):688–99.

Gibson, J., Chippindale, A., and Rice, W. (2002). The X chromosome is a hot spot for sexually antagonistic fitness variation. Proceedings of the Royal Society of London B: Biological Sciences, 269(1490):499–505.

Hedrick, P. W. (2012). What is the evidence for heterozygote advantage selection? Trends in Ecology & Evolution, 27(12):698–704.

Heliconius Genome Consortium (2012). Butterfly genome reveals promiscuous exchange of mimicry adaptations among species. Nature, 487(7405):94.

Höllinger, I. and Hermisson, J. (2017). Bounds to parapatric speciation: A Dobzhan-sky-Muller incompatibility model involving autosomes, X chromosomes, and mitochondria. Evolution, 71(5):1366–1380.

Jablonka, E. and Lamb, M. J. (1991). Sex Chromosomes and Speciation. Proceedings of the Royal Society B: Biological Sciences, 243(1308):203–208.

Johnson, N. A. and Lachance, J. (2012). The genetics of sex chromosomes: evolution and implications for hybrid incompatibility. Annals of the New York Academy of Sciences, 1256(1):E1–E22.

Knegt, B., Potter, T., Pearson, N., Sato, Y., Staudacher, H., Schimmel, B., Kiers, E., and Egas, M. (2017). Detection of genetic incompatibilities in non-model systems using simple genetic markers: hybrid breakdown in the haplodiploid spider mite tetranychus evansi. Heredity, 118(4):311.

Koevoets, T. and Beukeboom, L. (2009). Genetics of postzygotic isolation and Haldane’s rule in haplodiploids. Heredity, 102(1):16–23.

Kopp, M., Servedio, M. R., Mendelson, T. C., Safran, R. J., Rodríguez, R. L., Hauber, M. E., Scordato, E. C., Symes, L. B., Balakrishnan, C. N., Zonana, D. M., et al. (2017). Mechanisms of assortative mating in speciation with gene flow: Connecting theory and empirical research. The American Naturalist, 191(1).

Kraaijeveld, K. (2009). Male genes with nowhere to hide; sexual conflict in haplodiploids. Animal Biology, 59(4):403–415.

Kulmuni, J. and Pamilo, P. (2014). Introgression in hybrid ants is favored in females but selected against in males. Proceedings of the National Academy of Sciences, 111(35):12805–10.

Kulmuni, J., Seifert, B., and Pamilo, P. (2010). Segregation distortion causes large-scale differences between male and female genomes in hybrid ants. Proceedings of the National Academy of Sciences, 107(16):7371–6.

Kulmuni, J. and Westram, A. M. (2017). Intrinsic incompatibilities evolving as a by-product of divergent ecological selection: Considering them in empirical studies on divergence with gene flow. Molecular Ecology, 26(12):3093–3103.

Lewontin, R. (1964). The interaction of selection and linkage. i. general considerations; heterotic models. Genetics, 49-67(1):49.

Li, M.-H. and Merilä, J. (2010). Sex-specific population structure, natural selection, and linkage disequilibrium in a wild bird population as revealed by genome-wide microsatellite analyses. BMC evolutionary biology, 10(1):66.

Lima, T. G. (2014). Higher levels of sex chromosome heteromorphism are associated with markedly stronger reproductive isolation. Nature Communications, 5:4743.

Lohse, K. and Ross, L. (2015). What haplodiploids can teach us about hybridization and speciation. Molecular Ecology, 24(20):5075–5077.

Mallet, J. (2005). Hybridization as an invasion of the genome. Trends in Ecology and Evoluion, 20(5):229–237.

Montecinos, A. E., Guillemin, M.-L., Couceiro, L., Peters, A. F., Stoeckel, S., and Valero, M. (2017). Hybridization between two cryptic filamentous brown seaweeds along the shore: analysing pre- and postzygotic barriers in populations of individuals with varying ploidy levels. Molecular Ecology, 26(13):3497–3512.

Muller, H. (1942). Isolating mechanisms, evolution and temperature. In Biology Symposium, volume 6, pages 71–125.

Nagylaki, T. et al. (1992). Introduction to theoretical population genetics, volume 142. Springer-Verlag Berlin.

Orr, H. (1995). The population genetics of speciation: The evolution of hybrid incompatibilities. Genetics, 139(4):1805–1813.

Otto, S. P. and Day, T. (2007). A biologist’s guide to mathematical modeling in ecology and evolution, volume 13. Princeton University Press.

Paixão, T., Bassler, K. E., and Azevedo, R. B. R. (2014). Emergent speciation by multiple Dobzhansky-Muller incompatibilities. bioRxiv, page 8268.

Pamilo, P. (1979). Genic variation at sex-linked loci: Quantification of regular selection models. Hereditas, 91(1):129–133.

Pamilo, P. and Crozier, R. H. (1981). Genic variation in male haploids under deterministic selection. Genetics, 98(1):199–214.

Patten, M., Carioscia, S., and Linnen, C. (2015). Biased introgression of mitochondrial and nuclear genes: A comparison of diploid and haplodiploid systems. Molecular Ecology, 24(20):5200–5210.

Pischedda, A. and Chippindale, A. K. (2006). Intralocus sexual conflict diminishes the benefits of sexual selection. PLoS biology, 4(11):e356.

Presgraves, D. (2008). Sex chromosomes and speciation in *Drosophila*. Trends in Genetics, 24(7):336–343.

Runemark, A., Trier, C. N., Eroukhmanoff, F., Hermansen, J. S., Matschiner, M., Ravinet, M., Elgvin, T. O., and Saetre, G.-P. (2017). Variation and constraints in hybrid genome formation. bioRxiv, page 107508.

Sandor, C., Farnir, F., Hansoul, S., Coppieters, W., Meuwissen, T., and Georges, M. (2006). Linkage disequilibrium on the bovine x chromosome: characterization and use in quantitative trait locus mapping. Genetics, 173(3):1777–1786.

Schluter, D. (2009). Evidence for ecological speciation and its alternative. Science, 323(5915):737–741.

Schluter, D. and Conte, G. (2009). Genetics and ecological speciation. Proceedings of the National Academy of Sciences, 106(Supplement 1):9955–9962.

Schumer, M., Cui, R., Rosenthal, G. G., and Andolfatto, P. (2015). Reproductive isolation of hybrid populations driven by genetic incompatibilities. PLOS Genetics, 11(3):1–21.

Schumer, M., Rosenthal, G., and Andolfatto, P. (2014). How common is homoploid hybrid speciation? Evolution, 68(6V1553–1560

Schwarz, D., Matta, B., Shakir-Botteri, N., and McPheron, B. (2005). Host shift to an invasive plant triggers rapid animal hybrid speciation. Nature, 436(7050):546–9.

Seehausen, O., Butlin, R., Keller, I., Wagner, C., Boughman, J., Hohenlohe, P., Peichel, C., Saetre, G.-P., Bank, C., Brännström, A., Brelsford, A., Clarkson, C., Eroukhmanoff, F., Feder, J., Fischer, M., Foote, A., Franchini, P., et al. (2014). Genomics and the origin of species. Nature Reviews Genetics, 15(3):176–192.

Servedio, M. and Noor, M. (2003). The role of reinforcement in speciation: theory and data. Annual Review of Ecology, Evolution, and Systematics, 34(1):339–364.

Song, Y., Endepols, S., Klemann, N., Richter, D., Matuschka, F.-R., Shih, C.-H., Nachman, M., and Kohn, M. (2011). Adaptive introgression of anticoagulant rodent poison resistance by hybridization between old world mice. Current Biology, 21(15):1296–1301.

Suomalainen, E., Saura, A., and Lokki, J. (1987). Cytology and evolution in parthenogenesis. CRC Press, Boca Raton, Florida.

Wall, J. D., Andolfatto, P., and Przeworski, M. (2002). Testing models of selection and demography in *Drosophila simulans*. Genetics, 162(1):203–216.

Whitney, K., Randell, R., and Rieseberg, L. (2010). Adaptive introgression of abiotic tolerance traits in the sunflower *Helianthus annuus*. New Phytologist, 187(1):230–239.

Whitney, K. D., Broman, K. W., Kane, N. C., Hovick, S. M., Randell, R. A., and Rieseberg, L. H. (2015). Quantitative trait locus mapping identifies candidate alleles involved in adaptive introgression and range expansion in a wild sunflower. Molecular Ecology, 24(9):2194–2211.

Wolf, D., Takebayashi, N., and Rieseberg, L. (2001). Predicting the risk of extinction through hybridization. Conservation Biology, 15(4):1039–1053.

Wolfram Research, Inc. (2016). Mathematica v. 10.4.1.0. Champaign, Illinois, USA. https://www.wolfram.com.

## References

J Kulmuni and P Pamilo. Introgression in hybrid ants is favored in females but selected against in males. Proceedings of the National Academy of Sciences, 111(35):12805–10, 2014. doi: 10.1073/pnas.1323045111.

J Kulmuni, B Seifert, and P Pamilo. Segregation distortion causes large-scale differences between male and female genomes in hybrid ants. Proceedings of the National Academy of Sciences, 107(16):7371–6, 2010. doi: 10.1073/pnas.0912409107.

